# A genomic timescale for placental mammal evolution

**DOI:** 10.1101/2022.08.10.503388

**Authors:** Nicole M. Foley, Victor C. Mason, Andrew J. Harris, Kevin R. Bredemeyer, Joana Damas, Harris A. Lewin, Eduardo Eizirik, John Gatesy, Zoonomia Consortium, Mark S. Springer, William J. Murphy

**Author notes:** These authors contributed equally to this work (Victor C. Mason, Andrew J. Harris, Kevin R. Bredemeyer, Joana Damas).

## Abstract

The precise pattern and timing of speciation events that gave rise to all living placental mammals remain controversial. We provide a comprehensive phylogenetic analysis of genetic variation across an alignment of 241 placental mammal genome assemblies, addressing prior concerns regarding limited genomic sampling across species. We compared neutral genome-wide phylogenomic signal using concatenation and coalescent-based approaches, interrogated phylogenetic variation across chromosomes and analyzed extensive catalogs of structural variants. Interordinal relationships exhibit relatively low rates of phylogenomic conflict across diverse datasets and analytical methods. Conversely, X-chromosome versus autosome conflicts characterize multiple independent clades that radiated during the Cenozoic. Genomic timetrees reveal an accumulation of cladogenic events before and immediately following the KPg boundary implying important roles for Cretaceous continental vicariance and the KPg extinction in the placental radiation.

**One-Sentence Summary:** A comprehensive whole genome phylogeny of extant placental mammals reveals timing and patterns of ordinal diversification.

## Main Text

Placental mammals display a staggering breadth of morphological, karyotypic, and genomic diversity, rivaling or surpassing any other living vertebrate clade (*1–3*). This variation represents the culmination of one hundred million years of diversification and parallel adaptation to tumultuous changes in Earth’s environments, including catastrophic events like the Cretaceous-Paleogene (KPg) bolide impact. These different measures of diversity have impeded a complete reckoning of how and why modern placental mammal orders suddenly appeared in the Paleocene with scant paleontological signal preceding the KPg impact.

Prior studies have produced conflicting results regarding the timing and sequence of interordinal and intraordinal cladogenesis. As many as five models of placental mammal diversification have been proposed (*4, 5*), each implying different degrees of causality between the KPg extinction event and ordinal diversification. Each model is supported by molecular analyses of different sequence matrices that have been heavily biased toward short, evolutionarily-constrained protein-coding exons or ultraconserved non-coding sequences (*6–10*). Biased genomic sampling has hampered a full resolution of the placental mammal phylogeny and an understanding of the principal drivers of ordinal diversification.

Here we report a comprehensive analysis of phylogenomic signal from investigations of multiple genomic character types assayed from a hierarchical alignment (HAL) of 241 placental mammal whole-genome assemblies (*1, 11*). The HAL samples all placental mammal orders and represents 62% of placental families. The process and data structure that generated the HAL provide a statistically vetted whole genome assessment of synteny and sequence orthology, reducing the potential for phylogenetic reconstruction errors caused by ortholog misidentification observed in some previous studies (*12*). The resulting availability of per base estimates of genomic constraint (PhyloP scores) also allowed us to assess the impacts of natural selection on phylogenetic signal and enabled the rigorous application of coalescent approaches (*13*).

## Results

### Whole-Genome Phylogenies

We applied site pattern frequency-based coalescent methods implemented in the SVDquartets program to sample SNPs spaced by a minimum of 1kb to reduce the impacts of intralocus recombination and linkage. We estimated phylogenetic relationships for all species in the HAL alignment and for 65 taxon matrices that sample all ordinal lineages while minimizing missing data (table S1). Three versions of the 65-taxon alignment were analyzed to mitigate the reference-bias of MAF alignments that were extracted from the HAL (table S2): a human-referenced alignment (HRA), a dog-referenced alignment (DRA), and a root-referenced alignment (RRA) that was imputed from the inferred placental ancestor (*1*). Because of the absence of non-placental outgroups in our alignment the root position was assumed to be between Atlantogenata and Boreoeutheria (*5*) and remains an open question. To investigate the impact of selection, we also identified conserved, accelerated, and nearly-neutral evolving SNPs from a distribution of HRA sites ranked by PhyloP conservation scores across the 241-species alignment (*14*).

HRA coalescent trees estimated for 65 and 241 species from nearly-neutral PhyloP sites were highly resolved, with 96% and 97% of the quartets compatible with the inferred species trees, respectively (Fig. 1A, fig. S1A, and table S2). The 65-taxon accelerated sites tree was topologically identical to the nearly neutral tree (fig. S1B). The 65-taxon tree computed based on conserved sites (fig. S1C) differed only in the positions of Macroscelidea and Scandentia. The dog-referenced 65-taxon tree (fig S2A) was also identical to the nearly-neutral HRA topology, except for relationships within Afroinsectiphilia. The root-referenced tree (fig. S2B) differed from the human and dog referenced trees only by supporting an elephant+sirenian clade (Tethytheria) within Paenungulata (fig. S2). The HRA results were robust to different measures of missing data (fig. S3).

**Fig. 1.**
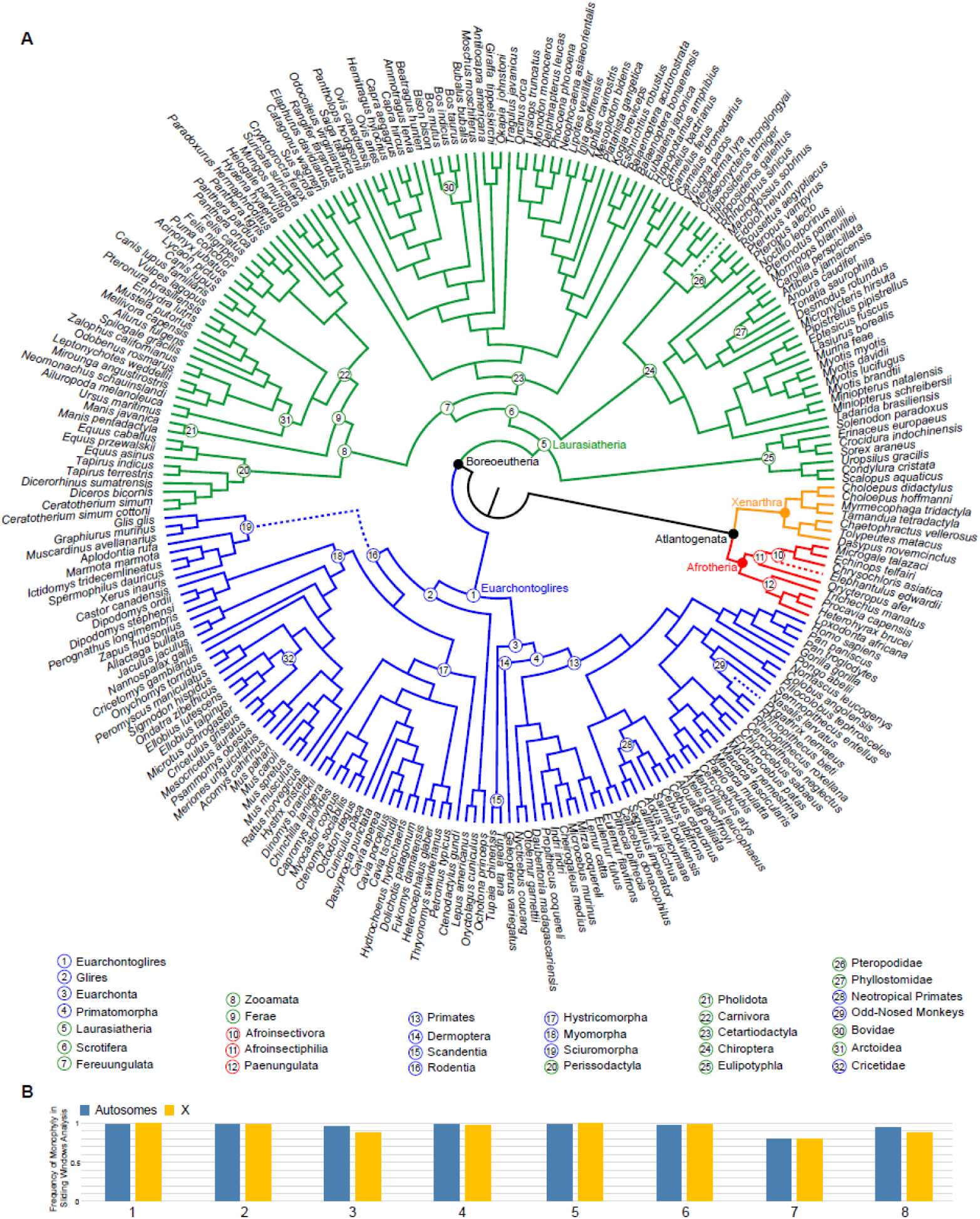
Placental mammal phylogeny based on coalescent analysis of nearly-neutral sites. (**A**) 50% Majority-rule consensus tree from a SVDquartets analysis of 411,110 genome-wide, nearly-neutral sites from the human referenced alignment of 241 species. Bootstrap support is 100% for all nodes. Superordinal clades are labelled and identified in four colors. Nodes corresponding to Boreoeutheria and Atlantogenata are indicated by black circles. (**B**) The frequency at which eight superordinal clades (numbered 1-8 in Fig. 1A) were recovered as monophyletic in 2,164 window-based maximum likelihood trees from representative autosomes (Chr1, Chr21 and Chr22) and ChrX. Dotted lines indicate relationships that differ from the concatenated Maximum Likelihood analysis.

The superordinal clades Euarchonta (primates, colugos, and treeshrews), Glires (rodents and lagomorphs), Scrotifera (bats, cetartiodactyls, perissodactyls, carnivorans, and pangolins), Fereuungulata (all scrotiferans excluding bats), and Zoomata (Ferae [carnivorans and pangolins] + Perissodactyla) were well supported in all analyses (Fig. 1), including those using sites at different extremes of selective constraint and missingness (i.e., the percentage of missing data per alignment column) (figs. S1 and S3). Concatenated analyses of the same SNP datasets generally were highly congruent with coalescent-based superordinal relationships (Fig. 1A and table S3), but within Afrotheria relationships among afroinsectiphilians were less well-resolved in a subset of the coalescent and concatenation analyses. More limited taxon sampling in this clade, higher percentages of missing data for some afrotherians, sequence alignment uncertainty, and/or long branches may contribute to the discordance observed for afroinsectiphilian relationships among different analyses (Table S1). Future high-quality genomic sampling of afrotherian biodiversity should be a priority.

### Genomic Distribution of Phylogenomic Signal

Coalescent-based approaches such as SVDquartets assume incomplete lineage sorting (ILS) but no interspecific gene flow. Concatenation methods assume that the most common phylogenetic signal represents the species tree. Both approaches typically mask signatures of ancestral hybridization or admixture (*15–17*). To address this problem, we generated 2,164 maximum likelihood trees for 228 species from 100-kilobase (kb) alignment windows (=locus trees) sampled across three human autosomes (Chr1, Chr21, and Chr22) and the X chromosome (ChrX) (table S4). These locus trees sample more than 95-Mb of predominantly (98%) non-coding alignment columns from chromosomes that sample a broad range of karyotypic attributes, including size, gene density, inferred historical recombination rate (Table 1), and ancestral gene order (*18–21*). The genomic segments corresponding to human Chr21 and Chr22 are frequently found near telomeres and on small chromosomes in the majority of placental mammal karyotypes (table S5) (*3, 21*), which is predictive of historically high meiotic recombination rate and gene tree conflict (*15, 16*). Conversely, the highly collinear X chromosome in mammals contains a large, conserved recombination coldspot and is expected to be enriched in signal that is consistent with the species trees across diverse clades (*16, 19*). Although resolved recombination maps are lacking for most placental mammal species, the correlation between biased GC conversion and meiotic recombination allows the local recombination rate to be approximated via estimates of GC content (*22*). We used TreeHouseExplorer (*23*) to visualize locus trees across autosomes and the X chromosome and regions of high and low GC content to identify chromosome-specific signatures of conflict that would not be apparent in the coalescence or concatenation (majority rule) analyses.

**Table 1.** Karyotypic features of four chromosomes selected for window-based phylogenetic analyses. Gene density refers to the percentage of coding bases for each chromosome in the human hg38 genome assembly.

| Chromosome | Size | Gene Density | Historical Recombination Rate |
| --- | --- | --- | --- |
| **ChrX** | 156,040,895 | 40.1% | Low |
| **Chr1** | 248,956,422 | 59.6% | Low-to-Moderate |
| **Chr21** | 46,709,983 | 48.6% | High |
| **Chr22** | 50,818,468 | 59.5% | High |

Superordinal relationships supported in the coalescent and concatenation trees were also recovered with high frequency in the locus trees distributed across chromosomes (Fig. 1B). Relationships within Laurasiatheria show very low conflict among locus trees, with the Zooamata clade occurring in 95% of autosomal and 89% of ChrX windows and >86% of high and low GC windows (Fig. 2 and table S6). The consistent recovery of the majority of clades among locus trees may be due to the increased number of informative sites. Notably, the high proportion of non-coding positions in our alignments (∼97%, Table 2) provide greater resolving power than coding exons (*24–27*).

**Fig. 2.**
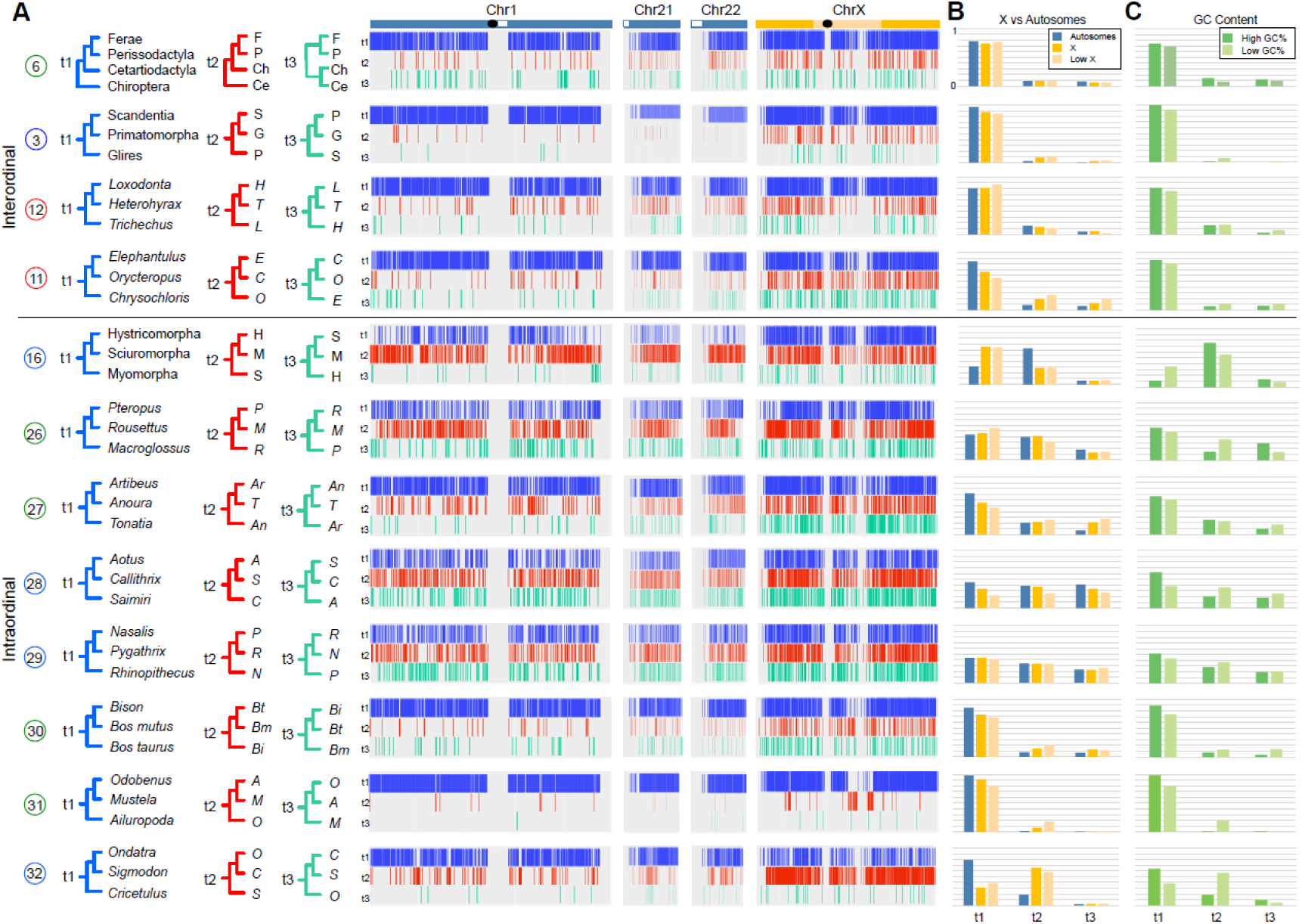
Contrasting patterns of phylogenomic discordance. (**A**) Distribution of phylogenomic signal from select clades (see table S5), visualized using TreeHouseExplorer (*23*) in 100-kb alignment windows along human Chr1, Chr21, Chr22 and ChrX. Vertical bars along each chromosome are color-coded to depict the distribution of the topology (t1 (blue), t2 (red) or t3 (green) – corresponding to topologies shown to the left of the panel) that was recovered in the locus window. Black ovals indicate approximate positions of centromeres and white boxes indicate heterochromatic regions (**B**) Frequency of each topology on the representative autosomes, ChrX and the low-recombining region of the X (*4*). (**C**) Relative topology frequencies in regions of high GC content (>55%) and low GC content (<35%). Note topological differences between ChrX and the autosomes, and corresponding GC content changes, for the primary intraordinal rodent clades, arctoid carnivorans, and cricetid rodents. Support for Zooamata is obtained by summing support for this clade across all three topologies in the top panel of this figure. An alternately colored version of this figure is also available (fig. S8).

**Table 2.**
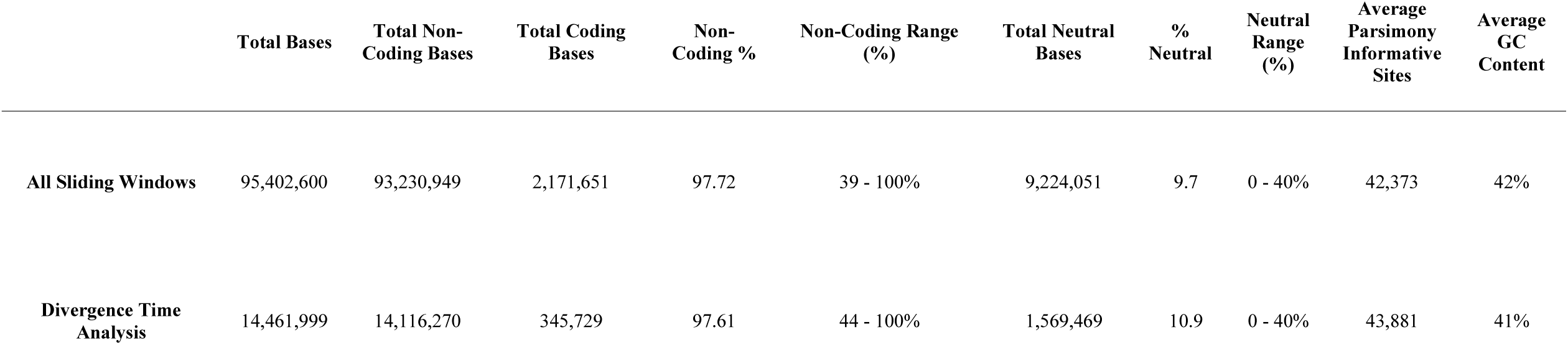
Summary of genomic features of sliding windows datasets used for phylogenomic and divergence time analyses.

### Rare Genomic Changes

We analyzed two independent sets of structural variants that evolve more slowly than nucleotide substitutions to provide an independent character evaluation of tree reconstruction-based results. We searched for deletions >10-bp in size that could potentially support all possible ordinal-level topologies within Laurasiatheria and Euarchontoglires (see Data S1 for ordinal definitions). Deletions provide significant statistical support for all superordinal relationships obtained by the genome-wide and locus tree analyses for Laurasiatheria and Euarchontoglires (Fig. 3 and table S7). The largest numbers of deletions were recovered for Scrotifera, Fereuungulata, and Zooamata (Fig. 3A), which were also supported without conflict by analyses of deletions on ChrX (which possesses the lowest rates of ILS). Euarchonta was the only hypothesis supported by deletions for the position of Scandentia (but see (*21*)).

**Fig. 3.**
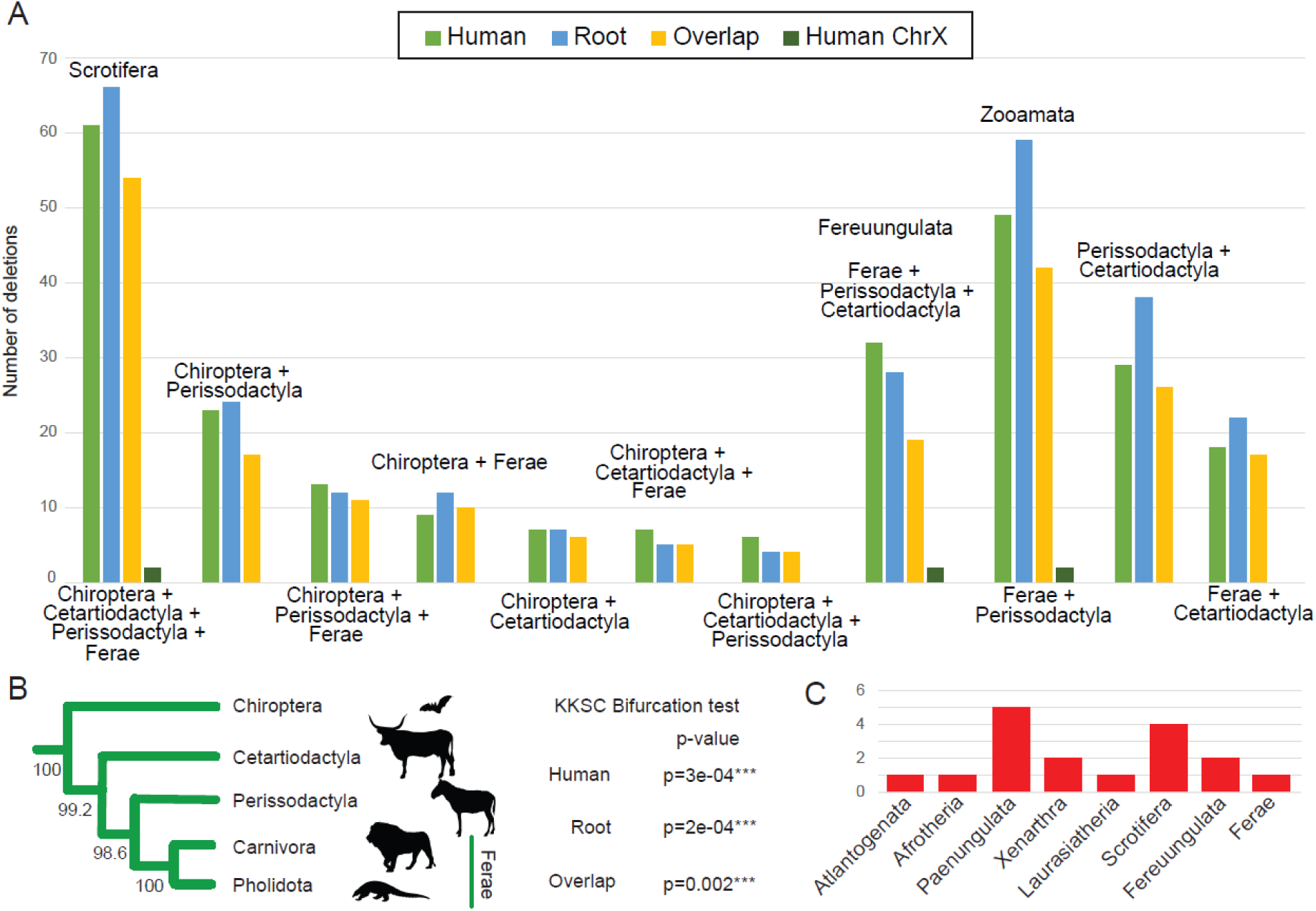
Rare genomic changes. (**A**) Number of deletions recovered in the HRA, RRA, in both the HRA and RRA, and on the HRA ChrX in support of all potential laurasiatherian hypotheses. Within Euarchontoglires, hundreds of raw deletions were recovered for Euarchonta, a subset of which were further validated (table S7). Glires + Primatomorpha and Glires + Scandentia were unsupported by the deletion analysis. (**B**) The topology inferred from the Kuritzin-Kischka-Schmitz-Churakov (KKSC) analysis (*48*) of deletions for Cetartiodactyla, Perissodactyla and Ferae (Carnivora + Pholidota) from the HRA, RRA and HRA/RRA overlap datasets. In all cases, the corresponding KKSC bifurcation test was significant, indicating a polytomy at this node was rejected. This topology was also recovered in an ASTRAL-BP analysis of the overlapping set of deletions (fig. S9). Bootstrap support values are shown for 500 replicates. (**C**) High-confidence chromosome breakpoints supporting the monophyly of select superordinal clades. No conflicting breakpoints were found for these nodes.

We also analyzed a set of phylogenetically informative chromosome breakpoints curated in an alignment of contiguous genome assemblies from members of 19 placental mammal orders (*28*). Although breakpoint reuse occurs at a frequency of about 10% across mammals (*20*), an analysis of unique chromosome rearrangements affirmed ordinal monophyly and supported a subset of superordinal clades also recovered by coalescent and window-based phylogenies and deletions, in addition to Atlantogenata (Fig. 3 and table S8). In summary, all analyses converged upon a resolved superordinal tree within Boreoeutheria, with low discordance among the basal nodes of Laurasiatheria and Euarchontoglires.

### Divergence Time & Ordinal Diversification

The paucity of genome-wide discordance in the Cretaceous superordinal phylogeny may be the signature of allopatric speciation processes that isolated small populations of placental mammal ancestors on different fragments of the Gondwanan and Laurasian landmasses. Previous gene-based studies of molecular divergence times have attributed early mammal diversification to continental fragmentation that resulted from a combination of plate tectonics and changes in global sea level (*29–31*). However, some phylogenomic studies (*8, 10, 32*) have produced point divergence estimates for the earliest superordinal branching events 10-15 million years younger and less compatible with vicariance-based hypotheses (fig. S4). These latter hypotheses fail to explain the hierarchical biogeographic pattern apparent in the four superordinal clades (*33*).

To test these competing hypotheses, we estimated molecular timetrees using MCMCtree in PAML (*34, 35*) from 316 independent 100-kb windows spread across the three autosomes and the X chromosome, using 37 soft-bounded fossil calibrations for 65-taxa (Fig. 4A, table S10 and fig. S5, S6). This approach allowed us to generate numerous independent datasets that sample adequate numbers of informative sites (table S7) and are not constrained by protein-coding gene size, mitigating the influence of locus tree error (*36*) and genomic undersampling, factors that have previously been demonstrated to bias divergence time estimates (*37*). Most (97.7%) of the sampled bases in these windows are non-coding (Table 2). The resulting age estimates were highly consistent across locus trees and chromosomes (Fig. 4B and fig. S7) and were robust to PhyloP classification (table S10), root age constraints, removal of large-bodied and long-lived mammals, and missingness (Fig. 5A and table S10). Estimated locus tree divergence times were consistent with those obtained from the concatenated 241-species nearly neutral dataset (Fig. 5A), which included an additional 23 fossil calibrations (tables S9 and S10).

**Fig. 4.**
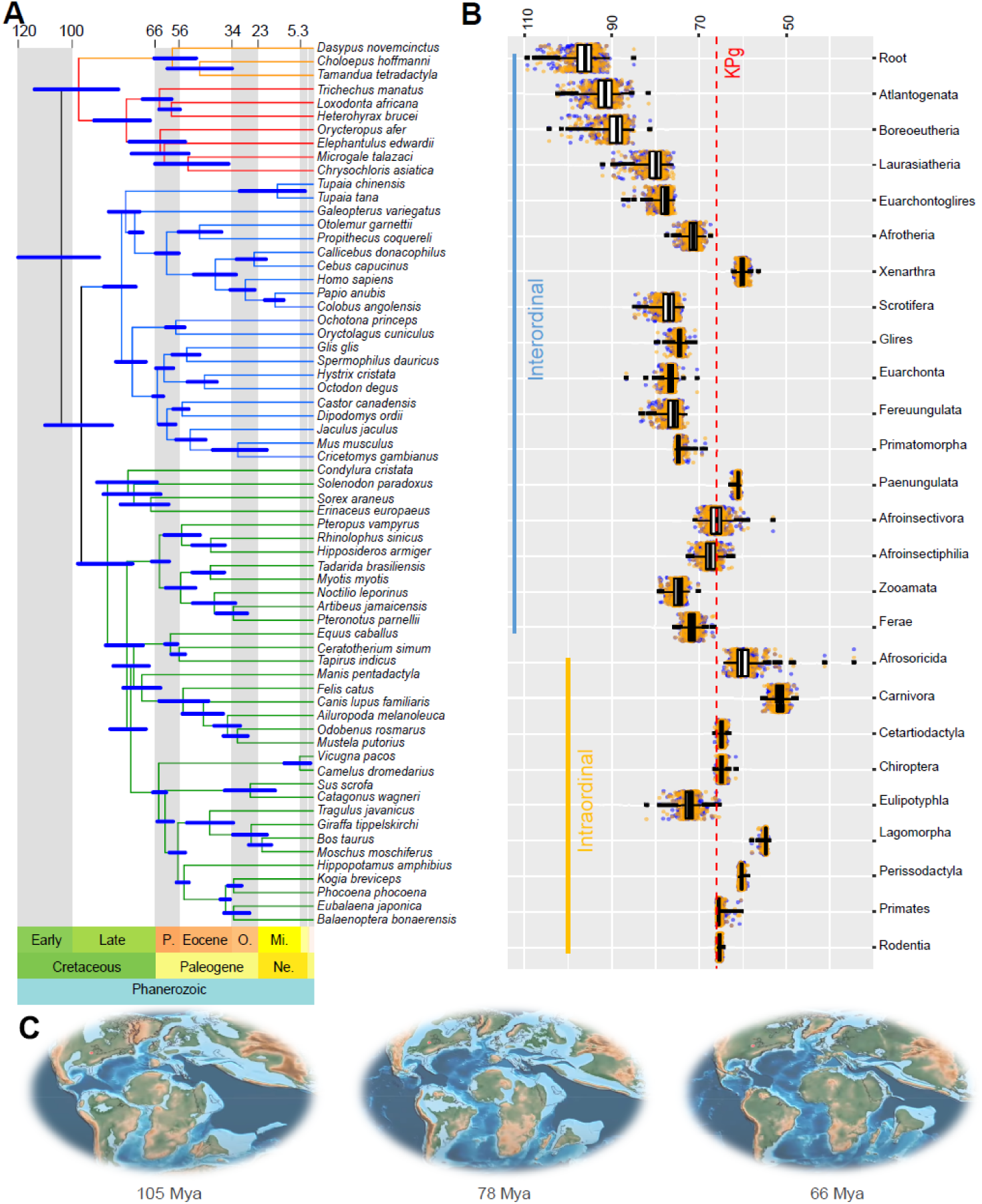
Genomic timescale for placental mammal diversification. Divergence times estimated with 37 fossil calibrations for interordinal and intraordinal diversification events in mammals. (**A**) A representative topology from ChrX showing divergence times and confidence intervals for 65-species estimated using the Benton2009 root constraint and the IRM clock model. (**B**) Genomic estimates for major placental mammal clades based on 316 100-kb windows using the Benton2009 + IRM analysis, distributed across Chr1, Chr21, Chr22, and ChrX. The boxplots summarize the mean and variation around the mean. The corresponding upper 95% confidence interval and lower 95% confidence interval are displayed as blue and orange circles, respectively, for each of the 316 estimates. The related minimum, maximum, mean, and median 95% confidence intervals are listed in Table S10. (**C**) Paleomaps (*38*) illustrate the extent of continental fragmentation and sea level rise at a series of time points during the Cretaceous.

**Fig. 5.**
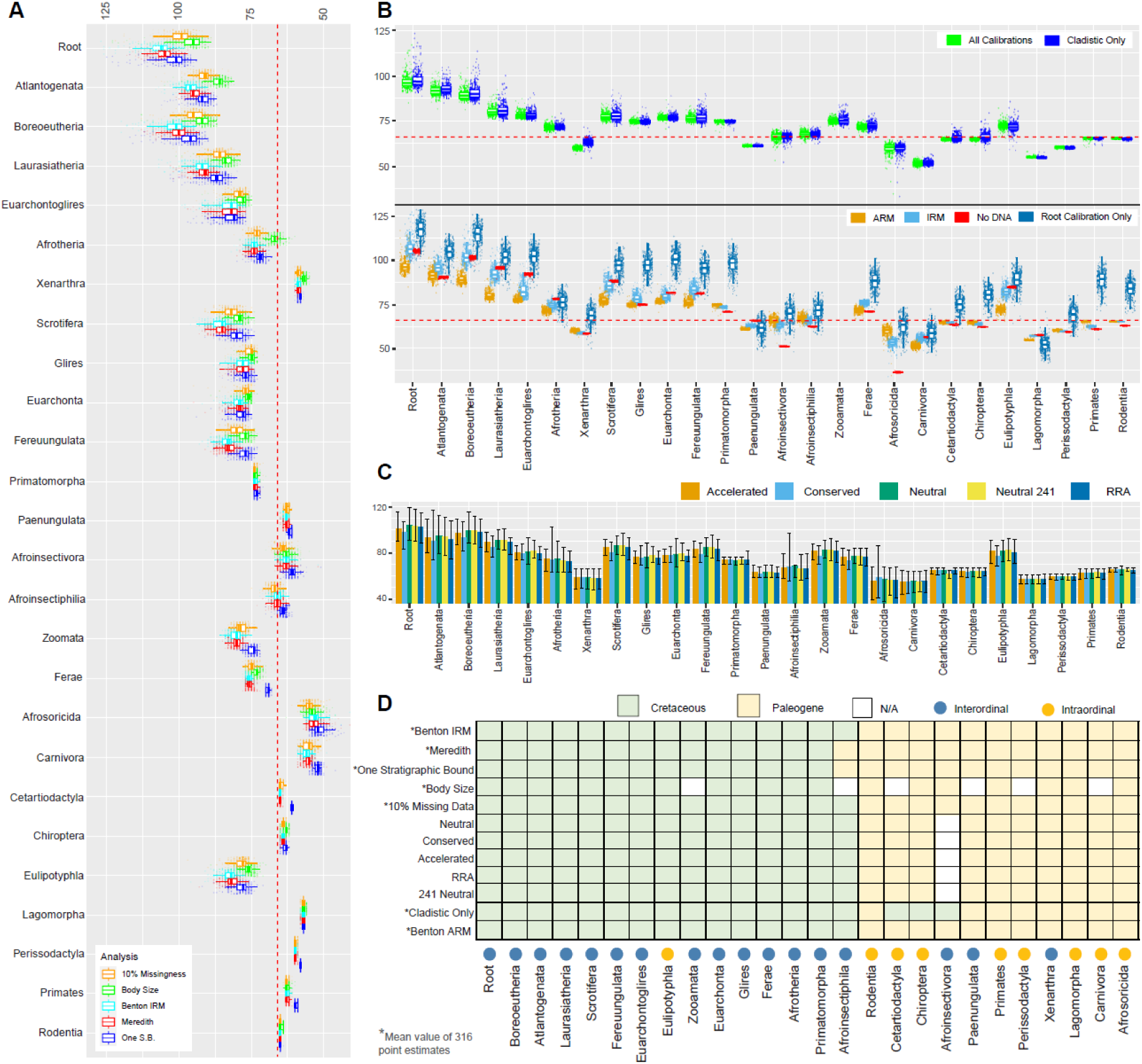
Divergence time sensitivity analyses. For analyses where 316 trees are used, point divergence time estimates for all 316 timetrees are displayed. The overlaid boxplots show the mean of 316 point estimates. The corresponding minimum, maximum, mean and median 95% confidence intervals are listed in Table S10. (**A)** Variation in node ages when the root constraint, stratigraphic bounds, correcting for body size, and missingness are varied. (**B)** Comparison of point estimates when the tree is fully calibrated using a combination of ‘cladistic’ (fossils assigned to a node based on a formal cladistic analysis) and ‘opinion’ fossil constraints relative to point estimates calibrated only with ‘cladistic’ fossils (see table S9). The lower portion of the plot compares divergence times estimates using the independent (IRM) or autocorrelated (ARM) rate models. The effective joint prior (No DNA) is compared to divergence times estimated when only the root of Placentalia is calibrated using the Benton 2009 soft bound upper constraint. (**C**) Comparison of point estimates and 95% confidence intervals for single tree datasets where selective pressure, genome alignment reference species, and the number of species are varied (Table S10). (**D)** The inferred ages of select interordinal (blue dots along x axis) and intraordinal divergences (yellow dots along x axis) across the range of sensitivity analyses are listed in Table S10.

Altogether our results support a hypothesis wherein continental fragmentation and sea-level changes likely played an important role in the superordinal diversification of placental mammals (*29, 31*). Under this hypothesis, the origin of placental mammals is placed at approximately 102 million years ago (Mya) (mean of 316 upper and lower 95% CI 90.4 – 114.5, all divergence-date estimates in main text correspond to the Benton Averaged dataset, see table S10 for full description and summary). The earliest divergences within Atlantogenata and Boreoeutheria also occurred in the Cretaceous Period at 94 Mya (95% CI 80.5 – 108.2) and 96 Mya (95% CI 86.5 – 105.9), respectively. The timing of these events coincides with Africa’s geological fragmentation from South America (∼110 Mya onwards) and with parts of Laurasia (*38*). Interordinal divergences within Laurasiatheria occurred between 81.6 and 73.6 Mya (95% CI 67.9 – 88.29), coinciding with the peak of Cretaceous land fragmentation due to elevated sea levels (∼97-75Mya) (*26, 33*). The origin of Euarchontoglires was dated 80.7 Mya (95% CI 75.0 – 88.3 Mya) and was followed by the afrotherian radiation that commenced at 73.0 Mya (95% CI 67.9 – 79.3 Mya).

We performed a suite of sensitivity analyses to demonstrate that these results were robust to variation in the underlying molecular dataset (Fig. 5A), the usage of different subsets of fossil calibrations (Fig. 5B), and the model of lineage-specific rate variation (Fig. 5C). Despite the minor observed differences in point time estimates across genomic windows, when we consider their uncertainty, a majority of analyses support the Long Fuse model of placental mammal diversification (Fig. 5D) (*39*). Our results contrast with many previous studies that instead support four alternative models of diversification (*4*). The remarkably consistent divergence time point estimates across locus trees may also be related to the high proportion of parsimony informative sites in our analyzed genomic windows. Marin and Hedges (*37*) suggested that genomic undersampling can result in biased divergence times. They used simulations to demonstrate that the number of sites required to recover divergence times accurately scales with the number of tips in a phylogeny. For example, roughly estimating from their regression analysis ∼4,000 variable sites are necessary to infer accurate divergence times for a tree containing 65 taxa. The number of parsimony informative sites in the genomic windows we sampled exceeds this threshold and contains, on average, 43,881 parsimony informative sites in the 65 species datasets alone (table S7) (*6*).

In contrast to strong evidence for superordinal divergences occurring almost entirely in the Cretaceous Period, intraordinal diversification mainly was restricted to the early Paleocene, immediately following the KPg extinction event, 65.3-53.6 Mya (95% CI 45.6 – 66.8) (Fig. 4B) (*40*). The Paleocene also saw the ordinal diversification of Xenarthra and the two primary afrotherian lineages, Paenungulata and Afroinsectiphilia. This result represents a molecular signature of the KPg extinction event influencing ordinal diversification. Only Eulipotyphla is estimated to have begun to diversify in the Cretaceous period (mean estimate =77.4 Mya) (95% CI 68.9 – 86.8). However, we demonstrate the sensitivity of some ordinal divergence estimates to different fossil calibration strategies (Table S10), highlighting the need for the development of improved divergence time models that account for molecular rate variation correlated with life-history traits.

### Phylogenomic Conflict in the Cenozoic Era

In contrast to the well-resolved lineage diversification events in the Cretaceous, Cenozoic branching events showed higher levels of phylogenomic discordance, which we hypothesize may have resulted from larger population sizes and markedly greater geographic continuity within and between continents at this time (Fig. 2) (*31*). The earliest radiations of New World and Old World primates show evenly distributed amounts of topological conflict across autosomal and ChrX locus trees and high and low partitions of GC content, both characteristic of ILS but not introgression (*13, 41*). By contrast, several other clades show strikingly different topological and GC content distributions between the autosomes and the X chromosome (Fig. 2), a pattern observed in cases of speciation with gene flow (*15, 16, 42, 43*). For example, the inferred species tree uniting sciuromorph and hystricomorph rodents is enriched on the X chromosome and the center of Chr1, regions of the ancestral placental mammal karyotype that are predicted to have historically lower rates of recombination. However, this topology is depleted on the small autosomes and the telomeric ends of Chr1, where ancestral reconstructions predict historically higher rates of recombination (tables 1 and S5) that lead to locus tree conflict.

A basal position for ursids is supported across most locus trees within arctoid carnivorans. However, it is noteworthy that there is strong enrichment for an ursid+musteloid clade found within two ChrX recombination coldspots that are enriched for the species tree in other carnivoran families (*16, 44*). We hypothesize that gene flow between the ancestors of musteloids and pinnipeds may have erased the species tree history across the autosomes, which was retained in the center of the low recombining region of ChrX mirroring observations in other animal clades (*15, 17*). Locus trees for cricetid rodents also reveal a very high disparity in ChrX versus autosomal signal, with ChrX enriched for a *Cricetulus*+*Ondatra* clade as the most probable species tree echoing findings from phylogenomic studies of other muroid rodents (*45*). Low GC content profiles similarly track the inferred species trees in each Cenozoic clade (Fig. 2) (*21, 46*). In summary, our findings highlight phylogenetically dispersed X-autosome discordance throughout the Paleogene and Neogene (Fig. 2 and table S10), a pattern absent throughout the first 25 million years of the superordinal placental mammal radiation.

## Discussion

George Gaylord Simpson (*47*) predicted that “*complete genetic analysis would provide the most priceless data for the mapping of this stream*”, referring to the resolution of mammalian phylogeny, a classic and recalcitrant problem in evolutionary biology. Our comprehensive analysis of the 241 placental mammal whole genome alignment confirms Simpson’s prediction. It establishes a standard for phylogenomics that maximizes the value of genome sequences at deep taxonomic levels and moves beyond constrained, gene-centric approaches (*1*). Based on the preponderance of evidence across multiple variants of divergence time estimation, we propose that the combination of two major Cretaceous events played a fundamental role in the successful radiation of crown placental mammals in the Paleogene. First, increased continental fragmentation promoted lineage isolation (Fig. 4C), followed by the most rapid episode of land emergence during the Mesozoic (*38*). This second event would have set the stage for the emergence of morphologically-diagnosable orders in the ecological vacuum that followed the mass extinction of non-avian dinosaurs 66 Mya. We envision a similar resolution of longstanding controversies across the Tree of Life with improved utilization of the historical information encoded within living genomes.

### Materials and Methods Summary

Genome-wide coalescence and concatenation phylogenies were generated using three differently referenced versions (human, dog, and inferred ancestor at the root) of the HAL alignment. Human referenced, single base pair resolved PhyloP scores were used to define genome-wide SNPs corresponding to accelerated, conserved, and neutrally evolving regions of the alignment to explore the impact of selective constraint on coalescent and concatenation-based phylogenomic inference. The conservation of karyotypic position across all placental mammals was used to infer the historical recombination rate for three autosomes (Chromosomes 1, 21, and 22) and the X chromosome to interrogate the role of genomic architecture and recombination in the distribution of phylogenomic signal for challenging to resolve nodes. ML trees were generated from consecutive 100kb windows across each chromosome for each clade examined. The frequency of each competing topology was calculated and compared across the X and autosomal locus trees and regions of high and low GC content (a proxy for recombination rate). Divergence time estimates were generated using MCMCtree in PAML and were calibrated using a suite of soft bounded fossil calibrations. Wide-ranging sensitivity analyses were performed, varying both the underlying molecular dataset and the fossil calibrations.

## Acknowledgments

We thank M. Dickens and the Texas A&M High Performance Research Computing Center for assistance, and M. Dong, D. Genereux and J. Johnson for facilitating data management. We also are grateful to the members of the Zoonomia Consortium (full list below), particularly K. Lindblad-Toh, E. Karlsson, and K. Pollard, for insightful and critical feedback.

## Author contributions

Conceptualization: WJM, MSS; Data curation: NMF, KRB, JD; Investigation: all authors; Project administration: WJM, HAL, MSS; Software: VCM, AJH, JD; Writing – original draft: WJM, NMF; Writing – review & editing: all authors.

## Data and materials availability

All datasets used in this analysis are available where indicated in the text. Scripts written as part of this study are available at https://github.com/VCMason/Foley2021 and also archived at https://doi.org/10.5281/zenodo.5793715. The HAL alignment is publicly available at https://cglgenomics.ucsc.edu/data/cactus/. Information regarding genome assemblies and specimen biosamples are provided in Ref. 1 and can be accessed at https://zoonomiaproject.org/the-data/. Human referenced PhyloP scores are publicly available at http://genome.ucsc.edu/cgi-bin/hgGateway?genome=Homo_sapiens&hubUrl=http://cgl.gi.ucsc.edu/data/cactus/241-mammalian-2020v2-hub/hub.txt. All other data including alignments, phylogenies, and Excel versions of the Supplementary Tables are available at the following repository <u>10.5281/zenodo.5823345</u>.

## Supplementary Materials

Materials and Methods

Supplementary Text

Figs. S1 to S9

Tables S1 to S10 References (*49–163*)

Data S1 to S2

