## SupplementaryMaterial for "A genomic timescale for placental mammal evolution"

**This PDF file includes:**

Materials and Methods
Supplementary Text
Figs. S1 to S9
Tables S1 to S10
Data S1 to S2

### Materials and Methods

Alignments

The following three versions of the alignment were used in different analyses to investigate the potential impact of reference bias: 1) human-referenced assembly (HRA), 2) dog-referenced assembly (DRA) and 3) root referenced assembly (RRA). MAF-formatted chromosome or scaffold alignments were extracted from the HAL alignment (available at https://cglgenomics.ucsc.edu/data/cactus/) with a parallelized version of the command hal2maf from halTools (*49*), using either *Homo sapiens*, *Canis lupus familiaris*, or the HAL inferred ancestor (*10*) at the root of the tree (fullTreeAnc239) as the reference. We removed duplicates when more than one sequence was present for a species in the alignment using the mafDuplicateFilter function (https://github.com/dentearl/mafTools). In such cases, we retained the sequences most similar to the consensus of all other sequences in that region. In situations where multiple sequences shared the same nucleotide identity to the consensus, one was randomly chosen for inclusion. We excluded protein coding-only analyses because visual inspection of alignments revealed some homology and local mis-alignment issues (e.g., a 5' end of an exon was not orthologous across pteropodids). A similar inspection of non-coding alignments revealed no overt homology errors. MAF-formatted alignments were then converted to fasta format using mafFilter (*50*) and alignment blocks stitched together using a custom script (https://github.com/VCMason/Foley2021) to form a chromosome-by-chromosome 241 species fasta alignment. Given that the root referenced version of the HAL alignment comprised a much larger number of scaffolds (> 11,000), only scaffolds ≥ 5000bps were converted to MAF format.

Three datasets were prepared from each of these alignments. The first alignment contained all 241 species from the original Zoonomia HAL alignment. The second alignment contained 228 species and reduced missingness. We constructed this alignment because 13 species consistently exhibited a higher degree of missingness and were removed based on a preliminary analysis of a subset of 100kb sliding window alignments and trees (see Table S1). A third dataset that sampled 65 species was constructed to estimate ordinal-level relationships and reduce computation time. The species in the reduced 65 species dataset were selected to represent major superordinal clades and index ordinal diversity in the phylogeny, while also maximizing nodes to which fossil calibrations could be applied. Taxa were selected that maximized quality where possible (e.g., selecting human, mouse, dog, cat, cow) and reduce missingness (Table S1).

Coalescence Analysis

SVDquartets (*51*) was chosen for the coalescence-based species tree approach to satisfy the assumption of no intra-locus recombination under the multispecies coalescent model (MSC). SNP sites were used to impute quartet trees which were then aggregated to infer a species tree. In total, nine datasets were analyzed using SVDquartets (Table S2). To determine if the results were heavily impacted by reference bias we pruned the HRA, DRA and RRA alignments to 65 species and compared phylogenies. We also examined the impact of missingness on phylogenetic inference by analyzing HRA datasets where the percentage of missingness was varied (10%, 25% and 50%).

Many published coalescence-based analyses have included only protein coding regions or ultraconserved elements, thereby violating the MSC's assumption of neutrality. To explore differences in trees due to selective constraint we leveraged per-base PhyloP scores (*52*) estimated across the human referenced alignment of 241 species (available from http://genome.ucsc.edu/cgi-bin/hgGateway?genome=Homo_sapiens&hubUrl=http://cgl.gi.ucsc.edu/data/cactus/241-

mammalian-2020v2-hub/hub.txt). Rather than constructing datasets based on extreme PhyloP scores, we used a quintile-based approach that allowed us to control for dataset size when comparing phylogenomic signal from genomic regions under different selective constraint. Genome-wide PhyloP scores were ordered and assigned to quintiles. The extremes of the quintile distribution were used to define conserved sites (1st quintile, containing positive scores 8.903 – 0.581) and accelerated sites (5th quintile, containing negative scores -20 – -0.457). The central, or 3rd, quintile was used to define nearly neutral sites, so called as they span values ranging from 0 – 0.268 (referred to as 'neutral' from here on). To prepare the final PhyloP score-based datasets, uninformative constant sites were removed. Neutral, conserved and accelerated datasets containing 65 species were analyzed individually. We also analyzed a neutral dataset containing all 241 species (Table S2).

SNP sites were extracted from each 241 species referenced fasta alignment (i.e., HRA, DRA and RRA) in vcf format using SNP-sites v2.5.1(*53*). The Y chromosome alignment was excluded from all analyses owing to the difficulty of assembling and accurately aligning this complex region using short read data (*54*, *55*). SNPs were thinned to a minimum of 1000bp between sites using VCFtools v1.16 (*56*) and converted to nexus format using the script vcf2phylip.py (https://github.com/edgardomortiz/vcf2phylip). By using SNPs that are fairly widely spaced, we aimed to account for the requirement of free recombination between loci that is an assumption of coalescent methods like SVDquartets. All datasets were further filtered on missingness, stripping columns from the alignment that contained more than 10% missingness. As such, while data were initially thinned to a minimum distance of 1000bp apart to avoid intralocus recombination, the final distance between most SNPs is likely much larger. Data from all chromosomes were concatenated using Geneious Prime 2019.04. A 50% majority rule tree was estimated using SVDquartets implemented in PAUP* v4a168 (*57*) using the QFM quartet amalgamation technique (*58*). The number of quartets sampled was dataset-dependent (see Table S2).

Maximum Likelihood Concatenation-based Analyses

Datasets listed in Table S2 were also concatenated and analyzed with IQ-TREE v1.6.10 (*59*) to generate maximum likelihood (ML) trees. Given that our datasets contained only SNPs or selected variable positions (PhyloP sites), we used the +ASC option to correct for ascertainment bias caused by the exclusion of constant sites in our analyses. Trees were analysed as a single partition using a GTR + I + G model of sequence evolution to facilitate the use of the +ASC option. The +ASC option is currently incompatible with complex mixture models. Trees were evaluated with 1000 bootstrap replicates using the ultrafast bootstrap approximation (*60*, *61*). The best scoring ML tree was selected for further comparative analyses. The impact of lineage-specific rate variation on divergence time estimation in mammals has been previously demonstrated (*6*). As such, we selected the GHOST model of rate variation (*62*), a mixture model that accounts for lineage-specific rate variation across sites. An ML tree was generated using IQ-TREE (*59*) under a GTR model of sequence evolution and the GHOST model of rate variation (*62*) (GTR*H4). The resulting topology was evaluated with 1000 bootstrap replicates using the ultrafast bootstrap approximation (*60*, *61*).

Distribution of Phylogenomic Signal

*Sliding Windows Analysis*

An emerging number of phylogenomic studies that sample divergent clades in the Tree of Life have demonstrated that phylogenetic signal is non-randomly distributed throughout the genome. In clades where species relationships have been difficult to resolve due to high gene tree conflict, studies have shown that the phylogenetic signal most consistent with the species tree has often been preserved and enriched on the X or Z chromosomes (*15–17*, *43*, *44*). This observation is particularly pronounced within conserved low recombining regions of the X chromosome (ChrX) in placental mammals (*16*). Conversely, discordant gene tree signal tends to be enriched near the telomeres of large chromosomes, or enriched on small chromosomes which have relatively elevated recombination rates (*2*, *15*). Most of these prior studies focused on recently diverged species groups with collinear genomes and resolved recombination maps, facilitating comparisons of phylogenomic signal and recombination rate. However, applying a similar approach across the placental mammal phylogeny is hindered twofold: first, by the lack of resolved recombination maps for most mammalian species, and second, mammalian autosomes have undergone considerable karyotypic rearrangement over evolutionary time (*3*, *63*). Despite these confounding issues, certain human chromosomal associations, and their relative positions within different species' karyotypes, have been maintained over deep stretches of mammalian evolution (*3*, *63*, *64*). A previous study based on multispecies comparative chromosome painting data and comparative gene mapping data presented evidence that human Chr1 was intact as a single chromosome in the most recent common ancestor of placental mammals (*18*). The gene order observed in human has also remained relatively stable since the placental mammal ancestor (*20*), allowing us to hypothesize that a lower historical recombination rate exists across the center of Chr1 when compared to its telomeric ends and to smaller autosomes. Chr21 and Chr22 are the smallest human autosomes, and despite millions of years of chromosomal rearrangements across the placental mammal phylogeny, syntenic fragments from these two chromosomes have typically occupied a distal or telomeric chromosomal position in the majority of living species for which comparative karyotypic data exist (Table S5). This novel observation allowed us to infer a historically high rate of recombination on these two chromosomes. In contrast, the X chromosome in placental mammals contains some of the highest (pseudoautosomal region = PAR) and lowest recombination rates (recombination desert) and has remained collinear in a large proportion of species. Indeed, previous studies have demonstrated strong conservation of the ChrX recombination landscape among species from different orders (*16*). As such, we chose Chr1, Chr21, Chr22, and ChrX for our sliding windows analysis given that our inference of their historical patterns of recombination rate were the most justifiable.

Human referenced Chr1, 21, 22 and ChrX alignments were trimmed to 228 species to reduce missingness across the alignment (Table S1) and divided into 100kb windows. We analyzed all windows for Chr21, Chr22, and ChrX, and every fifth window for Chr1 given its much larger size (Table 1). Alignment columns using the setting *-gt 0.50* to allow only 50% missingness per alignment column in trimAl and the alignment filtering script using parameters to accommodate filtering a combination of non-coding and coding regions: cutoff=0.80, Zscore=6, windsize=50, step=1 and *Homo sapiens* as the reference. Final alignment statistics were calculated per window and per species using AMAS.py (https://github.com/marekborowiec/AMAS) (*65*). After filtering, only windows 10kb or greater were retained for further analysis. The final number of windows per chromosome and the average length per window post-filtering are listed in Table S3. Missingness in filtered alignments was also summarized by species. These values were then averaged across all windows to assess missingness in a broader phylogenetic context (Table S1).

To estimate the proportion of coding and non-coding sequence represented in the filtered alignments, the human sequence was extracted from each alignment and mapped back to the human genome assembly (hg38) using minimap2 v2.10 with the settings *-ax asm5* (*66*). Resulting bam files were sorted and indexed using samtools v1.8 applying default settings (*67*). A bed file containing these coordinates was prepared using the genomecov function in bedtools with the options *-bga -split* (*68*). We chose the consensus human coding sequence (ccds) exon annotation (*69*) which comprises stable gene annotation sets that are consistently represented across the most commonly used genome browsers. The proportion of the filtered alignment that overlapped with human exonic sequence was determined using the intersect, sort and merge functions in bedtools. The proportion of neutral sites, those which overlapped with the nearly-neutral PhyloP quintile sites defined above, in the alignments were queried using the intersect, sort and merge functions in bedtools.

ML trees were generated for each window using IQ-TREE under a GTR model of sequence evolution and the GHOST model of rate variation (GTR*H4). Trees were evaluated with 1000 bootstrap replicates using the ultrafast bootstrap approximation. Given that the HAL alignment did not contain outgroups to placental mammals, we assumed the root position between Boreoeutheria and Atlantogenata in all analyses given its highly consistent recovery in the majority of studies to date (*6*, *8*, *9*, *70–72*).

*Chromosomal Distribution of Phylogenomic Signal*

Relationships for the most difficult to resolve superordinal nodes in the phylogeny (*5*, *73*) were summarized in four-taxon trees with three ingroup taxa and an outgroup (Fig. 2; Table S6). Sub-trees containing the four taxa required to test a given hypothesis (Table S6) were extracted from the 228 species sliding windows analysis trees using *tree_doctor* from the program PHAST v1.5 (*74*) and node information removed using gotree (https://github.com/evolbioinfo/gotree). The relative frequency of each topology was calculated for the autosomes and ChrX separately. Within ChrX we also partitioned the largest, historically conserved low recombining region, which corresponds approximately to the intervening sequence flanked by the genes *JADE3* (~47Mb) and *CHRDL1* (~110Mb) on human ChrX (*5*). Because detailed recombination maps are not available for most species, we also assayed topological frequency distributions based on GC content, a proxy for recombination rate. GC-biased gene conversion occurs during meiotic recombination and leads to strong correlations between recombination rate and GC content (*71*). As recombination rate can vary between species (*75*), GC content was calculated in alignment windows containing only the ingroup species for a given clade (Table S6).

Rare Genomic Changes

*Long Deletions*

Rare Genomic Changes (RGCs) are useful phylogenomic characters as they occur more infrequently than the majority of nucleotide substitutions and are therefore thought to be relatively homoplasy free (*76*). Our survey was restricted to deletions because MAF-formatted alignments output from the HAL alignment were aligned to a gapless reference sequence. While the HAL to MAF conversion tool does allow for users to allow gaps of a given length in the reference sequence, we did not elect to use this option to assess insertions because it would inherently introduce a length bias.

Counts of deletions were compiled per competing phylogenetic hypotheses for a given set of taxa. We focused on relationships among laurasiatherian and euarchontoglires orders as these are among the most controversial in the literature (*5*). Both the human (to assess laurasiatherian deletions) or dog (to assess deletions in Euarchontoglires) and the root referenced (for both) alignments were used to investigate potential reference bias. Deletions were surveyed using a script modified from Mason *et al* (*77*) (https://github.com/VCMason/Foley2021). The analysis was run using conservative parameters: minimum deletion length of 10bps, all taxa listed in the hypothesis file (Data S1) must share the deletion, and continuous sequence across the deletion was required in all outgroup taxa listed in the hypothesis file.

We also surveyed deletions supporting competing hypotheses among the major rodent lineages, however, none were found using the criteria listed above likely due to varying levels of genome completeness within and across clades: average BUSCO completeness among new Zoonomia sequenced rodent genomes was 65.37%, but ranged from 14.50% to 90.10% (*1*). We note this result may hinder our ability to fully evaluate hypotheses for the position of Scandentia, however, Euarchonta is recovered by all other tree-based analyses here and a previous study which examined genome-wide sets of indels (*78*).

All resulting deletions were visually inspected to ensure sufficient alignment quality surrounding a deletion. The following criteria were used: 1) additional phylogenetically informative deletions were only allowed within 10bp of either side of the focal deletion, 2) individual sequences flanking the deletion were allowed gaps within 10bp of the deletion (most likely representing local assembly errors or quality issues) if the sequence from a closely related congener was complete, and 3) a continuous sequence was required in all listed outgroups (*21*). This conservative approach did not result in rank order changes to the raw number of deletions recorded for any hypothesis. Hundreds of raw deletions were recovered supporting Euarchonta. In contrast, no deletions were recovered for the Glires + Primatomorpha (where Scandentia is basal) or the Glires + Scandentia hypotheses. Given this result, only a subset of Euarchonta deletions were further validated, as even a small number of validated deletions could be considered significant. For each candidate deletion we used blastn (*79*) to query an ingroup sequence (containing the deletion) against the corresponding NCBI assembly in order to confirm the candidate deletion was not an alignment artifact. Given that we observed a small amount of variation between differentially referenced alignments, we also determined the overlap between deletions called using the placental root ancestor and the human or dog referenced alignments as a maximally conservative dataset. For each hypothesis, the human sequence was extracted from the root and human or dog referenced alignments and were queried against the human genome (hg38). For each deletion the best hit was extracted and identified by the chromosomal coordinates of the top BLAT hit. Statistical support for competing phylogenetic hypotheses was evaluated using the KKSC test (*48*).

ASTRAL_BP uses low-homoplasy rare genomic changes to infer tree topologies in an ILS-aware framework (*80*). Previous studies have applied this approach to counts of retroposon insertions; here we applied the method to our Laurasiatheria dataset of deletions which overlap between the analyses of the HRA and RRA. The deletion dataset was converted to a presence/absence matrix. For the ASTRAL_BP analysis, the presence-absence matrix was converted into a series of incompletely resolved "gene trees", each representing a single deletion event, using the phylotools scripts (https://github.com/ekmolloy/phylotools). ASTRAL-II v4.11 (*81*) was used to estimate the frequency of each quartet derived from the deletion presence/absence matrix and a maximum quartet support supertree (MQSS) was computed using the exact mode (-x) and 500 bootstrap replicates.

As an additional check to our data, we investigated the impact of genome quality on the number of raw deletions recovered for each clade. The deletion detector script was re-run, requiring only one species that had a high-quality genome for each clade. For hypotheses within Euarchontoglires, the deletion had to be present in *Mus musculus*, *Homo sapiens*, *Oryctolagus cuniculus* and *Tupaia chinensis* for the raw deletion to be called. For hypotheses within Laurasiatheria, the deletion had to be present in *Felis catus*, *Bos taurus*, *Equus caballus* and *Macroglossus sobrinus* for the raw deletion to be called. For Laurasiatheria the rank order of the number of deletions identified per hypothesis did not differ from results obtained when all species per clade were required. A similar result was observed in Euarchontoglires. No deletions were recovered for the hypothesis where Scandentia is basal to Glires + Primatomorpha. A small number of raw deletions were recovered for Scandetia + Glires (n=31), while a large number of raw deletions were found for Euarchonta (n=1528). Despite this minor difference between analyses, the rank order and relative proportions of the deletions were the same indicating that variable genome quality among and within orders did not affect our results.

*Chromosome Breakpoints*

Chromosome breakpoints were inferred from a dataset comprising seven outgroups species (chicken, alligator, platypus, opossum, koala, wombat and tasmanian devil) and 27 ingroup species representing all extant placental mammal orders (*28*). Evolutionary breakpoint regions (EBRs), relative to the mammalian ancestor (*28*), were identified and classified following (*82*). The final EBR dataset was manually inspected to ensure accuracy (Table S8). Where the majority of species in a given clade contained a breakpoint that was absent in 1-2 species contained within that clade, the corresponding regions from the outlier genomes were manually checked for multiple contig joins and gaps indicative of poor local assembly quality.

Divergence Times

Divergence times were estimated using the approximate likelihood (*83*) calculation in MCMCtree from the PAML package (*35*). For each sliding window, individual ML trees and their corresponding alignments were pruned to 65 taxa (Table S1), sampling multiple members from all extant placental orders. Each analysis was calibrated using 37 soft bound fossil calibrations (Table S9). As MCMCtree requires an input topology, Phybin (*84*) was used to identify 316 trees across ChrX and the autosomes that shared an identical topology which was broadly congruent with relationships recovered in our whole genome coalescent and concatentation based analyses. Each of these trees was also characterized by a large number of parsimony informative sites (Table S7). Maximum and minimum soft-bound fossil calibrations cited by previous studies were reviewed and amended using updated fossil information and/or adjusting boundary age using the International Chronostratigraphic Chart v2020/03 where applicable (Table S9). The analyses were run with the HKY85 model of sequence evolution. Details listing the datasets, models, and number of trees used in all divergence time analyses are listed in Table S10.

*Sensitivity Analyses*

We evaluated the sensitivity of divergence time estimates to changing the root constraint. We first applied an upper bound of 131.5Mya ('Benton2009 constraint') based on the well documented fossil species *Eomaia* (*85*) and *Sinodelphys* (*86*), estimated to be Aptian or Barremian in age (*87*). A conservative estimate of 131.5Mya is used to reflect the uncertainty around the dating of the horizons from which these fossils are derived (*87*). We also evaluated a root constraint based upon upper 95% confidence intervals from a prior molecular study that reported a much younger point estimate of 116.81Mya (Meredith) (*6*). To investigate potential biases due to missing data we also repeated the analysis using the Benton2009 constraint while allowing only 10% missing data in the 316 previously identified alignments. After filtering, 241 alignments with ≥5kb of data were retained for further analysis.

Using the 316 sliding windows datasets and the Benton2009 root constraint, we performed an additional analysis for nodes where stratigraphic bounding was used to provide an upper calibration (Table S9). Here we applied stratigraphic bounding based on one stage instead of two, applied to the 316 sliding windows datasets and employing the Benton2009 root constraint. Conversely, we also performed additional sensitivity analyses where the softbound upper constraint for Primates, all calibrations in Rodentia, Cetartiodactyla, and Perissodactyla was arbitrarily extended to 75 Mya and the upper constraint on Primatomorpha was removed (Table S10) to assess the joint effect of relaxing upper constraints on these nodes. We also performed an analysis extending the softbound upper constraints for all calibrations in Rodentia, and removing the upper constraint on Primatomorpha. These latter two analyses were run using both the independent rates (IRM) and autocorrelated rates models (ARM), with a single upper constraint on the root of Placentalia and without DNA (Table S10).

Given that the considerable evolutionary rate variation predicted by life history correlates (body mass and lifespan) can confound molecular clock models (*88*), we also removed species with body masses of >10kg (*89*) and maximum lifespan >40 years (*90*) following (*91*). This resulted in the removal of 25 species (Table S1), including all perissodactyls and all but one cetartiodactyl, *Tragulus tragulus*. A divergence time analysis was also performed using the RRA variable site dataset described in Table S1 to investigate potential reference bias.

To assess the impact of selective constraint on divergence time estimates we also estimated divergence times for the conserved, neutral, and accelerated PhyloP datasets (see Coalescence section & Table S2). A 241 species time tree was generated using the HRA nearly neutral PhyloP dataset (see Table S2) and a concatenation-based ML tree calibrated using an expanded set of constraints (Table S9).

The impact of relaxed clock models on the inferred divergence times was compared using the results from the Benton2009 analysis using both the IRM and ARM using the 316 sliding windows datasets. Additional sensitivity analyses were carried out to compare the relative impact of the molecular data and fossil calibrations on our divergence time estimates. The Benton2009 analysis was run using only molecular data and a single calibration on the root or just fossil calibrations with no molecular data under an autocorrelated clock model. This allowed us to explore whether (a) our suite of soft-bounded fossil constraints were 'pre-determining' our inferred divergence times, (b) if fossil constraints were violated, and (c) to what extent the molecular data informed the results. Furthermore, fossil calibrations were classified as either 'opinion' or 'cladistic' depending on the method used to assign them to a given node in the phylogeny. Those labeled 'cladistic' are based on a formal cladistic analysis that assigned the fossil to a specific node (Table S9). We then performed analyses with the autocorrelated clock model using only soft bound fossil calibrations and applying only fossil calibrations classified as 'cladistic'.

In all cases, the MCMC chain was run for 200,000 generations sampling every tenth generation, with 2,000 trees discarded as burn-in. Divergence times were collated using MCMCtreeR (*92*) and point divergence time estimates were aggregated and averaged using dplyr (*93*) and ape (*94*). Plots to compare divergence times across all sensitivity analyses were generated and compared in R using the following packages ggplot2 (*95*), plotrix (*96*) and EvobiR (*97*).

### Zoonomia Consortium

Gregory Andrews[1]; Joel C. Armstrong[2]; Matteo Bianchi[3]; Bruce W. Birren[4]; Kevin R. Bredemeyer[5]; Ana M. Breit[6]; Matthew J. Christmas[3]; Hiram Clawson[2]; Joana Damas[7]; Federica Di Palma[8,9]; Mark Diekhans[2]; Michael X. Dong[3]; Eduardo Eizirik[10]; Kaili Fan[1]; Cornelia Fanter[11]; Nicole M. Foley[5]; Karin Forsberg-Nilsson[12,13]; Carlos J. Garcia[14]; John Gatesy[15]; Steven Gazal[16]; Diane P. Genereux[4]; Daniel Goodman[17]; Linda Goodman[18]; Jenna Grimshaw[14]; Michaela K. Halsey[14]; Andrew J. Harris[5]; Glenn Hickey[19]; Michael Hiller[20,21,22]; Allyson G. Hindle[11]; Robert M. Hubley[23]; Laura M. Huckins[24]; Graham M. Hughes[25]; Jeremy Johnson[4]; David Juan[26]; Irene M. Kaplow[27,28]; Elinor K. Karlsson[1,4,29]; Kathleen C. Keough[18,30,31]; Bogdan Kirilenko[20,21,22]; Klaus-Peter Koepfli[32,33,34]; Jennifer M. Korstian[14]; Amanda Kowalczyk[27,28]; Sergey V. Kozyrev[3]; Alyssa J. Lawler[4,28,35]; Colleen Lawless[25]; Danielle L. Levesque[6]; Harris A. Lewin[7,36,37]; Xue Li[1,4,38]; Yun Li[39]; Abigail Lind[30,31]; Kerstin Lindblad-Toh[3,4]; Ava Mackay-Smit[40]; Voichita D. Marinescu[3]; Tomas Marques-Bonet[41,42,43,44]; Victor C. Mason[45]; Jennifer R. S. Meadows[3]; Wynn K. Meyer[46]; Jill E. Moore[1]; Lucas R. Moreira[1,4]; Diana D. Moreno-Santillan[14]; Kathleen M. Morrill[1,4,38]; Gerard Muntané[26]; William J. Murphy[5]; Arcadi Navarro[41,43,47,48]; Martin Nweeia[49,50,51,52]; Austin Osmanski[14]; Benedict Paten[2]; Nicole S. Paulat[14]; Eric Pederson[3]; Andreas R. Pfenning[27,28]; BaDoi N. Phan[27,28,53]; Katherine S. Pollard[30,31,54]; Kavya Prasad[27]; Henry Pratt[1]; David A. Ray[14]; Steven K. Reilly[40]; Jeb R. Rosen[23]; Irina Ruf[55]; Louise Ryan[25]; Oliver A. Ryder[56,57]; Daniel E. Schäffer[27]; Aitor Serres[26]; Beth Shapiro[58,59]; Arian F. A. Smit[23]; Mark Springer[60]; Chaitanya Srinivasan[27]; Cynthia Steiner[56]; Jessica M. Storer[23]; Kevin A. M. Sullivan[14]; Patrick F. Sullivan[39,61]; Quan Sun[62]; Elisabeth Sundström[3]; Megan A. Supple[58]; Ross Swofford[4]; Jin Szatkiewicz[39]; Joy-El Talbot[63]; Emma Teeling[25]; Jason Turner-Maier[4]; Alejandro Valenzuela[26]; Franziska Wagner[64]; Ola Wallerman[3]; Chao Wang[3]; Juehan Wang[16]; Jia Wen[39]; Zhiping Weng[1]; Aryn P. Wilder[56]; Morgan E. Wirthlin[27,28,65]; James R. Xue[4,66]; Shuyang Yao[61]; Xiaomeng Zhang[4,27,28]

Affiliations:
[1]Program in Bioinformatics and Integrative Biology, UMass Chan Medical School; Worcester, MA 01605, USA
[2]Genomics Institute, University of California Santa Cruz; Santa Cruz, CA 95064, USA
[3]Department of Medical Biochemistry and Microbiology, Science for Life Laboratory, Uppsala University; Uppsala, 751 32, Sweden
[4]Broad Institute of MIT and Harvard; Cambridge, MA 02139, USA
[5]Veterinary Integrative Biosciences, Texas A&M University; College Station, TX 77843, USA
[6]School of Biology and Ecology, University of Maine; Orono, ME 04469, USA
[7]The Genome Center, University of California Davis; Davis, CA 95616, USA
[8]Genome British Columbia; Vancouver, BC, Canada
[9]School of Biological Sciences, University of East Anglia; Norwich, UK
[10]School of Health and Life Sciences, Pontifical Catholic University of Rio Grande do Sul; Porto Alegre, 90619-900, Brazil
[11]School of Life Sciences, University of Nevada Las Vegas; Las Vegas, NV 89154, USA
[12]Biodiscovery Institute, University of Nottingham; Nottingham, UK
[13]Department of Immunology, Genetics and Pathology, Science for Life Laboratory, Uppsala University; Uppsala, 751 85, Sweden
[14]Department of Biological Sciences, Texas Tech University; Lubbock, TX 79409, USA

[15]Division of Vertebrate Zoology, American Museum of Natural History; New York, NY 10024, USA
[16]Keck School of Medicine, University of Southern California; Los Angeles, CA 90033, USA
[17]Department of Immunology, University of California San Francisco; San Francisco, CA 94143, USA
[18]Fauna Bio Incorporated; Emeryville, CA 94608, USA
[19]Baskin School of Engineering, University of California Santa Cruz; Santa Cruz, CA 95064, USA
[20]Faculty of Biosciences, Goethe-University; 60438 Frankfurt, Germany
[21]LOEWE Centre for Translational Biodiversity Genomics; 60325 Frankfurt, Germany
[22]Senckenberg Research Institute; 60325 Frankfurt, Germany
[23]Institute for Systems Biology; Seattle, WA 98109, USA
[24]Department of Psychiatry, Icahn School of Medicine at Mount Sinai; New York, NY 10029, USA
[25]School of Biology and Environmental Science, University College Dublin; Belfield, Dublin 4, Ireland
[26]Department of Experimental and Health Sciences, Institute of Evolutionary Biology (UPF-CSIC), Universitat Pompeu Fabra; 08003, Barcelona, Spain
[27]Department of Computational Biology, School of Computer Science, Carnegie Mellon University; Pittsburgh, PA 15213, USA
[28]Neuroscience Institute, Carnegie Mellon University; Pittsburgh, PA 15213, USA
[29]Program in Molecular Medicine, UMass Chan Medical School; Worcester, MA 01605, USA
[30]Department of Epidemiology & Biostatistics, University of California San Francisco; San Francisco, CA 94158, USA
[31]Gladstone Institutes; San Francisco, CA 94158, USA
[32]Center for Species Survival, Smithsonian Conservation Biology Institute, National Zoological Park; Washington, DC 20008, USA
[33]Computer Technologies Laboratory, ITMO University; St. Petersburg 197101, Russia
[34]Smithsonian-Mason School of Conservation; Front Royal, VA 22630, USA
[35]Department of Biological Sciences, Mellon College of Science, Carnegie Mellon University; Pittsburgh, PA 15213, USA
[36]Department of Evolution and Ecology, University of California Davis; Davis, CA 95616, USA
[37]John Muir Institute for the Environment, University of California Davis; Davis, CA 95616, USA
[38]Morningside Graduate School of Biomedical Sciences, UMass Chan Medical School; Worcester, MA 01605, USA
[39]Department of Genetics, University of North Carolina Medical School; Chapel Hill, NC 27599, USA
[40]Department of Genetics, Yale School of Medicine; New Haven, CT 06510, USA
[41]Catalan Institution of Research and Advanced Studies (ICREA); 08010, Barcelona, Spain
[42]CNAG-CRG, Centre for Genomic Regulation, Barcelona Institute of Science and Technology (BIST); 08036, Barcelona, Spain
[43]Department of Medicine and LIfe Sciences, Institute of Evolutionary Biology (UPF-CSIC), Universitat Pompeu Fabra; 08003, Barcelona, Spain
[44]Institut Català de Paleontologia Miquel Crusafont, Universitat Autònoma de Barcelona; 08193, Cerdanyola del Vallès, Barcelona, Spain

[45]Institute of Cell Biology, University of Bern; 3012, Bern, Switzerland
[46]Department of Biological Sciences, Lehigh University; Bethlehem, PA 18015, USA
[47]BarcelonaBeta Brain Research Center, Pasqual Maragall Foundation; Barcelona, 08005, Spain
[48]CRG, Centre for Genomic Regulation, Barcelona Institute of Science and Technology (BIST); 08003, Barcelona, Spain
[49]Department of Comprehensive Care, School of Dental Medicine, Case Western Reserve University; Cleveland, OH 44106, USA
[50]Department of Vertebrate Zoology, Canadian Museum of Nature; Ottawa, Ontario K2P 2R1, Canada
[51]Department of Vertebrate Zoology, Smithsonian Institution; Washington, DC 20002, USA
[52]Narwhal Genome Initiative, Department of Restorative Dentistry and Biomaterials Sciences, Harvard School of Dental Medicine; Boston, MA 02115, USA
[53]Medical Scientist Training Program, University of Pittsburgh School of Medicine; Pittsburgh, PA 15261, USA
[54]Chan Zuckerberg Biohub; San Francisco, CA 94158, USA
[55]Division of Messel Research and Mammalogy, Senckenberg Research Institute and Natural History Museum Frankfurt; 60325 Frankfurt am Main, Germany
[56]Conservation Genetics, San Diego Zoo Wildlife Alliance; Escondido, CA 92027, USA
[57]Department of Evolution, Behavior and Ecology, School of Biological Sciences, University of California San Diego; La Jolla, CA 92039, USA
[58]Department of Ecology and Evolutionary Biology, University of California Santa Cruz; Santa Cruz, CA 95064, USA
[59]Howard Hughes Medical Institute, University of California Santa Cruz; Santa Cruz, CA 95064, USA
[60]Department of Evolution, Ecology and Organismal Biology, University of California Riverside; Riverside, CA 92521, USA
[61]Department of Medical Epidemiology and Biostatistics, Karolinska Institutet; Stockholm, Sweden
[62]Department of Biostatistics, University of North Carolina at Chapel Hill; Chapel Hill, NC, USA
[63]Iris Data Solutions, LLC; Orono, ME 04473, USA
[64]Museum of Zoology, Senckenberg Natural History Collections Dresden; 01109 Dresden, Germany
[65]Allen Institute for Brain Science; Seattle, WA 98109, USA
[66]Department of Organismic and Evolutionary Biology, Harvard University; Cambridge, MA 02138, USA

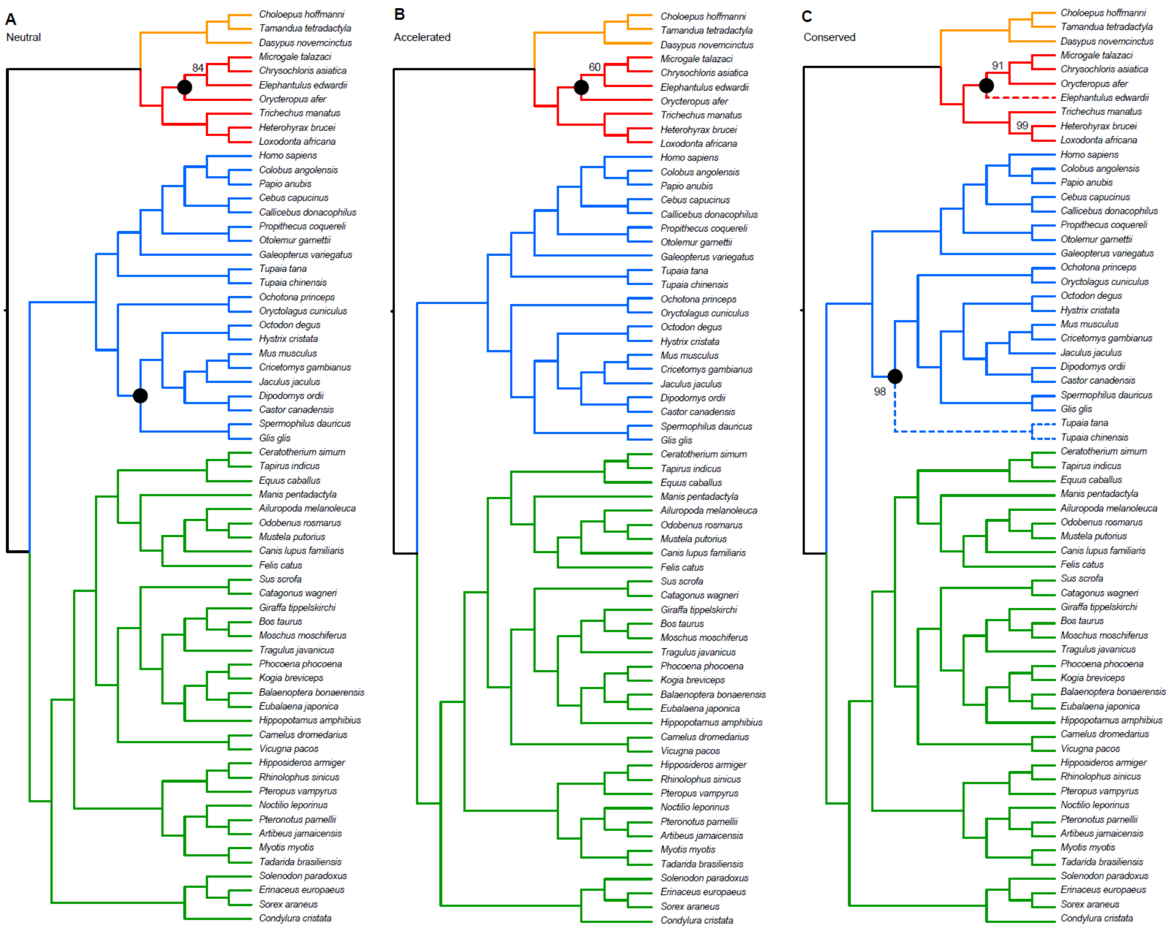

**Fig. S1.**

**Multispecies coalescent phylogenies of Neutral, Conserved and Accelerated PhyloP Sites**. 50% Majority Rule Trees from SVDquartets analysis of (**A**) 400,702 neutral, (**B**) 388,684 accelerated and (**C**) 408,790 conserved SNPs from the human referenced alignment (HRA) of 65 species. Bootstrap support is 100% unless otherwise indicated. Black nodes indicate nodes which differ in concatenated ML trees of the same dataset. Where alternative arrangements are indicated in Rodentia, concatenated ML trees support Sciuromorpha + Hystricomorpha and within Afrotheria (red) Orycteropus + *Elephantulus* is supported. Dashed lines indicate differences among datasets for the coalescent analyses.

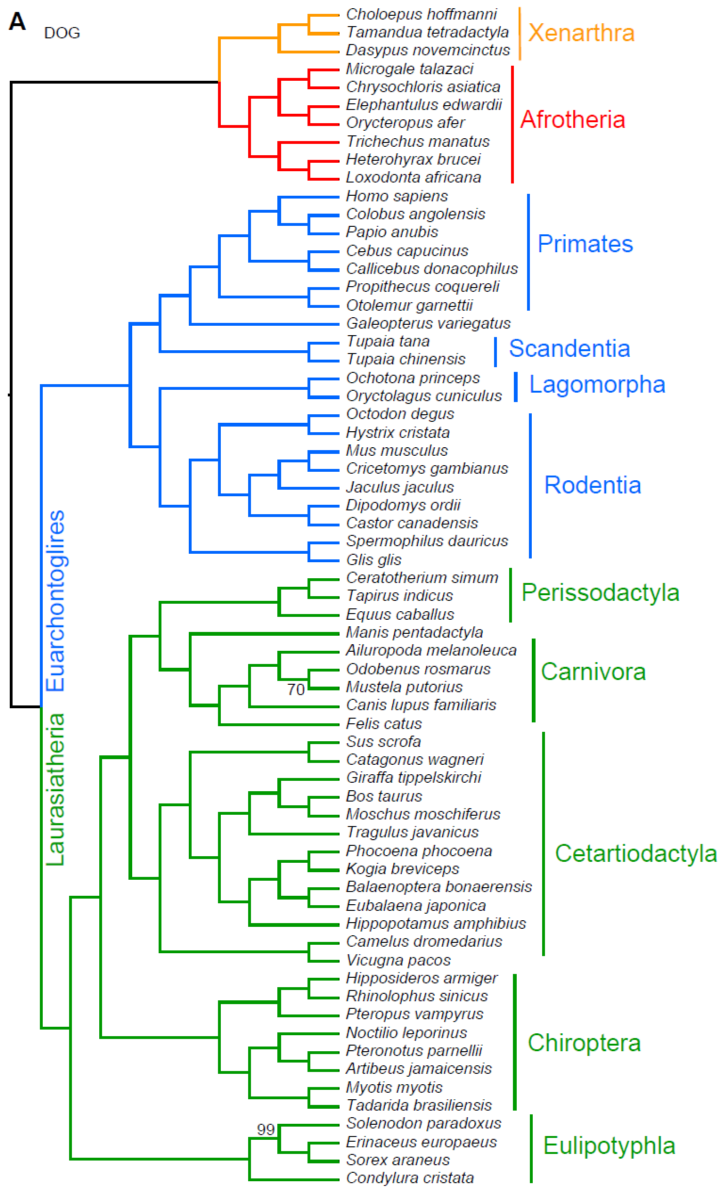

**Fig. S2.**

**Multispecies coalescent phylogenies from differently referenced versions of the HAL alignment with all sites (i.e., no classification based on PhyloP scores).** 50% Majority rule trees from SVDquartets analysis of (**A**) the dog referenced alignment (DRA) and (**B**) the alignment referenced at the inferred ancestor at the root of placental mammals (RRA). Bootstrap support is 100% unless otherwise indicated. Dashed lines indicate differences between the topologies displayed. Black circles indicate nodes which differ in concatenated ML trees of the same dataset. Within Rodentia, the RRA concatenated ML tree supports Sciuromorpha + Hystricomorpha. Within Afrotheria, the RRA concatenated ML trees support *Heterohyrax* + *Loxodonta*.

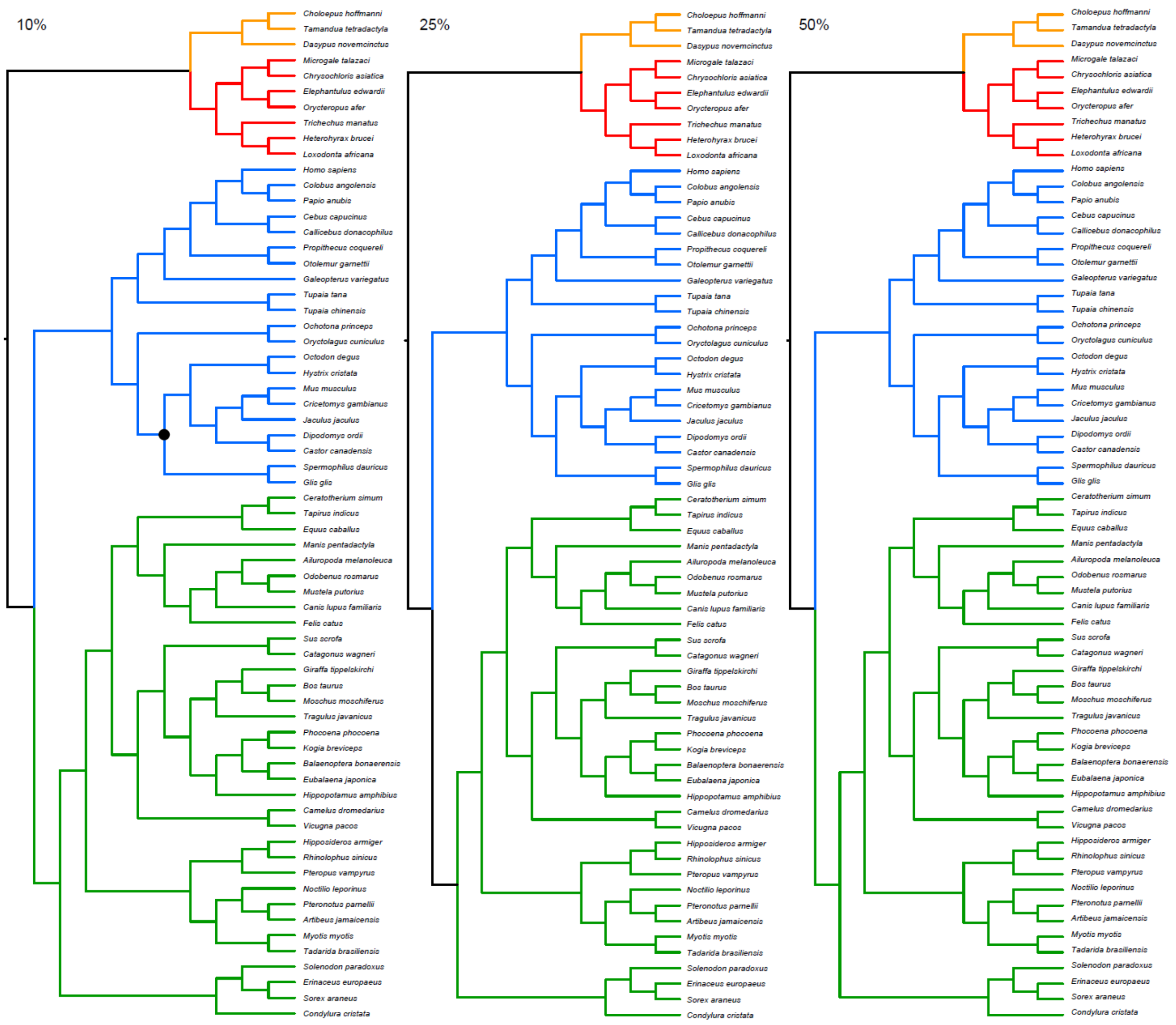

**Fig. S3.**
**Multispecies coalescent phylogenies based on the 65 taxon HRA of all SNPs with variable amounts of missing data**. The black dot in Rodentia indicates where a Sciuromorpha+Hystricomorpha clade is supported by the concatenated ML analysis of the 10% missing tree.

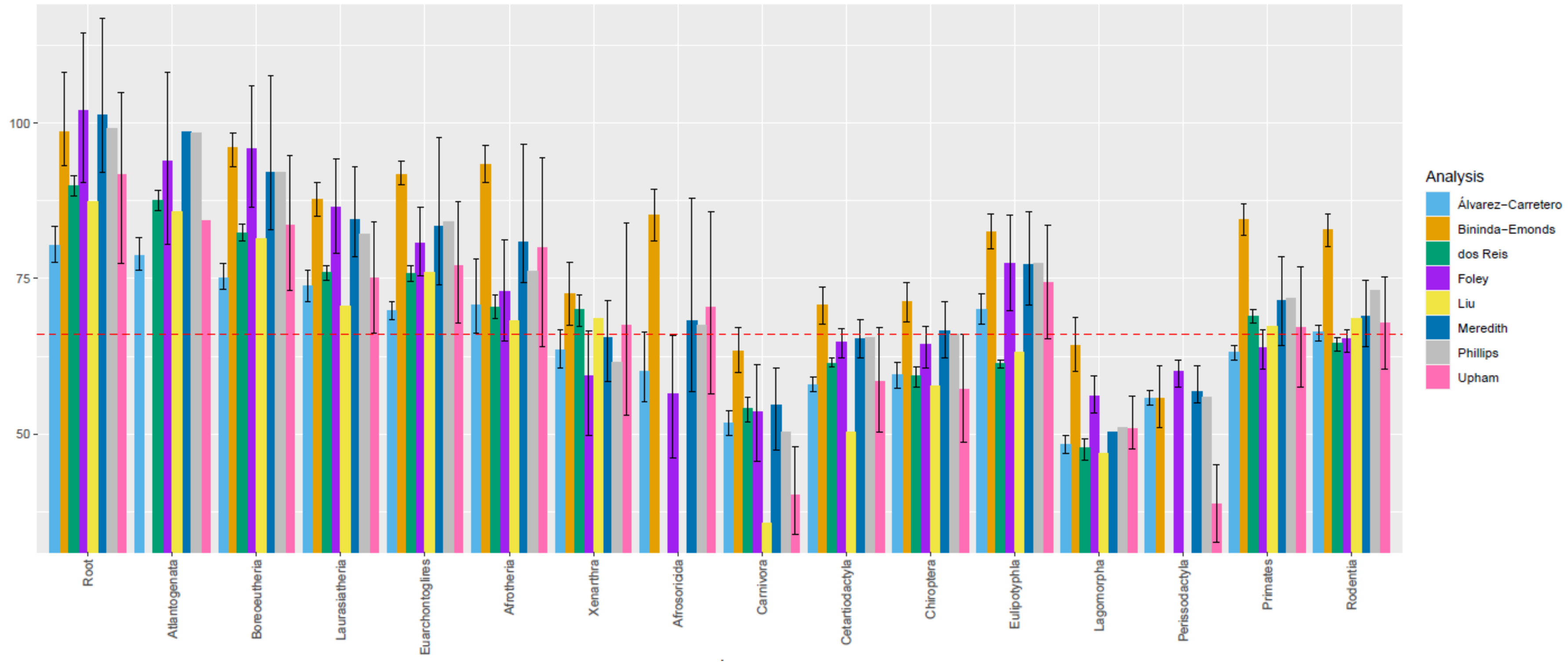

**Fig. S4.**

**Comparison of superordinal and ordinal divergence times**. We display the average of the Benton2009 IRM and ARM results for this study (Foley, purple). We used the Global Mean from Table S6 in Meredith et al. (*6*). We used ages from the Mer82c Posterior Tree from Phillips (*88*). Ages for Bininda-Emonds (*98*), Upham (*32*) and dos Reis (*7*) are from Table S3 in (*32*). Where nodes of interest were omitted from this table for the Upham dataset we supplemented it with ages from the associated VertLife database (*32*). Ages for the Liu et al. are from their supplementary data 9 (column F)(*8*). We used ages from the 4705sp_mean tree from Álvarez-Carretero et al. (*10*). 95% confidence intervals are shown where data was available.

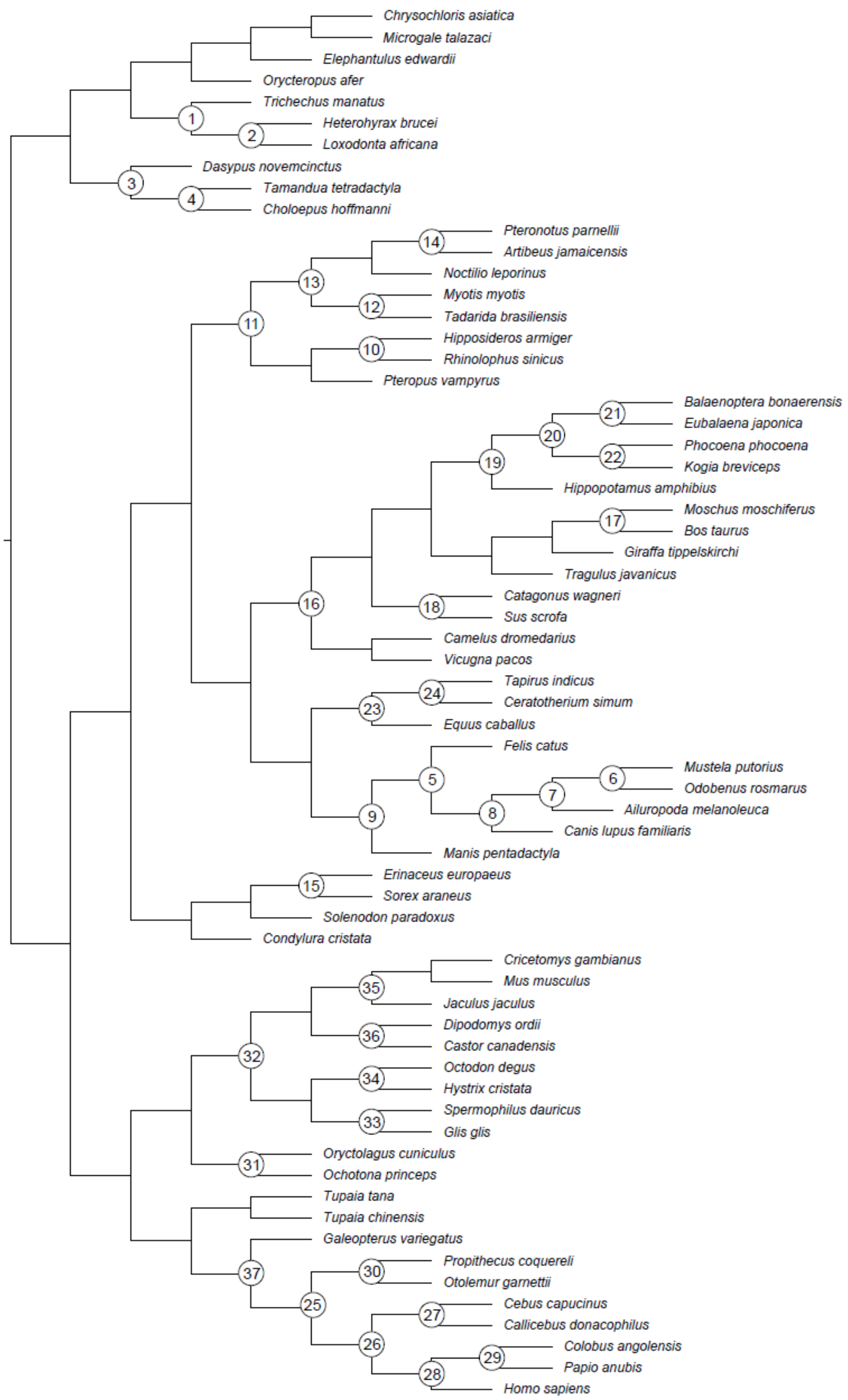

**Fig. S5.**

**Placement of fossil calibrations for the 65 species analysis.** Numbers at nodes correspond to constraints listed in table S9.

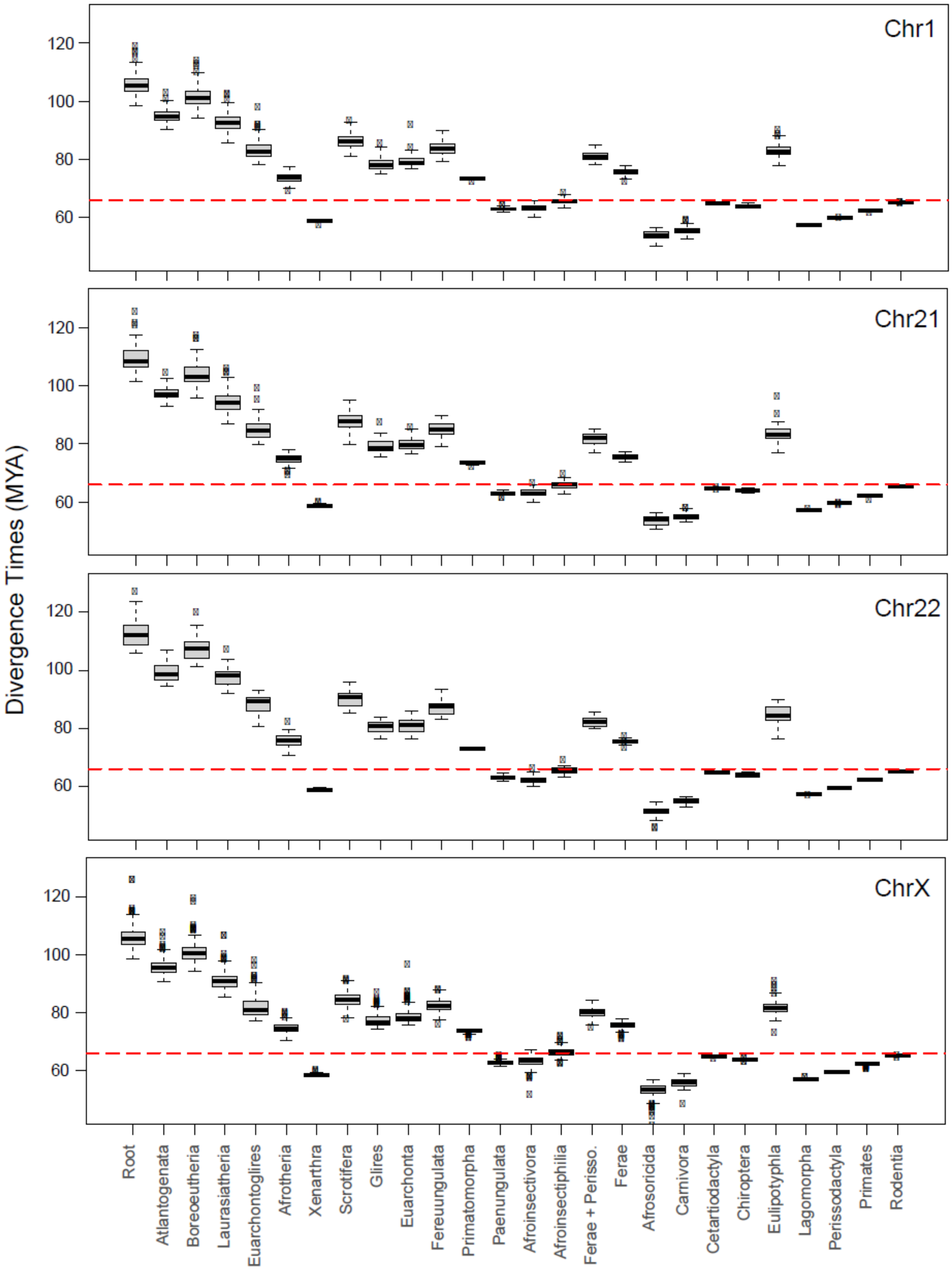

**Fig. S6.**

**Per chromosome divergence time estimates**. 100Kb sliding windows per chromosome using 131.5Mya Benton2009 constraint on the root. The red dashed line indicates the KPg boundary.

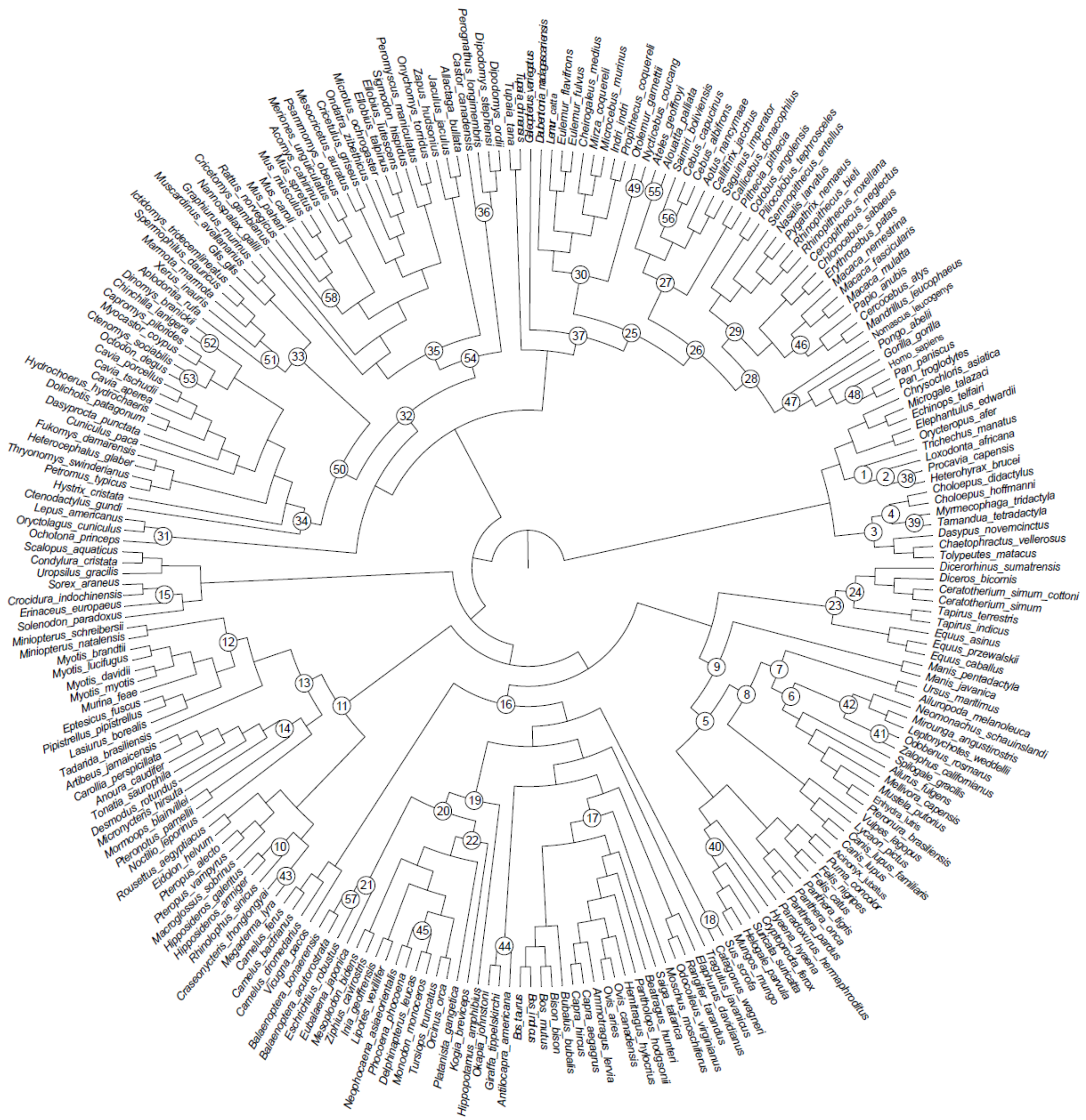

**Fig. S7.**

**Placement of fossil calibrations for the 241 species analysis.** Numbers at nodes correspond to constraints listed in table S9.

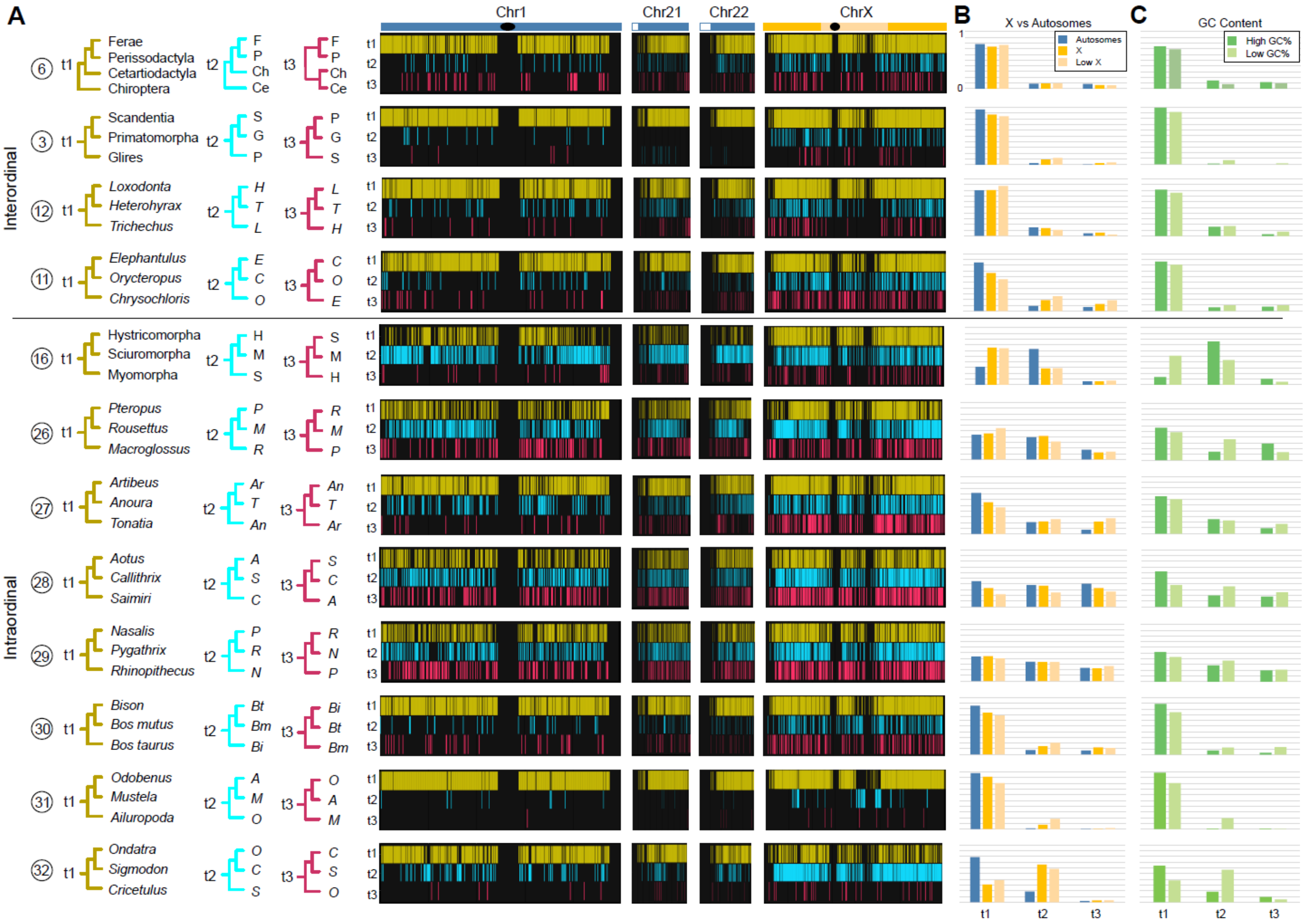

**Fig. S8.**

**Alternately colored version of Figure 2.**

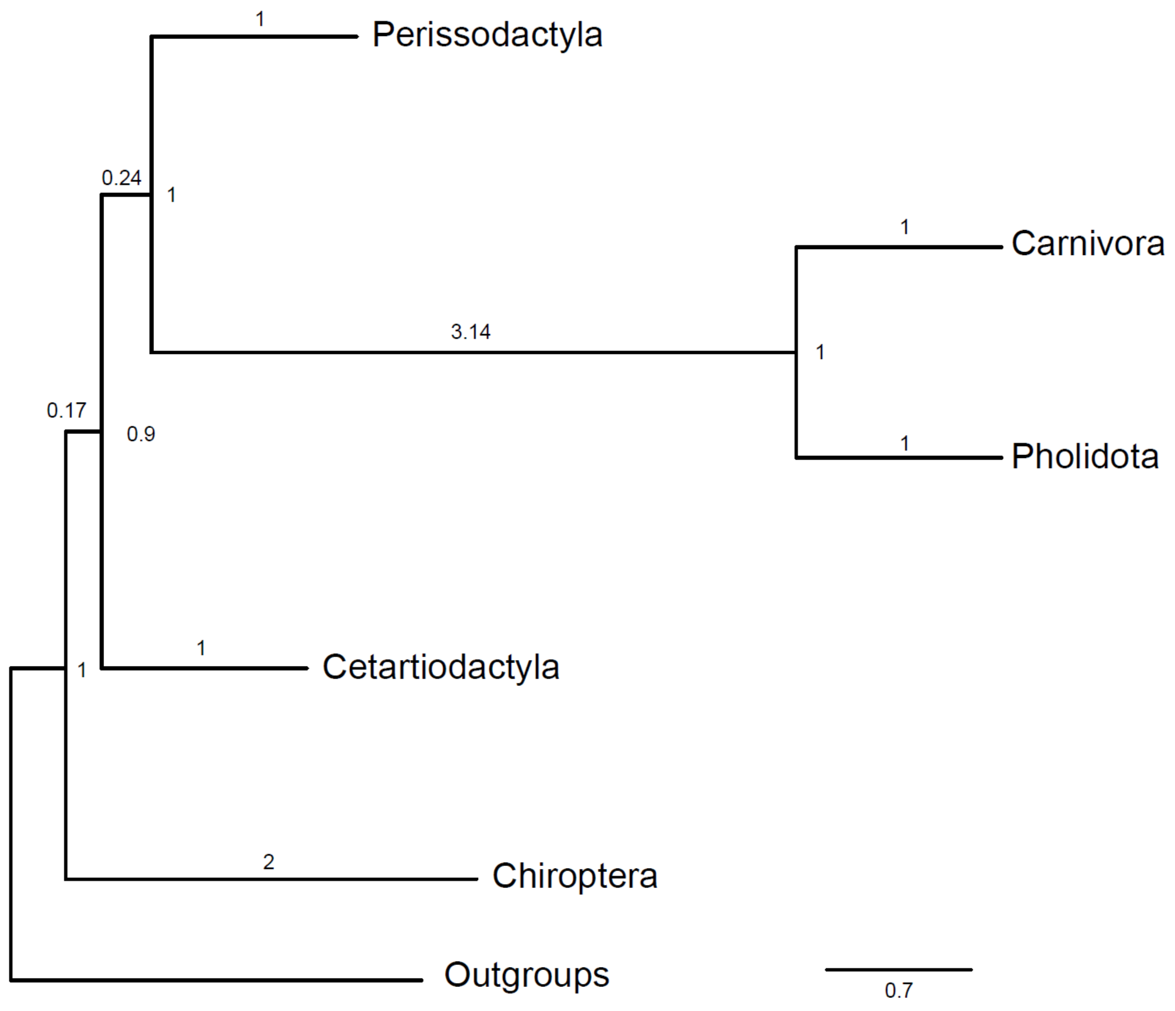

**Fig. S9.**

**ASTRAL_BP tree topology based on counts of deletions which overlap between the analyses of the HRA and RRA.** Local posterior probabilities are shown at nodes and branch lengths in coalescent units are shown on branches.

**Table S1.**

Species composition for phylogenomic datasets and missingness post-filtering.

| Species | Magnorder | Superorder | Order | Family | 65 species Dataset | 228 Species Dataset | Average Per Alignment Missingness Post Filtering for Sliding Windows Analysis |
|---|---|---|---|---|---|---|---|
| *Chrysochloris asiatica* | Atlantogenata | Afrotheria | Afrosoricida | Chrysochloridae | TRUE | TRUE | 41.5 |
| *Echinops telfairi* | Atlantogenata | Afrotheria | Afrosoricida | Tenrecidae | FALSE | TRUE | 44.4 |
| *Microgale talazaci* | Atlantogenata | Afrotheria | Afrosoricida | Tenrecidae | TRUE | TRUE | 44.0 |
| *Heterohyrax brucei* | Atlantogenata | Afrotheria | Hyracoidea | Procaviidae | TRUE | TRUE | 28.0 |
| *Procavia capensis* | Atlantogenata | Afrotheria | Hyracoidea | Procaviidae | FALSE | TRUE | 28.1 |
| *Elephantulus edwardii* | Atlantogenata | Afrotheria | Macroscelidea | Macroscelididae | TRUE | TRUE | 47.7 |
| *Loxodonta africana* | Atlantogenata | Afrotheria | Proboscidea | Elephantidae | TRUE | TRUE | 21.9 |
| *Trichechus manatus* | Atlantogenata | Afrotheria | Sirenia | Trichechidae | TRUE | TRUE | 22.5 |
| *Orycteropus afer* | Atlantogenata | Afrotheria | Tubulidentata | Orycteropodidae | TRUE | TRUE | 27.0 |
| *Chaetophractus vellerosus* | Atlantogenata | Xenarthra | Cingulata | Dasypodidae | FALSE | FALSE | |
| *Dasypus novemcinctus* | Atlantogenata | Xenarthra | Cingulata | Dasypodidae | TRUE | TRUE | 24.6 |
| *Tolypeutes matacus* | Atlantogenata | Xenarthra | Cingulata | Dasypodidae | FALSE | TRUE | 23.4 |
| *Choloepus didactylus* | Atlantogenata | Xenarthra | Pilosa | Megalonychidae | FALSE | FALSE | |
| *Choloepus hoffmanni* | Atlantogenata | Xenarthra | Pilosa | Megalonychidae | TRUE | TRUE | 20.9 |
| *Myrmecophaga tridactyla* | Atlantogenata | Xenarthra | Pilosa | Myrmecophagidae | FALSE | TRUE | 22.2 |
| *Tamandua tetradactyla* | Atlantogenata | Xenarthra | Pilosa | Myrmecophagidae | TRUE | TRUE | 22.4 |
| *Galeopterus variegatus* | Boreoeutheria | Euarchontoglires | Dermoptera | Cynocephalidae | TRUE | TRUE | 6.8 |
| *Lepus americanus* | Boreoeutheria | Euarchontoglires | Lagomorpha | Leporidae | FALSE | TRUE | 28.1 |
| *Oryctolagus cuniculus* | Boreoeutheria | Euarchontoglires | Lagomorpha | Leporidae | TRUE | TRUE | 29.2 |
| *Ochotona princeps* | Boreoeutheria | Euarchontoglires | Lagomorpha | Ochotonidae | TRUE | TRUE | 37.4 |
| *Aotus nancymaae* | Boreoeutheria | Euarchontoglires | Primates | Aotidae | FALSE | TRUE | 7.6 |
| *Alouatta palliata* | Boreoeutheria | Euarchontoglires | Primates | Atelidae | FALSE | TRUE | 6.9 |
| *Ateles geoffroyi* | Boreoeutheria | Euarchontoglires | Primates | Atelidae | FALSE | TRUE | 6.5 |
| *Callithrix jacchus* | Boreoeutheria | Euarchontoglires | Primates | Cebidae | FALSE | TRUE | 7.6 |
| *Cebus albifrons* | Boreoeutheria | Euarchontoglires | Primates | Cebidae | FALSE | TRUE | 9.4 |
| *Cebus capucinus* | Boreoeutheria | Euarchontoglires | Primates | Cebidae | TRUE | TRUE | 7.6 |
| *Saguinus imperator* | Boreoeutheria | Euarchontoglires | Primates | Cebidae | FALSE | TRUE | 7.8 |
| *Saimiri boliviensis* | Boreoeutheria | Euarchontoglires | Primates | Cebidae | FALSE | TRUE | 9.9 |

| Species | Magnorder | Superorder | Order | Family | 65 species Dataset | 228 Species Dataset | Average Per Alignment Missingness Post Filtering for Sliding Windows Analysis |
|---|---|---|---|---|---|---|---|
| *Cercocebus atys* | Boreoeutheria | Euarchontoglires | Primates | Cercopithecidae | FALSE | TRUE | 3.3 |
| *Cercopithecus neglectus* | Boreoeutheria | Euarchontoglires | Primates | Cercopithecidae | FALSE | FALSE | |
| *Chlorocebus sabaeus* | Boreoeutheria | Euarchontoglires | Primates | Cercopithecidae | FALSE | TRUE | 3.2 |
| *Colobus angolensis* | Boreoeutheria | Euarchontoglires | Primates | Cercopithecidae | TRUE | TRUE | 4.9 |
| *Erythrocebus patas* | Boreoeutheria | Euarchontoglires | Primates | Cercopithecidae | FALSE | TRUE | 3.7 |
| *Macaca fascicularis* | Boreoeutheria | Euarchontoglires | Primates | Cercopithecidae | FALSE | TRUE | 3.0 |
| *Macaca mulatta* | Boreoeutheria | Euarchontoglires | Primates | Cercopithecidae | FALSE | TRUE | 3.1 |
| *Macaca nemestrina* | Boreoeutheria | Euarchontoglires | Primates | Cercopithecidae | FALSE | TRUE | 3.0 |
| *Mandrillus leucophaeus* | Boreoeutheria | Euarchontoglires | Primates | Cercopithecidae | FALSE | TRUE | 4.1 |
| *Nasalis larvatus* | Boreoeutheria | Euarchontoglires | Primates | Cercopithecidae | FALSE | TRUE | 3.3 |
| *Papio anubis* | Boreoeutheria | Euarchontoglires | Primates | Cercopithecidae | TRUE | TRUE | 3.2 |
| *Piliocolobus tephrosceles* | Boreoeutheria | Euarchontoglires | Primates | Cercopithecidae | FALSE | TRUE | 4.0 |
| *Pygathrix nemaeus* | Boreoeutheria | Euarchontoglires | Primates | Cercopithecidae | FALSE | TRUE | 3.2 |
| *Rhinopithecus bieti* | Boreoeutheria | Euarchontoglires | Primates | Cercopithecidae | FALSE | TRUE | 5.4 |
| *Rhinopithecus roxellana* | Boreoeutheria | Euarchontoglires | Primates | Cercopithecidae | FALSE | TRUE | 3.4 |
| *Semnopithecus entellus* | Boreoeutheria | Euarchontoglires | Primates | Cercopithecidae | FALSE | TRUE | 5.1 |
| *Cheirogaleus medius* | Boreoeutheria | Euarchontoglires | Primates | Cheirogaleidae | FALSE | TRUE | 10.5 |
| *Microcebus murinus* | Boreoeutheria | Euarchontoglires | Primates | Cheirogaleidae | FALSE | TRUE | 12.1 |
| *Mirza coquereli* | Boreoeutheria | Euarchontoglires | Primates | Cheirogaleidae | FALSE | TRUE | 11.3 |
| *Daubentonia madagascariensis* | Boreoeutheria | Euarchontoglires | Primates | Daubentoniidae | FALSE | TRUE | 6.4 |
| *Otolemur garnettii* | Boreoeutheria | Euarchontoglires | Primates | Galagidae | TRUE | TRUE | 17.6 |
| *Gorilla gorilla* | Boreoeutheria | Euarchontoglires | Primates | Hominidae | FALSE | TRUE | 1.2 |
| *Homo sapiens* | Boreoeutheria | Euarchontoglires | Primates | Hominidae | TRUE | TRUE | 0.0 |
| *Pan paniscus* | Boreoeutheria | Euarchontoglires | Primates | Hominidae | FALSE | TRUE | 3.9 |
| *Pan troglodytes* | Boreoeutheria | Euarchontoglires | Primates | Hominidae | FALSE | TRUE | 2.0 |
| *Pongo abelii* | Boreoeutheria | Euarchontoglires | Primates | Hominidae | FALSE | TRUE | 1.3 |
| *Nomascus leucogenys* | Boreoeutheria | Euarchontoglires | Primates | Hylobatidae | FALSE | TRUE | 4.9 |
| *Indri indri* | Boreoeutheria | Euarchontoglires | Primates | Indridae | FALSE | TRUE | 10.0 |
| *Propithecus coquereli* | Boreoeutheria | Euarchontoglires | Primates | Indriidae | TRUE | TRUE | 13.8 |
| *Eulemur flavifrons* | Boreoeutheria | Euarchontoglires | Primates | Lemuridae | FALSE | TRUE | 11.4 |

| Species | Magnorder | Superorder | Order | Family | 65 species Dataset | 228 Species Dataset | Average Per Alignment Missingness Post Filtering for Sliding Windows Analysis |
|---|---|---|---|---|---|---|---|
| *Eulemur fulvus* | Boreoeutheria | Euarchontoglires | Primates | Lemuridae | FALSE | TRUE | 10.1 |
| *Lemur catta* | Boreoeutheria | Euarchontoglires | Primates | Lemuridae | FALSE | TRUE | 10.8 |
| *Nycticebus coucang* | Boreoeutheria | Euarchontoglires | Primates | Lorisidae | FALSE | TRUE | 18.6 |
| *Callicebus donacophilus* | Boreoeutheria | Euarchontoglires | Primates | Pitheciidae | TRUE | TRUE | 6.6 |
| *Pithecia pithecia* | Boreoeutheria | Euarchontoglires | Primates | Pitheciidae | FALSE | TRUE | 6.0 |
| *Aplodontia rufa* | Boreoeutheria | Euarchontoglires | Rodentia | Aplodontiidae | FALSE | TRUE | 23.9 |
| *Fukomys damarensis* | Boreoeutheria | Euarchontoglires | Rodentia | Bathyergidae | FALSE | TRUE | 30.4 |
| *Heterocephalus glaber* | Boreoeutheria | Euarchontoglires | Rodentia | Bathyergidae | FALSE | TRUE | 28.3 |
| *Capromys pilorides* | Boreoeutheria | Euarchontoglires | Rodentia | Capromyidae | FALSE | TRUE | 36.7 |
| *Castor canadensis* | Boreoeutheria | Euarchontoglires | Rodentia | Castoridae | TRUE | TRUE | 23.9 |
| *Cavia aperea* | Boreoeutheria | Euarchontoglires | Rodentia | Caviidae | FALSE | FALSE | |
| *Cavia porcellus* | Boreoeutheria | Euarchontoglires | Rodentia | Caviidae | FALSE | TRUE | 29.9 |
| *Cavia tschudii* | Boreoeutheria | Euarchontoglires | Rodentia | Caviidae | FALSE | TRUE | 28.9 |
| *Dolichotis patagonum* | Boreoeutheria | Euarchontoglires | Rodentia | Caviidae | FALSE | TRUE | 29.0 |
| *Hydrochoerus hydrochaeris* | Boreoeutheria | Euarchontoglires | Rodentia | Caviidae | FALSE | TRUE | 28.9 |
| *Chinchilla lanigera* | Boreoeutheria | Euarchontoglires | Rodentia | Chinchillidae | FALSE | TRUE | 26.1 |
| *Cricetulus griseus* | Boreoeutheria | Euarchontoglires | Rodentia | Cricetidae | FALSE | TRUE | 40.5 |
| *Ellobius lutescens* | Boreoeutheria | Euarchontoglires | Rodentia | Cricetidae | FALSE | TRUE | 42.7 |
| *Ellobius talpinus* | Boreoeutheria | Euarchontoglires | Rodentia | Cricetidae | FALSE | TRUE | 40.7 |
| *Mesocricetus auratus* | Boreoeutheria | Euarchontoglires | Rodentia | Cricetidae | FALSE | TRUE | 43.3 |
| *Microtus ochrogaster* | Boreoeutheria | Euarchontoglires | Rodentia | Cricetidae | FALSE | TRUE | 41.3 |
| *Ondatra zibethicus* | Boreoeutheria | Euarchontoglires | Rodentia | Cricetidae | FALSE | TRUE | 39.9 |
| *Onychomys torridus* | Boreoeutheria | Euarchontoglires | Rodentia | Cricetidae | FALSE | TRUE | 38.4 |
| *Peromyscus maniculatus* | Boreoeutheria | Euarchontoglires | Rodentia | Cricetidae | FALSE | TRUE | 37.3 |
| *Sigmodon hispidus* | Boreoeutheria | Euarchontoglires | Rodentia | Cricetidae | FALSE | TRUE | 41.2 |
| *Ctenodactylus gundi* | Boreoeutheria | Euarchontoglires | Rodentia | Ctenodactylidae | FALSE | TRUE | 27.9 |
| *Ctenomys sociabilis* | Boreoeutheria | Euarchontoglires | Rodentia | Ctenomyidae | FALSE | TRUE | 31.6 |
| *Cuniculus paca* | Boreoeutheria | Euarchontoglires | Rodentia | Cuniculidae | FALSE | TRUE | 34.9 |
| *Dasyprocta punctata* | Boreoeutheria | Euarchontoglires | Rodentia | Dasyproctidae | FALSE | TRUE | 26.3 |
| *Dinomys branickii* | Boreoeutheria | Euarchontoglires | Rodentia | Dinomyidae | FALSE | TRUE | 27.7 |

| Species | Magnorder | Superorder | Order | Family | 65 species Dataset | 228 Species Dataset | Average Per Alignment Missingness Post Filtering for Sliding Windows Analysis |
|---|---|---|---|---|---|---|---|
| *Allactaga bullata* | Boreoeutheria | Euarchontoglires | Rodentia | Dipodidae | FALSE | TRUE | 38.6 |
| *Jaculus jaculus* | Boreoeutheria | Euarchontoglires | Rodentia | Dipodidae | TRUE | TRUE | 40.0 |
| *Zapus hudsonius* | Boreoeutheria | Euarchontoglires | Rodentia | Dipodidae | FALSE | FALSE | |
| *Glis glis* | Boreoeutheria | Euarchontoglires | Rodentia | Gliridae | TRUE | TRUE | 26.0 |
| *Graphiurus murinus* | Boreoeutheria | Euarchontoglires | Rodentia | Gliridae | FALSE | TRUE | 27.7 |
| *Muscardinus avellanarius* | Boreoeutheria | Euarchontoglires | Rodentia | Gliridae | FALSE | TRUE | 29.6 |
| *Dipodomys ordii* | Boreoeutheria | Euarchontoglires | Rodentia | Heteromyidae | TRUE | TRUE | 39.2 |
| *Dipodomys stephensi* | Boreoeutheria | Euarchontoglires | Rodentia | Heteromyidae | FALSE | TRUE | 39.0 |
| *Perognathus longimembris* | Boreoeutheria | Euarchontoglires | Rodentia | Heteromyidae | FALSE | TRUE | 40.0 |
| *Hystrix cristata* | Boreoeutheria | Euarchontoglires | Rodentia | Hystricidae | TRUE | TRUE | 22.6 |
| *Acomys cahirinus* | Boreoeutheria | Euarchontoglires | Rodentia | Muridae | FALSE | TRUE | 41.5 |
| *Meriones unguiculatus* | Boreoeutheria | Euarchontoglires | Rodentia | Muridae | FALSE | FALSE | |
| *Mus caroli* | Boreoeutheria | Euarchontoglires | Rodentia | Muridae | FALSE | TRUE | 41.0 |
| *Mus musculus* | Boreoeutheria | Euarchontoglires | Rodentia | Muridae | TRUE | TRUE | 40.6 |
| *Mus pahari* | Boreoeutheria | Euarchontoglires | Rodentia | Muridae | FALSE | TRUE | 41.2 |
| *Mus spretus* | Boreoeutheria | Euarchontoglires | Rodentia | Muridae | FALSE | TRUE | 40.7 |
| *Psammomys obesus* | Boreoeutheria | Euarchontoglires | Rodentia | Muridae | FALSE | FALSE | |
| *Rattus norvegicus* | Boreoeutheria | Euarchontoglires | Rodentia | Muridae | FALSE | TRUE | 40.7 |
| *Myocastor coypus* | Boreoeutheria | Euarchontoglires | Rodentia | Myocastoridae | FALSE | TRUE | 32.8 |
| *Cricetomys gambianus* | Boreoeutheria | Euarchontoglires | Rodentia | Nesomyidae | TRUE | TRUE | 36.3 |
| *Octodon degus* | Boreoeutheria | Euarchontoglires | Rodentia | Octodontidae | TRUE | TRUE | 30.9 |
| *Petromus typicus* | Boreoeutheria | Euarchontoglires | Rodentia | Petromuridae | FALSE | TRUE | 33.2 |
| *Ictidomys tridecemlineatus* | Boreoeutheria | Euarchontoglires | Rodentia | Sciuridae | FALSE | TRUE | 23.2 |
| *Marmota marmota* | Boreoeutheria | Euarchontoglires | Rodentia | Sciuridae | FALSE | TRUE | 22.3 |
| *Spermophilus dauricus* | Boreoeutheria | Euarchontoglires | Rodentia | Sciuridae | TRUE | TRUE | 22.2 |
| *Xerus inauris* | Boreoeutheria | Euarchontoglires | Rodentia | Sciuridae | FALSE | TRUE | 20.2 |
| *Nannospalax galili* | Boreoeutheria | Euarchontoglires | Rodentia | Spalacidae | FALSE | TRUE | 31.9 |
| *Thryonomys swinderianus* | Boreoeutheria | Euarchontoglires | Rodentia | Thryonomyidae | FALSE | TRUE | 34.4 |
| *Tupaia chinensis* | Boreoeutheria | Euarchontoglires | Scandentia | Tupaiidae | TRUE | TRUE | 21.1 |
| *Tupaia tana* | Boreoeutheria | Euarchontoglires | Scandentia | Tupaiidae | TRUE | TRUE | 27.8 |

| Species | Magnorder | Superorder | Order | Family | 65 species Dataset | 228 Species Dataset | Average Per Alignment Missingness Post Filtering for Sliding Windows Analysis |
|---|---|---|---|---|---|---|---|
| *Ailurus fulgens* | Boreoeutheria | Laurasiatheria | Carnivora | Ailuridae | FALSE | TRUE | 13.1 |
| *Canis lupus* | Boreoeutheria | Laurasiatheria | Carnivora | Canidae | FALSE | TRUE | 11.7 |
| *Canis lupus familiaris* | Boreoeutheria | Laurasiatheria | Carnivora | Canidae | TRUE | TRUE | 12.2 |
| *Lycaon pictus* | Boreoeutheria | Laurasiatheria | Carnivora | Canidae | FALSE | TRUE | 17.3 |
| *Vulpes lagopus* | Boreoeutheria | Laurasiatheria | Carnivora | Canidae | FALSE | TRUE | 11.9 |
| *Cryptoprocta ferox* | Boreoeutheria | Laurasiatheria | Carnivora | Eupleridae | FALSE | TRUE | 12.3 |
| *Acinonyx jubatus* | Boreoeutheria | Laurasiatheria | Carnivora | Felidae | FALSE | TRUE | 13.1 |
| *Felis catus* | Boreoeutheria | Laurasiatheria | Carnivora | Felidae | TRUE | TRUE | 11.6 |
| *Felis nigripes* | Boreoeutheria | Laurasiatheria | Carnivora | Felidae | FALSE | FALSE | |
| *Panthera onca* | Boreoeutheria | Laurasiatheria | Carnivora | Felidae | FALSE | TRUE | 11.2 |
| *Panthera pardus* | Boreoeutheria | Laurasiatheria | Carnivora | Felidae | FALSE | TRUE | 11.0 |
| *Panthera tigris* | Boreoeutheria | Laurasiatheria | Carnivora | Felidae | FALSE | TRUE | 12.7 |
| *Puma concolor* | Boreoeutheria | Laurasiatheria | Carnivora | Felidae | FALSE | TRUE | 13.6 |
| *Helogale parvula* | Boreoeutheria | Laurasiatheria | Carnivora | Herpestidae | FALSE | TRUE | 14.3 |
| *Mungos mungo* | Boreoeutheria | Laurasiatheria | Carnivora | Herpestidae | FALSE | TRUE | 14.3 |
| *Suricata suricatta* | Boreoeutheria | Laurasiatheria | Carnivora | Herpestidae | FALSE | TRUE | 14.5 |
| *Hyaena hyaena* | Boreoeutheria | Laurasiatheria | Carnivora | Hyaenidae | FALSE | TRUE | 12.7 |
| *Spilogale gracilis* | Boreoeutheria | Laurasiatheria | Carnivora | Mephitidae | FALSE | TRUE | 14.3 |
| *Enhydra lutris* | Boreoeutheria | Laurasiatheria | Carnivora | Mustelidae | FALSE | TRUE | 12.7 |
| *Mellivora capensis* | Boreoeutheria | Laurasiatheria | Carnivora | Mustelidae | FALSE | TRUE | 13.0 |
| *Mustela putorius* | Boreoeutheria | Laurasiatheria | Carnivora | Mustelidae | TRUE | TRUE | 14.2 |
| *Pteronura brasiliensis* | Boreoeutheria | Laurasiatheria | Carnivora | Mustelidae | FALSE | TRUE | 12.8 |
| *Odobenus rosmarus* | Boreoeutheria | Laurasiatheria | Carnivora | Odobenidae | TRUE | TRUE | 10.5 |
| *Zalophus californianus* | Boreoeutheria | Laurasiatheria | Carnivora | Otariidae | FALSE | TRUE | 10.5 |
| *Leptonychotes weddellii* | Boreoeutheria | Laurasiatheria | Carnivora | Phocidae | FALSE | TRUE | 12.2 |
| *Mirounga angustirostris* | Boreoeutheria | Laurasiatheria | Carnivora | Phocidae | FALSE | TRUE | 10.9 |
| *Neomonachus schauinslandi* | Boreoeutheria | Laurasiatheria | Carnivora | Phocidae | FALSE | TRUE | 10.9 |
| *Ailuropoda melanoleuca* | Boreoeutheria | Laurasiatheria | Carnivora | Ursidae | TRUE | TRUE | 10.8 |
| *Ursus maritimus* | Boreoeutheria | Laurasiatheria | Carnivora | Ursidae | FALSE | TRUE | 12.2 |
| *Paradoxurus hermaphroditus* | Boreoeutheria | Laurasiatheria | Carnivora | Viverridae | FALSE | TRUE | 11.7 |

| Species | Magnorder | Superorder | Order | Family | 65 species Dataset | 228 Species Dataset | Average Per Alignment Missingness Post Filtering for Sliding Windows Analysis |
|---|---|---|---|---|---|---|---|
| *Antilocapra americana* | Boreoeutheria | Laurasiatheria | Cetartiodactyla | Antilocapridae | FALSE | TRUE | 16.5 |
| *Eubalaena japonica* | Boreoeutheria | Laurasiatheria | Cetartiodactyla | Balaenidae | TRUE | TRUE | 9.6 |
| *Balaenoptera acutorostrata* | Boreoeutheria | Laurasiatheria | Cetartiodactyla | Balaenopteridae | FALSE | TRUE | 10.2 |
| *Balaenoptera bonaerensis* | Boreoeutheria | Laurasiatheria | Cetartiodactyla | Balaenopteridae | TRUE | TRUE | 10.2 |
| *Ammotragus lervia* | Boreoeutheria | Laurasiatheria | Cetartiodactyla | Bovidae | FALSE | TRUE | 16.8 |
| *Beatragus hunteri* | Boreoeutheria | Laurasiatheria | Cetartiodactyla | Bovidae | FALSE | TRUE | 16.2 |
| *Bison bison* | Boreoeutheria | Laurasiatheria | Cetartiodactyla | Bovidae | FALSE | TRUE | 19.0 |
| *Bos indicus* | Boreoeutheria | Laurasiatheria | Cetartiodactyla | Bovidae | FALSE | FALSE | |
| *Bos mutus* | Boreoeutheria | Laurasiatheria | Cetartiodactyla | Bovidae | FALSE | TRUE | 15.8 |
| *Bos taurus* | Boreoeutheria | Laurasiatheria | Cetartiodactyla | Bovidae | TRUE | TRUE | 16.0 |
| *Bubalus bubalis* | Boreoeutheria | Laurasiatheria | Cetartiodactyla | Bovidae | FALSE | TRUE | 15.8 |
| *Capra aegagrus* | Boreoeutheria | Laurasiatheria | Cetartiodactyla | Bovidae | FALSE | TRUE | 17.2 |
| *Capra hircus* | Boreoeutheria | Laurasiatheria | Cetartiodactyla | Bovidae | FALSE | TRUE | 16.0 |
| *Hemitragus hylocrius* | Boreoeutheria | Laurasiatheria | Cetartiodactyla | Bovidae | FALSE | TRUE | 16.0 |
| *Ovis aries* | Boreoeutheria | Laurasiatheria | Cetartiodactyla | Bovidae | FALSE | TRUE | 16.2 |
| *Ovis canadensis* | Boreoeutheria | Laurasiatheria | Cetartiodactyla | Bovidae | FALSE | TRUE | 17.3 |
| *Pantholops hodgsonii* | Boreoeutheria | Laurasiatheria | Cetartiodactyla | Bovidae | FALSE | TRUE | 16.9 |
| *Saiga tatarica* | Boreoeutheria | Laurasiatheria | Cetartiodactyla | Bovidae | FALSE | FALSE | |
| *Camelus bactrianus* | Boreoeutheria | Laurasiatheria | Cetartiodactyla | Camelidae | FALSE | TRUE | 11.2 |
| *Camelus dromedarius* | Boreoeutheria | Laurasiatheria | Cetartiodactyla | Camelidae | TRUE | TRUE | 11.4 |
| *Camelus ferus* | Boreoeutheria | Laurasiatheria | Cetartiodactyla | Camelidae | FALSE | TRUE | 11.7 |
| *Vicugna pacos* | Boreoeutheria | Laurasiatheria | Cetartiodactyla | Camelidae | TRUE | TRUE | 10.9 |
| *Elaphurus davidianus* | Boreoeutheria | Laurasiatheria | Cetartiodactyla | Cervidae | FALSE | TRUE | 14.4 |
| *Odocoileus virginianus* | Boreoeutheria | Laurasiatheria | Cetartiodactyla | Cervidae | FALSE | TRUE | 15.4 |
| *Rangifer tarandus* | Boreoeutheria | Laurasiatheria | Cetartiodactyla | Cervidae | FALSE | TRUE | 14.5 |
| *Orcinus orca* | Boreoeutheria | Laurasiatheria | Cetartiodactyla | Delphinidae | FALSE | TRUE | 10.9 |
| *Tursiops truncatus* | Boreoeutheria | Laurasiatheria | Cetartiodactyla | Delphinidae | FALSE | TRUE | 15.6 |
| *Eschrichtius robustus* | Boreoeutheria | Laurasiatheria | Cetartiodactyla | Eschrichtiidae | FALSE | TRUE | 9.7 |
| *Giraffa tippelskirchi* | Boreoeutheria | Laurasiatheria | Cetartiodactyla | Giraffidae | TRUE | TRUE | 15.5 |
| *Okapia johnstoni* | Boreoeutheria | Laurasiatheria | Cetartiodactyla | Giraffidae | FALSE | TRUE | 16.0 |

| Species | Magnorder | Superorder | Order | Family | 65 species Dataset | 228 Species Dataset | Average Per Alignment Missingness Post Filtering for Sliding Windows Analysis |
|---|---|---|---|---|---|---|---|
| *Hippopotamus amphibius* | Boreoeutheria | Laurasiatheria | Cetartiodactyla | Hippopotamidae | TRUE | TRUE | 8.5 |
| *Inia geoffrensis* | Boreoeutheria | Laurasiatheria | Cetartiodactyla | Iniidae | FALSE | TRUE | 15.7 |
| *Lipotes vexillifer* | Boreoeutheria | Laurasiatheria | Cetartiodactyla | Lipotidae | FALSE | TRUE | 11.0 |
| *Delphinapterus leucas* | Boreoeutheria | Laurasiatheria | Cetartiodactyla | Monodontidae | FALSE | TRUE | 10.7 |
| *Monodon monoceros* | Boreoeutheria | Laurasiatheria | Cetartiodactyla | Monodontidae | FALSE | TRUE | 13.1 |
| *Moschus moschiferus* | Boreoeutheria | Laurasiatheria | Cetartiodactyla | Moschidae | TRUE | TRUE | 16.0 |
| *Neophocaena asiaeorientalis* | Boreoeutheria | Laurasiatheria | Cetartiodactyla | Phocoenidae | FALSE | TRUE | 11.4 |
| *Phocoena phocoena* | Boreoeutheria | Laurasiatheria | Cetartiodactyla | Phocoenidae | TRUE | TRUE | 11.1 |
| *Kogia breviceps* | Boreoeutheria | Laurasiatheria | Cetartiodactyla | Physeteridae | TRUE | TRUE | 11.3 |
| *Platanista gangetica* | Boreoeutheria | Laurasiatheria | Cetartiodactyla | Platanistidae | FALSE | TRUE | 11.7 |
| *Sus scrofa* | Boreoeutheria | Laurasiatheria | Cetartiodactyla | Suidae | TRUE | TRUE | 19.9 |
| *Catagonus wagneri* | Boreoeutheria | Laurasiatheria | Cetartiodactyla | Tayassuidae | TRUE | TRUE | 15.9 |
| *Tragulus javanicus* | Boreoeutheria | Laurasiatheria | Cetartiodactyla | Tragulidae | TRUE | TRUE | 19.8 |
| *Mesoplodon bidens* | Boreoeutheria | Laurasiatheria | Cetartiodactyla | Ziphiidae | FALSE | TRUE | 13.0 |
| *Ziphius cavirostris* | Boreoeutheria | Laurasiatheria | Cetartiodactyla | Ziphiidae | FALSE | FALSE | |
| *Craseonycteris thonglongyai* | Boreoeutheria | Laurasiatheria | Chiroptera | Craseonycteridae | FALSE | TRUE | 31.3 |
| *Hipposideros armiger* | Boreoeutheria | Laurasiatheria | Chiroptera | Hipposideridae | TRUE | TRUE | 20.3 |
| *Hipposideros galeritus* | Boreoeutheria | Laurasiatheria | Chiroptera | Hipposideridae | FALSE | TRUE | 19.0 |
| *Megaderma lyra* | Boreoeutheria | Laurasiatheria | Chiroptera | Megadermatidae | FALSE | TRUE | 23.9 |
| *Tadarida brasiliensis* | Boreoeutheria | Laurasiatheria | Chiroptera | Molossidae | TRUE | TRUE | 19.7 |
| *Mormoops blainvillei* | Boreoeutheria | Laurasiatheria | Chiroptera | Mormoopidae | FALSE | TRUE | 20.9 |
| *Pteronotus parnellii* | Boreoeutheria | Laurasiatheria | Chiroptera | Mormoopidae | TRUE | TRUE | 27.1 |
| *Noctilio leporinus* | Boreoeutheria | Laurasiatheria | Chiroptera | Noctilionidae | TRUE | TRUE | 26.7 |
| *Anoura caudifer* | Boreoeutheria | Laurasiatheria | Chiroptera | Phyllostomidae | FALSE | TRUE | 23.2 |
| *Artibeus jamaicensis* | Boreoeutheria | Laurasiatheria | Chiroptera | Phyllostomidae | TRUE | TRUE | 25.7 |
| *Carollia perspicillata* | Boreoeutheria | Laurasiatheria | Chiroptera | Phyllostomidae | FALSE | TRUE | 24.7 |
| *Desmodus rotundus* | Boreoeutheria | Laurasiatheria | Chiroptera | Phyllostomidae | FALSE | TRUE | 23.6 |
| *Micronycteris hirsuta* | Boreoeutheria | Laurasiatheria | Chiroptera | Phyllostomidae | FALSE | TRUE | 24.6 |
| *Tonatia saurophila* | Boreoeutheria | Laurasiatheria | Chiroptera | Phyllostomidae | FALSE | TRUE | 24.2 |
| *Eidolon helvum* | Boreoeutheria | Laurasiatheria | Chiroptera | Pteropodidae | FALSE | TRUE | 19.8 |

| Species | Magnorder | Superorder | Order | Family | 65 species Dataset | 228 Species Dataset | Average Per Alignment Missingness Post Filtering for Sliding Windows Analysis |
|---|---|---|---|---|---|---|---|
| *Macroglossus sobrinus* | Boreoeutheria | Laurasiatheria | Chiroptera | Pteropodidae | FALSE | TRUE | 19.4 |
| *Pteropus alecto* | Boreoeutheria | Laurasiatheria | Chiroptera | Pteropodidae | FALSE | TRUE | 16.9 |
| *Pteropus vampyrus* | Boreoeutheria | Laurasiatheria | Chiroptera | Pteropodidae | TRUE | TRUE | 18.0 |
| *Rousettus aegyptiacus* | Boreoeutheria | Laurasiatheria | Chiroptera | Pteropodidae | FALSE | TRUE | 18.7 |
| *Rhinolophus sinicus* | Boreoeutheria | Laurasiatheria | Chiroptera | Rhinolophidae | TRUE | TRUE | 18.7 |
| *Eptesicus fuscus* | Boreoeutheria | Laurasiatheria | Chiroptera | Vespertilionidae | FALSE | TRUE | 32.4 |
| *Lasiurus borealis* | Boreoeutheria | Laurasiatheria | Chiroptera | Vespertilionidae | FALSE | TRUE | 35.5 |
| *Miniopterus natalensis* | Boreoeutheria | Laurasiatheria | Chiroptera | Vespertilionidae | FALSE | TRUE | 25.8 |
| *Miniopterus schreibersii* | Boreoeutheria | Laurasiatheria | Chiroptera | Vespertilionidae | FALSE | TRUE | 25.3 |
| *Murina feae* | Boreoeutheria | Laurasiatheria | Chiroptera | Vespertilionidae | FALSE | TRUE | 31.7 |
| *Myotis brandtii* | Boreoeutheria | Laurasiatheria | Chiroptera | Vespertilionidae | FALSE | TRUE | 31.3 |
| *Myotis davidii* | Boreoeutheria | Laurasiatheria | Chiroptera | Vespertilionidae | FALSE | TRUE | 31.7 |
| *Myotis lucifugus* | Boreoeutheria | Laurasiatheria | Chiroptera | Vespertilionidae | FALSE | TRUE | 33.4 |
| *Myotis myotis* | Boreoeutheria | Laurasiatheria | Chiroptera | Vespertilionidae | TRUE | TRUE | 30.5 |
| *Pipistrellus pipistrellus* | Boreoeutheria | Laurasiatheria | Chiroptera | Vespertilionidae | FALSE | FALSE | |
| *Erinaceus europaeus* | Boreoeutheria | Laurasiatheria | Eulipotyphla | Erinaceidae | TRUE | TRUE | 53.5 |
| *Solenodon paradoxus* | Boreoeutheria | Laurasiatheria | Eulipotyphla | Solenodontidae | TRUE | TRUE | 19.6 |
| *Crocidura indochinensis* | Boreoeutheria | Laurasiatheria | Eulipotyphla | Soricidae | FALSE | FALSE | |
| *Sorex araneus* | Boreoeutheria | Laurasiatheria | Eulipotyphla | Soricidae | TRUE | TRUE | 52.7 |
| *Condylura cristata* | Boreoeutheria | Laurasiatheria | Eulipotyphla | Talpidae | TRUE | TRUE | |
| *Scalopus aquaticus* | Boreoeutheria | Laurasiatheria | Eulipotyphla | Talpidae | FALSE | TRUE | 32.0 |
| *Uropsilus gracilis* | Boreoeutheria | Laurasiatheria | Eulipotyphla | Talpidae | FALSE | TRUE | 33.6 |
| *Equus asinus* | Boreoeutheria | Laurasiatheria | Perissodactyla | Equidae | FALSE | TRUE | 5.9 |
| *Equus caballus* | Boreoeutheria | Laurasiatheria | Perissodactyla | Equidae | TRUE | TRUE | 6.8 |
| *Equus przewalskii* | Boreoeutheria | Laurasiatheria | Perissodactyla | Equidae | FALSE | TRUE | 7.1 |
| *Ceratotherium simum* | Boreoeutheria | Laurasiatheria | Perissodactyla | Rhinocerotidae | TRUE | TRUE | 5.5 |
| *Ceratotherium simum cottoni* | Boreoeutheria | Laurasiatheria | Perissodactyla | Rhinocerotidae | FALSE | TRUE | 5.5 |
| *Dicerorhinus sumatrensis* | Boreoeutheria | Laurasiatheria | Perissodactyla | Rhinocerotidae | FALSE | TRUE | 5.3 |
| *Diceros bicornis* | Boreoeutheria | Laurasiatheria | Perissodactyla | Rhinocerotidae | FALSE | TRUE | 4.9 |
| *Tapirus indicus* | Boreoeutheria | Laurasiatheria | Perissodactyla | Tapiridae | TRUE | TRUE | 5.2 |

| Species | Magnorder | Superorder | Order | Family | 65 species Dataset | 228 Species Dataset | Average Per Alignment Missingness Post Filtering for Sliding Windows Analysis |
|---|---|---|---|---|---|---|---|
| *Tapirus terrestris* | Boreoeutheria | Laurasiatheria | Perissodactyla | Tapiridae | FALSE | TRUE | 5.2 |
| *Manis javanica* | Boreoeutheria | Laurasiatheria | Pholidota | Manidae | FALSE | TRUE | 14.0 |
| *Manis pentadactyla* | Boreoeutheria | Laurasiatheria | Pholidota | Manidae | TRUE | TRUE | 20.9 |

**Table S2.**

Summary of datasets used for coalescence and concatenation based phylogenomic analysis using SVDquartets and IQ-TREE 2, respectively. Analyses are based on the human referenced alignment (HRA), the dog referenced alignment (DRA) or the inferred ancestor at the root referenced alignment (RRA).

| Dataset | Analysis | Alignment Reference | No. of species | Total SNPs | Data | Missingness | Sampled Quartets | Incompatible Quartets (%) | Compatible Quartets (%) |
|---|---|---|---|---|---|---|---|---|---|
| **HRA** | Coalescence & Concatenation | HRA | 65 | 232,123 | whole genome | 10% | 99% | 5.10% | 94.90% |
| **HRA** | Coalescence & Concatenation | HRA | 65 | 546,102 | whole genome | 25% | 99% | 3.79% | 96.21% |
| **HRA** | Coalescence & Concatenation | HRA | 65 | 796,581 | whole genome | 50% | 99% | 3.52% | 96.48% |
| **DRA** | Coalescence & Concatenation | DRA | 65 | 404,628 | whole genome | 10% | 99% | 3.96% | 96.05% |
| **RRA** | Coalescence & Concatenation | RRA | 65 | 396,620 | whole genome | 10% | 99% | 4.42% | 95.58% |
| **Conserved** | Coalescence & Concatenation | HRA | 65 | 408,790 | Sites within Conserved PhyloP Quintile | 10% | 99% | 4.71% | 95.29% |
| **Accelerated** | Coalescence & Concatenation | HRA | 65 | 388,684 | Sites within Accelerated PhyloP Quintile | 10% | 99% | 3.70% | 96.30% |
| **Neutral** | Coalescence & Concatenation | HRA | 65 | 400,702 | Sites within Neutral PhyloP Quintile | 10% | 99% | 3.66% | 96.34% |
| **Neutral** | Coalescence & Concatenation | HRA | 241 | 466,232 | Sites within Neutral PhyloP Quintile | 10% | All | 3.21% | 96.79% |

**Table S3.**
Summary of phylogenomic relationships for select clades across all analyses.

| | Coalescence | | | | | | | Concatenation | | | | | | | | | |
|---|---|---|---|---|---|---|---|---|---|---|---|---|---|---|---|---|---|
| | HRA - 65 | DRA - 65 | RRA - 65 | Conserved – HRA - 65 | Accelerated – HRA - 65 | Neutral – HRA - 65 | Neutral – HRA - 241 | HRA - 65 | DRA - 65 | RRA - 65 | Conserved – HRA - 65 | Accelerated – HRA - 65 | Neutral – HRA - 65 | Neutral -HRA - 241 | Sliding Windows - X | Sliding Windows - Autosomes | Rare Genomic Changes |
| **Laurasiatheria** | | | | | | | | | | | | | | | | | |
| Scrotifera | X | X | X | X | X | X | X | X | X | X | X | X | X | X | X | X | X |
| Fereunguulata | X | X | X | X | X | X | X | X | X | X | X | X | X | X | X | X | X |
| Zooamata | X | X | X | X | X | X | X | X | X | X | X | X | X | X | X | X | X |
| **Euarchontoglires** | | | | | | | | | | | | | | | | | |
| Euarchonta | X | X | X | | X | X | X | X | X | X | X | X | X | X | X | X | X |
| Sundatheria (Scandentia + Dermoptera) | | | | | | | | | | | | | | | | | |
| Scandentia + Glires | | | | X | | | | | | | | | | | | | |
| **Rodentia** | | | | | | | | | | | | | | | | | |
| Sciurimorpha (Myomorpha, Hystricomorpha) | X | X | X | X | X | X | X | X | X | | X | X | | | | X | / |
| Hystricomorpha (Sciurimorpha, Myomorpha) | | | | | | | | | | | | | | | | | / |
| Myomorpha (Sciurimorpha, Hystricomorpha) | | | | | | | | | | X | | | X | X | X | | / |

**Table S4.**

Per chromosome, post filtering, summary statistics for 100kb alignments used in sliding window analyses.

| | | Length | | Parsimony Informative Sites | | GC Content % | |
|---|---|---|---|---|---|---|---|
| **Chromosome** | **No. of Windows Analysed** | **Mean** | **Median** | **Mean** | **Median** | **Mean** | **Median** |
| **X** | 1027 | 42954 | 42450 | 41104 | 40430 | 39.3 | 38.7 |
| **1** | 427 | 49924 | 51340 | 48100 | 49672 | 42.7 | 41.9 |
| **21** | 361 | 43677 | 44047 | 42491 | 43053 | 40.9 | 39.2 |
| **22** | 349 | 40699 | 40199 | 38978 | 38330 | 50 | 51 |

**Table S5.**

Karyotypic position of human chromosomes 21 (HSA21) and 22 (HSA22) in 123 placental mammal genomes determined by Zoo-FISH (*3*). n(AUT) refers to the haploid number of autosomes in the karyotype. Red values indicate the HSA21 and/or HSA22 segments are located in a telomeric or distal chromosome arm position, or on one of the smallest chromosomes (smallest 25%) in the karyotype (also underlined).

| Order | Taxon | n (AUT) | 21 | 22a | 22b |
|---|---|---|---|---|---|
| **Afrosoricida** | *Chrysochloris asiatica* | 14 | 2q-prox | 4qtel | 4p-prox |
| **Sirenia** | *Trichechus manatus* | 23 | 1ptel | 7qdist | 16pcen |
| **Proboscidea** | *Elephas maximus* | 27 | 3ptel, 21q | 4qtel | 25p-prox |
| **Proboscidea** | *Loxodonta africana* | 27 | 3ptel, 21q | 4qtel | 25p-prox |
| **Tubulidentata** | *Orycteropus afer* | 9 | 2q-prox | 4qtel | 9qtel |
| **Macroscelidea** | *Macroscelides proboscideus* | 12 | 2q-prox | 3qtel | 9qtel |
| **Macroscelidea** | *Elephantulus rupestris* | 12 | 2q-prox | 4qtel | 11qtel |
| **Pilosa** | *Choloepus didactylus* | 32 | 5ptel | 2qtel | 28qtel+28p-prox |
| **Pilosa** | *Choloepus hoffmanni* | 24 | 6qtel | 11qtel | 23qtel |
| **Pilosa** | *Bradypus torquatus* | 24 | 2qtel | 5qtel | 22ptel |
| **Pilosa** | *Bradypus variegatus* | 26 | no signal | 6qtel | 15qprox |
| **Pilosa** | *Tamandua tetradactyla* | 26 | 10p | 4p | 8cen |
| **Cingulata** | *Euphractus sexcinctus* | 28 | 3qtel | 8qtel | 23p-prox |
| **Cingulata** | *Dasypus novemcinctus* | 31 | 4qtel+12qtel | 18qtel | 8ptel |
| **Scandentia** | *Tupaia minor* | 32 | 5p-prox | 7qtel | 28qtel |
| **Dermoptera** | *Galeopterus variegatus* | 27 | 1q-prox | 9qtel | 20qtel |
| **Primates** | *Nycticebus coucang* | 24 | 12qtel | 1ptel | 4ptel |
| | *Galago moholi* | 18 | 3ptel | 12ptel | 4q-prox |
| | *Otolemur crassicaudatus* | 30 | 10qtel | 24qtel | 2ptel |
| | *Eulemur macaco* | 21 | 1qtel | 5qtel | 13qtel |
| | *Eulemur fulvus* | 29 | 1qtel | 10qtel | 19qtel |
| | *Pithecia irrorata* | 23 | 19ptel | 8ptel | |
| | *Cacajao calvus* | 21 | 6ptel, 19qtel | 7ptel | 20 |

| Order | Taxon | n (AUT) | 21 | 22a | 22b |
|---|---|---|---|---|---|
| | *Callicebus lugens* | 7 | 4qdist | 2p-prox, 2cen | |
| | *Callicebus personatus* | 21 | 8ptel | 19qtel | |
| | *Aotus nancymaae* | 26 | 7ptel | 3p-prox | |
| | *Aotus lemurinus* | 27 | 4ptel | 6qtel | |
| | *Cebus albifrons* | 25 | 11ptel | 24tel | |
| | *Cebus apella* | 26 | 11ptel | 24tel | |
| | *Saimiri sciureus* | 21 | 21qtel | 19 | |
| | *Lagothrix lagotricha* | 30 | 23qtel | 30 | |
| | *Alouatta seniculus* | 22 | 9qtel | 5p-prox | |
| | *Alouatta sara* | 24 | 20qtel | 8ptel | |
| | *Alouatta belzebul* | 24 | 18q-prox | 6ptel | |
| | *Alouatta caraya* | 26 | 21qtel | 3ptel | |
| | *Alouatta fusca* | 22 | 16qtel | 8ptel | |
| | *Alouatta arctoidea* | 21 | 17 | 9ptel | |
| | *Ateles belzebuth* | 16 | 10ptel | 3ptel | |
| | *Ateles geoffroyi* | 16 | 11ptel | 3ptel | |
| | *Brachyteles arachnoides* | 30 | 23qtel | 30 | |
| | *Erythrocebus patas* | 26 | 10qtel | 26 | |
| | *Chlorocebus aethiops* | 29 | 2qtel | 19 | |
| | *Macaca mulatta* | 20 | 3ptel | 10ptel | |
| | *Pygathrix nemaeus/Colobus/Nasalis* | 21,22,23 | 21qtel | 21pq | |
| | *Symphalangus syndactylus* | 24 | 5qprox | 18ptel,18q-prox | |
| | *Hylobates lar* | 21 | 15qtel | 8q-prox | |
| | *Hoolock hoolock* | 18 | 2q-prox | 11p | |
| | *Hoolock leuconedys* | 18 | 2q-prox | 11ptel,13qtel | |
| | *Nomascus concolor* | 25 | 25 | 7ptel | |
| | *Pongo abelii* | 23 | 22 | 23 | |
| | *Gorilla gorilla* | 23 | 22 | 23 | |
| | *Pan troglodytes* | 23 | 22 | 23 | |
| | *Pan paniscus* | 23 | 22 | 23 | |

| Order | Taxon | n (AUT) | 21 | 22a | 22b |
|---|---|---|---|---|---|
| | *Homo sapiens* | 22 | 21 | 22 | |
| **Rodentia** | *Eliomys melanurus* | 23 | 3q-prox | 2qtel | 22ptel |
| | *Tamias sibiricus* | 18 | 7qtel | 2qtel | 12pdist |
| | *Marmota himalayana* | 18 | 10qtel | 4qtel | 1q-prox |
| | *Petaurista albiventer* | 18 | 14qtel | 8qtel | 7q-prox |
| | *Xerus erythropus* | 18 | 10qtel | 4qtel | 11p-prox |
| | *Sciurus carolinensis* | 19 | 15qtel | 10qtel | 1qdist,1p-prox |
| | *Castor fiber* | 23 | 2qtel | 6qtel | 11q |
| | *Sicista betulina* | 15 | 4qtel | 12ptel | 14ptel,15cen |
| | *Pedetes capensis* | 18 | 2ptel | 6qtel | 3p-prox |
| | *Spalax ehrenbergi* | 24-29 | 9qtel | 22qtel | 4p-prox |
| | *Mus musculus* | 19 | 16qtel | 5qprox, 10qdist | 11ptel, 15qprox, 16q-prox |
| | *Myocastor coypus* | 20 | 12p-prox | 17qtel | 16pdist |
| | *Cavia porcellus* | 31 | 14qprox | 17q-prox | 22qtel |
| | *Laonastes aenigmamus* | 18-20 | 11qtel | 2q-prox, 18qtel | 3qtel |
| **Lagomorpha** | *Ochotona forresti* | 26 | 11qtel | 7qtel | 25ptel |
| | *Oryctolagus cuniculus* | 21 | 14qtel | 4qtel | 21ptel |
| **Eulipotyphla** | *Neotracus sinensis* | 15 | 9qtel | 10qtel | 9pdist, 12q-prox |
| | *Hemiechinus auritus* | 23 | 7ptel | 20ptel | 6p-prox |
| | *Sorex araneus* | 9 | af,q-prox | ik,p-prox | tu,pdist |
| | *Sorex granarius* | 17 | 1pdist | 9qtel | 17qtel |
| | *Blarinella griselda* | 21 | 10qtel | 2p-prox | 18qdist |
| | *Talpa europaea* | 16 | 3qtel | 15qtel | 14qdist |
| | *Talpa altaica* | 16 | 4qtel | 15qtel | 14qtel |
| **Chiroptera** | *Megaderma spasma* | 18 | 5qtel | 2ptel | 7pdist |
| | *Aselliscus stoliczkanus* | 14 | 4qtel | 1ptel | 11pdist |
| | *Hipposideros larvatus* | 15 | 1qtel | 6qtel | 7p-prox |
| | *Eonycteris spelaea* | 17 | 4qtel | 3ptel | 8pdist |
| | *Glossophaga soricina* | 15 | 3qtel | 3ptel | 8qdist |
| | *Plecotus auritus* | 15 | 3/4qtel | 5/6qtel | 23qtel |

| Order | Taxon | n (AUT) | 21 | 22a | 22b |
|---|---|---|---|---|---|
| | *Taphozous melanopogon* | 20 | 4ptel | 3qtel | 8qdist |
| **Carnivora** | *Crocuta crocuta* | 19 | 4ptel | 15qtel | 10pdist |
| | *Paguma larvata* | 21 | 14ptel | 5qtel | 8pdist |
| | Felidae | 17/18 | C2ptel | B4qtel | D3pdist |
| | *Vulpes vulpes* | 16 | 15ptel | 16qprox | 10pdist |
| | *Canis lupus familiaris* | 38 | 27qtel | 26qtel | 10p-prox |
| | *Procyon lotor* | 18 | 1qtel | 6qtel | 12p-prox |
| | *Mustela vison* | 14 | 5qtel | 9qtel | 4p-prox |
| | *Martes foina* | 18 | 2qtel | 9qtel | 13q-prox |
| | *Mephitis mephitis* | 24 | 2ptel | 4ptel | 13cen, 21ptel |
| | *Ailuropoda melanoleuca* | 20 | 1cen | 15qtel | 12q-prox |
| | *Eumetopias jubatus* | 17 | 1qtel | 6qtel | 14p-dist |
| | *Odobenus rosmarus* | 15 | 2qtel | 8qtel | 14p-dist |
| | *Phoca vitulina* | 15 | 2qtel | 6qtel | 13p-dist |
| | *Pusa sibirica* | 15 | 2qtel | 6qtel | 13p-dist |
| **Pholidota** | *Manis javanica* | 18 | 3q-prox, 3qdist | 10qtel | 15qtel |
| | *Manis pentadactyla* | 19 | 2q-prox, 2qdist | 12qtel | 13qtel |
| **Cetartiodactyla** | *Camelus /Lama* | 36 | 1qtel | 12qtel | 32ptel |
| | *Sus scrofa* | 18 | 13qtel | 5ptel | 14pcen |
| | *Hippopotamus amphibius* | 17 | 11qtel | 7ptel | 9ptel |
| | *Eschrichtius robustus* | 21 | 5qtel | 8qtel | 9qdist |
| | *Neophocaena phocaenoides* | 21 | 5qtel | 8qtel | 9qdist |
| | *Tursiops truncatus* | 21 | 5qtel | 8qtel | 9qdist |
| | *Globicephala macrorhynchus* | 21 | 5qtel | 8qtel | 9qdist |
| | *Tragulus javanicus* | 15 | 5qtel | 1qtel | 6ptel |
| | *Moschus moschiferus* | 28 | 1ptel,1qtel | 5qprox, 5qtel | 20qtel |
| | *Bubalus bubalis* | 24 | 1qprox,1qtel | 4qtel | 17qtel |
| | *Bos taurus* | 29 | 1ptel | 5qprox, 5qtel | 17qtel |
| | *Pseudoryx nghetinhensis* | 24 | 1qtel | 8qtel | 14qtel |
| | *Ovibos moschatus* | 23 | 1pcen, 1qtel | 3q-prox, 2qtel | 1ptel |

| Order | Taxon | n (AUT) | 21 | 22a | 22b |
|---|---|---|---|---|---|
| | *Ovis aries* | 26 | 1pcen, 1qtel | 3qtel | 17qtel |
| | *Okapia johnstoni* | 21 | 7qtel | 15qtel | 6qtel |
| | *Giraffa camelopardalis* | 14 | 2qtel | 4qtel | 11ptel |
| | *Antilocapra americana* | 28 | 1qtel | 5qprox,5qtel | 24qtel |
| | *Muntiacus muntjak* | 3 | 1qcen, 3qcen | 1q-prox | 1p-prox |
| **Perissodactyla** | *Equus caballus* | 31 | 26qtel | 8ptel | 28qtel |
| | *Equus asinus* | 30 | 18qtel | 8qtel | 4ptel |
| | *Equus burchelli* | 21 | 20qtel | 21qtel | 13ptel |
| | *Equus grevyi* | 22 | 20 | 13qtel | 12p-prox |
| | *Tapirus indicus* | 25 | 21 | 22ptel | 7p-prox |
| | *Diceros bicornis* | 41 | 27 | 26qtel | 34ptel |

**Table S6.**

Species compositions for clade-based analysis of phylogenomic signal where a subset is shown in Fig. 2.

| | Ingroup 1 | Ingroup 2 | Ingroup 3 | Outgroup | Autosomes Frequency (%) | | | ChrX Frequency (%) | | |
|---|---|---|---|---|---|---|---|---|---|---|
| | | | | | **t1** | **t2** | **t3** | **t1** | **t2** | **t3** |
| **Paenungulata** | *Heterohyrax brucei* | *Loxodonta africana* | *Trichechus manatus* | *Orycteropus afer* | 80 | 15 | 5 | 80.0 | 14.0 | 6.0 |
| **Laurasiatheria** | *Equus caballus* | *Bos taurus* | *Felis catus* | *Macroglossus sobrunis* | 78.0 | 7.9 | 9.0 | 74.0 | 6.0 | 9.0 |
| **Scandentia** | *Tupaia chinensis* | *Homo sapiens* | *Mus musculus* | *Felis catus* | 96.7 | 2.7 | 0.4 | 88.0 | 9.0 | 3.0 |
| **Rodentia** | *Glis glis* | *Mus musculus* | *Heterocephalus glaber* | *Homo sapiens* | 63.4 | 30.5 | 6.0 | 29.0 | 65.0 | 6.0 |
| **Afroinsectophilia** | *Orycteropus afer* | *Chrysochloris asiatica* | *Elephantulus edwardii* | *Choloepus hoffmanii* | 84.8 | 8.6 | 6.5 | 67.0 | 19.0 | 12.0 |
| **Pteropodidae** | *Pteropus alecto* | *Macroglossus sobrinus* | *Rousettus aegyptiacus* | *Megaderma lyra* | 43.6 | 39.1 | 17.2 | 46.0 | 42.0 | 12.0 |
| **Phyllostomidae** | *Artibeus jamaicensis* | *Anoura caudifer* | *Tonatia saurophila* | *Macroglossus sobrunis* | 71.8 | 20.8 | 7.4 | 56.0 | 22.0 | 22.0 |
| **New World Primates** | *Aotus nancymaae* | *Callithrix jacchus* | *Saimiri boliviensis* | *Homo_sapiens* | 45.2 | 32.7 | 21.6 | 38.0 | 37.0 | 25.0 |
| **Old World Primates** | *Pygathrix nemaeus* | *Nasalis larvatus* | *Rhinopithecus roxellana* | *Homo_sapiens* | 43.0 | 34.2 | 22.9 | 44.0 | 34.0 | 23.0 |
| **Cricetidae** | *Cricetulus griseus* | *Ondatra zibethicus* | *Sigmodon hispidus* | *Mus musculus* | 79.1 | 18.6 | 2.0 | 31.0 | 66.0 | 3.0 |
| **Bovidae** | *Bos taurus* | *Bison bison* | *Bos mutus* | *Capra hircus* | 85.4 | 6.7 | 7.9 | 73.0 | 13.0 | 14.0 |
| **Carnivora** | *Ailuropoda melanoleuca* | *Odobenus rosmarus* | *Mustela putorius* | *Felis catus* | 98.1 | 1.3 | 0.5 | 91.0 | 8.0 | 1.0 |
| **Cetartiodactyla** | *Ammotragus lervia* | *Bos taurus* | *Capra hircus* | *Ovis aries* | 87.0 | 8.8 | 4.2 | 84.0 | 11.0 | 5.0 |
| **Cetartiodactyla** | *Capra aegagrus* | *Capra hircus* | *Ovis aries* | *Ammotragus lervia* | 87.0 | 8.8 | 4.2 | 84.0 | 10.0 | 5.0 |
| **Chiroptera** | *Macroglossus sobrinus* | *Hipposideros armiger* | *Myotis myotis* | *Bos taurus* | 99.7 | 0.0 | 0.0 | 99.0 | 0.2 | 0.5 |
| **Primates** | *Chlorocebus sabaeus* | *Macaca nemistrina* | *Papio anubis* | *Homo sapiens* | 99.4 | 0.6 | 0.1 | 99.5 | 0.1 | 0.2 |
| **Cetartiodactyla** | *Inia geoffrensis* | *Lipotes vexillifer* | *Tursiops truncatus* | *Mesoplodon bidens* | 90.7 | 5.5 | 3.2 | 85.0 | 7.0 | 7.0 |
| **Primates** | *Aotus nancymaae* | *Ateles geoffroyi* | *Callicebus donacophilus* | *Homo sapiens* | 81.0 | 16.2 | 2.8 | 96.0 | 2.0 | 2.0 |
| **Cetartiodactyla** | *Capra hircus* | *Hemitragus hylocrius* | *Ovis aries* | *Bos taurus* | 99.9 | 0.1 | 0.0 | 99.3 | 0.6 | 0.1 |
| **Primates** | *Homo sapiens* | *Pan paniscus* | *Pan troglodytes* | *Macaca fascicularis* | 96.7 | 1.9 | 1.4 | 99.8 | 0.0 | 0.2 |
| **Primates** | *Homo sapiens* | *Gorilla gorilla* | *Pan troglodytes* | *Macaca mulatta* | 95.5 | 2.7 | 1.8 | 99.3 | 0.5 | 0.2 |
| **Cetartiodactyla** | *Inia geoffrensis* | *Kogia breviceps* | *Orcinus orca* | *Platanista gangetica* | 96.1 | 0.9 | 2.5 | 93.3 | 2.2 | 3.3 |
| **Primates** | *Lemur catta* | *Mirza coquereli* | *Nycticebus coucang* | *Propithecus coquereli* | 79.3 | 12.6 | 33.2 | 90.0 | 5.0 | 5.0 |
| **Carnivora** | *Helogale parvula* | *Mungos mungo* | *Suricatta suricatta* | *Felis catus* | 90.1 | 5.5 | 4.3 | 85.0 | 8.0 | 7.0 |
| **Carnivora** | *Mellivora capensis* | *Pteronura brasiliensis* | *Mustela putorius* | *Ailuropoda melanleuca* | 99.8 | 0.1 | 0.1 | 99.3 | 0.3 | 0.4 |

| | Ingroup 1 | Ingroup 2 | Ingroup 3 | Outgroup | Autosomes Frequency (%) | | | ChrX Frequency (%) | | |
|---|---|---|---|---|---|---|---|---|---|---|
| | | | | | t1 | t2 | t3 | t1 | t2 | t3 |
| **Cetartiodactyla** | *Balaenoptera bonaerensis* | *Eschrichtius robustus* | *Eubalaena japonica* | *Hippopotamus amphibius* | 99.8 | 0.2 | 0.0 | 98.1 | 0.9 | 1.0 |
| **Cetartiodactyla** | *Kogia breviceps* | *Platanista gangetica* | *Mesoplodon bidens* | *Hippopotamus amphibius* | 96.4 | 2.5 | 1.1 | 93.5 | 3.4 | 3.1 |
| **Cetartiodactyla** | *Hemitragus hylocrius* | *Ovis aries* | *Ovis canadensis* | *Bos taurus* | 92.7 | 4.2 | 3.1 | 80.0 | 11.0 | 9.0 |
| **Primates** | *Cercocebus atys* | *Mandrillus leucophaeus* | *Papio anubis* | *Homo sapiens* | 87.1 | 7.2 | 5.8 | 89.6 | 4.7 | 5.7 |
| **Carnivora** | *Leptonychotes weddellii* | *Mirounga angustirostris* | *Neomanachus schauinslandi* | *Ailuropoda melanleuca* | 96.6 | 1.7 | 1.7 | 96.0 | 2.0 | 2.0 |
| **Perissodactyla** | *Ceratotherium simum* | *Dicerorhinus sumatrensis* | *Diceros bicornis* | *Equus caballus* | 100.0 | 0.0 | 0.0 | 99.7 | 0.2 | 0.1 |
| **Rodentia** | *Aplodontia rufa* | *Glis glis* | *Muscardinus avellanarius* | *Heterocephalus glaber* | 99.8 | 0.1 | 0.1 | 99.9 | 0.1 | 0.0 |

**Table S7.**

Location of deletions recovered in both the RRA and the HRA/DRA. Coordinates based on dividing human chromosomes into 10Mb windows with the start and end coordinates listed for each window. Window coordinates, as well as start and end sites are listed for deletions derived from the dog genome.

| Hypothesis | Reference | Chromosome | Window | Start | End | Length |
|---|---|---|---|---|---|---|
| **Chiroptera + Cetartiodactyla** | Human | chr20 | 30000000 | 1091244 | 1091254 | 10 |
| **Chiroptera + Cetartiodactyla** | Human | chr5 | 180000000 | 3867644 | 3867657 | 13 |
| **Chiroptera + Cetartiodactyla** | Human | chr20 | 40000000 | 6197154 | 6197167 | 13 |
| **Chiroptera + Cetartiodactyla** | Human | chr2 | 70000000 | 7627350 | 7627366 | 16 |
| **Chiroptera + Cetartiodactyla** | Human | chr12 | 110000000 | 498972 | 498988 | 16 |
| **Chiroptera + Cetartiodactyla** | Human | chr11 | 90000000 | 4819229 | 4819249 | 20 |
| **Chiroptera + Cetartiodactyla + Ferae** | Human | chr10 | 80000000 | 5487518 | 5487528 | 10 |
| **Chiroptera + Cetartiodactyla + Ferae** | Human | chr10 | 110000000 | 5769451 | 5769462 | 11 |
| **Chiroptera + Cetartiodactyla + Ferae** | Human | chr1 | 120000000 | 9218057 | 9218070 | 13 |
| **Chiroptera + Cetartiodactyla + Ferae** | Human | chr13 | 100000000 | 9859580 | 9859596 | 16 |
| **Chiroptera + Cetartiodactyla + Ferae** | Human | chr13 | 100000000 | 5630311 | 5630336 | 25 |
| **Chiroptera + Cetartiodactyla + Perissodactyla** | Human | chr2 | 140000000 | 6391426 | 6391436 | 10 |
| **Chiroptera + Cetartiodactyla + Perissodactyla** | Human | chr10 | 120000000 | 2750604 | 2750615 | 11 |
| **Chiroptera + Cetartiodactyla + Perissodactyla** | Human | chr11 | 40000000 | 860281 | 860292 | 11 |
| **Chiroptera + Cetartiodactyla + Perissodactyla** | Human | chr7 | 30000000 | 8799288 | 8799306 | 18 |
| **Chiroptera + Ferae** | Human | chr15 | 60000000 | 1275672 | 1275682 | 10 |
| **Chiroptera + Ferae** | Human | chr7 | 140000000 | 2885881 | 2885891 | 10 |
| **Chiroptera + Ferae** | Human | chr14 | 30000000 | 5478219 | 5478231 | 12 |
| **Chiroptera + Ferae** | Human | chr14 | 40000000 | 7267010 | 7267022 | 12 |
| **Chiroptera + Ferae** | Human | chr15 | 70000000 | 662811 | 662823 | 12 |
| **Chiroptera + Ferae** | Human | chr16 | 60000000 | 9662902 | 9662914 | 12 |
| **Chiroptera + Ferae** | Human | chr8 | 110000000 | 8114071 | 8114085 | 14 |
| **Chiroptera + Ferae** | Human | chr2 | 80000000 | 2026426 | 2026440 | 14 |

| Hypothesis | Reference | Chromosome | Window | Start | End | Length |
|---|---|---|---|---|---|---|
| **Chiroptera + Ferae** | Human | chr12 | 50000000 | 3096040 | 3096058 | 18 |
| **Chiroptera + Ferae** | Human | chr6 | 120000000 | 1824677 | 1824705 | 28 |
| **Chiroptera + Perissodactyla** | Human | chr9 | 120000000 | 7371989 | 7371999 | 10 |
| **Chiroptera + Perissodactyla** | Human | chr21 | 20000000 | 9803063 | 9803073 | 10 |
| **Chiroptera + Perissodactyla** | Human | chr6 | 150000000 | 9061859 | 9061870 | 11 |
| **Chiroptera + Perissodactyla** | Human | chr20 | 10000000 | 2296115 | 2296126 | 11 |
| **Chiroptera + Perissodactyla** | Human | chr16 | 30000000 | 1455428 | 1455440 | 12 |
| **Chiroptera + Perissodactyla** | Human | chr16 | 60000000 | 7234992 | 7235004 | 12 |
| **Chiroptera + Perissodactyla** | Human | chr2 | 60000000 | 9062292 | 9062305 | 13 |
| **Chiroptera + Perissodactyla** | Human | chr10 | 20000000 | 5778719 | 5778732 | 13 |
| **Chiroptera + Perissodactyla** | Human | chr14 | 70000000 | 9615757 | 9615770 | 13 |
| **Chiroptera + Perissodactyla** | Human | chr2 | 180000000 | 9300857 | 9300871 | 14 |
| **Chiroptera + Perissodactyla** | Human | chr2 | 230000000 | 6869114 | 6869128 | 14 |
| **Chiroptera + Perissodactyla** | Human | chr17 | 80000000 | 2152391 | 2152405 | 14 |
| **Chiroptera + Perissodactyla** | Human | chr5 | 120000000 | 2844574 | 2844589 | 15 |
| **Chiroptera + Perissodactyla** | Human | chr9 | 120000000 | 4551973 | 4551989 | 16 |
| **Chiroptera + Perissodactyla** | Human | chr9 | 70000000 | 9712340 | 9712357 | 17 |
| **Chiroptera + Perissodactyla** | Human | chr8 | 100000000 | 6006172 | 6006192 | 20 |
| **Chiroptera + Perissodactyla** | Human | chr4 | 60000000 | 7282328 | 7282350 | 22 |
| **Chiroptera + Perissodactyla + Ferae** | Human | chr2 | 80000000 | 2175705 | 2175715 | 10 |
| **Chiroptera + Perissodactyla + Ferae** | Human | chr5 | 20000000 | 1990651 | 1990661 | 10 |
| **Chiroptera + Perissodactyla + Ferae** | Human | chr4 | 190000000 | 9005954 | 9005965 | 11 |
| **Chiroptera + Perissodactyla + Ferae** | Human | chr13 | 80000000 | 5819053 | 5819064 | 11 |
| **Chiroptera + Perissodactyla + Ferae** | Human | chr1 | 60000000 | 566478 | 566490 | 12 |
| **Chiroptera + Perissodactyla + Ferae** | Human | chr2 | 150000000 | 7367110 | 7367122 | 12 |
| **Chiroptera + Perissodactyla + Ferae** | Human | chr16 | 20000000 | 5974176 | 5974188 | 12 |
| **Chiroptera + Perissodactyla + Ferae** | Human | chr4 | 150000000 | 4532452 | 4532465 | 13 |
| **Chiroptera + Perissodactyla + Ferae** | Human | chr6 | 100000000 | 522964 | 522978 | 14 |
| **Chiroptera + Perissodactyla + Ferae** | Human | chr10 | 110000000 | 1771807 | 1771821 | 14 |
| **Chiroptera + Perissodactyla + Ferae** | Human | chr6 | 90000000 | 9342036 | 9342056 | 20 |

| Hypothesis | Reference | Chromosome | Window | Start | End | Length |
|---|---|---|---|---|---|---|
| **Ferae + Cetartiodactyla** | Human | chr7 | 120000000 | 6674735 | 6674745 | 10 |
| **Ferae + Cetartiodactyla** | Human | chr10 | 100000000 | 2051815 | 2051825 | 10 |
| **Ferae + Cetartiodactyla** | Human | chr10 | 100000000 | 4209410 | 4209420 | 10 |
| **Ferae + Cetartiodactyla** | Human | chr10 | 120000000 | 6381173 | 6381183 | 10 |
| **Ferae + Cetartiodactyla** | Human | chr18 | 50000000 | 5254710 | 5254720 | 10 |
| **Ferae + Cetartiodactyla** | Human | chr6 | 120000000 | 4381834 | 4381845 | 11 |
| **Ferae + Cetartiodactyla** | Human | chr1 | 80000000 | 25509 | 25521 | 12 |
| **Ferae + Cetartiodactyla** | Human | chr13 | 60000000 | 3045918 | 3045930 | 12 |
| **Ferae + Cetartiodactyla** | Human | chr18 | 60000000 | 4249674 | 4249688 | 14 |
| **Ferae + Cetartiodactyla** | Human | chr3 | 20000000 | 8285602 | 8285617 | 15 |
| **Ferae + Cetartiodactyla** | Human | chr14 | 70000000 | 3283764 | 3283780 | 16 |
| **Ferae + Cetartiodactyla** | Human | chr16 | 20000000 | 2951751 | 2951768 | 17 |
| **Ferae + Cetartiodactyla** | Human | chr1 | 120000000 | 7329553 | 7329572 | 19 |
| **Ferae + Cetartiodactyla** | Human | chr5 | 150000000 | 9808266 | 9808285 | 19 |
| **Ferae + Cetartiodactyla** | Human | chr2 | 80000000 | 2928336 | 2928358 | 22 |
| **Ferae + Cetartiodactyla** | Human | chr9 | 10000000 | 4145378 | 4145400 | 22 |
| **Ferae + Cetartiodactyla** | Human | chr5 | 60000000 | 5916765 | 5916789 | 24 |
| **Ferae + Perissodactyla** | Human | chr1 | 70000000 | 560551 | 560561 | 10 |
| **Ferae + Perissodactyla** | Human | chr1 | 90000000 | 6207284 | 6207296 | 12 |
| **Ferae + Perissodactyla** | Human | chr1 | 40000000 | 3613657 | 3613670 | 13 |
| **Ferae + Perissodactyla** | Human | chr1 | 30000000 | 4257747 | 4257762 | 15 |
| **Ferae + Perissodactyla** | Human | chr10 | 20000000 | 7467914 | 7467931 | 17 |
| **Ferae + Perissodactyla** | Human | chr11 | 20000000 | 9535637 | 9535647 | 10 |
| **Ferae + Perissodactyla** | Human | chr11 | 20000000 | 7874155 | 7874165 | 10 |
| **Ferae + Perissodactyla** | Human | chr11 | 50000000 | 7579430 | 7579440 | 10 |
| **Ferae + Perissodactyla** | Human | chr11 | 120000000 | 3725279 | 3725296 | 17 |
| **Ferae + Perissodactyla** | Human | chr12 | 30000000 | 7967895 | 7967905 | 10 |
| **Ferae + Perissodactyla** | Human | chr13 | 110000000 | 4693946 | 4693956 | 10 |
| **Ferae + Perissodactyla** | Human | chr13 | 80000000 | 2294659 | 2294670 | 11 |
| **Ferae + Perissodactyla** | Human | chr13 | 60000000 | 9582514 | 9582526 | 12 |

| Hypothesis | Reference | Chromosome | Window | Start | End | Length |
|---|---|---|---|---|---|---|
| **Ferae + Perissodactyla** | Human | chr14 | 80000000 | 3794615 | 3794638 | 23 |
| **Ferae + Perissodactyla** | Human | chr16 | 50000000 | 6808470 | 6808480 | 10 |
| **Ferae + Perissodactyla** | Human | chr16 | 80000000 | 3703743 | 3703753 | 10 |
| **Ferae + Perissodactyla** | Human | chr18 | 30000000 | 6386521 | 6386531 | 10 |
| **Ferae + Perissodactyla** | Human | chr2 | 40000000 | 6956297 | 6956307 | 10 |
| **Ferae + Perissodactyla** | Human | chr2 | 150000000 | 3942793 | 3942804 | 11 |
| **Ferae + Perissodactyla** | Human | chr2 | 190000000 | 2103871 | 2103882 | 11 |
| **Ferae + Perissodactyla** | Human | chr2 | 230000000 | 2140864 | 2140876 | 12 |
| **Ferae + Perissodactyla** | Human | chr2 | 50000000 | 5898976 | 5898989 | 13 |
| **Ferae + Perissodactyla** | Human | chr2 | 200000000 | 9327385 | 9327398 | 13 |
| **Ferae + Perissodactyla** | Human | chr2 | 180000000 | 4284240 | 4284254 | 14 |
| **Ferae + Perissodactyla** | Human | chr20 | 10000000 | 7013296 | 7013307 | 11 |
| **Ferae + Perissodactyla** | Human | chr20 | 40000000 | 7840214 | 7840225 | 11 |
| **Ferae + Perissodactyla** | Human | chr20 | 20000000 | 9412052 | 9412068 | 16 |
| **Ferae + Perissodactyla** | Human | chr20 | 50000000 | 535973 | 535990 | 17 |
| **Ferae + Perissodactyla** | Human | chr3 | 110000000 | 7107843 | 7107853 | 10 |
| **Ferae + Perissodactyla** | Human | chr3 | 70000000 | 1008736 | 1008748 | 12 |
| **Ferae + Perissodactyla** | Human | chr3 | 160000000 | 7470055 | 7470078 | 23 |
| **Ferae + Perissodactyla** | Human | chr3 | 140000000 | 3644203 | 3644230 | 27 |
| **Ferae + Perissodactyla** | Human | chr4 | 20000000 | 9652264 | 9652274 | 10 |
| **Ferae + Perissodactyla** | Human | chr5 | 110000000 | 2835808 | 2835820 | 12 |
| **Ferae + Perissodactyla** | Human | chr5 | 150000000 | 3139495 | 3139509 | 14 |
| **Ferae + Perissodactyla** | Human | chr5 | 180000000 | 6594138 | 6594153 | 15 |
| **Ferae + Perissodactyla** | Human | chr6 | 160000000 | 2498919 | 2498932 | 13 |
| **Ferae + Perissodactyla** | Human | chr7 | 130000000 | 1923216 | 1923234 | 18 |
| **Ferae + Perissodactyla** | Human | chr8 | 60000000 | 5314550 | 5314560 | 10 |
| **Ferae + Perissodactyla** | Human | chr8 | 40000000 | 5592696 | 5592707 | 11 |
| **Ferae + Perissodactyla** | Human | chr8 | 130000000 | 622267 | 622282 | 15 |
| **Ferae + Perissodactyla** | Human | chrX | 130000000 | 4713541 | 4713555 | 14 |
| **Ferae + Perissodactyla + Cetartiodactyla** | Human | chr5 | 90000000 | 4559218 | 4559228 | 10 |

| Hypothesis | Reference | Chromosome | Window | Start | End | Length |
|---|---|---|---|---|---|---|
| **Ferae + Perissodactyla + Cetartiodactyla** | Human | chr5 | 140000000 | 2483950 | 2483960 | 10 |
| **Ferae + Perissodactyla + Cetartiodactyla** | Human | chr7 | 30000000 | 1366208 | 1366218 | 10 |
| **Ferae + Perissodactyla + Cetartiodactyla** | Human | chr17 | 10000000 | 2064784 | 2064794 | 10 |
| **Ferae + Perissodactyla + Cetartiodactyla** | Human | chr17 | 10000000 | 3840628 | 3840638 | 10 |
| **Ferae + Perissodactyla + Cetartiodactyla** | Human | chr17 | 20000000 | 3568938 | 3568948 | 10 |
| **Ferae + Perissodactyla + Cetartiodactyla** | Human | chr20 | 10000000 | 6698465 | 6698475 | 10 |
| **Ferae + Perissodactyla + Cetartiodactyla** | Human | chr12 | 60000000 | 9829553 | 9829565 | 12 |
| **Ferae + Perissodactyla + Cetartiodactyla** | Human | chr18 | 20000000 | 3042112 | 3042124 | 12 |
| **Ferae + Perissodactyla + Cetartiodactyla** | Human | chr15 | 70000000 | 2777743 | 2777756 | 13 |
| **Ferae + Perissodactyla + Cetartiodactyla** | Human | chr3 | 110000000 | 7947342 | 7947356 | 14 |
| **Ferae + Perissodactyla + Cetartiodactyla** | Human | chr12 | 110000000 | 5587263 | 5587277 | 14 |
| **Ferae + Perissodactyla + Cetartiodactyla** | Human | chr14 | 40000000 | 616197 | 616211 | 14 |
| **Ferae + Perissodactyla + Cetartiodactyla** | Human | chr17 | 50000000 | 6713969 | 6713983 | 14 |
| **Ferae + Perissodactyla + Cetartiodactyla** | Human | chr18 | 30000000 | 4125502 | 4125517 | 15 |
| **Ferae + Perissodactyla + Cetartiodactyla** | Human | chr12 | 60000000 | 4523973 | 4523989 | 16 |
| **Ferae + Perissodactyla + Cetartiodactyla** | Human | chr10 | 130000000 | 6333171 | 6333188 | 17 |
| **Ferae + Perissodactyla + Cetartiodactyla** | Human | chr7 | 120000000 | 6542789 | 6542809 | 20 |
| **Ferae + Perissodactyla + Cetartiodactyla** | Human | chr10 | 120000000 | 4687520 | 4687544 | 24 |
| **Perissodactyla + Cetartiodactyla** | Human | chr2 | 70000000 | 292297 | 292307 | 10 |
| **Perissodactyla + Cetartiodactyla** | Human | chr3 | 60000000 | 6251377 | 6251387 | 10 |
| **Perissodactyla + Cetartiodactyla** | Human | chr3 | 70000000 | 5594825 | 5594835 | 10 |
| **Perissodactyla + Cetartiodactyla** | Human | chr5 | 100000000 | 4562835 | 4562845 | 10 |
| **Perissodactyla + Cetartiodactyla** | Human | chr6 | 60000000 | 1265874 | 1265884 | 10 |
| **Perissodactyla + Cetartiodactyla** | Human | chr13 | 80000000 | 7928864 | 7928874 | 10 |
| **Perissodactyla + Cetartiodactyla** | Human | chr18 | 60000000 | 3197371 | 3197381 | 10 |
| **Perissodactyla + Cetartiodactyla** | Human | chr20 | 20000000 | 9252021 | 9252031 | 10 |
| **Perissodactyla + Cetartiodactyla** | Human | chr3 | 60000000 | 5539194 | 5539204 | 10 |
| **Perissodactyla + Cetartiodactyla** | Human | chr9 | 80000000 | 3556190 | 3556201 | 11 |
| **Perissodactyla + Cetartiodactyla** | Human | chr15 | 40000000 | 3343650 | 3343662 | 12 |
| **Perissodactyla + Cetartiodactyla** | Human | chr2 | 220000000 | 8587418 | 8587429 | 11 |

| Hypothesis | Reference | Chromosome | Window | Start | End | Length |
|---|---|---|---|---|---|---|
| **Perissodactyla + Cetartiodactyla** | Human | chr5 | 100000000 | 163045 | 163056 | 11 |
| **Perissodactyla + Cetartiodactyla** | Human | chr6 | 30000000 | 2321541 | 2321552 | 11 |
| **Perissodactyla + Cetartiodactyla** | Human | chr10 | 30000000 | 7111679 | 7111690 | 11 |
| **Perissodactyla + Cetartiodactyla** | Human | chr19 | 40000000 | 1815498 | 1815509 | 11 |
| **Perissodactyla + Cetartiodactyla** | Human | chr10 | 10000000 | 8435163 | 8435175 | 12 |
| **Perissodactyla + Cetartiodactyla** | Human | chr14 | 90000000 | 6858991 | 6859003 | 12 |
| **Perissodactyla + Cetartiodactyla** | Human | chr18 | 10000000 | 2796350 | 2796362 | 12 |
| **Perissodactyla + Cetartiodactyla** | Human | chr20 | 10000000 | 5923437 | 5923449 | 12 |
| **Perissodactyla + Cetartiodactyla** | Human | chr12 | 80000000 | 7190714 | 7190727 | 13 |
| **Perissodactyla + Cetartiodactyla** | Human | chr15 | 70000000 | 7747465 | 7747478 | 13 |
| **Perissodactyla + Cetartiodactyla** | Human | chr2 | 140000000 | 6136184 | 6136198 | 14 |
| **Perissodactyla + Cetartiodactyla** | Human | chr17 | 60000000 | 6615352 | 6615366 | 14 |
| **Perissodactyla + Cetartiodactyla** | Human | chr1 | 100000000 | 9580206 | 9580221 | 15 |
| **Perissodactyla + Cetartiodactyla** | Human | chr1 | 40000000 | 4160247 | 4160263 | 16 |
| **Perissodactyla + Cetartiodactyla** | Human | chr6 | 60000000 | 887061 | 887077 | 16 |
| **Perissodactyla + Cetartiodactyla** | Human | chr12 | 100000000 | 8898866 | 8898882 | 16 |
| **Perissodactyla + Cetartiodactyla** | Human | chr16 | 10000000 | 3243747 | 3243765 | 18 |
| **Scrotifera** | Human | chr10 | 20000000 | 9793359 | 9793370 | 11 |
| **Scrotifera** | Human | chr10 | 110000000 | 2233955 | 2233967 | 12 |
| **Scrotifera** | Human | chr10 | 120000000 | 1233868 | 1233883 | 15 |
| **Scrotifera** | Human | chr12 | 130000000 | 357803 | 357814 | 11 |
| **Scrotifera** | Human | chr12 | 80000000 | 1335311 | 1335323 | 12 |
| **Scrotifera** | Human | chr12 | 110000000 | 2475110 | 2475124 | 14 |
| **Scrotifera** | Human | chr14 | 80000000 | 9373773 | 9373783 | 10 |
| **Scrotifera** | Human | chr14 | 80000000 | 8362599 | 8362612 | 13 |
| **Scrotifera** | Human | chr14 | 70000000 | 4742285 | 4742306 | 21 |
| **Scrotifera** | Human | chr14 | 40000000 | 4965296 | 4965334 | 38 |
| **Scrotifera** | Human | chr17 | 60000000 | 9735191 | 9735204 | 13 |
| **Scrotifera** | Human | chr18 | 40000000 | 9972458 | 9972471 | 13 |
| **Scrotifera** | Human | chr18 | 60000000 | 6197277 | 6197293 | 16 |

| Hypothesis | Reference | Chromosome | Window | Start | End | Length |
|---|---|---|---|---|---|---|
| **Scrotifera** | Human | chr2 | 40000000 | 3524607 | 3524617 | 10 |
| **Scrotifera** | Human | chr2 | 120000000 | 6776384 | 6776394 | 10 |
| **Scrotifera** | Human | chr2 | 160000000 | 7305584 | 7305594 | 10 |
| **Scrotifera** | Human | chr2 | 100000000 | 9583129 | 9583140 | 11 |
| **Scrotifera** | Human | chr2 | 190000000 | 1850219 | 1850230 | 11 |
| **Scrotifera** | Human | chr2 | 230000000 | 1824653 | 1824664 | 11 |
| **Scrotifera** | Human | chr2 | 20000000 | 9700571 | 9700583 | 12 |
| **Scrotifera** | Human | chr2 | 110000000 | 2932816 | 2932829 | 13 |
| **Scrotifera** | Human | chr2 | 150000000 | 9676847 | 9676861 | 14 |
| **Scrotifera** | Human | chr2 | 180000000 | 3567912 | 3567926 | 14 |
| **Scrotifera** | Human | chr3 | 70000000 | 1679316 | 1679326 | 10 |
| **Scrotifera** | Human | chr3 | 70000000 | 6746912 | 6746925 | 13 |
| **Scrotifera** | Human | chr3 | 120000000 | 4718289 | 4718303 | 14 |
| **Scrotifera** | Human | chr3 | 130000000 | 3946950 | 3946965 | 15 |
| **Scrotifera** | Human | chr3 | 170000000 | 9088841 | 9088857 | 16 |
| **Scrotifera** | Human | chr4 | 150000000 | 7530559 | 7530569 | 10 |
| **Scrotifera** | Human | chr4 | 90000000 | 952091 | 952102 | 11 |
| **Scrotifera** | Human | chr4 | 60000000 | 5111722 | 5111737 | 15 |
| **Scrotifera** | Human | chr4 | 30000000 | 3937193 | 3937214 | 21 |
| **Scrotifera** | Human | chr4 | 190000000 | 1703159 | 1703183 | 24 |
| **Scrotifera** | Human | chr5 | 130000000 | 7762238 | 7762248 | 10 |
| **Scrotifera** | Human | chr5 | 40000000 | 2045181 | 2045193 | 12 |
| **Scrotifera** | Human | chr5 | 100000000 | 677227 | 677240 | 13 |
| **Scrotifera** | Human | chr5 | 70000000 | 1663779 | 1663798 | 19 |
| **Scrotifera** | Human | chr5 | 110000000 | 7951242 | 7951265 | 23 |
| **Scrotifera** | Human | chr6 | 50000000 | 4023132 | 4023143 | 11 |
| **Scrotifera** | Human | chr6 | 120000000 | 9328903 | 9328915 | 12 |
| **Scrotifera** | Human | chr6 | 50000000 | 2538073 | 2538086 | 13 |
| **Scrotifera** | Human | chr6 | 60000000 | 1152106 | 1152119 | 13 |
| **Scrotifera** | Human | chr6 | 100000000 | 63508 | 63522 | 14 |

| Hypothesis | Reference | Chromosome | Window | Start | End | Length |
|---|---|---|---|---|---|---|
| **Scrotifera** | Human | chr6 | 140000000 | 9906107 | 9906123 | 16 |
| **Scrotifera** | Human | chr7 | 110000000 | 8367586 | 8367596 | 10 |
| **Scrotifera** | Human | chr7 | 30000000 | 7873081 | 7873093 | 12 |
| **Scrotifera** | Human | chr7 | 90000000 | 870881 | 870900 | 19 |
| **Scrotifera** | Human | chr8 | 80000000 | 6783282 | 6783292 | 10 |
| **Scrotifera** | Human | chr8 | 110000000 | 7505181 | 7505194 | 13 |
| **Scrotifera** | Human | chr9 | 20000000 | 4330937 | 4330947 | 10 |
| **Scrotifera** | Human | chr9 | 120000000 | 6137497 | 6137507 | 10 |
| **Scrotifera** | Human | chr9 | 120000000 | 4573709 | 4573720 | 11 |
| **Scrotifera** | Human | chr9 | 130000000 | 961661 | 961673 | 12 |
| **Scrotifera** | Human | chr9 | 110000000 | 6889214 | 6889234 | 20 |
| **Euarchonta** | Dog | chr29 | 10000000-20000000 | 1628320 | 1628340 | 20 |
| **Euarchonta** | Dog | chr33 | 20000000-30000000 | 5289926 | 5289937 | 11 |
| **Euarchonta** | Dog | chrX | 80000000-90000000 | 1133041 | 1133057 | 16 |
| **Euarchonta** | Dog | chr2 | 30000000-40000000 | 7489734 | 7489755 | 21 |
| **Euarchonta** | Dog | chr6 | 30000000-40000000 | 3071133 | 3071154 | 21 |
| **Euarchonta** | Dog | chr8 | 40000000-50000000 | 5733621 | 5733636 | 15 |
| **Euarchonta** | Dog | chr10 | 0-10000000 | 8000914 | 8000931 | 17 |
| **Euarchonta** | Dog | chr15 | 60000000-64190966 | 1866483 | 1866504 | 21 |
| **Euarchonta** | Dog | chr26 | 10000000-20000000 | 4951780 | 4951791 | 11 |
| **Euarchonta** | Dog | chr6 | 50000000-60000000 | 1918005 | 1918015 | 17 |

**Table S8.**
List of breakpoints supporting ordinal and superordinal clades.

| Clade | Reference Chr. | Breakpoint |
|---|---|---|
| **Afrotheria** | 4 | 33242998--33242998=Breakpoints+0.147058823529412 |
| **Atlantogenata** | 8 | 7468577--7468577=Breakpoints+0.205882352941176 |
| **Ferae** | 9 | 45747286--45747286=Breakpoints+0.0588235294117647 |
| **Fereuungulata** | 7 | 77013930--77013930=Breakpoints+0.176470588235294 |
| **Fereuungulata** | 18 | 28356872--28356872=Breakpoints+0.176470588235294 |
| **Hominidae (Primates)** | 9 | 14639345--14639345=Breakpoints+0.0588235294117647 |
| **Hominidae (Primates)** | 10 | 31051431--31051431=Breakpoints+0.0588235294117647 |
| **Hominidae (Primates)** | 11 | 36265915--36265915=Breakpoints+0.0588235294117647 |
| **Hominidae (Primates)** | 18 | 37605607--37605607=Breakpoints+0.0735294117647059 |
| **Hominidae (Primates)** | 19 | 15761798--15761798=Breakpoints+0.0588235294117647 |
| **Hominidae (Primates)** | 19 | 18398292--18398292=Breakpoints+0.0735294117647059 |
| **Hominidae (Primates)** | 21 | 32387272--32387272=Breakpoints+0.0735294117647059 |
| **Hominidae (Primates)** | 25 | 1880175--1880175=Breakpoints+0.176470588235294 |
| **Hominidae (Primates)** | 25 | 2297381--2297381=Breakpoints+0.205882352941176 |
| **Hominidae (Primates)** | 26 | 17738539--17738539=Breakpoints+0.0735294117647059 |
| **Hominidae (Primates)** | 27 | 9028684--9028684=Breakpoints+0.0735294117647059 |
| **Hominidae (Primates)** | 32 | 14859021--14859021=Breakpoints+0.117647058823529 |
| **Hominidae (Primates)** | 37 | 2077878--2077878=Breakpoints+0.0882352941176471 |
| **Hominidae (Primates)** | 38 | 1146716--1146716=Breakpoints+0.0588235294117647 |
| **Hominidae (Primates)** | 38 | 3297188--3297188=Breakpoints+0.0588235294117647 |
| **Laurasiatheria** | 4 | 11803663--11803663=Breakpoints+0.338235294117647 |
| **Muridae (Rodentia)** | 1 | 11010363--11010363=Breakpoints+0.0735294117647059 |
| **Muridae (Rodentia)** | 1 | 73435141--73435141=Breakpoints+0.0588235294117647 |
| **Muridae (Rodentia)** | 1 | 110892617--110892617=Breakpoints+0.0735294117647059 |
| **Muridae (Rodentia)** | 2 | 50007766--50007766=Breakpoints+0.0588235294117647 |
| **Muridae (Rodentia)** | 2 | 69573856--69573856=Breakpoints+0.0588235294117647 |
| **Muridae (Rodentia)** | 2 | 78397655--78397655=Breakpoints+0.0588235294117647 |
| **Muridae (Rodentia)** | 2 | 81674357--81674357=Breakpoints+0.0588235294117647 |
| **Muridae (Rodentia)** | 2 | 88437468--88437468=Breakpoints+0.0588235294117647 |
| **Muridae (Rodentia)** | 3 | 83971173--83971173=Breakpoints+0.0882352941176471 |
| **Muridae (Rodentia)** | 3 | 93609187--93609187=Breakpoints+0.0588235294117647 |
| **Muridae (Rodentia)** | 4 | 2701825--2701825=Breakpoints+0.0735294117647059 |
| **Muridae (Rodentia)** | 4 | 19573477--19573477=Breakpoints+0.102941176470588 |
| **Muridae (Rodentia)** | 4 | 79141717--79528689=Breakpoints+0.0588235294117647 |
| **Muridae (Rodentia)** | 4 | 100791638--100791638=Breakpoints+0.0588235294117647 |
| **Muridae (Rodentia)** | 5 | 8010530--8010530=Breakpoints+0.0735294117647059 |
| **Muridae (Rodentia)** | 5 | 19573672--19573672=Breakpoints+0.0882352941176471 |

| Clade | Reference Chr. | Breakpoint |
|---|---|---|
| **Muridae (Rodentia)** | 5 | 83470174--83470174=Breakpoints+0.0735294117647059 |
| **Muridae (Rodentia)** | 6 | 15957068--15957068=Breakpoints+0.0588235294117647 |
| **Muridae (Rodentia)** | 6 | 24525634--24525634=Breakpoints+0.0588235294117647 |
| **Muridae (Rodentia)** | 6 | 47500112--47500112=Breakpoints+0.0588235294117647 |
| **Muridae (Rodentia)** | 7 | 24452975--24452975=Breakpoints+0.0735294117647059 |
| **Muridae (Rodentia)** | 7 | 35836670--35836670=Breakpoints+0.0588235294117647 |
| **Muridae (Rodentia)** | 7 | 49168639--49168639=Breakpoints+0.0588235294117647 |
| **Muridae (Rodentia)** | 7 | 59584580--59584580=Breakpoints+0.0588235294117647 |
| **Muridae (Rodentia)** | 7 | 84850905--84850905=Breakpoints+0.0735294117647059 |
| **Muridae (Rodentia)** | 7 | 98095291--98095291=Breakpoints+0.102941176470588 |
| **Muridae (Rodentia)** | 8 | 43286486--43286486=Breakpoints+0.0735294117647059 |
| **Muridae (Rodentia)** | 9 | 11456630--11456630=Breakpoints+0.0735294117647059 |
| **Muridae (Rodentia)** | 9 | 70918067--71456234=Breakpoints+0.176470588235294 |
| **Muridae (Rodentia)** | 10 | 28994225--28994225=Breakpoints+0.0735294117647059 |
| **Muridae (Rodentia)** | 10 | 59733800--59733800=Breakpoints+0.0735294117647059 |
| **Muridae (Rodentia)** | 11 | 33790608--33790608=Breakpoints+0.0588235294117647 |
| **Muridae (Rodentia)** | 12 | 25389851--25389851=Breakpoints+0.0588235294117647 |
| **Muridae (Rodentia)** | 12 | 26138109--26138109=Breakpoints+0.0588235294117647 |
| **Muridae (Rodentia)** | 12 | 42771572--42771572=Breakpoints+0.0735294117647059 |
| **Muridae (Rodentia)** | 12 | 52320511--52320511=Breakpoints+0.0588235294117647 |
| **Muridae (Rodentia)** | 15 | 25152799--25152799=Breakpoints+0.0735294117647059 |
| **Muridae (Rodentia)** | 15 | 42851923--42851923=Breakpoints+0.0588235294117647 |
| **Muridae (Rodentia)** | 16 | 1556982--1556982=Breakpoints+0.0735294117647059 |
| **Muridae (Rodentia)** | 16 | 41368842--41368842=Breakpoints+0.132352941176471 |
| **Muridae (Rodentia)** | 18 | 7500019--7500019=Breakpoints+0.102941176470588 |
| **Muridae (Rodentia)** | 20 | 7884837--7884837=Breakpoints+0.102941176470588 |
| **Muridae (Rodentia)** | 20 | 32149007--32486621=Breakpoints+0.102941176470588 |
| **Muridae (Rodentia)** | 21 | 17209575--17209575=Breakpoints+0.0588235294117647 |
| **Muridae (Rodentia)** | 22 | 21485725--21485725=Breakpoints+0.0882352941176471 |
| **Muridae (Rodentia)** | 24 | 6843751--6843751=Breakpoints+0.117647058823529 |
| **Muridae (Rodentia)** | 25 | 5944723--5944723=Breakpoints+0.0588235294117647 |
| **Muridae (Rodentia)** | 28 | 5698926--5698926=Breakpoints+0.0735294117647059 |
| **Muridae (Rodentia)** | 28 | 6019417--6019417=Breakpoints+0.0588235294117647 |
| **Muridae (Rodentia)** | 28 | 9813003--9813003=Breakpoints+0.0882352941176471 |
| **Muridae (Rodentia)** | 29 | 9667338--10395629=Breakpoints+0.0882352941176471 |
| **Muridae (Rodentia)** | 29 | 17027481--17027481=Breakpoints+0.0735294117647059 |
| **Paenungulata** | 4 | 36577499--36577499=Breakpoints+0.191176470588235 |
| **Paenungulata** | 6 | 90345447--90345447=Breakpoints+0.0882352941176471 |
| **Paenungulata** | 13 | 31487590--31487590=Breakpoints+0.0882352941176471 |
| **Paenungulata** | 22 | 19637272--19637272=Breakpoints+0.0882352941176471 |
| **Paenungulata** | 58 | 331491--331491=Breakpoints+0.117647058823529 |

| Clade | Reference Chr. | Breakpoint |
|---|---|---|
| **Perissodactyla** | 2 | 106070986--106070986=Breakpoints+0.0735294117647059 |
| **Perissodactyla** | 2 | 114747381--114747381=Breakpoints+0.0588235294117647 |
| **Perissodactyla** | 2 | 115184003--115184003=Breakpoints+0.102941176470588 |
| **Perissodactyla** | 3 | 60920090--60920090=Breakpoints+0.0588235294117647 |
| **Perissodactyla** | 3 | 61374454--61374454=Breakpoints+0.0588235294117647 |
| **Perissodactyla** | 4 | 63995357--63995357=Breakpoints+0.0735294117647059 |
| **Perissodactyla** | 9 | 49385716--49385716=Breakpoints+0.0735294117647059 |
| **Perissodactyla** | 12 | 32370297--32370297=Breakpoints+0.0735294117647059 |
| **Ruminantia** | 1 | 132815479--132815479=Breakpoints+0.0735294117647059 |
| **Ruminantia** | 2 | 28378951--28378951=Breakpoints+0.0735294117647059 |
| **Ruminantia** | 3 | 126947524--126947524=Breakpoints+0.0588235294117647 |
| **Ruminantia** | 5 | 10932766--10932766=Breakpoints+0.0588235294117647 |
| **Ruminantia** | 7 | 71464318--71464318=Breakpoints+0.0588235294117647 |
| **Ruminantia** | 8 | 12119376--12119376=Breakpoints+0.0588235294117647 |
| **Ruminantia** | 8 | 86094936--86094936=Breakpoints+0.0588235294117647 |
| **Ruminantia** | 8 | 86787815--86787815=Breakpoints+0.0882352941176471 |
| **Ruminantia** | 9 | 25884380--25884380=Breakpoints+0.0588235294117647 |
| **Ruminantia** | 9 | 50150313--50150313=Breakpoints+0.0735294117647059 |
| **Ruminantia** | 13 | 40234935--40234935=Breakpoints+0.0735294117647059 |
| **Ruminantia** | 14 | 47302083--47302083=Breakpoints+0.117647058823529 |
| **Ruminantia** | 14 | 48267382--48267382=Breakpoints+0.102941176470588 |
| **Ruminantia** | 16 | 743825--743825=Breakpoints+0.0882352941176471 |
| **Ruminantia** | 16 | 41982096--41982096=Breakpoints+0.132352941176471 |
| **Ruminantia** | 17 | 25906774--25906774=Breakpoints+0.0735294117647059 |
| **Ruminantia** | 20 | 2407886--2407886=Breakpoints+0.102941176470588 |
| **Ruminantia** | 21 | 31765587--31765587=Breakpoints+0.0882352941176471 |
| **Ruminantia** | 26 | 4926313--4926313=Breakpoints+0.0735294117647059 |
| **Ruminantia** | 33 | 13060329--13060329=Breakpoints+0.0735294117647059 |
| **Ruminantia** | 46 | 1635899--1635899=Breakpoints+0.117647058823529 |
| **Scrotifera** | 3 | 63595219--63595219=Breakpoints+0.308823529411765 |
| **Scrotifera** | 11 | 30843488--30843488=Breakpoints+0.294117647058824 |
| **Scrotifera** | 10 | 49546161--49546161=Breakpoints+0.25 |
| **Scrotifera** | 10 | 50121859--50121859=Breakpoints+0.25 |
| **Xenarthra** | 4 | 54455889--54455889=Breakpoints+0.0588235294117647 |
| **Xenarthra** | 13 | 4088996--4088996=Breakpoints+0.0588235294117647 |

**Table S9.**

Soft bounded fossil calibrations used to constrain divergence time analyses. Stratigraphic bounding indicates where two successive underlying, fossil-bearing chronologic units that did not contain any fossils from the clade of interest are used to define an upper bound. Phylogenetic bracketing uses the age of the oldest stem fossils up to two nodes below the divergence event to define a bound. Phylogenetic uncertainty allows for the possibility that a particular fossil might belong to the stem of a given clade.

| | **Clade** | **Oldest crown fossil** | **Minimum age** | **Maximum age** | **References for oldest fossil(s) and discussion of calibration** | **Comments on Minimum Constraint** | **Comments on Maximum Constraint** | **Type** |
|---|---|---|---|---|---|---|---|---|
| **1** | Paenungulata | *Eritherium azzouzorum* | 59.2 | 72.3 | (*99–101*) | *Eritherium* was originally described as a stem proboscidean (*99*), but is the sister group to Proboscidea, Sirenia, or Tethytheria in more recent analyses (*102*). | Stratigraphic Bounding | Cladistic |
| **2** | Proboscidea to Hyracoidea | *Phosphatherium escuilliei* is the oldest unequivocal stem proboscidean | 56 | 66 | | | | Cladistic |
| **3** | Xenarthra | *Riostegotherium* | 47.8 | 66 | (*101*, *103*, *104*) | | Phylogenetic Bracketing | Opinion |
| **4** | Pilosa | *Pseudoglyptodon chilensis* | 31.5 | 66 | (*101*) | | Phylogenetic Bracketing | Opinion |
| **5** | Carnivora | *Hesperocyon gregarious* | 37.71 | 66 | (*87*, *101*, *105*, *106*) | | Phylogenetic Uncertainty. Following (*101*) | Cladistic |
| **6** | Pinnipedia + Musteloidea | *Corumictis wolsani* is the oldest stem mustelid (Paterson et al. 2020) | 28 | 41.2 | (*107*) | *Corumictis wolsani* is the oldest stem mustelid and is known from the early Oligocene (Rupelian). The top of the early Oligocene is 27.82 Ma based on the International Chronostratigraphic Chart (v 2021/07) | Stratigraphic Bounding | Cladistic |
| **7** | Arctoidea | *Enaliarctos tedfordi* | 28.1 | 41.2 | (*6*, *108*, *109*) | Previously the ursid *Cephalogale* has been used to constrain the crown group Actoidea to a minimum of 28.1 Ma or the | Stratigraphic Bounding | Cladistic |

| | Clade | Oldest crown fossil | Minimum age | Maximum age | References for oldest fossil(s) and discussion of calibration | Comments on Minimum Constraint | Comments on Maximum Constraint | Type |
|---|---|---|---|---|---|---|---|---|
| | | | | | | top of the early Oligocene (Rupelian) (*101*). The oldest stem mustelid (*Corumictis wolsani*) is also known from the early Oligocene (*108*), as are fossil remains of the stem pinniped *Enaliarctos tedfordi* from the Yaquina Formation in Oregon (*109*) that were dated to 30.6 – 27.4 Ma based on a paleomagnetic and K:Ar study (*6*). We used a date of 27.82 Ma for the top of the early Oligocene based on the most recent version of the International Chronostratigraphic Chart (v 2021/07) | | |
| **8** | Caniformia | *Hesperocyon gregarious* | 37.71 | 56 | (*101*) | | Stratigraphic Bounding | Cladistic |
| **9** | Ostentoria/Ferae | *Ravenictis* | 64 | 83.8 | (*110*) | | Stratigraphic Bounding | Opinion |
| **10** | Hipposideridae + Rhinolophidae | *Hipposideros* | 37.8 | 56 | (*101*, *111*, *112*) | | Stratigraphic Bounding | Opinion |
| **11** | Chiroptera | *Tachypteron franzeni, Witwatia sigei, Dizzya exsultans* | 47.8 | 66 | (*88*, *101*, *113*, *114*) | | Phylogenetic uncertainty. Following (*101*) | Opinion |
| **12** | Molossidae + Vespertilionidae + Miniopteridae | *Wallia scalopidens* | 38 | 56 | (*101*, *112*, *115*) | | Stratigraphic Bounding | Opinion |
| **13** | Yangochiroptera | *Tachypteron franzeni, Witwatia sigei, Dizzya exsultans* | 47.8 | 61.6 | (*101*, *113*, *114*) | | Stratigraphic Bounding | Opinion |
| **14** | Phyllostomidae + Mormoopidae | *Mormoopid, gen. et sp. nov.* | 27.82 | 41.2 | (*101*, *111*, *116*, *117*) | | Stratigraphic Bounding | Opinion |

| | Clade | Oldest crown fossil | Minimum age | Maximum age | References for oldest fossil(s) and discussion of calibration | Comments on Minimum Constraint | Comments on Maximum Constraint | Type |
|---|---|---|---|---|---|---|---|---|
| **15** | Erinaceidae + Soricidae | *Adunator ladae/Litolestes ignotus* | 61.6 | 83.8 | (*6*, *7*, *87*, *101*, *118*) | While *Adunator ladae* most likely represents a stem erinaceid fossil, we also cite the more confidently assigned *Litolestes ignotus* following (*119*). This fossil is from the Ravenscrag Formation in Canada which is dated to the Tiffanian – the lower boundary of which is given as the Selandian (Paleobiology Database). This gives a minimum bound of 61.6 Ma in line with the ICS. (*119*) use the base of the Danian as the maximum bound, 66 Ma. | Stratigraphic Bounding | Opinion |
| **16** | Cetartiodactyla | *Himalayacetus* | 52.5 | 66 | (*101*) | Maximum age is based on the possibility that mesonychids are crown cetartiodactyls. | Phylogenetic Bracketing/Phylogenetic Uncertainty | Opinion |
| **17** | Moschidae to Bovidae | *Eotragus noyei* | 18 | 27.82 | (*101*, *120*) | | Stratigraphic Bounding. | Opinion |
| **18** | Suidae to Tayassuidae | *Hyotherium,* | 15.97 | 37.71 | (*101*, *121*) | | Phylogenetic Bracketing. Following (*101*) | Cladistic |
| **19** | Whippomorpha | *Himalayacetus* | 52.5 | 61.6 | (*101*) | | Stratigraphic Bounding. | Opinion |
| **20** | Cetacea | *Mystacodon selensis* | 36.4 | 47.8 | (*122*) | The oldest crown cetacean fossil is *Mystacodon selenensis* from the Late Eocene of Peru (*122*). This fossil gives a new minimum bound for this clade of 36.4 Ma – which (*122*) derived from assessing sedimentation rates, a result corroborated by biostratigraphic and biochronological analyses. | Stratigraphic Bounding. (*101*) used stratigraphic bounding of two stages to give a maximum bound of 47.8 Ma. | Cladistic |

| | Clade | Oldest crown fossil | Minimum age | Maximum age | References for oldest fossil(s) and discussion of calibration | Comments on Minimum Constraint | Comments on Maximum Constraint | Type |
|---|---|---|---|---|---|---|---|---|
| **21** | Mysticeti | *Moronocetus parvus* is a stem balaenid and has a minimum age of 15.97 Ma; maximum based on the age of the oldest stem mysticete (*Mystacodon*) | 15.97 | 36.4 | (*101*, *123*, *124*) | *Mauicetus parki* is older than *Morenocetus* (25.2-23.0 Ma) but may be crown mysticete (*125*) or a stem mysticete (*126*). | | Cladistic |
| **22** | Odontoceti | *Ferecetotherium* | 23.03 | 33.9 | (*127*, *128*) | | Stratigraphic Bounding. | Opinion |
| **23** | Perissodactyla | *Hyracotherium, Sifrhippus* | 56 | 61.6 | (*101*, *129*) | *Sifrhippus* is know from the Paleocene-Eocene Thermal Maximum (PETM) (*130*). | Stratigraphic Bounding. | Cladistic |
| **24** | Ceratomorpha | *Cambaylophus* | 53.7 | 61.6 | (*131*) | *Cambaylophus* from the early Eocene Cambay Shale formation from western India is the oldest known ceratomorph. The minimum for this node is given on the basis that Kapur and Bajpai (*131*) state that the mammals from this formation significantly predates the Second Eocene Thermal Maximum (SETM) dated at 53.7 Ma. The maximum bound for this node is given by stratigraphic bounding of two stages (Thanetian and Selandian), 61.6 Ma following (*132*). | Stratigraphic Bounding. | Cladistic |
| **25** | Primates | *Teilhardina brandti* is from the PETM of North America at ~56 Ma (Morse et al. 2019) | 56 | 66 | (*133*) | Calibration is actually for Haplorhini but *Tarsius* is missing from data set | Stratigraphic Bounding. | Cladistic |
| **26** | Anthropoidea | *Aegyptopithecus* | 27.82 | 56 | (*134*, *135*) | | | Cladistic |
| **27** | Platyrrhini | *Panamacebus transitus* | 20.76 | 37.3 | (*136*) | The oldest fossil attributable to this node is now *Panamacebus transitus* from the Las | Stratigraphic Bounding. | Cladistic |

| | Clade | Oldest crown fossil | Minimum age | Maximum age | References for oldest fossil(s) and discussion of calibration | Comments on Minimum Constraint | Comments on Maximum Constraint | Type |
|---|---|---|---|---|---|---|---|---|
| | | | | | | Cascadas Formation of the Panama Canal Basin (*136*). Magmatic Zircons from about 0.25m below the primate bearing horizon yield an age estimate for this layer of 20.93 ± 0.17 Ma. Conservatively, the upper bound of the Aquitanian should be used as the minimum bound for this node, 20.44 Ma. | | |
| **28** | Catarrhini | *Rukwapithecus fleaglei* | 25.22 | 37.71 | (*137*, *138*) | Stevens et al (*137*) describe *Rukwapithecus fleaglei* as the earliest crown catarrine and phylogenetic analyses place it within the Nyanzapithicine clade. Although this relationship is not robustly supported in Stevens et al.'s (*137*) analysis, later analyses aimed at placing the much more extensive cranial remains of *Nyanzapithecus alesi* among catarrhines, also recovered *Rukwapithecus* among the Nyanzapithicine clade (*138*). Based on high precision CA-TIMS U-Pb zircon dating this of this fossil locality provides a minimum constraint of 25.22 Ma (*137*) to this node. | Stratigraphic Bounding. | Cladistic |
| **29** | Cercopithecoidea | *Microcolobus turgenensis* | 12.47 | 20.44 | (*135*, *139*) | *Microcolobus tugenesis* is the oldest fossil attributable to this node (*119*). However a recently discovered specimen, from the | Stratigraphic Bounding. | Opinion |

| | Clade | Oldest crown fossil | Minimum age | Maximum age | References for oldest fossil(s) and discussion of calibration | Comments on Minimum Constraint | Comments on Maximum Constraint | Type |
|---|---|---|---|---|---|---|---|---|
| | | | | | | exceptionally calibrated Kabasero type section of the Ngoragora Formation, is estimated at the older age of 12.49 ± 0.02 Ma (*139*)compared to the more recent specimen referenced in Springer et al. (*135*). This yields a minimum constraint of 12.47 Ma for this node. | | |
| **30** | Strepsirrhini | *Saharagalago misrensis* | 37.71 | 56 | (*101*, *140*) | | Stratigraphic Bounding and Phylogenetic Uncertainty. | Cladistic |
| **31** | Lagomorpha | Tarsals from Vastan mine | 53.7 | 61.6 | (*88*, *101*, *131*, *141*) | | Stratigraphic Bounding and Phylogenetic Uncertainty. | Cladistic |
| **32** | Rodentia | *Acritoparamys, Paramys* | 56 | 66 | (*101*, *118*) | | Stratigraphic Bounding, Phylogenetic Uncertainty, Phylogenetic Bracketing. | Cladistic |
| **33** | Sciuromorpha | *Eogliravus* | 47.8 | 61.6 | (*6*, *73*, *101*) | | | Opinion |
| **34** | Hystricognathi | *Cachiyacuy contamanensis* | 40.94 | 56 | (*88*, *101*, *142*, *143*) | | Stratigraphic Bounding. | Cladistic |
| **35** | Myomorpha (=Muroidea to Dipodidae) | *Pappocricetodon* | 45 | 59.2 | (*101*, *144*) | | Stratigraphic Bounding. | Opinion |
| **36** | Castorimorpha | *Mattimys* | 52.4 | 61.6 | (*101*) | | Stratigraphic Bounding. | Opinion |
| **37** | Primatomorpha | *Purgatorius* | 65.5 | 75.8 | (*119*) | | | Cladistic |
| | **Additional Calibrations for the 241 species tree** | | | | | | | |
| **38** | Hyracoidea | *Dendrohyrax* | 6.08 | 11.63 | (*101*, *132*, *145*) | | | Opinion |
| **39** | Vermilingua (*Myrmecophaga* + *Tamandua*) | *Protamandua* | 15.97 | 56 | | | | Cladistic |

| | Clade | Oldest crown fossil | Minimu m age | Maximum age | References for oldest fossil(s) and discussion of calibration | Comments on Minimum Constraint | Comments on Maximum Constraint | Type |
|---|---|---|---|---|---|---|---|---|
| **40** | Herpestidae + Eupleridae + Hyaenidae | *Protictitherium* | 15.97 | 27.82 | | | | Opinion |
| **41** | Odobenidae + Otariidae | *Proneotherium* | 16.6 | 28.1 | (*146*) | | | Cladistic |
| **42** | Pinnipedia | *Desmatophoca brachycephala* | 20.44 | 33.9 | (*147*) | | | Cladistic |
| **43** | Craseonycteridae + Megadermatidae | *Saharaderma* | 33.9 | 47.8 | (*101*, *112*) | | Stratigraphic Bounding | Opinion |
| **44** | Giraffidae to Antilocapridae | *Canthumeryx* | 17.8 | 28.1 | | | | Opinion |
| **45** | Phocoenidae to Monodontidae | Pterophocaena nishinoi | 9.2 | 19.5 | (*148*, *149*) | Maximum follows McGowen et al. (*148*) and is based on the oldest known stem delphinoid (*Kentriodon pernix*) based on the Geisler et al. (*149*) analysis with constraints. | Opinion | |
| **46** | Papionini | *Macaca libyca* | 5.5 | 11.63 | (*135*) | | | Opinion |
| **47** | Hominoidea | *Sivapithecus* | 12.68 | 27 | (*135*, *150*) | The maximum age assumes that crown Hominidae is unlikely to be older than the stem hominoid *Kamoyapitheus* (*151*). The maximum age for this genus is 27.8 million years (*152*). | | Cladistic |
| **48** | *Homo* + *Pan* | *Ardipithecus ramidus kadabba* | 5.11 | 7.246 | (*87*, *135*) | | | Cladistic |
| **49** | Galagidae + Lorisidae | *Saharagalago misrensis* | 38 | 56 | (*101*, *140*) | | | Cladistic |
| **50** | Sciuromorpha to Hystricognathi | *Acritoparamys* | 56 | 66 | (*101*, *153–155*) | | | Opinion |
| **51** | Aplodontidae + Sciuridae | *Spurimus* | 45.7 | 59.2 | (*6*, *156*) | | | Opinion |
| **52** | Chinchillidae + Dinomyidae | *Eoviscaccia* | 24.5 | 38 | (*101*, *157*) | | | Opinion |

| | Clade | Oldest crown fossil | Minimum age | Maximum age | References for oldest fossil(s) and discussion of calibration | Comments on Minimum Constraint | Comments on Maximum Constraint | Type |
|---|---|---|---|---|---|---|---|---|
| **53** | Ctenomyidae + Octodontidae | *Xenodontomys, Palaeoctodon, Chasicomys* | 9.07 | 33.9 | (*101*, *157*, *158*) | | | Opinion |
| **54** | Castorimorpha + Anomaluromorpha + Myomorpha | *Erlianomys* | 54 | 66 | (*6*, *159*) | | | Opinion |
| **55** | Atelidae (*Ateles* + *Alouatta*) | *Stirtonia* | 12.8 | 20.44 | | First appearance of *Stirtonia* is at 12.8 Ma (*151*, *160*) | Stratigraphic Bounding. | Opinion |
| **56** | *Saimiri* to *Cebus* | *Neosaimiri fieldsi* | 11.8 | 20.44 | (*135*, *151*, *161*, *162*) | | Stratigraphic Bounding. | Opinion |
| **57** | Balaenopteroidea | *Plesiobalaenoptera* + *Incakujira anillodefuego* | 7.246 | 13.82 | (*163*) | The oldest fossils attributable to this node are *Plesiobalaenoptera quarantellii* from Northern Italy and *Incakujira anillodefuego* from Peru both from the Tortonian of the Late Miocene – yielding a minimum bound of 7.246 Ma. The paratype of *Incakujira* originates from the 'upper Isurus [Cosmopolitodus] zone', thought to most likely be the AGL level, which has been estimated age range of 7.5-7.3 Ma. Here we use the upper boundary of the Tortonian 7.246 Ma to reflect this uncertainty. | Stratigraphic Bounding. | Cladistic |
| **58** | *Rattus* + *Mus* | *Karnimata darwini* | 10.4 | 15.97 | (*73*) | | | Cladistic |
| **59** | Plancetalia (root) | *Eomaia, Sinodelphys* | | 131.5 | (*85-87*) | | An upper bound of 131.5Mya ('Benton2009 constraint') was placed on Placentalia (root) based on the well documented fossil species *Eomaia* (*85*) and *Sinodelphys* (*86*), estimated to be Aptian or | |

| | Clade | Oldest crown fossil | Minimum age | Maximum age | References for oldest fossil(s) and discussion of calibration | Comments on Minimum Constraint | Comments on Maximum Constraint | Type |
|---|---|---|---|---|---|---|---|---|
| **60** | Plancetalia (root) | | | 116.81 | (*6*) | | Barremian in age (*87*). A conservative estimate of 131.5Mya is used to reflect the uncertainty around the dating of the horizons from which these fossils are derived (*87*). The upper 95% confidence interval from (*6*) was used to place an upper bound on Placentalia (root). | |

**Table S10.**

Average divergence time estimates and 95% CI for sliding windows analyses when varying the underlying dataset and the constraint on the root. The constraint used to calibrate the Benton analysis was 131.5Mya (*87*). The upper 95% confidence intervals from another analysis also used to constrain the analysis, 116.81Mya from the Meredith analysis (*6*). Where the dataset analyzed contained 316 trees additional summary statistics are provided including minimum (min) and maximum (max) values, the standard deviation (std dev) around the average of 316 point estimates (average) as well as corresponding statistics for the lower 95% confidence interval (L95 CI) and upper 95% confidence interval (U95 CI).

| **Analysis** | **Model** | **Number of Trees** | **node** | **average** | **L95 CI mean** | **U95 CI mean** | **std dev** | **mean min** | **mean max** | **L95 CI min** | **U95 CI max** | **L95 CI median** | **U95 CI median** |
|---|---|---|---|---|---|---|---|---|---|---|---|---|---|
| 10% Missingness | IRM | 316 | Afroinsectiphilia | 66.86 | 57.46 | 78.62 | 1.70 | 63.24 | 75.12 | 53.26 | 87.57 | 57.49 | 78.37 |
| Accelerated | IRM | 1 | Afroinsectiphilia | 67.52 | 58.00 | 79.19 | | | | | | | |
| Benton | IRM | 316 | Afroinsectiphilia | 66.08 | 56.15 | 79.01 | 1.44 | 62.43 | 72.11 | 52.10 | 85.36 | 56.21 | 79.09 |
| Benton | ARM | 316 | Afroinsectiphilia | 67.45 | 62.63 | 73.12 | 1.59 | 62.36 | 72.41 | 56.37 | 76.95 | 62.77 | 73.36 |
| Benton | Averaged IRM & ARM | 1 | Afroinsectiphilia | 66.76 | 59.39 | 76.07 | | | | | | | |
| Cladistic | ARM | 316 | Afroinsectiphilia | 67.66 | 63.02 | 72.89 | 1.36 | 62.17 | 71.83 | 55.73 | 77.50 | 63.31 | 72.81 |
| Conserved | IRM | 1 | Afroinsectiphilia | 68.29 | 56.02 | 97.05 | | | | | | | |
| Meredith | IRM | 316 | Afroinsectiphilia | 65.87 | 56.16 | 78.52 | 1.41 | 62.39 | 72.52 | 51.99 | 85.78 | 56.20 | 78.60 |
| Neutral | IRM | 1 | Afroinsectiphilia | 68.56 | 58.91 | 69.57 | | | | | | | |
| Neutral 241 species | IRM | 1 | Afroinsectiphilia | 66.30 | 56.18 | 78.96 | | | | | | | |

| Analysis | Model | Number of Trees | node | average | L95 CI mean | U95 CI mean | std dev | mean min | mean max | L95 CI min | U95 CI max | L95 CI median | U95 CI median |
|---|---|---|---|---|---|---|---|---|---|---|---|---|---|
| No DNA | ARM | 316 | Afroinsectiphilia | 62.42 | 38.79 | 86.74 | 0.06 | 62.16 | 62.64 | 37.60 | 87.95 | 38.78 | 86.77 |
| One Stratigraphic Bound | IRM | 316 | Afroinsectiphilia | 64.20 | 55.12 | 75.20 | 1.19 | 60.93 | 69.97 | 52.07 | 83.05 | 55.20 | 75.15 |
| Root | IRM | 1 | Afroinsectiphilia | 66.18 | 57.99 | 79.32 | | | | | | | |
| Root Calibration Only | ARM | 316 | Afroinsectiphilia | 71.26 | 51.55 | 89.61 | 4.64 | 57.71 | 82.07 | 31.43 | 97.99 | 51.65 | 89.97 |
| all Benton All6 Extended | ARM | 316 | Afroinsectiphilia | 69.14 | 63.72 | 75.52 | 1.70 | 62.51 | 76.06 | 56.94 | 84.88 | 63.96 | 75.54 |
| Benton All6 Extended | IRM | 316 | Afroinsectiphilia | 69.82 | 57.71 | 83.86 | 1.65 | 65.47 | 77.00 | 52.84 | 91.22 | 57.75 | 84.29 |
| Benton All6 Extended NoDNA | ARM | 316 | Afroinsectiphilia | 64.15 | 34.88 | 89.95 | 0.13 | 63.72 | 64.86 | 31.49 | 102.80 | 34.92 | 89.82 |
| all Benton All6 Extended NoDNA | IRM | 316 | Afroinsectiphilia | 64.14 | 34.85 | 89.92 | 0.08 | 63.64 | 64.80 | 31.64 | 92.88 | 34.92 | 89.95 |
| all Benton Rodentia Extended | ARM | 316 | Afroinsectiphilia | 68.19 | 62.87 | 74.64 | 1.67 | 62.69 | 74.16 | 56.19 | 80.06 | 62.91 | 74.76 |
| all Benton Rodentia Extended | IRM | 316 | Afroinsectiphilia | 68.41 | 56.83 | 82.65 | 1.81 | 64.06 | 76.23 | 53.10 | 90.27 | 56.86 | 82.83 |
| all Benton Rodentia Extended NoDNA | ARM | 316 | Afroinsectiphilia | 63.92 | 35.72 | 94.34 | 0.22 | 63.20 | 65.62 | 25.11 | 107.65 | 35.85 | 94.09 |
| all Benton Rodentia Extended NoDNA | IRM | 316 | Afroinsectiphilia | 63.90 | 35.91 | 94.00 | 0.14 | 63.51 | 65.55 | 31.81 | 98.68 | 35.85 | 94.06 |
| 10% Missingness | IRM | 316 | Afroinsectivora | 64.09 | 54.32 | 76.34 | 1.84 | 57.94 | 68.92 | 44.03 | 80.04 | 54.46 | 76.57 |
| Accelerated | IRM | 1 | Afroinsectivora | NA | NA | NA | | | | | | | |
| Benton | IRM | 316 | Afroinsectivora | 63.16 | 52.85 | 76.42 | 1.73 | 51.89 | 67.13 | 22.42 | 81.49 | 53.10 | 76.75 |

| Analysis | Model | Number of Trees | node | average | L95 CI mean | U95 CI mean | std dev | mean min | mean max | L95 CI min | U95 CI max | L95 CI median | U95 CI median |
|---|---|---|---|---|---|---|---|---|---|---|---|---|---|
| Benton | ARM | 316 | Afroinsectivora | 65.93 | 61.01 | 71.71 | 2.00 | 53.04 | 70.91 | 22.73 | 75.59 | 61.43 | 72.00 |
| Benton | Averaged IRM & ARM | 1 | Afroinsectivora | 64.55 | 56.93 | 74.07 | | | | | | | |
| Cladistic | ARM | 316 | Afroinsectivora | 66.11 | 61.35 | 71.42 | 1.81 | 52.03 | 69.89 | 25.45 | 75.54 | 61.94 | 71.48 |
| Conserved | IRM | 1 | Afroinsectivora | NA | NA | NA | | | | | | | |
| Meredith | IRM | 316 | Afroinsectivora | 62.96 | 52.83 | 75.96 | 1.73 | 51.95 | 67.29 | 24.04 | 82.80 | 53.11 | 76.40 |
| Neutral | IRM | 1 | Afroinsectivora | NA | NA | NA | | | | | | | |
| Neutral 241 species | IRM | 1 | Afroinsectivora | NA | NA | NA | | | | | | | |
| No DNA | ARM | 316 | Afroinsectivora | 51.08 | 24.60 | 68.51 | 0.08 | 50.60 | 51.72 | 23.39 | 70.78 | 24.62 | 68.50 |
| One Stratigraphic Bound | IRM | 316 | Afroinsectivora | 61.29 | 51.97 | 72.78 | 1.51 | 50.57 | 64.84 | 23.17 | 77.65 | 52.38 | 72.94 |
| Root | IRM | 1 | Afroinsectivora | NA | NA | NA | | | | | | | |
| Root Calibration Only | ARM | 316 | Afroinsectivora | 69.45 | 50.03 | 87.61 | 4.97 | 48.10 | 81.14 | 10.18 | 96.18 | 50.39 | 88.09 |
| all Benton All6 Extended | ARM | 316 | Afroinsectivora | 67.53 | 61.99 | 72.25 | 2.08 | 53.08 | 73.38 | 23.42 | 84.24 | 62.60 | 73.12 |
| Benton All6 Extended | IRM | 316 | Afroinsectivora | 66.77 | 54.30 | 81.04 | 1.94 | 53.20 | 71.81 | 26.32 | 87.37 | 54.44 | 81.40 |
| Benton All6 Extended NoDNA | ARM | 316 | Afroinsectivora | 51.52 | 26.14 | 73.21 | 0.09 | 51.14 | 52.12 | 24.51 | 74.29 | 26.15 | 73.20 |
| all Benton All6 Extended NoDNA | IRM | 316 | Afroinsectivora | 51.51 | 26.11 | 73.21 | 0.06 | 51.29 | 51.88 | 25.45 | 73.90 | 26.12 | 73.22 |

| Analysis | Model | Number of Trees | node | average | L95 CI mean | U95 CI mean | std dev | mean min | mean max | L95 CI min | U95 CI max | L95 CI median | U95 CI median |
|---|---|---|---|---|---|---|---|---|---|---|---|---|---|
| all Benton Rodentia Extended | ARM | 316 | Afroinsectivora | 66.64 | 61.22 | 73.14 | 2.04 | 53.12 | 72.05 | 23.78 | 77.82 | 61.62 | 73.27 |
| all Benton Rodentia Extended | IRM | 316 | Afroinsectivora | 65.40 | 53.53 | 79.93 | 2.03 | 52.78 | 70.96 | 26.27 | 86.10 | 53.62 | 80.10 |
| all Benton Rodentia Extended NoDNA | ARM | 316 | Afroinsectivora | 51.45 | 25.25 | 71.53 | 0.09 | 51.01 | 52.13 | 24.46 | 72.69 | 25.22 | 71.52 |
| all Benton Rodentia Extended NoDNA | IRM | 316 | Afroinsectivora | 51.46 | 25.29 | 71.54 | 0.23 | 51.11 | 55.04 | 23.41 | 73.66 | 25.26 | 71.55 |
| Body Size | IRM | 316 | Afroinsectivora | 62.58 | 48.98 | 77.06 | 1.84 | 54.97 | 70.37 | 31.91 | 89.16 | 49.22 | 76.96 |
| 10% Missingness | IRM | 316 | Afrosoricida | 54.54 | 39.98 | 66.64 | 2.08 | 48.14 | 60.63 | 32.32 | 76.45 | 40.21 | 66.44 |
| Accelerated | IRM | 1 | Afrosoricida | 55.73 | 39.45 | 67.75 | | | | | | | |
| Benton | IRM | 316 | Afrosoricida | 53.14 | 37.99 | 65.92 | 2.38 | 33.95 | 56.79 | 5.97 | 69.07 | 38.14 | 66.24 |
| Benton | ARM | 316 | Afrosoricida | 59.65 | 54.11 | 65.66 | 2.90 | 34.55 | 64.43 | 6.24 | 69.68 | 54.93 | 66.11 |
| Benton | Averaged IRM & ARM | 1 | Afrosoricida | 56.40 | 46.05 | 65.79 | | | | | | | |
| Cladistic | ARM | 316 | Afrosoricida | 59.75 | 54.32 | 65.61 | 2.85 | 33.07 | 64.49 | 2.46 | 69.38 | 54.98 | 66.03 |
| Conserved | IRM | 1 | Afrosoricida | 58.39 | 40.97 | 86.52 | | | | | | | |
| Meredith | IRM | 316 | Afrosoricida | 53.03 | 38.13 | 65.77 | 2.42 | 34.06 | 56.71 | 6.43 | 68.93 | 38.35 | 66.15 |
| Neutral | IRM | 1 | Afrosoricida | 57.20 | 37.83 | 68.23 | | | | | | | |
| Neutral 241 species | IRM | 1 | Afrosoricida | 56.30 | 42.98 | 67.11 | | | | | | | |

| Analysis | Model | Number of Trees | node | average | L95 CI mean | U95 CI mean | std dev | mean min | mean max | L95 CI min | U95 CI max | L95 CI median | U95 CI median |
|---|---|---|---|---|---|---|---|---|---|---|---|---|---|
| No DNA | ARM | 316 | Afrosoricida | 36.39 | 9.49 | 62.54 | 0.10 | 36.03 | 37.45 | 8.17 | 64.28 | 9.47 | 62.49 |
| One Stratigraphic Bound | IRM | 316 | Afrosoricida | 51.58 | 36.76 | 63.80 | 2.41 | 32.80 | 55.36 | 6.55 | 66.46 | 36.77 | 63.76 |
| Root | IRM | 1 | Afrosoricida | 56.39 | 40.90 | 68.08 | | | | | | | |
| Root Calibration Only | ARM | 316 | Afrosoricida | 62.36 | 44.43 | 79.40 | 5.43 | 23.62 | 73.52 | 0.00 | 89.95 | 44.80 | 80.11 |
| all Benton All6 Extended | ARM | 316 | Afrosoricida | 60.86 | 54.89 | 72.22 | 2.94 | 34.15 | 66.77 | 5.68 | 86.21 | 55.76 | 68.98 |
| Benton All6 Extended | IRM | 316 | Afrosoricida | 56.13 | 39.80 | 71.17 | 2.53 | 34.44 | 60.55 | 3.32 | 76.45 | 39.97 | 71.41 |
| Benton All6 Extended NoDNA | ARM | 316 | Afrosoricida | 36.07 | 8.53 | 65.09 | 0.09 | 35.75 | 36.54 | 7.27 | 66.11 | 8.53 | 65.08 |
| all Benton All6 Extended NoDNA | IRM | 316 | Afrosoricida | 36.06 | 8.54 | 65.12 | 0.07 | 35.86 | 36.61 | 7.31 | 66.34 | 8.50 | 65.11 |
| all Benton Rodentia Extended | ARM | 316 | Afrosoricida | 60.17 | 54.19 | 66.77 | 2.92 | 34.42 | 65.74 | 7.54 | 72.54 | 54.90 | 66.83 |
| all Benton Rodentia Extended | IRM | 316 | Afrosoricida | 55.07 | 38.44 | 69.13 | 2.54 | 34.27 | 59.46 | 4.04 | 76.87 | 38.49 | 68.72 |
| all Benton Rodentia Extended NoDNA | ARM | 316 | Afrosoricida | 36.31 | 9.41 | 63.89 | 0.09 | 35.97 | 36.96 | 7.94 | 65.56 | 9.43 | 63.93 |
| all Benton Rodentia Extended NoDNA | IRM | 316 | Afrosoricida | 36.32 | 9.38 | 63.87 | 0.15 | 36.02 | 38.52 | 7.78 | 65.70 | 9.43 | 63.89 |
| Body Size | IRM | 316 | Afrosoricida | 54.25 | 38.93 | 67.78 | 2.03 | 37.68 | 62.57 | 3.59 | 81.95 | 38.94 | 67.20 |
| 10% Missingness | IRM | 316 | Afrotheria | 73.35 | 63.67 | 83.13 | 1.82 | 66.70 | 81.22 | 61.63 | 96.72 | 63.61 | 82.97 |
| Accelerated | IRM | 1 | Afrotheria | 74.96 | 64.11 | 83.43 | | | | | | | |

| Analysis | Model | Number of Trees | node | average | L95 CI mean | U95 CI mean | std dev | mean min | mean max | L95 CI min | U95 CI max | L95 CI median | U95 CI median |
|---|---|---|---|---|---|---|---|---|---|---|---|---|---|
| Benton | IRM | 316 | Afrotheria | 74.46 | 63.88 | 85.32 | 1.83 | 69.34 | 82.37 | 62.23 | 97.02 | 63.76 | 85.03 |
| Benton | ARM | 316 | Afrotheria | 71.40 | 65.98 | 77.24 | 1.34 | 67.39 | 77.27 | 63.42 | 85.60 | 65.59 | 76.90 |
| Benton | Averaged IRM & ARM | 1 | Afrotheria | 72.93 | 64.93 | 81.28 | | | | | | | |
| Cladistic | ARM | 316 | Afrotheria | 71.67 | 66.55 | 77.21 | 1.19 | 68.23 | 79.04 | 63.66 | 88.78 | 66.58 | 76.86 |
| Conserved | IRM | 1 | Afrotheria | 74.04 | 63.25 | 102.69 | | | | | | | |
| Meredith | IRM | 316 | Afrotheria | 74.13 | 63.85 | 84.63 | 1.72 | 69.05 | 80.37 | 62.34 | 93.28 | 63.75 | 84.51 |
| Neutral | IRM | 1 | Afrotheria | 75.29 | 63.07 | 66.43 | | | | | | | |
| Neutral 241 species | IRM | 1 | Afrotheria | 73.59 | 63.24 | 84.92 | | | | | | | |
| No DNA | ARM | 316 | Afrotheria | 78.12 | 59.67 | 103.07 | 0.16 | 77.07 | 78.50 | 59.27 | 103.99 | 59.67 | 103.10 |
| One Stratigraphic Bound | IRM | 316 | Afrotheria | 72.15 | 63.43 | 81.86 | 1.53 | 68.34 | 79.95 | 61.27 | 94.30 | 63.35 | 81.48 |
| Root | IRM | 1 | Afrotheria | 72.96 | 63.57 | 82.01 | | | | | | | |
| Root Calibration Only | ARM | 316 | Afrotheria | 75.93 | 55.24 | 94.98 | 4.58 | 61.76 | 86.53 | 33.30 | 104.91 | 55.34 | 95.06 |
| all Benton All6 Extended | ARM | 316 | Afrotheria | 73.38 | 67.57 | 90.41 | 1.64 | 70.26 | 80.88 | 65.08 | 121.87 | 67.44 | 81.74 |
| Benton All6 Extended | IRM | 316 | Afrotheria | 77.58 | 66.45 | 90.38 | 2.08 | 72.25 | 85.27 | 63.77 | 100.41 | 66.25 | 90.35 |
| Benton All6 Extended NoDNA | ARM | 316 | Afrotheria | 79.94 | 60.66 | 113.61 | 0.37 | 78.03 | 82.90 | 60.05 | 124.07 | 60.60 | 113.53 |

| Analysis | Model | Number of Trees | node | average | L95 CI mean | U95 CI mean | std dev | mean min | mean max | L95 CI min | U95 CI max | L95 CI median | U95 CI median |
|---|---|---|---|---|---|---|---|---|---|---|---|---|---|
| all Benton All6 Extended NoDNA | IRM | 316 | Afrotheria | 79.95 | 60.59 | 113.65 | 0.24 | 77.48 | 81.30 | 59.99 | 121.21 | 60.60 | 113.59 |
| all Benton Rodentia Extended | ARM | 316 | Afrotheria | 72.27 | 66.41 | 79.03 | 1.48 | 68.90 | 78.37 | 64.04 | 87.61 | 66.06 | 78.87 |
| all Benton Rodentia Extended | IRM | 316 | Afrotheria | 76.41 | 64.99 | 88.78 | 2.07 | 70.55 | 83.92 | 62.75 | 98.59 | 64.77 | 88.69 |
| all Benton Rodentia Extended NoDNA | ARM | 316 | Afrotheria | 81.95 | 60.10 | 115.79 | 0.80 | 80.39 | 88.51 | 59.31 | 151.60 | 60.03 | 115.32 |
| all Benton Rodentia Extended NoDNA | IRM | 316 | Afrotheria | 81.80 | 60.11 | 115.44 | 0.41 | 77.02 | 84.46 | 59.58 | 130.75 | 60.05 | 115.26 |
| Body Size | IRM | 316 | Afrotheria | 67.16 | 53.64 | 81.63 | 2.29 | 59.92 | 76.88 | 44.59 | 95.64 | 53.77 | 81.24 |
| 10% Missingness | IRM | 316 | Atlantogenata | 91.67 | 77.21 | 107.06 | 3.49 | 82.56 | 108.25 | 72.21 | 126.92 | 76.78 | 106.39 |
| Accelerated | IRM | 1 | Atlantogenata | 93.17 | 80.36 | 109.59 | | | | | | | |
| Benton | IRM | 316 | Atlantogenata | 96.07 | 79.80 | 113.87 | 3.04 | 90.13 | 107.75 | 74.46 | 126.32 | 79.49 | 113.57 |
| Benton | ARM | 316 | Atlantogenata | 91.72 | 81.25 | 102.52 | 2.77 | 81.54 | 102.45 | 74.67 | 119.13 | 80.98 | 101.86 |
| Benton | Averaged IRM & ARM | 1 | Atlantogenata | 93.90 | 80.52 | 108.19 | | | | | | | |
| Cladistic | ARM | 316 | Atlantogenata | 92.90 | 82.18 | 104.08 | 2.98 | 84.84 | 104.72 | 76.91 | 122.39 | 81.86 | 103.07 |
| Conserved | IRM | 1 | Atlantogenata | 90.26 | 74.46 | 117.23 | | | | | | | |
| Meredith | IRM | 316 | Atlantogenata | 94.63 | 80.09 | 110.08 | 2.12 | 89.42 | 101.41 | 75.18 | 116.23 | 79.90 | 110.21 |
| Neutral | IRM | 1 | Atlantogenata | 94.73 | 79.70 | 112.40 | | | | | | | |

| Analysis | Model | Number of Trees | node | average | L95 CI mean | U95 CI mean | std dev | mean min | mean max | L95 CI min | U95 CI max | L95 CI median | U95 CI median |
|---|---|---|---|---|---|---|---|---|---|---|---|---|---|
| Neutral 241 species | IRM | 1 | Atlantogenata | 94.31 | 77.70 | 111.26 | | | | | | | |
| No DNA | ARM | 316 | Atlantogenata | 90.39 | 63.78 | 121.18 | 0.37 | 87.46 | 91.09 | 63.16 | 122.32 | 63.78 | 121.19 |
| One Stratigraphic Bound | IRM | 316 | Atlantogenata | 92.22 | 77.40 | 108.58 | 3.10 | 87.19 | 105.65 | 73.13 | 125.06 | 77.06 | 107.99 |
| Root | IRM | 1 | Atlantogenata | 91.71 | 75.06 | 107.89 | | | | | | | |
| Root Calibration Only | ARM | 316 | Atlantogenata | 104.16 | 78.74 | 125.27 | 5.38 | 86.09 | 115.12 | 47.69 | 134.68 | 79.67 | 125.68 |
| all Benton All6 Extended | ARM | 316 | Atlantogenata | 96.93 | 85.87 | 108.81 | 3.04 | 87.80 | 107.41 | 80.66 | 123.44 | 85.55 | 107.92 |
| Benton All6 Extended | IRM | 316 | Atlantogenata | 102.10 | 84.87 | 120.45 | 2.69 | 96.91 | 111.15 | 80.56 | 129.25 | 84.48 | 120.43 |
| Benton All6 Extended NoDNA | ARM | 316 | Atlantogenata | 94.33 | 66.41 | 127.01 | 0.67 | 91.19 | 101.04 | 64.98 | 133.60 | 66.31 | 126.92 |
| all Benton All6 Extended NoDNA | IRM | 316 | Atlantogenata | 94.34 | 66.37 | 127.11 | 0.44 | 88.96 | 96.28 | 65.02 | 130.30 | 66.35 | 127.01 |
| all Benton Rodentia Extended | ARM | 316 | Atlantogenata | 94.08 | 82.72 | 106.51 | 2.95 | 83.94 | 105.19 | 76.74 | 122.18 | 82.56 | 105.97 |
| all Benton Rodentia Extended | IRM | 316 | Atlantogenata | 100.02 | 82.92 | 118.44 | 2.84 | 93.58 | 109.82 | 78.94 | 135.29 | 82.57 | 118.36 |
| all Benton Rodentia Extended NoDNA | ARM | 316 | Atlantogenata | 96.06 | 64.95 | 127.28 | 1.02 | 94.04 | 105.38 | 60.96 | 154.03 | 64.89 | 127.05 |
| all Benton Rodentia Extended NoDNA | IRM | 316 | Atlantogenata | 95.84 | 64.89 | 127.05 | 0.55 | 91.42 | 99.73 | 64.14 | 131.72 | 64.85 | 126.95 |
| Body Size | IRM | 316 | Atlantogenata | 86.89 | 72.89 | 101.83 | 3.53 | 81.35 | 105.01 | 69.35 | 125.87 | 72.55 | 100.92 |
| 10% Missingness | IRM | 316 | Boreoeutheria | 95.07 | 84.26 | 106.94 | 4.04 | 86.15 | 113.38 | 78.39 | 128.50 | 83.71 | 106.38 |

| Analysis | Model | Number of Trees | node | average | L95 CI mean | U95 CI mean | std dev | mean min | mean max | L95 CI min | U95 CI max | L95 CI median | U95 CI median |
|---|---|---|---|---|---|---|---|---|---|---|---|---|---|
| Accelerated | IRM | 1 | Boreoeutheria | 97.14 | 87.01 | 109.25 | | | | | | | |
| Benton | IRM | 316 | Boreoeutheria | 102.04 | 89.24 | 115.96 | 4.54 | 94.11 | 120.22 | 83.14 | 133.02 | 88.43 | 115.31 |
| Benton | ARM | 316 | Boreoeutheria | 89.53 | 83.73 | 95.88 | 2.97 | 81.18 | 104.38 | 77.39 | 117.45 | 83.31 | 94.82 |
| Benton | Averaged IRM & ARM | 1 | Boreoeutheria | 95.79 | 86.48 | 105.92 | | | | | | | |
| Cladistic | ARM | 316 | Boreoeutheria | 90.76 | 83.18 | 99.10 | 4.35 | 84.07 | 113.49 | 78.39 | 127.48 | 82.50 | 97.81 |
| Conserved | IRM | 1 | Boreoeutheria | 93.45 | 82.06 | 107.60 | | | | | | | |
| Meredith | IRM | 316 | Boreoeutheria | 100.73 | 89.43 | 112.55 | 3.51 | 92.76 | 113.27 | 83.03 | 123.11 | 88.83 | 112.47 |
| Neutral | IRM | 1 | Boreoeutheria | 99.64 | 88.79 | 115.48 | | | | | | | |
| Neutral 241 species | IRM | 1 | Boreoeutheria | 99.26 | 88.12 | 111.85 | | | | | | | |
| No DNA | ARM | 316 | Boreoeutheria | 101.50 | 74.03 | 129.70 | 0.62 | 96.61 | 102.89 | 66.40 | 131.42 | 74.17 | 129.77 |
| One Stratigraphic Bound | IRM | 316 | Boreoeutheria | 97.28 | 86.09 | 109.71 | 4.65 | 89.62 | 117.01 | 80.30 | 130.52 | 85.43 | 108.71 |
| Root | IRM | 1 | Boreoeutheria | 98.07 | 85.17 | 109.94 | | | | | | | |
| Root Calibration Only | ARM | 316 | Boreoeutheria | 114.10 | 87.52 | 135.27 | 5.97 | 93.79 | 126.15 | 53.16 | 146.12 | 88.80 | 135.33 |
| all Benton All6 Extended | ARM | 316 | Boreoeutheria | 99.85 | 92.17 | 92.49 | 4.24 | 92.04 | 117.89 | 86.06 | 129.09 | 91.42 | 104.30 |
| Benton All6 Extended | IRM | 316 | Boreoeutheria | 110.03 | 97.65 | 123.22 | 4.17 | 102.08 | 124.85 | 91.40 | 136.46 | 96.85 | 122.92 |

| Analysis | Model | Number of Trees | node | average | L95 CI mean | U95 CI mean | std dev | mean min | mean max | L95 CI min | U95 CI max | L95 CI median | U95 CI median |
|---|---|---|---|---|---|---|---|---|---|---|---|---|---|
| Benton All6 Extended NoDNA | ARM | 316 | Boreoeutheria | 108.74 | 79.70 | 133.23 | 0.89 | 103.40 | 118.58 | 74.75 | 135.16 | 79.72 | 133.27 |
| all Benton All6 Extended NoDNA | IRM | 316 | Boreoeutheria | 108.74 | 79.66 | 133.30 | 0.59 | 100.95 | 111.39 | 74.45 | 135.44 | 79.69 | 133.31 |
| all Benton Rodentia Extended | ARM | 316 | Boreoeutheria | 93.96 | 86.36 | 102.08 | 3.69 | 85.47 | 109.82 | 80.45 | 122.94 | 85.69 | 101.16 |
| all Benton Rodentia Extended | IRM | 316 | Boreoeutheria | 106.99 | 94.37 | 120.57 | 4.15 | 98.99 | 122.67 | 88.13 | 134.59 | 93.48 | 120.08 |
| all Benton Rodentia Extended NoDNA | ARM | 316 | Boreoeutheria | 108.35 | 78.71 | 133.35 | 0.99 | 105.12 | 116.57 | 72.58 | 155.85 | 78.68 | 133.26 |
| all Benton Rodentia Extended NoDNA | IRM | 316 | Boreoeutheria | 108.15 | 78.57 | 133.28 | 0.60 | 102.09 | 111.87 | 75.08 | 137.39 | 78.60 | 133.25 |
| Body Size | IRM | 316 | Boreoeutheria | 92.41 | 83.23 | 102.47 | 4.21 | 86.68 | 112.85 | 78.93 | 127.03 | 82.45 | 101.56 |
| 10% Missingness | IRM | 316 | Carnivora | 55.27 | 45.24 | 64.92 | 1.87 | 48.19 | 59.26 | 39.57 | 66.59 | 44.99 | 65.13 |
| Accelerated | IRM | 1 | Carnivora | 55.05 | 44.41 | 64.90 | | | | | | | |
| Benton | IRM | 316 | Carnivora | 55.66 | 45.54 | 65.29 | 1.41 | 48.77 | 59.28 | 39.91 | 66.63 | 45.28 | 65.34 |
| Benton | ARM | 316 | Carnivora | 51.47 | 45.67 | 57.19 | 1.22 | 47.64 | 55.38 | 42.85 | 61.53 | 45.58 | 57.18 |
| Benton | Averaged IRM & ARM | 1 | Carnivora | 53.56 | 45.61 | 61.24 | | | | | | | |
| Cladistic | ARM | 316 | Carnivora | 51.82 | 46.16 | 57.69 | 1.16 | 48.64 | 56.50 | 43.35 | 64.65 | 46.04 | 57.71 |
| Conserved | IRM | 1 | Carnivora | 54.62 | 44.71 | 63.88 | | | | | | | |
| Meredith | IRM | 316 | Carnivora | 55.56 | 45.46 | 65.22 | 1.41 | 48.68 | 59.23 | 39.84 | 66.61 | 45.12 | 65.22 |

| Analysis | Model | Number of Trees | node | average | L95 CI mean | U95 CI mean | std dev | mean min | mean max | L95 CI min | U95 CI max | L95 CI median | U95 CI median |
|---|---|---|---|---|---|---|---|---|---|---|---|---|---|
| Neutral | IRM | 1 | Carnivora | 55.42 | 45.64 | 64.45 | | | | | | | |
| Neutral 241 species | IRM | 1 | Carnivora | 54.99 | 46.13 | 64.01 | | | | | | | |
| No DNA | ARM | 316 | Carnivora | 56.36 | 43.81 | 66.58 | 0.03 | 56.30 | 56.70 | 43.22 | 66.73 | 43.81 | 66.58 |
| One Stratigraphic Bound | IRM | 316 | Carnivora | 52.42 | 43.38 | 61.38 | 1.08 | 45.84 | 55.17 | 39.36 | 63.86 | 43.26 | 61.41 |
| Root | IRM | 1 | Carnivora | 55.55 | 45.42 | 64.89 | | | | | | | |
| Root Calibration Only | ARM | 316 | Carnivora | 58.46 | 41.63 | 74.60 | 4.19 | 46.05 | 72.35 | 25.11 | 87.63 | 41.80 | 74.53 |
| all Benton All6 Extended | ARM | 316 | Carnivora | 53.86 | 47.03 | 64.65 | 1.49 | 49.70 | 59.27 | 43.73 | 76.28 | 46.83 | 61.07 |
| Benton All6 Extended | IRM | 316 | Carnivora | 56.46 | 46.64 | 66.24 | 1.58 | 49.21 | 60.25 | 40.25 | 67.19 | 46.40 | 66.32 |
| Benton All6 Extended NoDNA | ARM | 316 | Carnivora | 56.50 | 43.80 | 66.76 | 0.04 | 55.85 | 56.66 | 43.62 | 66.92 | 43.80 | 66.76 |
| all Benton All6 Extended NoDNA | IRM | 316 | Carnivora | 56.50 | 43.80 | 66.77 | 0.03 | 56.10 | 56.57 | 43.14 | 66.93 | 43.80 | 66.77 |
| all Benton Rodentia Extended | ARM | 316 | Carnivora | 51.83 | 45.83 | 57.73 | 1.26 | 47.68 | 55.95 | 42.83 | 62.99 | 45.74 | 57.76 |
| all Benton Rodentia Extended | IRM | 316 | Carnivora | 55.53 | 45.71 | 65.49 | 1.57 | 48.42 | 59.64 | 39.95 | 66.96 | 45.45 | 65.74 |
| all Benton Rodentia Extended NoDNA | ARM | 316 | Carnivora | 56.29 | 43.74 | 66.59 | 0.02 | 56.23 | 56.36 | 43.58 | 66.76 | 43.73 | 66.59 |
| all Benton Rodentia Extended NoDNA | IRM | 316 | Carnivora | 56.29 | 43.74 | 66.59 | 0.02 | 56.02 | 56.32 | 43.57 | 66.72 | 43.74 | 66.59 |
| 10% Missingness | IRM | 316 | Cetartiodactyla | 64.63 | 61.58 | 66.94 | 0.24 | 63.37 | 65.30 | 59.50 | 67.28 | 61.57 | 66.94 |

| **Analysis** | **Model** | **Number of Trees** | **node** | **average** | **L95 CI mean** | **U95 CI mean** | **std dev** | **mean min** | **mean max** | **L95 CI min** | **U95 CI max** | **L95 CI median** | **U95 CI median** |
|---|---|---|---|---|---|---|---|---|---|---|---|---|---|
| Accelerated | IRM | 1 | Cetartiodactyla | 64.79 | 62.01 | 66.98 | | | | | | | |
| Benton | IRM | 316 | Cetartiodactyla | 64.80 | 61.88 | 67.01 | 0.18 | 64.30 | 65.30 | 61.03 | 67.32 | 61.87 | 67.00 |
| Benton | ARM | 316 | Cetartiodactyla | 64.84 | 62.75 | 66.76 | 0.48 | 63.25 | 66.48 | 61.11 | 68.35 | 62.73 | 66.73 |
| Benton | Averaged IRM & ARM | 1 | Cetartiodactyla | 64.82 | 62.32 | 66.88 | | | | | | | |
| Cladistic | ARM | 316 | Cetartiodactyla | 65.72 | 61.21 | 70.53 | 1.77 | 62.61 | 73.14 | 58.98 | 80.57 | 60.95 | 70.09 |
| Conserved | IRM | 1 | Cetartiodactyla | 64.59 | 61.43 | 66.98 | | | | | | | |
| Meredith | IRM | 316 | Cetartiodactyla | 64.79 | 61.87 | 67.01 | 0.18 | 64.28 | 65.26 | 61.01 | 67.28 | 61.85 | 67.00 |
| Neutral | IRM | 1 | Cetartiodactyla | 64.89 | 61.90 | 67.07 | | | | | | | |
| Neutral 241 species | IRM | 1 | Cetartiodactyla | 61.76 | 58.33 | 64.88 | | | | | | | |
| No DNA | ARM | 316 | Cetartiodactyla | 63.57 | 58.68 | 66.89 | 0.02 | 63.49 | 63.72 | 58.50 | 67.02 | 58.68 | 66.88 |
| One Stratigraphic Bound | IRM | 316 | Cetartiodactyla | 61.07 | 59.33 | 62.40 | 0.08 | 60.84 | 61.32 | 58.81 | 62.62 | 59.32 | 62.40 |
| Root | IRM | 1 | Cetartiodactyla | 64.78 | 61.92 | 67.02 | | | | | | | |
| Root Calibration Only | ARM | 316 | Cetartiodactyla | 74.70 | 55.98 | 91.30 | 4.45 | 61.58 | 85.55 | 33.93 | 106.27 | 56.50 | 91.47 |
| all Benton All6 Extended | ARM | 316 | Cetartiodactyla | 70.04 | 66.26 | 71.92 | 1.34 | 66.28 | 73.94 | 62.90 | 76.36 | 66.09 | 72.93 |
| Benton All6 Extended | IRM | 316 | Cetartiodactyla | 71.60 | 66.29 | 76.02 | 0.53 | 70.01 | 73.25 | 63.93 | 76.71 | 66.31 | 76.01 |

| Analysis | Model | Number of Trees | node | average | L95 CI mean | U95 CI mean | std dev | mean min | mean max | L95 CI min | U95 CI max | L95 CI median | U95 CI median |
|---|---|---|---|---|---|---|---|---|---|---|---|---|---|
| Benton All6 Extended NoDNA | ARM | 316 | Cetartiodactyla | 70.77 | 62.97 | 76.29 | 0.03 | 70.60 | 70.99 | 62.74 | 76.47 | 62.97 | 76.29 |
| all Benton All6 Extended NoDNA | IRM | 316 | Cetartiodactyla | 70.77 | 62.97 | 76.30 | 0.02 | 70.58 | 70.85 | 61.85 | 76.46 | 62.97 | 76.30 |
| all Benton Rodentia Extended | ARM | 316 | Cetartiodactyla | 65.36 | 63.29 | 67.15 | 0.38 | 64.02 | 66.71 | 61.61 | 68.77 | 63.28 | 67.11 |
| all Benton Rodentia Extended | IRM | 316 | Cetartiodactyla | 65.02 | 62.13 | 67.16 | 0.18 | 64.44 | 65.51 | 60.90 | 67.43 | 62.12 | 67.15 |
| all Benton Rodentia Extended NoDNA | ARM | 316 | Cetartiodactyla | 63.71 | 58.82 | 66.94 | 0.02 | 63.42 | 63.76 | 58.29 | 67.05 | 58.82 | 66.94 |
| all Benton Rodentia Extended NoDNA | IRM | 316 | Cetartiodactyla | 63.72 | 58.85 | 66.94 | 0.13 | 63.69 | 65.98 | 58.70 | 67.17 | 58.83 | 66.94 |
| 10% Missingness | IRM | 316 | Chiroptera | 63.81 | 59.32 | 67.22 | 0.40 | 62.44 | 65.15 | 57.17 | 67.88 | 59.20 | 67.20 |
| Accelerated | IRM | 1 | Chiroptera | 63.82 | 59.19 | 67.09 | | | | | | | |
| Benton | IRM | 316 | Chiroptera | 63.91 | 59.49 | 67.27 | 0.36 | 63.12 | 65.15 | 58.06 | 67.82 | 59.37 | 67.24 |
| Benton | ARM | 316 | Chiroptera | 64.86 | 61.86 | 67.26 | 0.54 | 61.62 | 66.35 | 58.56 | 69.23 | 61.87 | 67.25 |
| Benton | Averaged IRM & ARM | 1 | Chiroptera | 64.38 | 60.68 | 67.26 | | | | | | | |
| Cladistic | ARM | 316 | Chiroptera | 66.28 | 60.12 | 72.41 | 2.32 | 58.68 | 79.32 | 51.30 | 89.50 | 59.76 | 71.89 |
| Conserved | IRM | 1 | Chiroptera | 63.64 | 58.74 | 67.21 | | | | | | | |
| Meredith | IRM | 316 | Chiroptera | 63.90 | 59.47 | 67.26 | 0.37 | 62.88 | 65.05 | 57.59 | 67.79 | 59.37 | 67.24 |
| Neutral | IRM | 1 | Chiroptera | 63.97 | 59.48 | 67.45 | | | | | | | |

| Analysis | Model | Number of Trees | node | average | L95 CI mean | U95 CI mean | std dev | mean min | mean max | L95 CI min | U95 CI max | L95 CI median | U95 CI median |
|---|---|---|---|---|---|---|---|---|---|---|---|---|---|
| Neutral 241 species | IRM | 1 | Chiroptera | 63.90 | 59.07 | 67.38 | | | | | | | |
| No DNA | ARM | 316 | Chiroptera | 62.30 | 56.04 | 67.02 | 0.03 | 62.25 | 62.75 | 55.90 | 67.19 | 56.04 | 67.02 |
| One Stratigraphic Bound | IRM | 316 | Chiroptera | 63.35 | 58.51 | 66.99 | 0.50 | 62.09 | 64.91 | 56.68 | 67.71 | 58.36 | 66.93 |
| Root | IRM | 1 | Chiroptera | 64.15 | 59.99 | 67.33 | | | | | | | |
| Root Calibration Only | ARM | 316 | Chiroptera | 79.96 | 60.26 | 97.19 | 4.51 | 66.04 | 92.74 | 36.48 | 109.13 | 61.06 | 97.46 |
| all Benton All6 Extended | ARM | 316 | Chiroptera | 65.74 | 63.19 | 75.54 | 0.34 | 64.48 | 69.02 | 61.34 | 105.58 | 63.19 | 68.32 |
| Benton All6 Extended | IRM | 316 | Chiroptera | 64.49 | 60.18 | 67.54 | 0.33 | 63.61 | 65.48 | 58.30 | 68.14 | 60.08 | 67.52 |
| Benton All6 Extended NoDNA | ARM | 316 | Chiroptera | 62.33 | 55.51 | 67.11 | 0.01 | 62.29 | 62.44 | 55.34 | 67.22 | 55.52 | 67.11 |
| all Benton All6 Extended NoDNA | IRM | 316 | Chiroptera | 62.33 | 55.52 | 67.11 | 0.01 | 62.21 | 62.42 | 54.97 | 67.23 | 55.52 | 67.11 |
| all Benton Rodentia Extended | ARM | 316 | Chiroptera | 65.17 | 62.11 | 67.63 | 0.46 | 62.61 | 66.56 | 59.22 | 69.48 | 62.08 | 67.63 |
| all Benton Rodentia Extended | IRM | 316 | Chiroptera | 64.22 | 59.78 | 67.46 | 0.37 | 63.28 | 65.34 | 57.87 | 67.98 | 59.67 | 67.44 |
| all Benton Rodentia Extended NoDNA | ARM | 316 | Chiroptera | 62.36 | 55.89 | 67.07 | 0.02 | 62.14 | 62.41 | 55.75 | 67.18 | 55.89 | 67.07 |
| all Benton Rodentia Extended NoDNA | IRM | 316 | Chiroptera | 62.36 | 55.88 | 67.06 | 0.03 | 62.34 | 62.95 | 55.60 | 67.27 | 55.88 | 67.06 |
| Body Size | IRM | 316 | Chiroptera | 63.12 | 58.05 | 67.00 | 0.66 | 61.62 | 65.09 | 55.90 | 67.79 | 57.77 | 66.94 |
| 10% Missingness | IRM | 316 | Euarchonta | 77.29 | 71.58 | 83.29 | 2.32 | 74.11 | 94.83 | 67.26 | 107.17 | 71.46 | 82.19 |

| Analysis | Model | Number of Trees | node | average | L95 CI mean | U95 CI mean | std dev | mean min | mean max | L95 CI min | U95 CI max | L95 CI median | U95 CI median |
|---|---|---|---|---|---|---|---|---|---|---|---|---|---|
| Accelerated | IRM | 1 | Euarchonta | 77.69 | 72.21 | 83.75 | | | | | | | |
| Benton | IRM | 316 | Euarchonta | 79.37 | 72.05 | 87.96 | 2.67 | 75.60 | 96.62 | 68.33 | 109.50 | 71.80 | 87.18 |
| Benton | ARM | 316 | Euarchonta | 76.68 | 74.17 | 78.89 | 1.16 | 70.47 | 86.67 | 67.15 | 92.51 | 74.12 | 78.51 |
| Benton | Averaged IRM & ARM | 1 | Euarchonta | 78.02 | 73.11 | 83.42 | | | | | | | |
| Cladistic | ARM | 316 | Euarchonta | 76.77 | 73.98 | 79.19 | 1.45 | 71.47 | 88.84 | 68.01 | 95.55 | 73.93 | 78.68 |
| Conserved | IRM | 1 | Euarchonta | 77.83 | 72.00 | 85.62 | | | | | | | |
| Meredith | IRM | 316 | Euarchonta | 79.25 | 72.07 | 87.58 | 2.49 | 75.58 | 95.29 | 68.29 | 106.24 | 71.87 | 87.16 |
| Neutral | IRM | 1 | Euarchonta | 78.53 | 68.17 | 92.35 | | | | | | | |
| Neutral 241 species | IRM | 1 | Euarchonta | 79.43 | 73.39 | 88.64 | | | | | | | |
| No DNA | ARM | 316 | Euarchonta | 81.61 | 65.38 | 102.38 | 0.20 | 80.29 | 82.03 | 65.04 | 103.11 | 65.38 | 102.41 |
| One Stratigraphic Bound | IRM | 316 | Euarchonta | 78.77 | 71.04 | 86.84 | 2.30 | 74.86 | 94.03 | 67.39 | 106.27 | 70.82 | 86.06 |
| Root | IRM | 1 | Euarchonta | 77.40 | 72.44 | 82.59 | | | | | | | |
| Root Calibration Only | ARM | 316 | Euarchonta | 100.20 | 76.61 | 119.52 | 5.37 | 80.36 | 113.56 | 46.49 | 129.08 | 77.61 | 119.92 |
| all Benton All6 Extended | ARM | 316 | Euarchonta | 85.86 | 81.19 | 91.19 | 2.31 | 78.39 | 97.09 | 72.73 | 106.49 | 80.85 | 90.60 |
| Benton All6 Extended | IRM | 316 | Euarchonta | 93.22 | 82.60 | 104.47 | 2.38 | 83.38 | 106.75 | 70.31 | 119.05 | 82.38 | 104.29 |

| Analysis | Model | Number of Trees | node | average | L95 CI mean | U95 CI mean | std dev | mean min | mean max | L95 CI min | U95 CI max | L95 CI median | U95 CI median |
|---|---|---|---|---|---|---|---|---|---|---|---|---|---|
| Benton All6 Extended NoDNA | ARM | 316 | Euarchonta | 94.16 | 67.93 | 123.66 | 0.92 | 87.48 | 103.14 | 65.39 | 133.32 | 67.89 | 123.51 |
| all Benton All6 Extended NoDNA | IRM | 316 | Euarchonta | 94.17 | 67.83 | 123.74 | 0.59 | 87.13 | 97.05 | 64.78 | 130.06 | 67.79 | 123.62 |
| all Benton Rodentia Extended | ARM | 316 | Euarchonta | 81.18 | 76.02 | 86.69 | 2.20 | 71.80 | 90.20 | 68.07 | 98.36 | 75.78 | 86.21 |
| all Benton Rodentia Extended | IRM | 316 | Euarchonta | 90.20 | 79.16 | 101.78 | 2.26 | 76.66 | 103.07 | 68.43 | 115.99 | 79.32 | 101.65 |
| all Benton Rodentia Extended NoDNA | ARM | 316 | Euarchonta | 93.49 | 65.70 | 122.97 | 1.41 | 91.54 | 107.88 | 62.78 | 152.39 | 65.27 | 122.38 |
| all Benton Rodentia Extended NoDNA | IRM | 316 | Euarchonta | 93.24 | 65.49 | 122.55 | 0.76 | 90.03 | 100.30 | 64.19 | 130.85 | 65.23 | 122.26 |
| Body Size | IRM | 316 | Euarchonta | 76.63 | 71.62 | 81.32 | 1.72 | 72.67 | 89.00 | 67.17 | 97.87 | 71.44 | 80.45 |
| 10% Missingness | IRM | 316 | Euarchontoglires | 79.50 | 73.52 | 86.12 | 2.99 | 75.59 | 96.93 | 70.99 | 109.21 | 73.11 | 84.84 |
| Accelerated | IRM | 1 | Euarchontoglires | 80.17 | 73.85 | 86.55 | | | | | | | |
| Benton | IRM | 316 | Euarchontoglires | 83.31 | 75.40 | 92.56 | 4.22 | 77.11 | 99.42 | 71.54 | 112.46 | 74.66 | 91.58 |
| Benton | ARM | 316 | Euarchontoglires | 78.07 | 75.33 | 80.56 | 1.67 | 75.39 | 87.25 | 72.56 | 93.12 | 75.06 | 79.91 |
| Benton | Averaged IRM & ARM | 1 | Euarchontoglires | 80.69 | 75.36 | 86.56 | | | | | | | |
| Cladistic | ARM | 316 | Euarchontoglires | 78.25 | 75.13 | 81.05 | 2.15 | 75.04 | 90.42 | 72.17 | 97.43 | 74.82 | 80.20 |
| Conserved | IRM | 1 | Euarchontoglires | 79.78 | 73.95 | 87.86 | | | | | | | |
| Meredith | IRM | 316 | Euarchontoglires | 83.09 | 75.43 | 91.96 | 3.89 | 77.10 | 97.33 | 71.95 | 107.97 | 74.68 | 91.20 |

| Analysis | Model | Number of Trees | node | average | L95 CI mean | U95 CI mean | std dev | mean min | mean max | L95 CI min | U95 CI max | L95 CI median | U95 CI median |
|---|---|---|---|---|---|---|---|---|---|---|---|---|---|
| Neutral | IRM | 1 | Euarchontoglires | 80.89 | 70.13 | 93.67 | | | | | | | |
| Neutral 241 species | IRM | 1 | Euarchontoglires | 82.06 | 75.81 | 90.84 | | | | | | | |
| No DNA | ARM | 316 | Euarchontoglires | 92.16 | 69.43 | 119.06 | 0.41 | 89.03 | 93.05 | 66.03 | 120.61 | 69.50 | 119.06 |
| One Stratigraphic Bound | IRM | 316 | Euarchontoglires | 82.54 | 74.71 | 91.26 | 3.75 | 76.86 | 97.35 | 70.36 | 110.47 | 74.20 | 90.18 |
| Root | IRM | 1 | Euarchontoglires | 79.72 | 74.04 | 85.56 | | | | | | | |
| Root Calibration Only | ARM | 316 | Euarchontoglires | 101.83 | 77.89 | 121.35 | 5.35 | 84.23 | 114.91 | 47.01 | 130.64 | 79.03 | 121.60 |
| all Benton All6 Extended | ARM | 316 | Euarchontoglires | 87.43 | 82.50 | 89.66 | 2.94 | 82.55 | 97.79 | 78.58 | 105.18 | 81.95 | 89.80 |
| Benton All6 Extended | IRM | 316 | Euarchontoglires | 96.38 | 85.94 | 107.48 | 3.49 | 90.13 | 108.93 | 81.19 | 121.15 | 85.41 | 106.89 |
| Benton All6 Extended NoDNA | ARM | 316 | Euarchontoglires | 102.45 | 75.36 | 128.17 | 0.85 | 98.15 | 111.65 | 72.93 | 133.61 | 75.23 | 128.16 |
| all Benton All6 Extended NoDNA | IRM | 316 | Euarchontoglires | 102.46 | 75.30 | 128.29 | 0.57 | 95.18 | 104.77 | 71.82 | 134.23 | 75.22 | 128.21 |
| all Benton Rodentia Extended | ARM | 316 | Euarchontoglires | 82.72 | 77.32 | 88.49 | 2.74 | 77.29 | 92.70 | 72.75 | 101.45 | 76.79 | 87.75 |
| all Benton Rodentia Extended | IRM | 316 | Euarchontoglires | 93.44 | 83.07 | 104.50 | 3.23 | 86.51 | 105.12 | 78.14 | 118.17 | 82.49 | 103.95 |
| all Benton Rodentia Extended NoDNA | ARM | 316 | Euarchontoglires | 102.14 | 75.17 | 128.52 | 1.12 | 99.62 | 112.96 | 72.49 | 155.09 | 75.15 | 128.35 |
| all Benton Rodentia Extended NoDNA | IRM | 316 | Euarchontoglires | 101.93 | 75.07 | 128.41 | 0.63 | 97.26 | 107.04 | 73.33 | 136.59 | 75.05 | 128.28 |
| Body Size | IRM | 316 | Euarchontoglires | 78.87 | 73.66 | 84.00 | 2.79 | 75.12 | 92.29 | 71.15 | 104.01 | 73.14 | 82.64 |

| Analysis | Model | Number of Trees | node | average | L95 CI mean | U95 CI mean | std dev | mean min | mean max | L95 CI min | U95 CI max | L95 CI median | U95 CI median |
|---|---|---|---|---|---|---|---|---|---|---|---|---|---|
| 10% Missingness | IRM | 316 | Eulipotyphla | 78.74 | 69.38 | 88.94 | 2.70 | 71.70 | 97.33 | 62.87 | 112.04 | 69.46 | 88.68 |
| Accelerated | IRM | 1 | Eulipotyphla | 81.64 | 70.69 | 91.23 | | | | | | | |
| Benton | IRM | 316 | Eulipotyphla | 82.49 | 71.11 | 94.52 | 2.73 | 73.13 | 96.42 | 62.17 | 111.39 | 71.32 | 94.27 |
| Benton | ARM | 316 | Eulipotyphla | 72.26 | 68.72 | 76.00 | 1.86 | 65.43 | 82.05 | 62.29 | 92.30 | 68.93 | 75.79 |
| Benton | Averaged IRM & ARM | 1 | Eulipotyphla | 77.37 | 69.91 | 85.26 | | | | | | | |
| Cladistic | ARM | 316 | Eulipotyphla | 71.30 | 64.92 | 77.72 | 2.85 | 58.60 | 85.62 | 41.55 | 103.00 | 65.31 | 77.16 |
| Conserved | IRM | 1 | Eulipotyphla | 77.11 | 68.92 | 86.80 | | | | | | | |
| Meredith | IRM | 316 | Eulipotyphla | 81.90 | 71.10 | 93.18 | 2.39 | 72.92 | 93.26 | 61.98 | 105.66 | 71.14 | 92.92 |
| Neutral | IRM | 1 | Eulipotyphla | 81.58 | 72.55 | 90.77 | | | | | | | |
| Neutral 241 species | IRM | 1 | Eulipotyphla | 82.66 | 73.20 | 93.02 | | | | | | | |
| No DNA | ARM | 316 | Eulipotyphla | 84.67 | 62.54 | 110.01 | 0.35 | 80.97 | 85.34 | 61.51 | 110.88 | 62.54 | 110.02 |
| One Stratigraphic Bound | IRM | 316 | Eulipotyphla | 78.55 | 69.80 | 87.89 | 2.64 | 70.63 | 91.77 | 62.03 | 106.41 | 69.80 | 87.33 |
| Root | IRM | 1 | Eulipotyphla | 80.55 | 71.02 | 91.86 | | | | | | | |
| Root Calibration Only | ARM | 316 | Eulipotyphla | 88.48 | 67.03 | 106.81 | 5.61 | 64.20 | 101.62 | 41.05 | 118.86 | 68.00 | 107.30 |
| all Benton All6 Extended | ARM | 316 | Eulipotyphla | 78.21 | 72.84 | 83.46 | 2.60 | 69.44 | 91.08 | 64.36 | 102.90 | 72.92 | 83.14 |

| Analysis | Model | Number of Trees | node | average | L95 CI mean | U95 CI mean | std dev | mean min | mean max | L95 CI min | U95 CI max | L95 CI median | U95 CI median |
|---|---|---|---|---|---|---|---|---|---|---|---|---|---|
| Benton All6 Extended | IRM | 316 | Eulipotyphla | 85.77 | 73.35 | 97.79 | 2.74 | 76.54 | 100.52 | 64.45 | 114.03 | 73.13 | 97.32 |
| Benton All6 Extended NoDNA | ARM | 316 | Eulipotyphla | 87.46 | 63.60 | 119.53 | 0.57 | 84.31 | 92.82 | 62.35 | 127.26 | 63.57 | 119.42 |
| all Benton All6 Extended NoDNA | IRM | 316 | Eulipotyphla | 87.45 | 63.59 | 119.61 | 0.37 | 83.25 | 89.35 | 62.85 | 128.22 | 63.53 | 119.52 |
| all Benton Rodentia Extended | ARM | 316 | Eulipotyphla | 73.94 | 69.63 | 78.60 | 2.10 | 66.68 | 84.03 | 62.63 | 94.07 | 69.73 | 78.33 |
| all Benton Rodentia Extended | IRM | 316 | Eulipotyphla | 84.11 | 72.23 | 95.95 | 2.71 | 74.68 | 98.04 | 63.18 | 112.89 | 72.03 | 95.43 |
| all Benton Rodentia Extended NoDNA | ARM | 316 | Eulipotyphla | 89.40 | 63.05 | 120.75 | 1.12 | 87.91 | 99.61 | 61.87 | 153.90 | 63.00 | 120.36 |
| all Benton Rodentia Extended NoDNA | IRM | 316 | Eulipotyphla | 89.15 | 63.01 | 120.38 | 0.80 | 77.69 | 93.47 | 62.22 | 130.77 | 62.98 | 120.29 |
| Body Size | IRM | 316 | Eulipotyphla | 76.43 | 68.77 | 84.57 | 2.31 | 69.80 | 88.68 | 61.80 | 101.21 | 68.81 | 84.05 |
| 10% Missingness | IRM | 316 | Ferae | 74.85 | 67.32 | 82.64 | 1.31 | 69.72 | 77.76 | 62.96 | 84.45 | 67.58 | 82.97 |
| Accelerated | IRM | 1 | Ferae | 76.61 | 69.47 | 83.87 | | | | | | | |
| Benton | IRM | 316 | Ferae | 75.65 | 67.28 | 83.75 | 1.09 | 71.17 | 77.81 | 63.17 | 84.60 | 67.43 | 83.95 |
| Benton | ARM | 316 | Ferae | 71.59 | 68.50 | 74.91 | 1.23 | 66.62 | 75.46 | 63.52 | 80.38 | 68.73 | 74.72 |
| Benton | Averaged IRM & ARM | 1 | Ferae | 73.62 | 67.89 | 79.33 | | | | | | | |
| Cladistic | ARM | 316 | Ferae | 72.10 | 67.18 | 77.40 | 1.73 | 65.55 | 79.06 | 58.17 | 88.31 | 67.24 | 76.90 |
| Conserved | IRM | 1 | Ferae | 73.26 | 65.73 | 82.86 | | | | | | | |

| Analysis | Model | Number of Trees | node | average | L95 CI mean | U95 CI mean | std dev | mean min | mean max | L95 CI min | U95 CI max | L95 CI median | U95 CI median |
|---|---|---|---|---|---|---|---|---|---|---|---|---|---|
| Meredith | IRM | 316 | Ferae | 75.43 | 67.11 | 83.65 | 1.04 | 70.92 | 77.98 | 63.12 | 84.56 | 67.32 | 83.88 |
| Neutral | IRM | 1 | Ferae | 77.51 | 71.00 | 84.00 | | | | | | | |
| Neutral 241 species | IRM | 1 | Ferae | 77.27 | 69.77 | 84.52 | | | | | | | |
| No DNA | ARM | 316 | Ferae | 70.88 | 63.02 | 82.92 | 0.10 | 70.59 | 71.44 | 62.76 | 83.31 | 63.02 | 82.92 |
| One Stratigraphic Bound | IRM | 316 | Ferae | 69.28 | 64.75 | 72.59 | 0.34 | 68.09 | 69.88 | 63.97 | 72.81 | 64.71 | 72.60 |
| Root | IRM | 1 | Ferae | 76.41 | 68.59 | 84.06 | | | | | | | |
| Root Calibration Only | ARM | 316 | Ferae | 87.87 | 66.87 | 105.52 | 5.06 | 72.28 | 101.03 | 40.38 | 117.78 | 67.59 | 106.05 |
| all Benton All6 Extended | ARM | 316 | Ferae | 77.80 | 73.35 | 83.61 | 1.50 | 72.54 | 82.10 | 65.09 | 90.88 | 73.37 | 83.36 |
| Benton All6 Extended | IRM | 316 | Ferae | 78.47 | 69.59 | 84.72 | 1.30 | 72.86 | 80.47 | 64.18 | 85.29 | 69.65 | 84.77 |
| Benton All6 Extended NoDNA | ARM | 316 | Ferae | 70.34 | 63.18 | 82.06 | 0.06 | 70.16 | 70.93 | 63.01 | 82.72 | 63.18 | 82.06 |
| all Benton All6 Extended NoDNA | IRM | 316 | Ferae | 70.34 | 63.18 | 82.06 | 0.03 | 70.08 | 70.55 | 63.05 | 82.66 | 63.18 | 82.06 |
| all Benton Rodentia Extended | ARM | 316 | Ferae | 73.12 | 69.49 | 76.98 | 1.34 | 67.50 | 78.00 | 64.01 | 83.83 | 69.75 | 76.93 |
| all Benton Rodentia Extended | IRM | 316 | Ferae | 75.98 | 67.42 | 84.17 | 1.44 | 69.13 | 79.14 | 62.94 | 84.79 | 67.53 | 84.26 |
| all Benton Rodentia Extended NoDNA | ARM | 316 | Ferae | 70.22 | 62.94 | 82.46 | 0.10 | 69.95 | 71.11 | 62.77 | 83.65 | 62.94 | 82.45 |
| all Benton Rodentia Extended NoDNA | IRM | 316 | Ferae | 70.20 | 62.93 | 82.44 | 0.14 | 67.83 | 70.48 | 62.63 | 82.67 | 62.93 | 82.44 |

| **Analysis** | **Model** | **Number of Trees** | **node** | **average** | **L95 CI mean** | **U95 CI mean** | **std dev** | **mean min** | **mean max** | **L95 CI min** | **U95 CI max** | **L95 CI median** | **U95 CI median** |
|---|---|---|---|---|---|---|---|---|---|---|---|---|---|
| Body Size | IRM | 316 | Ferae | 73.47 | 65.52 | 81.30 | 1.43 | 68.28 | 78.12 | 62.70 | 84.67 | 65.59 | 81.28 |
| 10% Missingness | IRM | 316 | Ferae + Perissodactyla | 78.66 | 70.81 | 86.66 | 1.94 | 73.18 | 83.45 | 64.90 | 92.25 | 70.75 | 86.69 |
| Accelerated | IRM | 1 | Ferae + Perissodactyla | 81.48 | 74.08 | 88.29 | | | | | | | |
| Benton | IRM | 316 | Ferae + Perissodactyla | 80.66 | 71.50 | 89.36 | 1.81 | 74.92 | 85.75 | 66.57 | 96.69 | 71.40 | 89.05 |
| Benton | ARM | 316 | Ferae + Perissodactyla | 74.82 | 71.68 | 78.27 | 1.29 | 70.20 | 78.96 | 68.22 | 84.98 | 71.59 | 78.14 |
| Benton | Averaged IRM & ARM | 1 | Ferae + Perissodactyla | 77.74 | 71.59 | 83.81 | | | | | | | |
| Cladistic | ARM | 316 | Ferae + Perissodactyla | 75.51 | 70.36 | 81.12 | 2.30 | 70.68 | 84.32 | 67.84 | 94.22 | 69.99 | 80.47 |
| Conserved | IRM | 1 | Ferae + Perissodactyla | 77.49 | 70.02 | 86.61 | | | | | | | |
| Meredith | IRM | 316 | Ferae + Perissodactyla | 80.33 | 71.38 | 88.84 | 1.57 | 74.82 | 85.10 | 65.86 | 94.17 | 71.38 | 88.58 |
| Neutral | IRM | 1 | Ferae + Perissodactyla | 82.73 | 75.61 | 90.99 | | | | | | | |
| Neutral 241 species | IRM | 1 | Ferae + Perissodactyla | 82.84 | 73.60 | 92.75 | | | | | | | |
| No DNA | ARM | 316 | Ferae + Perissodactyla | 75.72 | 63.25 | 91.79 | 0.16 | 74.32 | 76.26 | 62.76 | 92.45 | 63.26 | 91.82 |
| One Stratigraphic Bound | IRM | 316 | Ferae + Perissodactyla | 74.64 | 68.29 | 81.47 | 1.49 | 71.76 | 79.82 | 65.72 | 89.77 | 68.22 | 81.08 |
| Root | IRM | 1 | Ferae + Perissodactyla | 81.63 | 72.12 | 90.22 | | | | | | | |
| Root Calibration Only | ARM | 316 | Ferae + Perissodactyla | 92.84 | 70.90 | 111.08 | 4.94 | 76.14 | 105.73 | 42.62 | 121.64 | 72.07 | 111.46 |

| Analysis | Model | Number of Trees | node | average | L95 CI mean | U95 CI mean | std dev | mean min | mean max | L95 CI min | U95 CI max | L95 CI median | U95 CI median |
|---|---|---|---|---|---|---|---|---|---|---|---|---|---|
| all Benton All6 Extended | ARM | 316 | Ferae + Perissodactyla | 81.76 | 77.10 | 87.12 | 1.94 | 76.53 | 88.34 | 72.70 | 94.29 | 76.97 | 86.88 |
| Benton All6 Extended | IRM | 316 | Ferae + Perissodactyla | 84.64 | 75.01 | 93.16 | 1.70 | 78.80 | 89.58 | 69.82 | 101.55 | 74.71 | 92.49 |
| Benton All6 Extended NoDNA | ARM | 316 | Ferae + Perissodactyla | 78.21 | 63.91 | 103.97 | 0.31 | 76.56 | 80.27 | 63.54 | 114.02 | 63.90 | 103.81 |
| all Benton All6 Extended NoDNA | IRM | 316 | Ferae + Perissodactyla | 78.22 | 63.89 | 104.01 | 0.20 | 76.26 | 79.27 | 63.61 | 110.01 | 63.89 | 103.92 |
| all Benton Rodentia Extended | ARM | 316 | Ferae + Perissodactyla | 76.75 | 72.95 | 80.82 | 1.47 | 71.93 | 82.84 | 69.44 | 88.71 | 72.84 | 80.56 |
| all Benton Rodentia Extended | IRM | 316 | Ferae + Perissodactyla | 81.36 | 71.51 | 90.44 | 2.00 | 74.56 | 87.22 | 66.07 | 98.48 | 71.43 | 89.87 |
| all Benton Rodentia Extended NoDNA | ARM | 316 | Ferae + Perissodactyla | 77.75 | 63.07 | 105.40 | 0.76 | 76.78 | 84.53 | 62.72 | 149.61 | 63.04 | 104.70 |
| all Benton Rodentia Extended NoDNA | IRM | 316 | Ferae + Perissodactyla | 77.60 | 63.04 | 104.78 | 0.38 | 72.36 | 79.49 | 62.55 | 116.66 | 63.02 | 104.57 |
| 10% Missingness | IRM | 316 | Fereuungulata | 80.34 | 72.27 | 88.50 | 2.46 | 74.20 | 88.79 | 69.02 | 99.80 | 72.00 | 88.25 |
| Accelerated | IRM | 1 | Fereuungulata | 83.46 | 76.22 | 90.66 | | | | | | | |
| Benton | IRM | 316 | Fereuungulata | 83.52 | 74.04 | 92.60 | 2.65 | 76.13 | 93.33 | 69.81 | 105.22 | 73.54 | 92.02 |
| Benton | ARM | 316 | Fereuungulata | 76.25 | 72.92 | 79.89 | 1.70 | 72.64 | 83.31 | 70.47 | 89.80 | 72.71 | 79.51 |
| Benton | Averaged IRM & ARM | 1 | Fereuungulata | 79.89 | 73.48 | 86.25 | | | | | | | |
| Cladistic | ARM | 316 | Fereuungulata | 77.00 | 71.64 | 82.85 | 2.87 | 71.29 | 90.01 | 68.50 | 99.50 | 71.20 | 82.04 |
| Conserved | IRM | 1 | Fereuungulata | 79.04 | 72.01 | 88.71 | | | | | | | |

| Analysis | Model | Number of Trees | node | average | L95 CI mean | U95 CI mean | std dev | mean min | mean max | L95 CI min | U95 CI max | L95 CI median | U95 CI median |
|---|---|---|---|---|---|---|---|---|---|---|---|---|---|
| Meredith | IRM | 316 | Fereuungulata | 83.13 | 73.92 | 91.91 | 2.29 | 76.04 | 90.92 | 69.72 | 100.48 | 73.58 | 91.59 |
| Neutral | IRM | 1 | Fereuungulata | 84.65 | 77.60 | 93.14 | | | | | | | |
| Neutral 241 species | IRM | 1 | Fereuungulata | 84.85 | 75.74 | 95.46 | | | | | | | |
| No DNA | ARM | 316 | Fereuungulata | 81.37 | 64.32 | 102.17 | 0.28 | 78.44 | 81.94 | 63.85 | 102.91 | 64.32 | 102.19 |
| One Stratigraphic Bound | IRM | 316 | Fereuungulata | 77.66 | 70.75 | 85.17 | 2.54 | 73.22 | 87.92 | 68.36 | 99.08 | 70.46 | 84.60 |
| Root | IRM | 1 | Fereuungulata | 83.43 | 73.13 | 92.02 | | | | | | | |
| Root Calibration Only | ARM | 316 | Fereuungulata | 95.03 | 72.65 | 113.59 | 5.08 | 77.97 | 108.07 | 43.48 | 123.39 | 73.57 | 113.75 |
| all Benton All6 Extended | ARM | 316 | Fereuungulata | 83.51 | 78.63 | 88.38 | 2.54 | 77.70 | 93.46 | 73.75 | 101.07 | 78.27 | 87.79 |
| Benton All6 Extended | IRM | 316 | Fereuungulata | 87.84 | 78.64 | 96.77 | 2.55 | 82.40 | 97.67 | 72.02 | 108.96 | 78.54 | 95.91 |
| Benton All6 Extended NoDNA | ARM | 316 | Fereuungulata | 87.03 | 67.66 | 116.59 | 0.60 | 83.86 | 91.52 | 66.46 | 124.06 | 67.64 | 116.42 |
| all Benton All6 Extended NoDNA | IRM | 316 | Fereuungulata | 87.03 | 67.63 | 116.69 | 0.38 | 82.96 | 89.12 | 66.07 | 128.18 | 67.60 | 116.54 |
| all Benton Rodentia Extended | ARM | 316 | Fereuungulata | 78.37 | 74.32 | 82.71 | 1.98 | 74.85 | 86.96 | 71.71 | 94.62 | 74.04 | 82.16 |
| all Benton Rodentia Extended | IRM | 316 | Fereuungulata | 84.34 | 74.19 | 93.83 | 2.78 | 77.43 | 94.94 | 70.14 | 107.62 | 73.78 | 93.08 |
| all Benton Rodentia Extended NoDNA | ARM | 316 | Fereuungulata | 85.39 | 64.48 | 115.76 | 1.07 | 83.75 | 95.02 | 63.94 | 152.00 | 64.44 | 115.24 |
| all Benton Rodentia Extended NoDNA | IRM | 316 | Fereuungulata | 85.16 | 64.46 | 115.36 | 0.53 | 79.36 | 88.83 | 63.95 | 130.75 | 64.44 | 115.17 |

| Analysis | Model | Number of Trees | node | average | L95 CI mean | U95 CI mean | std dev | mean min | mean max | L95 CI min | U95 CI max | L95 CI median | U95 CI median |
|---|---|---|---|---|---|---|---|---|---|---|---|---|---|
| Body Size | IRM | 316 | Fereuungulata | 77.74 | 69.86 | 85.90 | 2.24 | 72.97 | 86.09 | 65.40 | 96.29 | 69.75 | 85.64 |
| 10% Missingness | IRM | 316 | Glires | 75.76 | 70.02 | 81.97 | 1.81 | 71.08 | 83.87 | 63.88 | 97.03 | 70.01 | 81.03 |
| Accelerated | IRM | 1 | Glires | 76.26 | 71.07 | 82.15 | | | | | | | |
| Benton | IRM | 316 | Glires | 78.17 | 70.77 | 86.74 | 2.48 | 74.45 | 87.54 | 68.68 | 100.83 | 70.58 | 86.36 |
| Benton | ARM | 316 | Glires | 74.56 | 72.21 | 76.71 | 0.90 | 70.97 | 79.76 | 68.74 | 84.98 | 72.21 | 76.40 |
| Benton | Averaged IRM & ARM | 1 | Glires | 76.36 | 71.49 | 81.72 | | | | | | | |
| Cladistic | ARM | 316 | Glires | 74.59 | 71.96 | 76.96 | 1.20 | 71.29 | 82.63 | 68.82 | 89.31 | 71.88 | 76.57 |
| Conserved | IRM | 1 | Glires | 75.68 | 69.87 | 86.80 | | | | | | | |
| Meredith | IRM | 316 | Glires | 78.01 | 70.79 | 86.32 | 2.27 | 74.07 | 86.19 | 68.78 | 97.63 | 70.59 | 86.00 |
| Neutral | IRM | 1 | Glires | 76.71 | 67.07 | 89.16 | | | | | | | |
| Neutral 241 species | IRM | 1 | Glires | 77.79 | 71.85 | 85.85 | | | | | | | |
| No DNA | ARM | 316 | Glires | 74.79 | 59.15 | 98.37 | 0.14 | 73.70 | 75.19 | 58.70 | 99.02 | 59.15 | 98.43 |
| One Stratigraphic Bound | IRM | 316 | Glires | 77.21 | 69.74 | 85.46 | 2.30 | 73.49 | 86.08 | 67.17 | 99.09 | 69.52 | 84.83 |
| Root | IRM | 1 | Glires | 75.83 | 70.53 | 81.50 | | | | | | | |
| Root Calibration Only | ARM | 316 | Glires | 97.09 | 74.12 | 116.08 | 5.12 | 81.68 | 109.80 | 44.96 | 126.23 | 74.90 | 116.51 |

| Analysis | Model | Number of Trees | node | average | L95 CI mean | U95 CI mean | std dev | mean min | mean max | L95 CI min | U95 CI max | L95 CI median | U95 CI median |
|---|---|---|---|---|---|---|---|---|---|---|---|---|---|
| all Benton All6 Extended | ARM | 316 | Glires | 83.05 | 78.71 | 78.59 | 1.72 | 78.37 | 88.87 | 73.81 | 95.17 | 78.58 | 85.92 |
| Benton All6 Extended | IRM | 316 | Glires | 89.93 | 79.98 | 100.72 | 1.99 | 86.45 | 96.55 | 76.72 | 109.77 | 79.83 | 100.41 |
| Benton All6 Extended NoDNA | ARM | 316 | Glires | 87.83 | 66.46 | 119.14 | 0.58 | 84.58 | 93.01 | 64.65 | 125.92 | 66.45 | 119.03 |
| all Benton All6 Extended NoDNA | IRM | 316 | Glires | 87.85 | 66.50 | 119.35 | 0.37 | 83.63 | 89.75 | 65.22 | 128.21 | 66.49 | 119.24 |
| all Benton Rodentia Extended | ARM | 316 | Glires | 78.87 | 74.04 | 84.00 | 1.74 | 75.05 | 84.55 | 71.01 | 91.33 | 73.83 | 83.82 |
| all Benton Rodentia Extended | IRM | 316 | Glires | 87.27 | 77.82 | 97.73 | 1.73 | 83.17 | 93.28 | 75.19 | 106.29 | 77.70 | 97.40 |
| all Benton Rodentia Extended NoDNA | ARM | 316 | Glires | 88.32 | 63.55 | 119.37 | 1.02 | 86.95 | 99.62 | 60.36 | 149.31 | 63.49 | 118.89 |
| all Benton Rodentia Extended NoDNA | IRM | 316 | Glires | 88.15 | 63.54 | 119.02 | 0.50 | 85.96 | 92.97 | 62.33 | 130.85 | 63.51 | 118.89 |
| Body Size | IRM | 316 | Glires | 77.74 | 69.86 | 85.90 | 2.24 | 72.97 | 86.09 | 65.40 | 96.29 | 69.75 | 85.64 |
| 10% Missingness | IRM | 316 | Lagomorpha | 57.19 | 53.58 | 61.37 | 0.20 | 56.69 | 58.11 | 53.43 | 61.75 | 53.57 | 61.36 |
| Accelerated | IRM | 1 | Lagomorpha | 57.22 | 53.54 | 61.28 | | | | | | | |
| Benton | IRM | 316 | Lagomorpha | 57.31 | 53.61 | 61.43 | 0.17 | 56.81 | 58.07 | 53.48 | 61.71 | 53.61 | 61.43 |
| Benton | ARM | 316 | Lagomorpha | 54.81 | 53.12 | 57.21 | 0.37 | 54.15 | 57.93 | 52.93 | 61.75 | 53.11 | 57.04 |
| Benton | Averaged IRM & ARM | 1 | Lagomorpha | 56.06 | 53.36 | 59.32 | | | | | | | |
| Cladistic | ARM | 316 | Lagomorpha | 54.59 | 53.05 | 56.83 | 0.35 | 54.10 | 57.73 | 52.89 | 61.62 | 53.04 | 56.63 |

| Analysis | Model | Number of Trees | node | average | L95 CI mean | U95 CI mean | std dev | mean min | mean max | L95 CI min | U95 CI max | L95 CI median | U95 CI median |
|---|---|---|---|---|---|---|---|---|---|---|---|---|---|
| Conserved | IRM | 1 | Lagomorpha | 57.06 | 53.54 | 61.29 | | | | | | | |
| Meredith | IRM | 316 | Lagomorpha | 57.30 | 53.61 | 61.43 | 0.16 | 56.83 | 58.04 | 53.48 | 61.74 | 53.61 | 61.43 |
| Neutral | IRM | 1 | Lagomorpha | 57.17 | 53.55 | 61.37 | | | | | | | |
| Neutral 241 species | IRM | 1 | Lagomorpha | 57.11 | 53.57 | 61.34 | | | | | | | |
| No DNA | ARM | 316 | Lagomorpha | 57.73 | 53.75 | 61.62 | 0.01 | 57.71 | 57.76 | 53.69 | 61.67 | 53.75 | 61.62 |
| One Stratigraphic Bound | IRM | 316 | Lagomorpha | 57.12 | 53.56 | 61.26 | 0.15 | 56.68 | 57.83 | 53.44 | 61.68 | 53.56 | 61.25 |
| Root | IRM | 1 | Lagomorpha | 57.33 | 53.63 | 61.48 | | | | | | | |
| Root Calibration Only | ARM | 316 | Lagomorpha | 52.23 | 33.74 | 70.78 | 3.93 | 41.79 | 70.88 | 0.00 | 98.42 | 34.32 | 70.55 |
| all Benton All6 Extended | ARM | 316 | Lagomorpha | 55.47 | 52.06 | 71.05 | 1.05 | 53.95 | 66.00 | 51.05 | 106.16 | 52.04 | 60.11 |
| Benton All6 Extended | IRM | 316 | Lagomorpha | 62.24 | 53.13 | 72.66 | 0.58 | 60.68 | 65.72 | 52.77 | 75.25 | 53.12 | 72.68 |
| Benton All6 Extended NoDNA | ARM | 316 | Lagomorpha | 64.87 | 53.92 | 75.05 | 0.04 | 64.78 | 65.36 | 53.76 | 75.29 | 53.89 | 75.08 |
| all Benton All6 Extended NoDNA | IRM | 316 | Lagomorpha | 64.87 | 53.89 | 75.08 | 0.02 | 64.83 | 64.92 | 53.74 | 75.19 | 53.89 | 75.08 |
| all Benton Rodentia Extended | ARM | 316 | Lagomorpha | 55.43 | 53.18 | 58.65 | 0.43 | 54.65 | 57.85 | 53.03 | 61.76 | 53.16 | 58.51 |
| all Benton Rodentia Extended | IRM | 316 | Lagomorpha | 57.56 | 53.70 | 61.55 | 0.08 | 57.25 | 57.94 | 53.59 | 61.73 | 53.69 | 61.54 |
| all Benton Rodentia Extended NoDNA | ARM | 316 | Lagomorpha | 57.71 | 53.73 | 61.62 | 0.01 | 57.61 | 57.73 | 53.67 | 61.65 | 53.73 | 61.62 |

| **Analysis** | **Model** | **Number of Trees** | **node** | **average** | **L95 CI mean** | **U95 CI mean** | **std dev** | **mean min** | **mean max** | **L95 CI min** | **U95 CI max** | **L95 CI median** | **U95 CI median** |
|---|---|---|---|---|---|---|---|---|---|---|---|---|---|
| all Benton Rodentia Extended NoDNA | IRM | 316 | Lagomorpha | 57.71 | 53.73 | 61.62 | 0.01 | 57.61 | 57.73 | 53.68 | 61.67 | 53.73 | 61.62 |
| Body Size | IRM | 316 | Lagomorpha | 56.66 | 53.42 | 60.86 | 0.22 | 56.16 | 58.04 | 53.25 | 61.69 | 53.41 | 60.85 |
| 10% Missingness | IRM | 316 | Laurasiatheria | 86.74 | 77.49 | 96.41 | 3.55 | 79.38 | 102.00 | 72.32 | 116.33 | 76.97 | 95.85 |
| Accelerated | IRM | 1 | Laurasiatheria | 89.72 | 81.34 | 97.65 | | | | | | | |
| Benton | IRM | 316 | Laurasiatheria | 92.53 | 81.60 | 103.87 | 3.97 | 85.32 | 107.38 | 75.36 | 120.56 | 81.00 | 103.01 |
| Benton | ARM | 316 | Laurasiatheria | 80.34 | 76.33 | 84.72 | 2.55 | 75.80 | 92.14 | 72.76 | 101.33 | 76.00 | 83.82 |
| Benton | Averaged IRM & ARM | 1 | Laurasiatheria | 86.43 | 78.97 | 94.29 | | | | | | | |
| Cladistic | ARM | 316 | Laurasiatheria | 81.15 | 75.18 | 87.63 | 3.76 | 74.66 | 99.93 | 70.58 | 111.08 | 74.66 | 86.45 |
| Conserved | IRM | 1 | Laurasiatheria | 84.86 | 76.78 | 94.96 | | | | | | | |
| Meredith | IRM | 316 | Laurasiatheria | 91.69 | 81.59 | 102.00 | 3.27 | 84.30 | 102.60 | 75.27 | 112.81 | 81.10 | 101.57 |
| Neutral | IRM | 1 | Laurasiatheria | 90.78 | 83.05 | 100.37 | | | | | | | |
| Neutral 241 species | IRM | 1 | Laurasiatheria | 91.28 | 82.36 | 100.75 | | | | | | | |
| No DNA | ARM | 316 | Laurasiatheria | 95.85 | 70.72 | 122.94 | 0.61 | 89.54 | 96.81 | 66.17 | 124.66 | 70.84 | 122.97 |
| One Stratigraphic Bound | IRM | 316 | Laurasiatheria | 87.45 | 78.55 | 97.21 | 4.03 | 80.47 | 103.37 | 73.31 | 117.44 | 77.94 | 96.01 |
| Root | IRM | 1 | Laurasiatheria | 89.46 | 79.14 | 93.56 | | | | | | | |

| Analysis | Model | Number of Trees | node | average | L95 CI mean | U95 CI mean | std dev | mean min | mean max | L95 CI min | U95 CI max | L95 CI median | U95 CI median |
|---|---|---|---|---|---|---|---|---|---|---|---|---|---|
| Root Calibration Only | ARM | 316 | Laurasiatheria | 101.13 | 77.43 | 120.55 | 5.51 | 83.05 | 113.73 | 46.37 | 129.23 | 78.38 | 120.58 |
| all Benton All6 Extended | ARM | 316 | Laurasiatheria | 88.59 | 82.92 | 94.77 | 3.61 | 81.33 | 103.46 | 76.90 | 112.21 | 82.36 | 93.62 |
| Benton All6 Extended | IRM | 316 | Laurasiatheria | 97.51 | 86.94 | 108.53 | 3.89 | 90.84 | 111.33 | 79.98 | 123.39 | 86.42 | 107.70 |
| Benton All6 Extended NoDNA | ARM | 316 | Laurasiatheria | 102.70 | 74.52 | 128.36 | 0.83 | 98.38 | 111.28 | 72.19 | 133.61 | 74.40 | 128.32 |
| all Benton All6 Extended NoDNA | IRM | 316 | Laurasiatheria | 102.71 | 74.42 | 128.45 | 0.55 | 95.66 | 105.01 | 72.33 | 134.24 | 74.32 | 128.34 |
| all Benton Rodentia Extended | ARM | 316 | Laurasiatheria | 83.12 | 78.09 | 88.47 | 3.02 | 77.99 | 95.37 | 73.81 | 105.08 | 77.59 | 87.46 |
| all Benton Rodentia Extended | IRM | 316 | Laurasiatheria | 94.59 | 83.21 | 106.07 | 3.98 | 86.85 | 108.62 | 76.59 | 121.61 | 82.68 | 105.15 |
| all Benton Rodentia Extended NoDNA | ARM | 316 | Laurasiatheria | 102.20 | 73.13 | 128.95 | 1.11 | 99.66 | 112.41 | 68.42 | 157.14 | 73.14 | 128.78 |
| all Benton Rodentia Extended NoDNA | IRM | 316 | Laurasiatheria | 101.97 | 73.00 | 128.79 | 0.65 | 97.25 | 106.75 | 68.96 | 136.68 | 72.98 | 128.70 |
| Body Size | IRM | 316 | Laurasiatheria | 84.19 | 76.12 | 92.70 | 3.49 | 78.74 | 101.02 | 71.27 | 112.85 | 75.52 | 91.99 |
| 10% Missingness | IRM | 316 | Paenungulata | 62.90 | 58.70 | 68.52 | 0.65 | 61.16 | 65.63 | 58.36 | 72.34 | 58.68 | 68.45 |
| Accelerated | IRM | 1 | Paenungulata | 63.30 | 58.69 | 69.54 | | | | | | | |
| Benton | IRM | 316 | Paenungulata | 62.92 | 58.73 | 68.42 | 0.58 | 61.56 | 65.28 | 58.47 | 71.83 | 58.71 | 68.32 |
| Benton | ARM | 316 | Paenungulata | 61.14 | 58.35 | 65.07 | 0.37 | 60.29 | 62.72 | 57.89 | 67.92 | 58.33 | 65.02 |
| Benton | Averaged IRM & ARM | 1 | Paenungulata | 62.03 | 58.54 | 66.75 | | | | | | | |

| Analysis | Model | Number of Trees | node | average | L95 CI mean | U95 CI mean | std dev | mean min | mean max | L95 CI min | U95 CI max | L95 CI median | U95 CI median |
|---|---|---|---|---|---|---|---|---|---|---|---|---|---|
| Cladistic | ARM | 316 | Paenungulata | 61.17 | 58.48 | 64.85 | 0.35 | 60.52 | 63.18 | 58.07 | 68.70 | 58.48 | 64.71 |
| Conserved | IRM | 1 | Paenungulata | 62.10 | 58.39 | 67.72 | | | | | | | |
| Meredith | IRM | 316 | Paenungulata | 62.86 | 58.72 | 68.32 | 0.57 | 61.40 | 65.13 | 58.42 | 71.83 | 58.71 | 68.28 |
| Neutral | IRM | 1 | Paenungulata | 63.54 | 58.91 | 69.57 | | | | | | | |
| Neutral 241 species | IRM | 1 | Paenungulata | 63.05 | 58.49 | 69.67 | | | | | | | |
| No DNA | ARM | 316 | Paenungulata | 65.95 | 59.43 | 72.42 | 0.04 | 65.82 | 66.46 | 59.20 | 72.50 | 59.43 | 72.41 |
| One Stratigraphic Bound | IRM | 316 | Paenungulata | 61.54 | 58.95 | 64.72 | 0.36 | 60.53 | 62.70 | 58.80 | 66.03 | 58.94 | 64.76 |
| Root | IRM | 1 | Paenungulata | 62.50 | 58.65 | 67.70 | | | | | | | |
| Root Calibration Only | ARM | 316 | Paenungulata | 61.70 | 43.74 | 78.84 | 4.26 | 48.46 | 71.33 | 26.80 | 87.89 | 43.91 | 79.11 |
| all Benton All6 Extended | ARM | 316 | Paenungulata | 61.83 | 58.58 | 66.82 | 0.58 | 60.82 | 64.59 | 58.19 | 69.93 | 58.53 | 66.84 |
| Benton All6 Extended | IRM | 316 | Paenungulata | 64.13 | 58.93 | 69.68 | 0.72 | 62.55 | 66.79 | 58.67 | 72.37 | 58.89 | 69.68 |
| Benton All6 Extended NoDNA | ARM | 316 | Paenungulata | 66.22 | 59.76 | 72.44 | 0.02 | 66.09 | 66.31 | 59.60 | 72.53 | 59.76 | 72.44 |
| all Benton All6 Extended NoDNA | IRM | 316 | Paenungulata | 66.22 | 59.76 | 72.44 | 0.01 | 66.08 | 66.25 | 59.67 | 72.52 | 59.76 | 72.44 |
| all Benton Rodentia Extended | ARM | 316 | Paenungulata | 61.44 | 58.34 | 66.05 | 0.44 | 60.35 | 63.26 | 57.92 | 69.29 | 58.31 | 66.00 |
| all Benton Rodentia Extended | IRM | 316 | Paenungulata | 63.65 | 58.81 | 69.51 | 0.75 | 61.96 | 66.56 | 58.53 | 72.39 | 58.77 | 69.50 |

| Analysis | Model | Number of Trees | node | average | L95 CI mean | U95 CI mean | std dev | mean min | mean max | L95 CI min | U95 CI max | L95 CI median | U95 CI median |
|---|---|---|---|---|---|---|---|---|---|---|---|---|---|
| all Benton Rodentia Extended NoDNA | ARM | 316 | Paenungulata | 66.15 | 59.59 | 72.45 | 0.03 | 65.86 | 66.25 | 59.45 | 72.52 | 59.59 | 72.46 |
| all Benton Rodentia Extended NoDNA | IRM | 316 | Paenungulata | 66.15 | 59.59 | 72.45 | 0.01 | 66.09 | 66.20 | 59.52 | 72.53 | 59.59 | 72.45 |
| 10% Missingness | IRM | 316 | Perissodactyla | 59.66 | 56.75 | 61.83 | 0.18 | 59.16 | 60.18 | 56.22 | 61.93 | 56.74 | 61.83 |
| Accelerated | IRM | 1 | Perissodactyla | 59.77 | 56.84 | 61.85 | | | | | | | |
| Benton | IRM | 316 | Perissodactyla | 59.69 | 56.78 | 61.84 | 0.14 | 59.23 | 60.19 | 56.31 | 61.94 | 56.77 | 61.83 |
| Benton | ARM | 316 | Perissodactyla | 60.27 | 58.24 | 61.84 | 0.35 | 58.88 | 61.14 | 56.50 | 62.06 | 58.26 | 61.84 |
| Benton | Averaged IRM & ARM | 1 | Perissodactyla | 59.98 | 57.51 | 61.84 | | | | | | | |
| Cladistic | ARM | 316 | Perissodactyla | 60.08 | 58.04 | 61.80 | 0.38 | 58.99 | 61.14 | 56.47 | 62.04 | 58.05 | 61.80 |
| Conserved | IRM | 1 | Perissodactyla | 59.57 | 56.65 | 61.80 | | | | | | | |
| Meredith | IRM | 316 | Perissodactyla | 59.69 | 56.78 | 61.83 | 0.14 | 59.25 | 60.12 | 56.34 | 61.92 | 56.77 | 61.84 |
| Neutral | IRM | 1 | Perissodactyla | 59.77 | 57.00 | 61.82 | | | | | | | |
| Neutral 241 species | IRM | 1 | Perissodactyla | 59.55 | 56.63 | 61.82 | | | | | | | |
| No DNA | ARM | 316 | Perissodactyla | 59.40 | 56.36 | 61.77 | 0.00 | 59.38 | 59.43 | 56.32 | 61.81 | 56.36 | 61.77 |
| One Stratigraphic Bound | IRM | 316 | Perissodactyla | 58.03 | 56.30 | 59.32 | 0.07 | 57.83 | 58.26 | 56.12 | 59.38 | 56.29 | 59.32 |
| Root | IRM | 1 | Perissodactyla | 59.68 | 56.72 | 61.82 | | | | | | | |

| Analysis | Model | Number of Trees | node | average | L95 CI mean | U95 CI mean | std dev | mean min | mean max | L95 CI min | U95 CI max | L95 CI median | U95 CI median |
|---|---|---|---|---|---|---|---|---|---|---|---|---|---|
| Root Calibration Only | ARM | 316 | Perissodactyla | 68.96 | 50.15 | 86.51 | 4.49 | 56.85 | 80.14 | 31.59 | 98.93 | 50.56 | 86.96 |
| all Benton All6 Extended | ARM | 316 | Perissodactyla | 64.04 | 59.63 | 68.06 | 1.19 | 60.78 | 68.43 | 56.88 | 72.56 | 59.50 | 67.92 |
| Benton All6 Extended | IRM | 316 | Perissodactyla | 64.11 | 56.82 | 71.56 | 0.94 | 61.30 | 66.78 | 55.68 | 74.77 | 56.69 | 71.62 |
| Benton All6 Extended NoDNA | ARM | 316 | Perissodactyla | 65.73 | 57.25 | 74.92 | 0.02 | 65.66 | 65.83 | 56.97 | 75.09 | 57.26 | 74.93 |
| all Benton All6 Extended NoDNA | IRM | 316 | Perissodactyla | 65.73 | 57.26 | 74.93 | 0.01 | 65.67 | 65.79 | 56.95 | 75.09 | 57.26 | 74.94 |
| all Benton Rodentia Extended | ARM | 316 | Perissodactyla | 60.37 | 58.37 | 61.88 | 0.35 | 58.98 | 61.16 | 56.54 | 62.07 | 58.40 | 61.88 |
| all Benton Rodentia Extended | IRM | 316 | Perissodactyla | 59.57 | 56.71 | 61.81 | 0.16 | 59.05 | 60.15 | 56.25 | 61.92 | 56.69 | 61.81 |
| all Benton Rodentia Extended NoDNA | ARM | 316 | Perissodactyla | 59.38 | 56.33 | 61.77 | 0.01 | 59.28 | 59.40 | 56.26 | 61.80 | 56.33 | 61.77 |
| all Benton Rodentia Extended NoDNA | IRM | 316 | Perissodactyla | 59.38 | 56.33 | 61.77 | 0.01 | 59.32 | 59.39 | 56.27 | 61.86 | 56.33 | 61.77 |
| 10% Missingness | IRM | 316 | Primates | 62.65 | 57.68 | 66.41 | 0.34 | 60.75 | 63.56 | 55.91 | 66.64 | 57.68 | 66.42 |
| Accelerated | IRM | 1 | Primates | 62.57 | 57.57 | 66.34 | | | | | | | |
| Benton | IRM | 316 | Primates | 62.46 | 57.47 | 66.36 | 0.31 | 60.73 | 63.01 | 55.88 | 66.51 | 57.50 | 66.37 |
| Benton | ARM | 316 | Primates | 65.42 | 63.38 | 67.10 | 0.69 | 60.40 | 66.05 | 55.89 | 67.72 | 63.71 | 67.17 |
| Benton | Averaged IRM & ARM | 1 | Primates | 63.94 | 60.42 | 66.73 | | | | | | | |
| Cladistic | ARM | 316 | Primates | 65.41 | 63.45 | 67.04 | 0.60 | 61.37 | 66.03 | 57.09 | 67.65 | 63.70 | 67.07 |

| Analysis | Model | Number of Trees | node | average | L95 CI mean | U95 CI mean | std dev | mean min | mean max | L95 CI min | U95 CI max | L95 CI median | U95 CI median |
|---|---|---|---|---|---|---|---|---|---|---|---|---|---|
| Conserved | IRM | 1 | Primates | 62.79 | 58.09 | 66.44 | | | | | | | |
| Meredith | IRM | 316 | Primates | 62.46 | 57.47 | 66.36 | 0.32 | 60.81 | 63.01 | 55.92 | 66.52 | 57.50 | 66.38 |
| Neutral | IRM | 1 | Primates | 62.64 | 57.83 | 66.40 | | | | | | | |
| Neutral 241 species | IRM | 1 | Primates | 63.48 | 58.87 | 66.59 | | | | | | | |
| No DNA | ARM | 316 | Primates | 60.94 | 56.01 | 65.93 | 0.01 | 60.91 | 60.98 | 55.95 | 66.03 | 56.00 | 65.93 |
| One Stratigraphic Bound | IRM | 316 | Primates | 59.51 | 56.43 | 61.80 | 0.10 | 59.04 | 60.32 | 56.13 | 61.88 | 56.42 | 61.80 |
| Root | IRM | 1 | Primates | 62.52 | 57.49 | 66.44 | | | | | | | |
| Root Calibration Only | ARM | 316 | Primates | 88.47 | 67.12 | 106.66 | 5.72 | 55.17 | 101.89 | 34.39 | 117.21 | 67.92 | 107.57 |
| all Benton All6 Extended | ARM | 316 | Primates | 73.94 | 70.81 | 76.51 | 1.06 | 65.55 | 74.97 | 58.97 | 77.47 | 71.25 | 76.59 |
| Benton All6 Extended | IRM | 316 | Primates | 70.41 | 62.64 | 76.07 | 0.53 | 67.55 | 71.45 | 58.21 | 76.36 | 62.80 | 76.09 |
| Benton All6 Extended NoDNA | ARM | 316 | Primates | 65.48 | 56.07 | 74.98 | 0.09 | 64.70 | 66.42 | 55.82 | 75.15 | 56.08 | 74.99 |
| all Benton All6 Extended NoDNA | IRM | 316 | Primates | 65.48 | 56.08 | 74.99 | 0.05 | 64.87 | 65.74 | 55.84 | 75.13 | 56.09 | 75.00 |
| all Benton Rodentia Extended | ARM | 316 | Primates | 65.63 | 63.55 | 67.33 | 0.37 | 62.14 | 66.05 | 57.59 | 67.76 | 63.71 | 67.37 |
| all Benton Rodentia Extended | IRM | 316 | Primates | 63.14 | 57.86 | 66.61 | 0.22 | 62.02 | 63.59 | 56.69 | 66.76 | 57.87 | 66.62 |
| all Benton Rodentia Extended NoDNA | ARM | 316 | Primates | 60.97 | 56.02 | 65.97 | 0.02 | 60.78 | 61.01 | 55.94 | 66.05 | 56.02 | 65.97 |

| Analysis | Model | Number of Trees | node | average | L95 CI mean | U95 CI mean | std dev | mean min | mean max | L95 CI min | U95 CI max | L95 CI median | U95 CI median |
|---|---|---|---|---|---|---|---|---|---|---|---|---|---|
| all Benton Rodentia Extended NoDNA | IRM | 316 | Primates | 60.97 | 56.02 | 65.98 | 0.03 | 60.92 | 61.38 | 55.96 | 66.09 | 56.02 | 65.98 |
| Body Size | IRM | 316 | Primates | 62.66 | 57.70 | 66.40 | 0.51 | 60.13 | 63.72 | 55.79 | 66.65 | 57.73 | 66.44 |
| 10% Missingness | IRM | 316 | Primatomorpha | 73.79 | 69.83 | 76.47 | 0.48 | 70.67 | 74.64 | 65.91 | 76.66 | 70.07 | 76.48 |
| Accelerated | IRM | 1 | Primatomorpha | 74.00 | 70.31 | 76.52 | | | | | | | |
| Benton | IRM | 316 | Primatomorpha | 73.49 | 69.11 | 76.45 | 0.42 | 71.57 | 74.47 | 66.30 | 76.63 | 69.19 | 76.46 |
| Benton | ARM | 316 | Primatomorpha | 74.54 | 72.39 | 76.27 | 0.72 | 68.62 | 75.53 | 65.43 | 76.88 | 72.58 | 76.32 |
| Benton | Averaged IRM & ARM | 1 | Primatomorpha | 74.01 | 70.75 | 76.36 | | | | | | | |
| Cladistic | ARM | 316 | Primatomorpha | 74.51 | 72.25 | 76.30 | 0.58 | 69.47 | 75.53 | 66.27 | 76.86 | 72.31 | 76.32 |
| Conserved | IRM | 1 | Primatomorpha | 73.96 | 70.34 | 76.46 | | | | | | | |
| Meredith | IRM | 316 | Primatomorpha | 73.48 | 69.10 | 76.45 | 0.42 | 71.51 | 74.44 | 66.29 | 76.62 | 69.18 | 76.45 |
| Neutral | IRM | 1 | Primatomorpha | 73.54 | 69.37 | 76.56 | | | | | | | |
| Neutral 241 species | IRM | 1 | Primatomorpha | 74.17 | 70.55 | 76.60 | | | | | | | |
| No DNA | ARM | 316 | Primatomorpha | 70.75 | 65.49 | 75.82 | 0.03 | 70.57 | 70.81 | 65.44 | 75.88 | 65.49 | 75.82 |
| One Stratigraphic Bound | IRM | 316 | Primatomorpha | 73.24 | 68.19 | 76.44 | 0.48 | 70.76 | 74.29 | 65.62 | 76.64 | 68.21 | 76.45 |
| Root | IRM | 1 | Primatomorpha | 73.98 | 70.53 | 81.50 | | | | | | | |

| Analysis | Model | Number of Trees | node | average | L95 CI mean | U95 CI mean | std dev | mean min | mean max | L95 CI min | U95 CI max | L95 CI median | U95 CI median |
|---|---|---|---|---|---|---|---|---|---|---|---|---|---|
| Root Calibration Only | ARM | 316 | Primatomorpha | 97.77 | 74.67 | 116.78 | 5.59 | 71.21 | 112.01 | 45.36 | 127.92 | 75.60 | 117.36 |
| all Benton All6 Extended | ARM | 316 | Primatomorpha | 83.50 | 79.24 | 87.90 | 1.73 | 76.16 | 91.16 | 70.58 | 98.22 | 79.13 | 87.44 |
| Benton All6 Extended | IRM | 316 | Primatomorpha | 88.78 | 78.09 | 99.96 | 1.56 | 79.24 | 95.72 | 66.57 | 109.73 | 78.30 | 99.77 |
| Benton All6 Extended NoDNA | ARM | 316 | Primatomorpha | 87.13 | 62.74 | 118.21 | 0.97 | 79.46 | 95.22 | 60.49 | 133.12 | 62.65 | 117.87 |
| all Benton All6 Extended NoDNA | IRM | 316 | Primatomorpha | 87.15 | 62.64 | 118.28 | 0.64 | 80.39 | 90.51 | 60.30 | 128.07 | 62.57 | 118.04 |
| all Benton Rodentia Extended | ARM | 316 | Primatomorpha | 78.72 | 73.97 | 83.83 | 1.81 | 69.70 | 85.81 | 66.17 | 93.34 | 73.96 | 83.65 |
| all Benton Rodentia Extended | IRM | 316 | Primatomorpha | 85.64 | 73.82 | 97.37 | 1.90 | 72.91 | 92.10 | 64.89 | 105.41 | 74.71 | 97.28 |
| all Benton Rodentia Extended NoDNA | ARM | 316 | Primatomorpha | 86.01 | 61.71 | 118.02 | 1.70 | 84.24 | 103.49 | 60.20 | 151.45 | 61.55 | 117.40 |
| all Benton Rodentia Extended NoDNA | IRM | 316 | Primatomorpha | 85.74 | 61.64 | 117.61 | 0.90 | 82.82 | 94.46 | 60.09 | 130.77 | 61.56 | 117.36 |
| Body Size | IRM | 316 | Primatomorpha | 73.44 | 69.49 | 76.34 | 0.49 | 70.18 | 74.48 | 65.36 | 76.61 | 69.70 | 76.35 |
| 10% Missingness | IRM | 316 | Rodentia | 65.13 | 62.71 | 66.84 | 0.11 | 64.73 | 65.45 | 61.93 | 67.00 | 62.69 | 66.84 |
| Accelerated | IRM | 1 | Rodentia | 65.40 | 63.21 | 66.97 | | | | | | | |
| Benton | IRM | 316 | Rodentia | 65.22 | 62.88 | 66.89 | 0.10 | 64.89 | 65.49 | 62.11 | 67.04 | 62.87 | 66.88 |
| Benton | ARM | 316 | Rodentia | 65.36 | 63.52 | 66.76 | 0.26 | 64.63 | 66.02 | 62.34 | 67.42 | 63.52 | 66.74 |
| Benton | Averaged IRM & ARM | 1 | Rodentia | 65.29 | 63.20 | 66.82 | | | | | | | |

| Analysis | Model | Number of Trees | node | average | L95 CI mean | U95 CI mean | std dev | mean min | mean max | L95 CI min | U95 CI max | L95 CI median | U95 CI median |
|---|---|---|---|---|---|---|---|---|---|---|---|---|---|
| Cladistic | ARM | 316 | Rodentia | 65.03 | 62.72 | 66.74 | 0.36 | 63.47 | 66.03 | 60.40 | 67.45 | 62.72 | 66.72 |
| Conserved | IRM | 1 | Rodentia | 65.41 | 63.11 | 66.98 | | | | | | | |
| Meredith | IRM | 316 | Rodentia | 65.22 | 62.88 | 66.88 | 0.09 | 64.88 | 65.49 | 62.12 | 67.04 | 62.88 | 66.88 |
| Neutral | IRM | 1 | Rodentia | 65.57 | 61.37 | 68.76 | | | | | | | |
| Neutral 241 species | IRM | 1 | Rodentia | 65.73 | 64.08 | 67.05 | | | | | | | |
| No DNA | ARM | 316 | Rodentia | 62.96 | 58.25 | 66.48 | 0.02 | 62.91 | 63.20 | 58.06 | 66.56 | 58.25 | 66.48 |
| One Stratigraphic Bound | IRM | 316 | Rodentia | 64.89 | 62.30 | 66.73 | 0.15 | 64.43 | 65.33 | 61.14 | 66.96 | 62.27 | 66.73 |
| Root | IRM | 1 | Rodentia | 65.15 | 62.60 | 66.90 | | | | | | | |
| Root Calibration Only | ARM | 316 | Rodentia | 83.43 | 63.13 | 101.02 | 4.63 | 69.77 | 94.40 | 39.03 | 110.08 | 63.75 | 101.45 |
| all Benton All6 Extended | ARM | 316 | Rodentia | 70.02 | 66.77 | 73.36 | 1.27 | 67.47 | 74.19 | 64.97 | 81.99 | 66.52 | 73.22 |
| Benton All6 Extended | IRM | 316 | Rodentia | 73.44 | 69.05 | 76.57 | 0.25 | 72.86 | 74.25 | 67.92 | 77.14 | 69.05 | 76.57 |
| Benton All6 Extended NoDNA | ARM | 316 | Rodentia | 71.21 | 63.54 | 76.25 | 0.07 | 70.16 | 71.48 | 62.65 | 76.57 | 63.54 | 76.25 |
| all Benton All6 Extended NoDNA | IRM | 316 | Rodentia | 71.22 | 63.53 | 76.24 | 0.02 | 71.15 | 71.27 | 63.22 | 76.36 | 63.54 | 76.24 |
| all Benton Rodentia Extended | ARM | 316 | Rodentia | 69.05 | 65.18 | 73.37 | 1.30 | 66.67 | 73.82 | 62.83 | 76.49 | 65.04 | 73.26 |
| all Benton Rodentia Extended | IRM | 316 | Rodentia | 73.20 | 68.27 | 76.59 | 0.28 | 72.60 | 74.39 | 67.11 | 77.29 | 68.23 | 76.56 |

| Analysis | Model | Number of Trees | node | average | L95 CI mean | U95 CI mean | std dev | mean min | mean max | L95 CI min | U95 CI max | L95 CI median | U95 CI median |
|---|---|---|---|---|---|---|---|---|---|---|---|---|---|
| all Benton Rodentia Extended NoDNA | ARM | 316 | Rodentia | 70.68 | 62.18 | 76.18 | 0.12 | 68.97 | 71.06 | 59.29 | 76.37 | 62.20 | 76.18 |
| all Benton Rodentia Extended NoDNA | IRM | 316 | Rodentia | 70.70 | 62.21 | 76.18 | 0.06 | 70.59 | 71.58 | 61.86 | 76.44 | 62.20 | 76.17 |
| Body Size | IRM | 316 | Rodentia | 64.51 | 61.05 | 66.76 | 0.19 | 63.75 | 65.09 | 59.30 | 66.96 | 61.02 | 66.76 |
| 10% Missingness | IRM | 316 | Placentalia (root) | 99.85 | 87.16 | 113.75 | 4.63 | 89.19 | 121.56 | 80.16 | 135.35 | 86.51 | 113.13 |
| Accelerated | IRM | 1 | Placentalia (root) | 101.48 | 90.15 | 115.98 | | | | | | | |
| Benton | IRM | 316 | Placentalia (root) | 107.01 | 92.52 | 122.95 | 4.85 | 98.34 | 127.36 | 85.49 | 139.40 | 91.59 | 122.50 |
| Benton | ARM | 316 | Placentalia (root) | 96.86 | 88.36 | 106.05 | 3.88 | 84.88 | 116.96 | 79.49 | 131.73 | 87.85 | 105.09 |
| Benton | Averaged IRM & ARM | 1 | Placentalia (root) | 101.94 | 90.44 | 114.50 | | | | | | | |
| Cladistic | ARM | 316 | Placentalia (root) | 98.24 | 88.84 | 108.39 | 4.85 | 88.62 | 123.34 | 81.99 | 135.30 | 88.05 | 106.77 |
| Conserved | IRM | 1 | Placentalia (root) | 98.21 | 83.48 | 107.60 | | | | | | | |
| Meredith | IRM | 316 | Placentalia (root) | 105.25 | 93.06 | 117.81 | 3.47 | 97.07 | 117.83 | 85.85 | 128.37 | 92.53 | 117.73 |
| Neutral | IRM | 1 | Placentalia (root) | 104.45 | 91.46 | 119.68 | | | | | | | |
| Neutral 241 species | IRM | 1 | Placentalia (root) | 103.80 | 90.70 | 118.45 | | | | | | | |
| No DNA | ARM | 316 | Placentalia (root) | 105.36 | 74.98 | 134.04 | 0.69 | 99.71 | 107.03 | 66.52 | 136.17 | 75.13 | 134.05 |
| One Stratigraphic Bound | IRM | 316 | Placentalia (root) | 102.24 | 89.04 | 116.90 | 5.03 | 94.20 | 124.75 | 83.02 | 136.97 | 88.22 | 116.05 |

| **Analysis** | **Model** | **Number of Trees** | **node** | **average** | **L95 CI mean** | **U95 CI mean** | **std dev** | **mean min** | **mean max** | **L95 CI min** | **U95 CI max** | **L95 CI median** | **U95 CI median** |
|---|---|---|---|---|---|---|---|---|---|---|---|---|---|
| Root | IRM | 1 | Placentalia (root) | 102.73 | 88.99 | 115.01 | | | | | | | |
| Root Calibration Only | ARM | 316 | Placentalia (root) | 116.83 | 89.73 | 137.83 | 6.03 | 96.07 | 128.76 | 54.45 | 149.17 | 90.61 | 137.96 |
| all Benton All6 Extended | ARM | 316 | Placentalia (root) | 105.51 | 95.85 | 115.94 | 4.70 | 94.47 | 126.06 | 87.72 | 136.51 | 95.04 | 114.82 |
| Benton All6 Extended | IRM | 316 | Placentalia (root) | 115.08 | 101.25 | 130.02 | 4.30 | 106.59 | 130.81 | 94.09 | 142.24 | 100.50 | 130.40 |
| Benton All6 Extended NoDNA | ARM | 316 | Placentalia (root) | 111.99 | 80.75 | 136.43 | 0.92 | 106.12 | 122.34 | 75.02 | 139.55 | 80.71 | 136.43 |
| all Benton All6 Extended NoDNA | IRM | 316 | Placentalia (root) | 112.00 | 80.74 | 136.50 | 0.62 | 103.88 | 114.87 | 74.47 | 138.47 | 80.76 | 136.50 |
| all Benton Rodentia Extended | ARM | 316 | Placentalia (root) | 100.73 | 90.87 | 111.43 | 4.35 | 88.61 | 120.70 | 82.42 | 134.06 | 90.17 | 110.44 |
| all Benton Rodentia Extended | IRM | 316 | Placentalia (root) | 112.21 | 98.03 | 127.74 | 4.41 | 102.86 | 129.28 | 90.42 | 140.71 | 96.91 | 127.72 |
| all Benton Rodentia Extended NoDNA | ARM | 316 | Placentalia (root) | 111.63 | 79.76 | 136.44 | 0.95 | 108.03 | 118.48 | 72.60 | 156.55 | 79.85 | 136.41 |
| all Benton Rodentia Extended NoDNA | IRM | 316 | Placentalia (root) | 111.44 | 79.73 | 136.48 | 0.60 | 104.66 | 114.60 | 76.36 | 139.12 | 79.82 | 136.50 |
| Body Size | IRM | 316 | Placentalia (root) | 96.29 | 85.35 | 108.20 | 4.96 | 89.39 | 123.02 | 80.54 | 135.67 | 84.47 | 106.97 |
| 10% Missingness | IRM | 316 | Scrotifera | 81.86 | 73.53 | 90.27 | 2.91 | 74.53 | 91.28 | 69.80 | 103.62 | 72.96 | 89.80 |
| Accelerated | IRM | 1 | Scrotifera | 85.01 | 77.80 | 91.99 | | | | | | | |
| Benton | IRM | 316 | Scrotifera | 85.91 | 76.07 | 95.43 | 3.16 | 77.73 | 96.01 | 71.04 | 108.18 | 75.54 | 94.83 |
| Benton | ARM | 316 | Scrotifera | 77.33 | 73.83 | 81.15 | 1.96 | 73.36 | 84.75 | 71.15 | 92.05 | 73.55 | 80.66 |

| Analysis | Model | Number of Trees | node | average | L95 CI mean | U95 CI mean | std dev | mean min | mean max | L95 CI min | U95 CI max | L95 CI median | U95 CI median |
|---|---|---|---|---|---|---|---|---|---|---|---|---|---|
| Benton | Averaged IRM & ARM | 1 | Scrotifera | 81.62 | 74.95 | 88.29 | | | | | | | |
| Cladistic | ARM | 316 | Scrotifera | 78.09 | 72.57 | 84.10 | 3.12 | 72.02 | 92.09 | 69.12 | 101.76 | 72.16 | 83.23 |
| Conserved | IRM | 1 | Scrotifera | 80.34 | 73.00 | 89.79 | | | | | | | |
| Meredith | IRM | 316 | Scrotifera | 85.46 | 75.95 | 94.57 | 2.74 | 77.64 | 93.45 | 70.94 | 103.22 | 75.49 | 94.26 |
| Neutral | IRM | 1 | Scrotifera | 86.28 | 79.03 | 95.16 | | | | | | | |
| Neutral 241 species | IRM | 1 | Scrotifera | 86.68 | 78.23 | 97.40 | | | | | | | |
| No DNA | ARM | 316 | Scrotifera | 88.05 | 65.59 | 112.17 | 0.43 | 83.37 | 88.80 | 65.03 | 113.00 | 65.59 | 112.22 |
| One Stratigraphic Bound | IRM | 316 | Scrotifera | 80.20 | 72.75 | 88.21 | 3.14 | 74.17 | 91.05 | 69.57 | 101.94 | 72.21 | 87.52 |
| Root | IRM | 1 | Scrotifera | 84.94 | 74.28 | 93.56 | | | | | | | |
| Root Calibration Only | ARM | 316 | Scrotifera | 96.68 | 73.94 | 115.48 | 5.17 | 80.54 | 109.72 | 44.44 | 125.17 | 74.90 | 115.50 |
| all Benton All6 Extended | ARM | 316 | Scrotifera | 84.84 | 79.76 | 90.07 | 2.88 | 78.40 | 95.04 | 74.44 | 102.95 | 79.37 | 89.44 |
| Benton All6 Extended | IRM | 316 | Scrotifera | 90.29 | 80.81 | 99.68 | 3.08 | 83.40 | 100.43 | 73.27 | 111.89 | 80.56 | 98.80 |
| Benton All6 Extended NoDNA | ARM | 316 | Scrotifera | 94.77 | 70.39 | 122.98 | 0.74 | 91.01 | 101.42 | 68.88 | 133.31 | 70.34 | 122.85 |
| all Benton All6 Extended NoDNA | IRM | 316 | Scrotifera | 94.77 | 70.38 | 123.09 | 0.49 | 89.02 | 97.19 | 68.10 | 128.24 | 70.36 | 123.01 |
| all Benton Rodentia Extended | ARM | 316 | Scrotifera | 79.61 | 75.32 | 84.20 | 2.30 | 75.44 | 88.47 | 72.28 | 96.48 | 74.88 | 83.50 |

| Analysis | Model | Number of Trees | node | average | L95 CI mean | U95 CI mean | std dev | mean min | mean max | L95 CI min | U95 CI max | L95 CI median | U95 CI median |
|---|---|---|---|---|---|---|---|---|---|---|---|---|---|
| all Benton Rodentia Extended | IRM | 316 | Scrotifera | 86.81 | 76.34 | 96.75 | 3.29 | 79.13 | 97.73 | 71.43 | 110.28 | 75.95 | 95.97 |
| all Benton Rodentia Extended NoDNA | ARM | 316 | Scrotifera | 93.52 | 67.02 | 122.83 | 1.23 | 91.68 | 104.45 | 65.60 | 153.72 | 66.94 | 122.40 |
| all Benton Rodentia Extended NoDNA | IRM | 316 | Scrotifera | 93.26 | 66.92 | 122.51 | 0.63 | 87.60 | 97.86 | 65.49 | 130.79 | 66.89 | 122.40 |
| Body Size | IRM | 316 | Scrotifera | 79.63 | 71.72 | 87.85 | 2.58 | 73.86 | 89.16 | 67.88 | 100.56 | 71.36 | 87.33 |
| 10% Missingness | IRM | 316 | Xenarthra | 58.66 | 48.85 | 66.35 | 0.44 | 57.42 | 61.44 | 47.75 | 67.15 | 48.78 | 66.35 |
| Accelerated | IRM | 1 | Xenarthra | 58.69 | 48.80 | 66.27 | | | | | | | |
| Benton | IRM | 316 | Xenarthra | 58.69 | 48.88 | 66.36 | 0.38 | 57.60 | 60.34 | 48.30 | 66.77 | 48.81 | 66.35 |
| Benton | ARM | 316 | Xenarthra | 60.11 | 50.63 | 66.83 | 0.74 | 56.40 | 62.18 | 47.62 | 67.56 | 50.51 | 66.84 |
| Benton | Averaged IRM & ARM | 1 | Xenarthra | 59.40 | 49.75 | 66.60 | | | | | | | |
| Cladistic | ARM | 316 | Xenarthra | 63.31 | 49.75 | 75.52 | 1.68 | 54.36 | 71.50 | 36.10 | 85.15 | 50.33 | 75.33 |
| Conserved | IRM | 1 | Xenarthra | 59.02 | 49.19 | 66.41 | | | | | | | |
| Meredith | IRM | 316 | Xenarthra | 58.62 | 48.82 | 66.34 | 0.36 | 57.76 | 60.32 | 48.16 | 66.83 | 48.76 | 66.33 |
| Neutral | IRM | 1 | Xenarthra | 58.94 | 48.59 | 66.43 | | | | | | | |
| Neutral 241 species | IRM | 1 | Xenarthra | 57.73 | 48.11 | 66.15 | | | | | | | |
| No DNA | ARM | 316 | Xenarthra | 58.42 | 48.50 | 66.41 | 0.02 | 58.35 | 58.49 | 48.38 | 66.55 | 48.50 | 66.41 |

| Analysis | Model | Number of Trees | node | average | L95 CI mean | U95 CI mean | std dev | mean min | mean max | L95 CI min | U95 CI max | L95 CI median | U95 CI median |
|---|---|---|---|---|---|---|---|---|---|---|---|---|---|
| One Stratigraphic Bound | IRM | 316 | Xenarthra | 58.30 | 48.70 | 66.33 | 0.38 | 57.30 | 59.93 | 47.60 | 66.75 | 48.63 | 66.32 |
| Root | IRM | 1 | Xenarthra | 57.94 | 48.39 | 66.20 | | | | | | | |
| Root Calibration Only | ARM | 316 | Xenarthra | 68.49 | 47.09 | 89.28 | 4.69 | 49.92 | 80.70 | 28.41 | 101.63 | 47.59 | 89.42 |
| all Benton All6 Extended | ARM | 316 | Xenarthra | 60.88 | 51.43 | 67.11 | 0.76 | 57.64 | 63.06 | 48.60 | 67.69 | 51.35 | 67.11 |
| Benton All6 Extended | IRM | 316 | Xenarthra | 59.11 | 49.07 | 66.66 | 0.44 | 57.87 | 60.77 | 48.22 | 67.16 | 48.99 | 66.64 |
| Benton All6 Extended NoDNA | ARM | 316 | Xenarthra | 58.48 | 48.45 | 66.45 | 0.02 | 58.42 | 58.62 | 48.34 | 66.55 | 48.45 | 66.45 |
| all Benton All6 Extended NoDNA | IRM | 316 | Xenarthra | 58.47 | 48.45 | 66.45 | 0.02 | 58.14 | 58.53 | 48.34 | 66.55 | 48.45 | 66.45 |
| all Benton Rodentia Extended | ARM | 316 | Xenarthra | 60.22 | 50.69 | 66.95 | 0.72 | 57.04 | 62.26 | 47.97 | 67.60 | 50.55 | 66.96 |
| all Benton Rodentia Extended | IRM | 316 | Xenarthra | 58.77 | 48.91 | 66.54 | 0.42 | 57.74 | 60.27 | 48.16 | 67.00 | 48.81 | 66.52 |
| all Benton Rodentia Extended NoDNA | ARM | 316 | Xenarthra | 58.40 | 48.45 | 66.41 | 0.05 | 57.54 | 58.45 | 48.08 | 66.56 | 48.45 | 66.41 |
| all Benton Rodentia Extended NoDNA | IRM | 316 | Xenarthra | 58.40 | 48.45 | 66.41 | 0.02 | 58.32 | 58.74 | 48.28 | 66.53 | 48.45 | 66.41 |
| Body Size | IRM | 316 | Xenarthra | 56.76 | 47.79 | 65.40 | 0.67 | 55.19 | 59.81 | 47.21 | 66.64 | 47.65 | 65.36 |

### Data S1.

Hypotheses file used to find deletions supporting alternative hypotheses. Each hypothesis is named using a fasta-like notation - >hypothesis_1 All species in the test clade | Required species from the test clade | Outgroups. The first section contains all species associated with the clade to be tested, the second section lists the species that are required to share the deletion for the deletion to be reported, this allows for the exclusion of genomes which are low coverage or less continuous. The final section lists outgroups, which are required to have continuous sequence through candidate deletion region.

>Scrotifera
Manis_pentadactyla, Manis_javanica, Ursus_maritimus, Ailuropoda_melanoleuca, Neomonachus_schauinslandi, Mirounga_angustirostris, Leptonychotes_weddellii, Odobenus_rosmarus, Zalophus_californianus, Spilogale_gracilis, Ailurus_fulgens, Mellivora_capensis, Mustela_putorius, Enhydra_lutris, Pteronura_brasiliensis, Vulpes_lagopus, Canis_lupus_familiaris, Canis_lupus, Lycaon_pictus, Acinonyx_jubatus, Puma_concolor, Felis_nigripes, Felis_catus, Panthera_onca, Panthera_pardus, Panthera_tigris, Paradoxurus_hermaphroditus, Hyaena_hyaena, Helogale_parvula, Suricata_suricatta, Mungos_mungo, Cryptoprocta_ferox, Ceratotherium_simum_cottoni, Ceratotherium_simum, Diceros_bicornis, Dicerorhinus_sumatrensis, Tapirus_terrestris, Tapirus_indicus, Equus_asinus, Equus_przewalskii, Equus_caballus, Sus_scrofa, Catagonus_wagneri, Elaphurus_davidianus, Rangifer_tarandus, Odocoileus_virginianus, Antilocapra_americana, Giraffa_tippelskirchi, Okapia_johnstoni, Saiga_tatarica, Pantholops_hodgsonii, Ovis_canadensis, Ovis_aries, Capra_aegagrus, Capra_hircus, Hemitragus_hylocrius, Ammotragus_lervia, Beatragus_hunteri, Bison_bison, Bos_mutus, Bos_indicus, Bos_taurus, Bubalus_bubalis, Moschus_moschiferus, Tragulus_javanicus, Orcinus_orca, Tursiops_truncatus, Monodon_monoceros, Delphinapterus_leucas, Phocoena_phocoena, Neophocaena_asiaeorientalis, Lipotes_vexillifer, Inia_geoffrensis, Ziphius_cavirostris, Mesoplodon_bidens, Platanista_gangetica, Kogia_breviceps, Eschrichtius_robustus, Balaenoptera_acutorostrata, Balaenoptera_bonaerensis, Eubalaena_japonica, Hippopotamus_amphibius, Camelus_bactrianus, Camelus_ferus, Camelus_dromedarius, Vicugna_pacos, Megaderma_lyra, Craseonycteris_thonglongyai, Hipposideros_armiger, Hipposideros_galeritus, Rhinolophus_sinicus, Macroglossus_sobrinus, Eidolon_helvum, Pteropus_vampyrus, Pteropus_alecto, Rousettus_aegyptiacus, Noctilio_leporinus, Pteronotus_parnellii, Mormoops_blainvillei, Carollia_perspicillata, Artibeus_jamaicensis, Anoura_caudifer, Tonatia_saurophila, Micronycteris_hirsuta, Desmodus_rotundus, Pipistrellus_pipistrellus, Eptesicus_fuscus, Lasiurus_borealis, Murina_feae, Myotis_myotis, Myotis_davidii, Myotis_lucifugus, Myotis_brandtii, Miniopterus_natalensis, Miniopterus_schreibersii, Tadarida_brasiliensis | Manis_pentadactyla, Manis_javanica, Ursus_maritimus, Ailuropoda_melanoleuca, Neomonachus_schauinslandi, Mirounga_angustirostris, Leptonychotes_weddellii, Odobenus_rosmarus, Zalophus_californianus, Spilogale_gracilis, Ailurus_fulgens, Mellivora_capensis, Mustela_putorius, Enhydra_lutris, Pteronura_brasiliensis, Vulpes_lagopus, Canis_lupus_familiaris, Canis_lupus, Lycaon_pictus, Acinonyx_jubatus, Puma_concolor, Felis_nigripes, Felis_catus, Panthera_onca, Panthera_pardus, Panthera_tigris, Paradoxurus_hermaphroditus, Hyaena_hyaena, Helogale_parvula, Suricata_suricatta, Mungos_mungo, Cryptoprocta_ferox, Ceratotherium_simum_cottoni, Ceratotherium_simum, Diceros_bicornis, Dicerorhinus_sumatrensis, Tapirus_terrestris, Tapirus_indicus, Equus_asinus, Equus_przewalskii, Equus_caballus, Sus_scrofa, Catagonus_wagneri, Elaphurus_davidianus, Rangifer_tarandus, Odocoileus_virginianus, Antilocapra_americana, Giraffa_tippelskirchi, Okapia_johnstoni, Saiga_tatarica, Pantholops_hodgsonii, Ovis_canadensis, Ovis_aries, Capra_aegagrus, Capra_hircus, Hemitragus_hylocrius, Ammotragus_lervia, Beatragus_hunteri, Bison_bison, Bos_mutus, Bos_indicus, Bos_taurus, Bubalus_bubalis, Moschus_moschiferus, Tragulus_javanicus, Orcinus_orca, Tursiops_truncatus, Monodon_monoceros, Delphinapterus_leucas, Phocoena_phocoena, Neophocaena_asiaeorientalis, Lipotes_vexillifer, Inia_geoffrensis, Ziphius_cavirostris, Mesoplodon_bidens, Platanista_gangetica, Kogia_breviceps, Eschrichtius_robustus, Balaenoptera_acutorostrata, Balaenoptera_bonaerensis, Eubalaena_japonica, Hippopotamus_amphibius, Camelus_bactrianus, Camelus_ferus, Camelus_dromedarius, Vicugna_pacos, Megaderma_lyra, Craseonycteris_thonglongyai, Hipposideros_armiger, Hipposideros_galeritus, Rhinolophus_sinicus, Macroglossus_sobrinus, Eidolon_helvum, Pteropus_vampyrus, Pteropus_alecto, Rousettus_aegyptiacus, Noctilio_leporinus, Pteronotus_parnellii, Mormoops_blainvillei, Carollia_perspicillata, Artibeus_jamaicensis, Anoura_caudifer, Tonatia_saurophila, Micronycteris_hirsuta, Desmodus_rotundus, Pipistrellus_pipistrellus, Eptesicus_fuscus, Lasiurus_borealis, Murina_feae, Myotis_myotis, Myotis_davidii, Myotis_lucifugus, Myotis_brandtii, Miniopterus_natalensis, Miniopterus_schreibersii, Tadarida_brasiliensis | Erinaceus_europaeus, Solenodon_paradoxus, Homo_sapiens, Mus_musculus

>Fereuungulata
Manis_pentadactyla, Manis_javanica, Ursus_maritimus, Ailuropoda_melanoleuca, Neomonachus_schauinslandi, Mirounga_angustirostris, Leptonychotes_weddellii, Odobenus_rosmarus, Zalophus_californianus, Spilogale_gracilis, Ailurus_fulgens, Mellivora_capensis, Mustela_putorius, Enhydra_lutris, Pteronura_brasiliensis, Vulpes_lagopus, Canis_lupus_familiaris, Canis_lupus, Lycaon_pictus, Acinonyx_jubatus, Puma_concolor, Felis_nigripes, Felis_catus, Panthera_onca, Panthera_pardus, Panthera_tigris, Paradoxurus_hermaphroditus, Hyaena_hyaena, Helogale_parvula, Suricata_suricatta, Mungos_mungo, Cryptoprocta_ferox, Ceratotherium_simum_cottoni, Ceratotherium_simum, Diceros_bicornis, Dicerorhinus_sumatrensis, Tapirus_terrestris, Tapirus_indicus, Equus_asinus, Equus_przewalskii, Equus_caballus, Sus_scrofa, Catagonus_wagneri, Elaphurus_davidianus, Rangifer_tarandus, Odocoileus_virginianus, Antilocapra_americana, Giraffa_tippelskirchi, Okapia_johnstoni, Saiga_tatarica, Pantholops_hodgsonii, Ovis_canadensis, Ovis_aries, Capra_aegagrus, Capra_hircus, Hemitragus_hylocrius, Ammotragus_lervia, Beatragus_hunteri, Bison_bison, Bos_mutus, Bos_indicus, Bos_taurus, Bubalus_bubalis, Moschus_moschiferus, Tragulus_javanicus, Orcinus_orca, Tursiops_truncatus, Monodon_monoceros, Delphinapterus_leucas, Phocoena_phocoena, Neophocaena_asiaeorientalis, Lipotes_vexillifer, Inia_geoffrensis, Ziphius_cavirostris, Mesoplodon_bidens, Platanista_gangetica, Kogia_breviceps, Eschrichtius_robustus, Balaenoptera_acutorostrata, Balaenoptera_bonaerensis, Eubalaena_japonica, Hippopotamus_amphibius, Camelus_bactrianus, Camelus_ferus, Camelus_dromedarius, Vicugna_pacos | Manis_pentadactyla, Manis_javanica, Ursus_maritimus, Ailuropoda_melanoleuca, Neomonachus_schauinslandi, Mirounga_angustirostris, Leptonychotes_weddellii, Odobenus_rosmarus, Zalophus_californianus, Spilogale_gracilis, Ailurus_fulgens, Mellivora_capensis, Mustela_putorius, Enhydra_lutris, Pteronura_brasiliensis, Vulpes_lagopus, Canis_lupus_familiaris, Canis_lupus, Lycaon_pictus, Acinonyx_jubatus, Puma_concolor, Felis_nigripes, Felis_catus, Panthera_onca, Panthera_pardus, Panthera_tigris, Paradoxurus_hermaphroditus, Hyaena_hyaena, Helogale_parvula,

Suricata_suricatta, Mungos_mungo, Cryptoprocta_ferox, Ceratotherium_simum_cottoni, Ceratotherium_simum, Diceros_bicornis, Dicerorhinus_sumatrensis, Tapirus_terrestris, Tapirus_indicus, Equus_asinus, Equus_przewalskii, Equus_caballus, Sus_scrofa, Catagonus_wagneri, Elaphurus_davidianus, Rangifer_tarandus, Odocoileus_virginianus, Antilocapra_americana, Giraffa_tippelskirchi, Okapia_johnstoni, Saiga_tatarica, Pantholops_hodgsonii, Ovis_canadensis, Ovis_aries, Capra_aegagrus, Capra_hircus, Hemitragus_hylocrius, Ammotragus_lervia, Beatragus_hunteri, Bison_bison, Bos_mutus, Bos_indicus, Bos_taurus, Bubalus_bubalis, Moschus_moschiferus, Tragulus_javanicus, Orcinus_orca, Tursiops_truncatus, Monodon_monoceros, Delphinapterus_leucas, Phocoena_phocoena, Neophocaena_asiaeorientalis, Lipotes_vexillifer, Inia_geoffrensis, Ziphius_cavirostris, Mesoplodon_bidens, Platanista_gangetica, Kogia_breviceps, Eschrichtius_robustus, Balaenoptera_acutorostrata, Balaenoptera_bonaerensis, Eubalaena_japonica, Hippopotamus_amphibius, Camelus_bactrianus, Camelus_ferus, Camelus_dromedarius, Vicugna_pacos | Macroglossus_sobrinus, Erinaceus_europaeus, Homo_sapiens, Mus_musculus

>Euungulata_PerissodactylaCetartiodactyla
Ceratotherium_simum_cottoni, Ceratotherium_simum, Diceros_bicornis, Dicerorhinus_sumatrensis, Tapirus_terrestris, Tapirus_indicus, Equus_asinus, Equus_przewalskii, Equus_caballus, Sus_scrofa, Catagonus_wagneri, Elaphurus_davidianus, Rangifer_tarandus, Odocoileus_virginianus, Antilocapra_americana, Giraffa_tippelskirchi, Okapia_johnstoni, Saiga_tatarica, Pantholops_hodgsonii, Ovis_canadensis, Ovis_aries, Capra_aegagrus, Capra_hircus, Hemitragus_hylocrius, Ammotragus_lervia, Beatragus_hunteri, Bison_bison, Bos_mutus, Bos_indicus, Bos_taurus, Bubalus_bubalis, Moschus_moschiferus, Tragulus_javanicus, Orcinus_orca, Tursiops_truncatus, Monodon_monoceros, Delphinapterus_leucas, Phocoena_phocoena, Neophocaena_asiaeorientalis, Lipotes_vexillifer, Inia_geoffrensis, Ziphius_cavirostris, Mesoplodon_bidens, Platanista_gangetica, Kogia_breviceps, Eschrichtius_robustus, Balaenoptera_acutorostrata, Balaenoptera_bonaerensis, Eubalaena_japonica, Hippopotamus_amphibius, Camelus_bactrianus, Camelus_ferus, Camelus_dromedarius, Vicugna_pacos | Ceratotherium_simum_cottoni, Ceratotherium_simum, Diceros_bicornis, Dicerorhinus_sumatrensis, Tapirus_terrestris, Tapirus_indicus, Equus_asinus, Equus_przewalskii, Equus_caballus, Sus_scrofa, Catagonus_wagneri, Elaphurus_davidianus, Rangifer_tarandus, Odocoileus_virginianus, Antilocapra_americana, Giraffa_tippelskirchi, Okapia_johnstoni, Saiga_tatarica, Pantholops_hodgsonii, Ovis_canadensis, Ovis_aries, Capra_aegagrus, Capra_hircus, Hemitragus_hylocrius, Ammotragus_lervia, Beatragus_hunteri, Bison_bison, Bos_mutus, Bos_indicus, Bos_taurus, Bubalus_bubalis, Moschus_moschiferus, Tragulus_javanicus, Orcinus_orca, Tursiops_truncatus, Monodon_monoceros, Delphinapterus_leucas, Phocoena_phocoena, Neophocaena_asiaeorientalis, Lipotes_vexillifer, Inia_geoffrensis, Ziphius_cavirostris, Mesoplodon_bidens, Platanista_gangetica, Kogia_breviceps, Eschrichtius_robustus, Balaenoptera_acutorostrata, Balaenoptera_bonaerensis, Eubalaena_japonica, Hippopotamus_amphibius, Camelus_bactrianus, Camelus_ferus, Camelus_dromedarius, Vicugna_pacos | Canis_lupus_familiaris, Felis_catus, Macroglossus_sobrinus, Erinaceus_europaeus, Homo_sapiens, Mus_musculus

>Ferae_Cetartiodactyla
Manis_pentadactyla, Manis_javanica, Ursus_maritimus, Ailuropoda_melanoleuca, Neomonachus_schauinslandi, Mirounga_angustirostris, Leptonychotes_weddellii, Odobenus_rosmarus, Zalophus_californianus, Spilogale_gracilis, Ailurus_fulgens, Mellivora_capensis, Mustela_putorius, Enhydra_lutris, Pteronura_brasiliensis, Vulpes_lagopus, Canis_lupus_familiaris, Canis_lupus, Lycaon_pictus, Acinonyx_jubatus, Puma_concolor, Felis_nigripes, Felis_catus, Panthera_onca, Panthera_pardus, Panthera_tigris, Paradoxurus_hermaphroditus, Hyaena_hyaena, Helogale_parvula, Suricata_suricatta, Mungos_mungo, Cryptoprocta_ferox, Sus_scrofa, Catagonus_wagneri, Elaphurus_davidianus, Rangifer_tarandus, Odocoileus_virginianus, Antilocapra_americana, Giraffa_tippelskirchi, Okapia_johnstoni, Saiga_tatarica, Pantholops_hodgsonii, Ovis_canadensis, Ovis_aries, Capra_aegagrus, Capra_hircus, Hemitragus_hylocrius, Ammotragus_lervia, Beatragus_hunteri, Bison_bison, Bos_mutus, Bos_indicus, Bos_taurus, Bubalus_bubalis, Moschus_moschiferus, Tragulus_javanicus, Orcinus_orca, Tursiops_truncatus, Monodon_monoceros, Delphinapterus_leucas, Phocoena_phocoena, Neophocaena_asiaeorientalis, Lipotes_vexillifer, Inia_geoffrensis, Ziphius_cavirostris, Mesoplodon_bidens, Platanista_gangetica, Kogia_breviceps, Eschrichtius_robustus, Balaenoptera_acutorostrata, Balaenoptera_bonaerensis, Eubalaena_japonica, Hippopotamus_amphibius, Camelus_bactrianus, Camelus_ferus, Camelus_dromedarius, Vicugna_pacos | Manis_pentadactyla, Manis_javanica, Ursus_maritimus, Ailuropoda_melanoleuca, Neomonachus_schauinslandi, Mirounga_angustirostris, Leptonychotes_weddellii, Odobenus_rosmarus, Zalophus_californianus, Spilogale_gracilis, Ailurus_fulgens, Mellivora_capensis, Mustela_putorius, Enhydra_lutris, Pteronura_brasiliensis, Vulpes_lagopus, Canis_lupus_familiaris, Canis_lupus, Lycaon_pictus, Acinonyx_jubatus, Puma_concolor, Felis_nigripes, Felis_catus, Panthera_onca, Panthera_pardus, Panthera_tigris, Paradoxurus_hermaphroditus, Hyaena_hyaena, Helogale_parvula, Suricata_suricatta, Mungos_mungo, Cryptoprocta_ferox, Sus_scrofa, Catagonus_wagneri, Elaphurus_davidianus, Rangifer_tarandus, Odocoileus_virginianus, Antilocapra_americana, Giraffa_tippelskirchi, Okapia_johnstoni, Saiga_tatarica, Pantholops_hodgsonii, Ovis_canadensis, Ovis_aries, Capra_aegagrus, Capra_hircus, Hemitragus_hylocrius, Ammotragus_lervia, Beatragus_hunteri, Bison_bison, Bos_mutus, Bos_indicus, Bos_taurus, Bubalus_bubalis, Moschus_moschiferus, Tragulus_javanicus, Orcinus_orca, Tursiops_truncatus, Monodon_monoceros, Delphinapterus_leucas, Phocoena_phocoena, Neophocaena_asiaeorientalis, Lipotes_vexillifer, Inia_geoffrensis, Ziphius_cavirostris, Mesoplodon_bidens, Platanista_gangetica, Kogia_breviceps, Eschrichtius_robustus, Balaenoptera_acutorostrata, Balaenoptera_bonaerensis, Eubalaena_japonica, Hippopotamus_amphibius, Camelus_bactrianus, Camelus_ferus, Camelus_dromedarius, Vicugna_pacos | Equus_caballus, Macroglossus_sobrinus, Erinaceus_europaeus, Homo_sapiens, Mus_musculus

>Ferae_Perissodactyla
Manis_pentadactyla, Manis_javanica, Ursus_maritimus, Ailuropoda_melanoleuca, Neomonachus_schauinslandi, Mirounga_angustirostris, Leptonychotes_weddellii, Odobenus_rosmarus, Zalophus_californianus, Spilogale_gracilis, Ailurus_fulgens, Mellivora_capensis, Mustela_putorius, Enhydra_lutris, Pteronura_brasiliensis, Vulpes_lagopus, Canis_lupus_familiaris, Canis_lupus, Lycaon_pictus, Acinonyx_jubatus, Puma_concolor, Felis_nigripes, Felis_catus, Panthera_onca, Panthera_pardus, Panthera_tigris, Paradoxurus_hermaphroditus, Hyaena_hyaena, Helogale_parvula, Suricata_suricatta, Mungos_mungo, Cryptoprocta_ferox, Ceratotherium_simum_cottoni, Ceratotherium_simum, Diceros_bicornis, Dicerorhinus_sumatrensis, Tapirus_terrestris, Tapirus_indicus, Equus_asinus, Equus_przewalskii, Equus_caballus | Manis_pentadactyla, Manis_javanica, Ursus_maritimus, Ailuropoda_melanoleuca, Neomonachus_schauinslandi, Mirounga_angustirostris, Leptonychotes_weddellii, Odobenus_rosmarus, Zalophus_californianus, Spilogale_gracilis, Ailurus_fulgens, Mellivora_capensis, Mustela_putorius, Enhydra_lutris, Pteronura_brasiliensis, Vulpes_lagopus, Canis_lupus_familiaris, Canis_lupus, Lycaon_pictus, Acinonyx_jubatus, Puma_concolor, Felis_nigripes, Felis_catus, Panthera_onca, Panthera_pardus, Panthera_tigris, Paradoxurus_hermaphroditus, Hyaena_hyaena, Helogale_parvula, Suricata_suricatta, Mungos_mungo, Cryptoprocta_ferox, Ceratotherium_simum_cottoni, Ceratotherium_simum, Diceros_bicornis, Dicerorhinus_sumatrensis, Tapirus_terrestris, Tapirus_indicus, Equus_asinus, Equus_przewalskii, Equus_caballus | Bos_taurus, Macroglossus_sobrinus, Erinaceus_europaeus, Homo_sapiens, Mus_musculus

>Pegasoferae
Manis_pentadactyla, Manis_javanica, Ursus_maritimus, Ailuropoda_melanoleuca, Neomonachus_schauinslandi, Mirounga_angustirostris, Leptonychotes_weddellii, Odobenus_rosmarus, Zalophus_californianus, Spilogale_gracilis, Ailurus_fulgens, Mellivora_capensis, Mustela_putorius, Enhydra_lutris, Pteronura_brasiliensis, Vulpes_lagopus, Canis_lupus_familiaris, Canis_lupus, Lycaon_pictus, Acinonyx_jubatus, Puma_concolor, Felis_nigripes, Felis_catus, Panthera_onca, Panthera_pardus, Panthera_tigris, Paradoxurus_hermaphroditus, Hyaena_hyaena, Helogale_parvula, Suricata_suricatta, Mungos_mungo, Cryptoprocta_ferox, Ceratotherium_simum_cottoni, Ceratotherium_simum, Diceros_bicornis, Dicerorhinus_sumatrensis, Tapirus_terrestris, Tapirus_indicus, Equus_asinus, Equus_przewalskii, Equus_caballus, Megaderma_lyra, Craseonycteris_thonglongyai, Hipposideros_armiger, Hipposideros_galeritus, Rhinolophus_sinicus, Macroglossus_sobrinus, Eidolon_helvum, Pteropus_vampyrus, Pteropus_alecto, Rousettus_aegyptiacus, Noctilio_leporinus, Pteronotus_parnellii, Mormoops_blainvillei, Carollia_perspicillata, Artibeus_jamaicensis, Anoura_caudifer, Tonatia_saurophila, Micronycteris_hirsuta, Desmodus_rotundus, Pipistrellus_pipistrellus, Eptesicus_fuscus, Lasiurus_borealis, Murina_feae, Myotis_myotis, Myotis_davidii, Myotis_lucifugus, Myotis_brandtii, Miniopterus_natalensis, Miniopterus_schreibersii, Tadarida_brasiliensis | Manis_pentadactyla, Manis_javanica, Ursus_maritimus, Ailuropoda_melanoleuca, Neomonachus_schauinslandi, Mirounga_angustirostris, Leptonychotes_weddellii, Odobenus_rosmarus, Zalophus_californianus, Spilogale_gracilis, Ailurus_fulgens, Mellivora_capensis, Mustela_putorius, Enhydra_lutris, Pteronura_brasiliensis, Vulpes_lagopus, Canis_lupus_familiaris, Canis_lupus, Lycaon_pictus, Acinonyx_jubatus, Puma_concolor, Felis_nigripes, Felis_catus, Panthera_onca, Panthera_pardus, Panthera_tigris, Paradoxurus_hermaphroditus, Hyaena_hyaena, Helogale_parvula, Suricata_suricatta, Mungos_mungo, Cryptoprocta_ferox, Ceratotherium_simum_cottoni, Ceratotherium_simum, Diceros_bicornis, Dicerorhinus_sumatrensis, Tapirus_terrestris, Tapirus_indicus, Equus_asinus, Equus_przewalskii, Equus_caballus, Megaderma_lyra, Craseonycteris_thonglongyai, Hipposideros_armiger, Hipposideros_galeritus, Rhinolophus_sinicus, Macroglossus_sobrinus, Eidolon_helvum, Pteropus_vampyrus, Pteropus_alecto, Rousettus_aegyptiacus, Noctilio_leporinus, Pteronotus_parnellii, Mormoops_blainvillei, Carollia_perspicillata, Artibeus_jamaicensis, Anoura_caudifer, Tonatia_saurophila, Micronycteris_hirsuta, Desmodus_rotundus, Pipistrellus_pipistrellus, Eptesicus_fuscus, Lasiurus_borealis, Murina_feae, Myotis_myotis, Myotis_davidii, Myotis_lucifugus, Myotis_brandtii, Miniopterus_natalensis, Miniopterus_schreibersii, Tadarida_brasiliensis | Sus_scrofa, Bos_taurus, Balaenoptera_bonaerensis, Erinaceus_europaeus, Homo_sapiens, Mus_musculus

>Chiroptera_Ferae
Megaderma_lyra, Craseonycteris_thonglongyai, Hipposideros_armiger, Hipposideros_galeritus, Rhinolophus_sinicus, Macroglossus_sobrinus, Eidolon_helvum, Pteropus_vampyrus, Pteropus_alecto, Rousettus_aegyptiacus, Noctilio_leporinus, Pteronotus_parnellii, Mormoops_blainvillei, Carollia_perspicillata, Artibeus_jamaicensis, Anoura_caudifer, Tonatia_saurophila, Micronycteris_hirsuta, Desmodus_rotundus, Pipistrellus_pipistrellus, Eptesicus_fuscus, Lasiurus_borealis, Murina_feae, Myotis_myotis, Myotis_davidii, Myotis_lucifugus, Myotis_brandtii, Miniopterus_natalensis, Miniopterus_schreibersii, Tadarida_brasiliensis, Manis_pentadactyla, Manis_javanica, Ursus_maritimus, Ailuropoda_melanoleuca, Neomonachus_schauinslandi, Mirounga_angustirostris, Leptonychotes_weddellii, Odobenus_rosmarus, Zalophus_californianus, Spilogale_gracilis, Ailurus_fulgens, Mellivora_capensis, Mustela_putorius, Enhydra_lutris, Pteronura_brasiliensis, Vulpes_lagopus, Canis_lupus_familiaris, Canis_lupus, Lycaon_pictus, Acinonyx_jubatus, Puma_concolor, Felis_nigripes, Felis_catus, Panthera_onca, Panthera_pardus, Panthera_tigris, Paradoxurus_hermaphroditus, Hyaena_hyaena, Helogale_parvula, Suricata_suricatta, Mungos_mungo, Cryptoprocta_ferox | Megaderma_lyra, Craseonycteris_thonglongyai, Hipposideros_armiger, Hipposideros_galeritus, Rhinolophus_sinicus, Macroglossus_sobrinus, Eidolon_helvum, Pteropus_vampyrus, Pteropus_alecto, Rousettus_aegyptiacus, Noctilio_leporinus, Pteronotus_parnellii, Mormoops_blainvillei, Carollia_perspicillata, Artibeus_jamaicensis, Anoura_caudifer, Tonatia_saurophila, Micronycteris_hirsuta, Desmodus_rotundus, Pipistrellus_pipistrellus, Eptesicus_fuscus, Lasiurus_borealis, Murina_feae, Myotis_myotis, Myotis_davidii, Myotis_lucifugus, Myotis_brandtii, Miniopterus_natalensis, Miniopterus_schreibersii, Tadarida_brasiliensis, Manis_pentadactyla, Manis_javanica, Ursus_maritimus, Ailuropoda_melanoleuca, Neomonachus_schauinslandi, Mirounga_angustirostris, Leptonychotes_weddellii, Odobenus_rosmarus, Zalophus_californianus, Spilogale_gracilis, Ailurus_fulgens, Mellivora_capensis, Mustela_putorius, Enhydra_lutris, Pteronura_brasiliensis, Vulpes_lagopus, Canis_lupus_familiaris, Canis_lupus, Lycaon_pictus, Acinonyx_jubatus, Puma_concolor, Felis_nigripes, Felis_catus, Panthera_onca, Panthera_pardus, Panthera_tigris, Paradoxurus_hermaphroditus, Hyaena_hyaena, Helogale_parvula, Suricata_suricatta, Mungos_mungo, Cryptoprocta_ferox | Bos_taurus, Equus_caballus, Erinaceus_europaeus, Homo_sapiens, Mus_musculus

>Chiroptera_Perissodactyla
Megaderma_lyra, Craseonycteris_thonglongyai, Hipposideros_armiger, Hipposideros_galeritus, Rhinolophus_sinicus, Macroglossus_sobrinus, Eidolon_helvum, Pteropus_vampyrus, Pteropus_alecto, Rousettus_aegyptiacus, Noctilio_leporinus, Pteronotus_parnellii, Mormoops_blainvillei, Carollia_perspicillata, Artibeus_jamaicensis, Anoura_caudifer, Tonatia_saurophila, Micronycteris_hirsuta, Desmodus_rotundus, Pipistrellus_pipistrellus, Eptesicus_fuscus, Lasiurus_borealis, Murina_feae, Myotis_myotis, Myotis_davidii, Myotis_lucifugus, Myotis_brandtii, Miniopterus_natalensis, Miniopterus_schreibersii, Tadarida_brasiliensis,Ceratotherium_simum_cottoni, Ceratotherium_simum, Diceros_bicornis, Dicerorhinus_sumatrensis, Tapirus_terrestris, Tapirus_indicus, Equus_asinus, Equus_przewalskii, Equus_caballus | Megaderma_lyra, Craseonycteris_thonglongyai, Hipposideros_armiger, Hipposideros_galeritus, Rhinolophus_sinicus, Macroglossus_sobrinus, Eidolon_helvum, Pteropus_vampyrus, Pteropus_alecto, Rousettus_aegyptiacus, Noctilio_leporinus, Pteronotus_parnellii, Mormoops_blainvillei, Carollia_perspicillata, Artibeus_jamaicensis, Anoura_caudifer, Tonatia_saurophila, Micronycteris_hirsuta, Desmodus_rotundus, Pipistrellus_pipistrellus, Eptesicus_fuscus, Lasiurus_borealis, Murina_feae, Myotis_myotis, Myotis_davidii, Myotis_lucifugus, Myotis_brandtii, Miniopterus_natalensis, Miniopterus_schreibersii, Tadarida_brasiliensis,Ceratotherium_simum_cottoni, Ceratotherium_simum, Diceros_bicornis, Dicerorhinus_sumatrensis, Tapirus_terrestris, Tapirus_indicus, Equus_asinus, Equus_przewalskii, Equus_caballus | Bos_taurus, Felis_catus, Erinaceus_europaeus, Homo_sapiens, Mus_musculus

>Chiroptera_Cetartiodactyla
Megaderma_lyra, Craseonycteris_thonglongyai, Hipposideros_armiger, Hipposideros_galeritus, Rhinolophus_sinicus, Macroglossus_sobrinus, Eidolon_helvum, Pteropus_vampyrus, Pteropus_alecto, Rousettus_aegyptiacus, Noctilio_leporinus, Pteronotus_parnellii, Mormoops_blainvillei, Carollia_perspicillata, Artibeus_jamaicensis, Anoura_caudifer, Tonatia_saurophila, Micronycteris_hirsuta, Desmodus_rotundus, Pipistrellus_pipistrellus, Eptesicus_fuscus, Lasiurus_borealis, Murina_feae, Myotis_myotis, Myotis_davidii, Myotis_lucifugus, Myotis_brandtii, Miniopterus_natalensis, Miniopterus_schreibersii, Tadarida_brasiliensis, Sus_scrofa, Catagonus_wagneri, Elaphurus_davidianus, Rangifer_tarandus, Odocoileus_virginianus, Antilocapra_americana, Giraffa_tippelskirchi, Okapia_johnstoni, Saiga_tatarica, Pantholops_hodgsonii, Ovis_canadensis, Ovis_aries, Capra_aegagrus, Capra_hircus, Hemitragus_hylocrius, Ammotragus_lervia,

Beatragus_hunteri, Bison_bison, Bos_mutus, Bos_indicus, Bos_taurus, Bubalus_bubalis, Moschus_moschiferus, Tragulus_javanicus, Orcinus_orca, Tursiops_truncatus, Monodon_monoceros, Delphinapterus_leucas, Phocoena_phocoena, Neophocaena_asiaeorientalis, Lipotes_vexillifer, Inia_geoffrensis, Ziphius_cavirostris, Mesoplodon_bidens, Platanista_gangetica, Kogia_breviceps, Eschrichtius_robustus, Balaenoptera_acutorostrata, Balaenoptera_bonaerensis, Eubalaena_japonica, Hippopotamus_amphibius, Camelus_bactrianus, Camelus_ferus, Camelus_dromedarius, Vicugna_pacos | Megaderma_lyra, Craseonycteris_thonglongyai, Hipposideros_armiger, Hipposideros_galeritus, Rhinolophus_sinicus, Macroglossus_sobrinus, Eidolon_helvum, Pteropus_vampyrus, Pteropus_alecto, Rousettus_aegyptiacus, Noctilio_leporinus, Pteronotus_parnellii, Mormoops_blainvillei, Carollia_perspicillata, Artibeus_jamaicensis, Anoura_caudifer, Tonatia_saurophila, Micronycteris_hirsuta, Desmodus_rotundus, Pipistrellus_pipistrellus, Eptesicus_fuscus, Lasiurus_borealis, Murina_feae, Myotis_myotis, Myotis_davidii, Myotis_lucifugus, Myotis_brandtii, Miniopterus_natalensis, Miniopterus_schreibersii, Tadarida_brasiliensis, Sus_scrofa, Catagonus_wagneri, Elaphurus_davidianus, Rangifer_tarandus, Odocoileus_virginianus, Antilocapra_americana, Giraffa_tippelskirchi, Okapia_johnstoni, Saiga_tatarica, Pantholops_hodgsonii, Ovis_canadensis, Ovis_aries, Capra_aegagrus, Capra_hircus, Hemitragus_hylocrius, Ammotragus_lervia, Beatragus_hunteri, Bison_bison, Bos_mutus, Bos_indicus, Bos_taurus, Bubalus_bubalis, Moschus_moschiferus, Tragulus_javanicus, Orcinus_orca, Tursiops_truncatus, Monodon_monoceros, Delphinapterus_leucas, Phocoena_phocoena, Neophocaena_asiaeorientalis, Lipotes_vexillifer, Inia_geoffrensis, Ziphius_cavirostris, Mesoplodon_bidens, Platanista_gangetica, Kogia_breviceps, Eschrichtius_robustus, Balaenoptera_acutorostrata, Balaenoptera_bonaerensis, Eubalaena_japonica, Hippopotamus_amphibius, Camelus_bactrianus, Camelus_ferus, Camelus_dromedarius, Vicugna_pacos | Equus_caballus, Felis_catus, Erinaceus_europaeus, Homo_sapiens, Mus_musculus

>Chiroptera_Perissodactyla_Cetartiodactyla
Megaderma_lyra, Craseonycteris_thonglongyai, Hipposideros_armiger, Hipposideros_galeritus, Rhinolophus_sinicus, Macroglossus_sobrinus, Eidolon_helvum, Pteropus_vampyrus, Pteropus_alecto, Rousettus_aegyptiacus, Noctilio_leporinus, Pteronotus_parnellii, Mormoops_blainvillei, Carollia_perspicillata, Artibeus_jamaicensis, Anoura_caudifer, Tonatia_saurophila, Micronycteris_hirsuta, Desmodus_rotundus, Pipistrellus_pipistrellus, Eptesicus_fuscus, Lasiurus_borealis, Murina_feae, Myotis_myotis, Myotis_davidii, Myotis_lucifugus, Myotis_brandtii, Miniopterus_natalensis, Miniopterus_schreibersii, Tadarida_brasiliensis, Ceratotherium_simum_cottoni, Ceratotherium_simum, Diceros_bicornis, Dicerorhinus_sumatrensis, Tapirus_terrestris, Tapirus_indicus, Equus_asinus, Equus_przewalskii, Equus_caballus, Sus_scrofa, Catagonus_wagneri, Elaphurus_davidianus, Rangifer_tarandus, Odocoileus_virginianus, Antilocapra_americana, Giraffa_tippelskirchi, Okapia_johnstoni, Saiga_tatarica, Pantholops_hodgsonii, Ovis_canadensis, Ovis_aries, Capra_aegagrus, Capra_hircus, Hemitragus_hylocrius, Ammotragus_lervia, Beatragus_hunteri, Bison_bison, Bos_mutus, Bos_indicus, Bos_taurus, Bubalus_bubalis, Moschus_moschiferus, Tragulus_javanicus, Orcinus_orca, Tursiops_truncatus, Monodon_monoceros, Delphinapterus_leucas, Phocoena_phocoena, Neophocaena_asiaeorientalis, Lipotes_vexillifer, Inia_geoffrensis, Ziphius_cavirostris, Mesoplodon_bidens, Platanista_gangetica, Kogia_breviceps, Eschrichtius_robustus, Balaenoptera_acutorostrata, Balaenoptera_bonaerensis, Eubalaena_japonica, Hippopotamus_amphibius, Camelus_bactrianus, Camelus_ferus, Camelus_dromedarius, Vicugna_pacos | Megaderma_lyra, Craseonycteris_thonglongyai, Hipposideros_armiger, Hipposideros_galeritus, Rhinolophus_sinicus, Macroglossus_sobrinus, Eidolon_helvum, Pteropus_vampyrus, Pteropus_alecto, Rousettus_aegyptiacus, Noctilio_leporinus, Pteronotus_parnellii, Mormoops_blainvillei, Carollia_perspicillata, Artibeus_jamaicensis, Anoura_caudifer, Tonatia_saurophila, Micronycteris_hirsuta, Desmodus_rotundus, Pipistrellus_pipistrellus, Eptesicus_fuscus, Lasiurus_borealis, Murina_feae, Myotis_myotis, Myotis_davidii, Myotis_lucifugus, Myotis_brandtii, Miniopterus_natalensis, Miniopterus_schreibersii, Tadarida_brasiliensis, Ceratotherium_simum_cottoni, Ceratotherium_simum, Diceros_bicornis, Dicerorhinus_sumatrensis, Tapirus_terrestris, Tapirus_indicus, Equus_asinus, Equus_przewalskii, Equus_caballus, Sus_scrofa, Catagonus_wagneri, Elaphurus_davidianus, Rangifer_tarandus, Odocoileus_virginianus, Antilocapra_americana, Giraffa_tippelskirchi, Okapia_johnstoni, Saiga_tatarica, Pantholops_hodgsonii, Ovis_canadensis, Ovis_aries, Capra_aegagrus, Capra_hircus, Hemitragus_hylocrius, Ammotragus_lervia, Beatragus_hunteri, Bison_bison, Bos_mutus, Bos_indicus, Bos_taurus, Bubalus_bubalis, Moschus_moschiferus, Tragulus_javanicus, Orcinus_orca, Tursiops_truncatus, Monodon_monoceros, Delphinapterus_leucas, Phocoena_phocoena, Neophocaena_asiaeorientalis, Lipotes_vexillifer, Inia_geoffrensis, Ziphius_cavirostris, Mesoplodon_bidens, Platanista_gangetica, Kogia_breviceps, Eschrichtius_robustus, Balaenoptera_acutorostrata, Balaenoptera_bonaerensis, Eubalaena_japonica, Hippopotamus_amphibius, Camelus_bactrianus, Camelus_ferus, Camelus_dromedarius, Vicugna_pacos | Felis_catus, Erinaceus_europaeus, Homo_sapiens, Mus_musculus

>Chiroptera_Perissodactyla_Ferae
Megaderma_lyra, Craseonycteris_thonglongyai, Hipposideros_armiger, Hipposideros_galeritus, Rhinolophus_sinicus, Macroglossus_sobrinus, Eidolon_helvum, Pteropus_vampyrus, Pteropus_alecto, Rousettus_aegyptiacus, Noctilio_leporinus, Pteronotus_parnellii, Mormoops_blainvillei, Carollia_perspicillata, Artibeus_jamaicensis, Anoura_caudifer, Tonatia_saurophila, Micronycteris_hirsuta, Desmodus_rotundus, Pipistrellus_pipistrellus, Eptesicus_fuscus, Lasiurus_borealis, Murina_feae, Myotis_myotis, Myotis_davidii, Myotis_lucifugus, Myotis_brandtii, Miniopterus_natalensis, Miniopterus_schreibersii, Tadarida_brasiliensis, Ceratotherium_simum_cottoni, Ceratotherium_simum, Diceros_bicornis, Dicerorhinus_sumatrensis, Tapirus_terrestris, Tapirus_indicus, Equus_asinus, Equus_przewalskii, Equus_caballus, Manis_pentadactyla, Manis_javanica, Ursus_maritimus, Ailuropoda_melanoleuca, Neomonachus_schauinslandi, Mirounga_angustirostris, Leptonychotes_weddellii, Odobenus_rosmarus, Zalophus_californianus, Spilogale_gracilis, Ailurus_fulgens, Mellivora_capensis, Mustela_putorius, Enhydra_lutris, Pteronura_brasiliensis, Vulpes_lagopus, Canis_lupus_familiaris, Canis_lupus, Lycaon_pictus, Acinonyx_jubatus, Puma_concolor, Felis_nigripes, Felis_catus, Panthera_onca, Panthera_pardus, Panthera_tigris, Paradoxurus_hermaphroditus, Hyaena_hyaena, Helogale_parvula, Suricata_suricatta, Mungos_mungo, Cryptoprocta_ferox | Megaderma_lyra, Craseonycteris_thonglongyai, Hipposideros_armiger, Hipposideros_galeritus, Rhinolophus_sinicus, Macroglossus_sobrinus, Eidolon_helvum, Pteropus_vampyrus, Pteropus_alecto, Rousettus_aegyptiacus, Noctilio_leporinus, Pteronotus_parnellii, Mormoops_blainvillei, Carollia_perspicillata, Artibeus_jamaicensis, Anoura_caudifer, Tonatia_saurophila, Micronycteris_hirsuta, Desmodus_rotundus, Pipistrellus_pipistrellus, Eptesicus_fuscus, Lasiurus_borealis, Murina_feae, Myotis_myotis, Myotis_davidii, Myotis_lucifugus, Myotis_brandtii, Miniopterus_natalensis, Miniopterus_schreibersii, Tadarida_brasiliensis, Ceratotherium_simum_cottoni, Ceratotherium_simum, Diceros_bicornis, Dicerorhinus_sumatrensis, Tapirus_terrestris, Tapirus_indicus, Equus_asinus, Equus_przewalskii, Equus_caballus, Manis_pentadactyla, Manis_javanica, Ursus_maritimus, Ailuropoda_melanoleuca, Neomonachus_schauinslandi, Mirounga_angustirostris, Leptonychotes_weddellii, Odobenus_rosmarus, Zalophus_californianus, Spilogale_gracilis, Ailurus_fulgens, Mellivora_capensis, Mustela_putorius, Enhydra_lutris, Pteronura_brasiliensis, Vulpes_lagopus, Canis_lupus_familiaris, Canis_lupus, Lycaon_pictus, Acinonyx_jubatus, Puma_concolor, Felis_nigripes, Felis_catus, Panthera_onca, Panthera_pardus, Panthera_tigris, Paradoxurus_hermaphroditus, Hyaena_hyaena, Helogale_parvula, Suricata_suricatta, Mungos_mungo, Cryptoprocta_ferox | Bos_taurus, Erinaceus_europaeus, Homo_sapiens, Mus_musculus

>Chiroptera_Cetartiodactyla_Ferae
Megaderma_lyra, Craseonycteris_thonglongyai, Hipposideros_armiger, Hipposideros_galeritus, Rhinolophus_sinicus, Macroglossus_sobrinus, Eidolon_helvum, Pteropus_vampyrus, Pteropus_alecto, Rousettus_aegyptiacus, Noctilio_leporinus, Pteronotus_parnellii, Mormoops_blainvillei, Carollia_perspicillata, Artibeus_jamaicensis, Anoura_caudifer, Tonatia_saurophila, Micronycteris_hirsuta, Desmodus_rotundus, Pipistrellus_pipistrellus, Eptesicus_fuscus, Lasiurus_borealis, Murina_feae, Myotis_myotis, Myotis_davidii, Myotis_lucifugus, Myotis_brandtii, Miniopterus_natalensis, Miniopterus_schreibersii, Tadarida_brasiliensis, Sus_scrofa, Catagonus_wagneri, Elaphurus_davidianus, Rangifer_tarandus, Odocoileus_virginianus, Antilocapra_americana, Giraffa_tippelskirchi, Okapia_johnstoni, Saiga_tatarica, Pantholops_hodgsonii, Ovis_canadensis, Ovis_aries, Capra_aegagrus, Capra_hircus, Hemitragus_hylocrius, Ammotragus_lervia, Beatragus_hunteri, Bison_bison, Bos_mutus, Bos_indicus, Bos_taurus, Bubalus_bubalis, Moschus_moschiferus, Tragulus_javanicus, Orcinus_orca, Tursiops_truncatus, Monodon_monoceros, Delphinapterus_leucas, Phocoena_phocoena, Neophocaena_asiaeorientalis, Lipotes_vexillifer, Inia_geoffrensis, Ziphius_cavirostris, Mesoplodon_bidens, Platanista_gangetica, Kogia_breviceps, Eschrichtius_robustus, Balaenoptera_acutorostrata, Balaenoptera_bonaerensis, Eubalaena_japonica, Hippopotamus_amphibius, Camelus_bactrianus, Camelus_ferus, Camelus_dromedarius, Vicugna_pacos, Manis_pentadactyla, Manis_javanica, Ursus_maritimus, Ailuropoda_melanoleuca, Neomonachus_schauinslandi, Mirounga_angustirostris, Leptonychotes_weddellii, Odobenus_rosmarus, Zalophus_californianus, Spilogale_gracilis, Ailurus_fulgens, Mellivora_capensis, Mustela_putorius, Enhydra_lutris, Pteronura_brasiliensis, Vulpes_lagopus, Canis_lupus_familiaris, Canis_lupus, Lycaon_pictus, Acinonyx_jubatus, Puma_concolor, Felis_nigripes, Felis_catus, Panthera_onca, Panthera_pardus, Panthera_tigris, Paradoxurus_hermaphroditus, Hyaena_hyaena, Helogale_parvula, Suricata_suricatta, Mungos_mungo, Cryptoprocta_ferox | Megaderma_lyra, Craseonycteris_thonglongyai, Hipposideros_armiger, Hipposideros_galeritus, Rhinolophus_sinicus, Macroglossus_sobrinus, Eidolon_helvum, Pteropus_vampyrus, Pteropus_alecto, Rousettus_aegyptiacus, Noctilio_leporinus, Pteronotus_parnellii, Mormoops_blainvillei, Carollia_perspicillata, Artibeus_jamaicensis, Anoura_caudifer, Tonatia_saurophila, Micronycteris_hirsuta, Desmodus_rotundus, Pipistrellus_pipistrellus, Eptesicus_fuscus, Lasiurus_borealis, Murina_feae, Myotis_myotis, Myotis_davidii, Myotis_lucifugus, Myotis_brandtii, Miniopterus_natalensis, Miniopterus_schreibersii, Tadarida_brasiliensis, Sus_scrofa, Catagonus_wagneri, Elaphurus_davidianus, Rangifer_tarandus, Odocoileus_virginianus, Antilocapra_americana, Giraffa_tippelskirchi, Okapia_johnstoni, Saiga_tatarica, Pantholops_hodgsonii, Ovis_canadensis, Ovis_aries, Capra_aegagrus, Capra_hircus, Hemitragus_hylocrius, Ammotragus_lervia, Beatragus_hunteri, Bison_bison, Bos_mutus, Bos_indicus, Bos_taurus, Bubalus_bubalis, Moschus_moschiferus, Tragulus_javanicus, Orcinus_orca, Tursiops_truncatus, Monodon_monoceros, Delphinapterus_leucas, Phocoena_phocoena, Neophocaena_asiaeorientalis, Lipotes_vexillifer, Inia_geoffrensis, Ziphius_cavirostris, Mesoplodon_bidens, Platanista_gangetica, Kogia_breviceps, Eschrichtius_robustus, Balaenoptera_acutorostrata, Balaenoptera_bonaerensis, Eubalaena_japonica, Hippopotamus_amphibius, Camelus_bactrianus, Camelus_ferus, Camelus_dromedarius, Vicugna_pacos, Manis_pentadactyla, Manis_javanica, Ursus_maritimus, Ailuropoda_melanoleuca, Neomonachus_schauinslandi, Mirounga_angustirostris, Leptonychotes_weddellii, Odobenus_rosmarus, Zalophus_californianus, Spilogale_gracilis, Ailurus_fulgens, Mellivora_capensis, Mustela_putorius, Enhydra_lutris, Pteronura_brasiliensis, Vulpes_lagopus, Canis_lupus_familiaris, Canis_lupus, Lycaon_pictus, Acinonyx_jubatus, Puma_concolor, Felis_nigripes, Felis_catus, Panthera_onca, Panthera_pardus, Panthera_tigris, Paradoxurus_hermaphroditus, Hyaena_hyaena, Helogale_parvula, Suricata_suricatta, Mungos_mungo, Cryptoprocta_ferox | Equus_caballus, Erinaceus_europaeus, Homo_sapiens, Mus_musculus

>Euarchonta
Homo_sapiens, Pan_paniscus, Pan_troglodytes, Gorilla_gorilla, Pongo_abelii, Nomascus_leucogenys, Colobus_angolensis, Piliocolobus_tephrosceles, Semnopithecus_entellus, Nasalis_larvatus, Pygathrix_nemaeus, Rhinopithecus_bieti, Rhinopithecus_roxellana, Cercopithecus_neglectus, Chlorocebus_sabaeus, Erythrocebus_patas, Macaca_nemestrina, Macaca_fascicularis, Macaca_mulatta, Papio_anubis, Cercocebus_atys, Mandrillus_leucophaeus, Ateles_geoffroyi, Alouatta_palliata, Cebus_capucinus, Cebus_albifrons, Saimiri_boliviensis, Aotus_nancymaae, Callithrix_jacchus, Saguinus_imperator, Callicebus_donacophilus, Pithecia_pithecia, Eulemur_flavifrons, Eulemur_fulvus, Lemur_catta, Mirza_coquereli, Microcebus_murinus, Cheirogaleus_medius, Indri_indri, Propithecus_coquereli, Daubentonia_madagascariensis, Otolemur_garnettii, Nycticebus_coucang, Galeopterus_variegatus, Tupaia_tana, Tupaia_chinensis | Homo_sapiens, Pan_paniscus, Pan_troglodytes, Gorilla_gorilla, Pongo_abelii, Nomascus_leucogenys, Colobus_angolensis, Piliocolobus_tephrosceles, Semnopithecus_entellus, Nasalis_larvatus, Pygathrix_nemaeus, Rhinopithecus_bieti, Rhinopithecus_roxellana, Cercopithecus_neglectus, Chlorocebus_sabaeus, Erythrocebus_patas, Macaca_nemestrina, Macaca_fascicularis, Macaca_mulatta, Papio_anubis, Cercocebus_atys, Mandrillus_leucophaeus, Ateles_geoffroyi, Alouatta_palliata, Cebus_capucinus, Cebus_albifrons, Saimiri_boliviensis, Aotus_nancymaae, Callithrix_jacchus, Saguinus_imperator, Callicebus_donacophilus, Pithecia_pithecia, Eulemur_flavifrons, Eulemur_fulvus, Lemur_catta, Mirza_coquereli, Microcebus_murinus, Cheirogaleus_medius, Indri_indri, Propithecus_coquereli, Daubentonia_madagascariensis, Otolemur_garnettii, Nycticebus_coucang, Galeopterus_variegatus, Tupaia_tana, Tupaia_chinensis | Mus_musculus, Rattus_norvegicus, Oryctolagus_cuniculus, Canis_lupus_familiaris, Felis_catus, Bos_taurus

>Glires_Scandentia
Ochotona_princeps, Oryctolagus_cuniculus, Lepus_americanus, Ctenodactylus_gundi, Petromus_typicus, Thryonomys_swinderianus, Heterocephalus_glaber, Fukomys_damarensis, Dolichotis_patagonum, Hydrochoerus_hydrochaeris, Cavia_tschudii, Cavia_porcellus, Cavia_aperea, Dasyprocta_punctata, Cuniculus_paca, Octodon_degus, Ctenomys_sociabilis, Myocastor_coypus, Capromys_pilorides, Chinchilla_lanigera, Dinomys_branickii, Hystrix_cristata, Rattus_norvegicus, Mus_musculus, Mus_spretus, Mus_caroli, Mus_pahari, Acomys_cahirinus, Meriones_unguiculatus, Psammomys_obesus, Mesocricetus_auratus, Cricetulus_griseus, Microtus_ochrogaster, Ondatra_zibethicus, Ellobius_talpinus, Ellobius_lutescens, Sigmodon_hispidus, Peromyscus_maniculatus, Onychomys_torridus, Cricetomys_gambianus, Nannospalax_galili, Jaculus_jaculus, Allactaga_bullata, Zapus_hudsonius, Perognathus_longimembris, Dipodomys_stephensi, Dipodomys_ordii, Castor_canadensis, Xerus_inauris, Spermophilus_dauricus, Ictidomys_tridecemlineatus, Marmota_marmota, Aplodontia_rufa, Muscardinus_avellanarius, Glis_glis, Graphiurus_murinus, Tupaia_tana, Tupaia_chinensis | Ochotona_princeps, Oryctolagus_cuniculus, Lepus_americanus, Ctenodactylus_gundi, Petromus_typicus, Thryonomys_swinderianus, Heterocephalus_glaber, Fukomys_damarensis, Dolichotis_patagonum, Hydrochoerus_hydrochaeris, Cavia_tschudii, Cavia_porcellus, Cavia_aperea, Dasyprocta_punctata, Cuniculus_paca, Octodon_degus, Ctenomys_sociabilis, Myocastor_coypus, Capromys_pilorides, Chinchilla_lanigera, Dinomys_branickii, Hystrix_cristata, Rattus_norvegicus, Mus_musculus, Mus_spretus, Mus_caroli, Mus_pahari, Acomys_cahirinus, Meriones_unguiculatus, Psammomys_obesus, Mesocricetus_auratus, Cricetulus_griseus, Microtus_ochrogaster, Ondatra_zibethicus, Ellobius_talpinus, Ellobius_lutescens, Sigmodon_hispidus, Peromyscus_maniculatus, Onychomys_torridus, Cricetomys_gambianus, Nannospalax_galili, Jaculus_jaculus, Allactaga_bullata, Zapus_hudsonius, Perognathus_longimembris, Dipodomys_stephensi, Dipodomys_ordii, Castor_canadensis, Xerus_inauris, Spermophilus_dauricus, Ictidomys_tridecemlineatus,

Marmota_marmota, Aplodontia_rufa, Muscardinus_avellanarius, Glis_glis, Graphiurus_murinus, Tupaia_tana, Tupaia_chinensis | Homo_sapiens, Gorilla_gorilla, Canis_lupus_familiaris, Felis_catus, Bos_taurus, Loxodonta_africana

>GliresPrimatomorpha
Homo_sapiens, Pan_paniscus, Pan_troglodytes, Gorilla_gorilla, Pongo_abelii, Nomascus_leucogenys, Colobus_angolensis, Piliocolobus_tephrosceles, Semnopithecus_entellus, Nasalis_larvatus, Pygathrix_nemaeus, Rhinopithecus_bieti, Rhinopithecus_roxellana, Cercopithecus_neglectus, Chlorocebus_sabaeus, Erythrocebus_patas, Macaca_nemestrina, Macaca_fascicularis, Macaca_mulatta, Papio_anubis, Cercocebus_atys, Mandrillus_leucophaeus, Ateles_geoffroyi, Alouatta_palliata, Cebus_capucinus, Cebus_albifrons, Saimiri_boliviensis, Aotus_nancymaae, Callithrix_jacchus, Saguinus_imperator, Callicebus_donacophilus, Pithecia_pithecia, Eulemur_flavifrons, Eulemur_fulvus, Lemur_catta, Mirza_coquereli, Microcebus_murinus, Cheirogaleus_medius, Indri_indri, Propithecus_coquereli, Daubentonia_madagascariensis, Otolemur_garnettii, Nycticebus_coucang, Galeopterus_variegatus, Ochotona_princeps, Oryctolagus_cuniculus, Lepus_americanus, Ctenodactylus_gundi, Petromus_typicus, Thryonomys_swinderianus, Heterocephalus_glaber, Fukomys_damarensis, Dolichotis_patagonum, Hydrochoerus_hydrochaeris, Cavia_tschudii, Cavia_porcellus, Cavia_aperea, Dasyprocta_punctata, Cuniculus_paca, Octodon_degus, Ctenomys_sociabilis, Myocastor_coypus, Capromys_pilorides, Chinchilla_lanigera, Dinomys_branickii, Hystrix_cristata, Rattus_norvegicus, Mus_musculus, Mus_spretus, Mus_caroli, Mus_pahari, Acomys_cahirinus, Meriones_unguiculatus, Psammomys_obesus, Mesocricetus_auratus, Cricetulus_griseus, Microtus_ochrogaster, Ondatra_zibethicus, Ellobius_talpinus, Ellobius_lutescens, Sigmodon_hispidus, Peromyscus_maniculatus, Onychomys_torridus, Cricetomys_gambianus, Nannospalax_galili, Jaculus_jaculus, Allactaga_bullata, Zapus_hudsonius, Perognathus_longimembris, Dipodomys_stephensi, Dipodomys_ordii, Castor_canadensis, Xerus_inauris, Spermophilus_dauricus, Ictidomys_tridecemlineatus, Marmota_marmota, Aplodontia_rufa, Muscardinus_avellanarius, Glis_glis, Graphiurus_murinus | Homo_sapiens, Pan_paniscus, Pan_troglodytes, Gorilla_gorilla, Pongo_abelii, Nomascus_leucogenys, Colobus_angolensis, Piliocolobus_tephrosceles, Semnopithecus_entellus, Nasalis_larvatus, Pygathrix_nemaeus, Rhinopithecus_bieti, Rhinopithecus_roxellana, Cercopithecus_neglectus, Chlorocebus_sabaeus, Erythrocebus_patas, Macaca_nemestrina, Macaca_fascicularis, Macaca_mulatta, Papio_anubis, Cercocebus_atys, Mandrillus_leucophaeus, Ateles_geoffroyi, Alouatta_palliata, Cebus_capucinus, Cebus_albifrons, Saimiri_boliviensis, Aotus_nancymaae, Callithrix_jacchus, Saguinus_imperator, Callicebus_donacophilus, Pithecia_pithecia, Eulemur_flavifrons, Eulemur_fulvus, Lemur_catta, Mirza_coquereli, Microcebus_murinus, Cheirogaleus_medius, Indri_indri, Propithecus_coquereli, Daubentonia_madagascariensis, Otolemur_garnettii, Nycticebus_coucang, Galeopterus_variegatus, Ochotona_princeps, Oryctolagus_cuniculus, Lepus_americanus, Ctenodactylus_gundi, Petromus_typicus, Thryonomys_swinderianus, Heterocephalus_glaber, Fukomys_damarensis, Dolichotis_patagonum, Hydrochoerus_hydrochaeris, Cavia_tschudii, Cavia_porcellus, Cavia_aperea, Dasyprocta_punctata, Cuniculus_paca, Octodon_degus, Ctenomys_sociabilis, Myocastor_coypus, Capromys_pilorides, Chinchilla_lanigera, Dinomys_branickii, Hystrix_cristata, Rattus_norvegicus, Mus_musculus, Mus_spretus, Mus_caroli, Mus_pahari, Acomys_cahirinus, Meriones_unguiculatus, Psammomys_obesus, Mesocricetus_auratus, Cricetulus_griseus, Microtus_ochrogaster, Ondatra_zibethicus, Ellobius_talpinus, Ellobius_lutescens, Sigmodon_hispidus, Peromyscus_maniculatus, Onychomys_torridus, Cricetomys_gambianus, Nannospalax_galili, Jaculus_jaculus, Allactaga_bullata, Zapus_hudsonius, Perognathus_longimembris, Dipodomys_stephensi, Dipodomys_ordii, Castor_canadensis, Xerus_inauris, Spermophilus_dauricus, Ictidomys_tridecemlineatus, Marmota_marmota, Aplodontia_rufa, Muscardinus_avellanarius, Glis_glis, Graphiurus_murinus | Tupaia_chinensis, Tupaia_tana, Canis_lupus_familiaris, Felis_catus, Bos_taurus

### Data S2.

Timetree for the 241 species nearly-neutral dataset.

#NEXUS
BEGIN TREES;

UTREE 1 = (((((((((Dicerorhinus_sumatrensis: 0.137485, (Diceros_bicornis: 0.063043, (Ceratotherium_simum_cottoni: 0.005107, Ceratotherium_simum: 0.005107) [&95%HPD={0.0016225, 0.0096101}]: 0.057936) [&95%HPD={0.0170198, 0.205012}]: 0.074442) [&95%HPD={0.0503577, 0.262802}]: 0.420239, (Tapirus_terrestris: 0.077115, Tapirus_indicus: 0.077115) [&95%HPD={0.0244868, 0.157223}]: 0.480609) [&95%HPD={0.534167, 0.591706}]: 0.037790, (Equus_asinus: 0.038325, (Equus_przewalskii: 0.007432, Equus_caballus: 0.007432) [&95%HPD={0.0022457, 0.0139398}]: 0.030893) [&95%HPD={0.0143881, 0.0667745}]: 0.557189) [&95%HPD={0.566334, 0.618168}]: 0.232899, ((Manis_pentadactyla: 0.112908, Manis_javanica: 0.112908) [&95%HPD={0.0348647, 0.228649}]: 0.659790, ((((Ursus_maritimus: 0.105632, Ailuropoda_melanoleuca: 0.105632) [&95%HPD={0.0319043, 0.190797}]: 0.257165, (((Neomonachus_schauinslandi: 0.046555, (Mirounga_angustirostris: 0.037416, Leptonychotes_weddellii: 0.037416) [&95%HPD={0.0131316, 0.0658575}]: 0.009139) [&95%HPD={0.020402, 0.0754745}]: 0.194747, (Odobenus_rosmarus: 0.179560, Zalophus_californianus: 0.179560) [&95%HPD={0.156954, 0.217415}]: 0.061742) [&95%HPD={0.199561, 0.295971}]: 0.098170, (Spilogale_gracilis: 0.266550, (Ailurus_fulgens: 0.236400, (Mellivora_capensis: 0.126881, (Mustela_putorius: 0.087764, (Enhydra_lutris: 0.057115, Pteronura_brasiliensis: 0.057115) [&95%HPD={0.0301147, 0.0886302}]: 0.030649) [&95%HPD={0.0471448, 0.13241}]: 0.039117) [&95%HPD={0.0748725, 0.179675}]: 0.109519) [&95%HPD={0.175838, 0.295344}]: 0.030150) [&95%HPD={0.206467, 0.323914}]: 0.072922) [&95%HPD={0.284833, 0.390903}]: 0.023324) [&95%HPD={0.310071, 0.414275}]: 0.086318, (Vulpes_lagopus: 0.066789, (Lycaon_pictus: 0.021539, (Canis_lupus_familiaris: 0.005352, Canis_lupus: 0.005352) [&95%HPD={0.0017551, 0.0100162}]: 0.016187) [&95%HPD={0.0092837, 0.0370327}]: 0.045250) [&95%HPD={0.0284286, 0.115817}]: 0.382325) [&95%HPD={0.376238, 0.524865}]: 0.100799, ((((Acinonyx_jubatus: 0.031897, Puma_concolor: 0.031897) [&95%HPD={0.0134304, 0.0524504}]: 0.019787, (Felis_nigripes: 0.025878, Felis_catus: 0.025878) [&95%HPD={0.0100102, 0.0440883}]: 0.025806) [&95%HPD={0.0311255, 0.0745526}]: 0.020755, (Panthera_tigris: 0.026768, (Panthera_onca: 0.017864, Panthera_pardus: 0.017864) [&95%HPD={0.0069619, 0.0303392}]: 0.008904) [&95%HPD={0.0124635, 0.0417449}]: 0.045672) [&95%HPD={0.0452663, 0.102966}]: 0.281585, (Paradoxurus_hermaphroditus: 0.323778, (Hyaena_hyaena: 0.246347, (Cryptoprocta_ferox: 0.186192, (Suricata_suricatta: 0.081729, (Helogale_parvula: 0.062804, Mungos_mungo: 0.062804) [&95%HPD={0.0275206, 0.104491}]: 0.018925) [&95%HPD={0.0440712, 0.127455}]: 0.104463) [&95%HPD={0.131441, 0.239775}]: 0.060154) [&95%HPD={0.19842, 0.284647}]: 0.077431) [&95%HPD={0.250557, 0.394387}]: 0.030247) [&95%HPD={0.268674, 0.428748}]: 0.195889) [&95%HPD={0.461276, 0.640111}]: 0.222784) [&95%HPD={0.697713, 0.845177}]: 0.055714) [&95%HPD={0.735971, 0.927451}]: 0.020122, (((Sus_scrofa: 0.277169, Catagonus_wagneri: 0.277169) [&95%HPD={0.171957, 0.379169}]: 0.340457, ((Tragulus_javanicus: 0.407718, (((Elaphurus_davidianus: 0.111165, (Rangifer_tarandus: 0.047322, Odocoileus_virginianus: 0.047322) [&95%HPD={0.0191801, 0.0778132}]: 0.063844) [&95%HPD={0.0629282, 0.169056}]: 0.118358, (Moschus_moschiferus: 0.205839, ((Saiga_tatarica: 0.132865, (Beatragus_hunteri: 0.106178, (Pantholops_hodgsonii: 0.078481, ((Hemitragus_hylocrius: 0.029674, (Ovis_canadensis: 0.019879, Ovis_aries: 0.019879) [&95%HPD={0.0088948, 0.0321918}]: 0.009795) [&95%HPD={0.0169778, 0.0441087}]: 0.026875, (Ammotragus_lervia: 0.044189, (Capra_aegagrus: 0.008084, Capra_hircus: 0.008084) [&95%HPD={0.0025699, 0.0148974}]: 0.036105) [&95%HPD={0.0259226, 0.0668736}]: 0.012360) [&95%HPD={0.0379542, 0.0784571}]: 0.021932) [&95%HPD={0.05473, 0.10469}]: 0.027698) [&95%HPD={0.0753918, 0.140981}]: 0.026686) [&95%HPD={0.0975447, 0.170676}]: 0.039573, (Bubalus_bubalis: 0.089537, ((Bison_bison: 0.019869, Bos_mutus: 0.019869) [&95%HPD={0.0080816, 0.0330696}]: 0.008605, (Bos_indicus: 0.010641, Bos_taurus: 0.010641) [&95%HPD={0.0035768, 0.0194401}]: 0.017833) [&95%HPD={0.0145812, 0.0440361}]: 0.061063) [&95%HPD={0.0465314, 0.145445}]: 0.082901) [&95%HPD={0.138647, 0.220339}]: 0.033401) [&95%HPD={0.176786, 0.252011}]: 0.023685) [&95%HPD={0.19356, 0.274817}]: 0.017945, (Antilocapra_americana: 0.229614, (Giraffa_tippelskirchi: 0.099270, Okapia_johnstoni: 0.099270) [&95%HPD={0.0346767, 0.176201}]: 0.130344) [&95%HPD={0.189147, 0.279662}]: 0.017855) [&95%HPD={0.204729, 0.29255}]: 0.160250) [&95%HPD={0.319002, 0.499097}]: 0.161884, (Hippopotamus_amphibius: 0.538132, ((Kogia_breviceps: 0.274888, (Platanista_gangetica: 0.262674, ((((Orcinus_orca: 0.048634, Tursiops_truncatus: 0.048634) [&95%HPD={0.0157244, 0.0866757}]: 0.077294, ((Monodon_monoceros: 0.024855, Delphinapterus_leucas: 0.024855) [&95%HPD={0.0069564, 0.0486067}]: 0.073990, (Phocoena_phocoena: 0.021434, Neophocaena_asiaeorientalis: 0.021434) [&95%HPD={0.006006, 0.0412412}]: 0.077411) [&95%HPD={0.0808085, 0.123489}]: 0.027082) [&95%HPD={0.094779, 0.16426}]: 0.045778, (Lipotes_vexillifer: 0.159497, Inia_geoffrensis: 0.159497) [&95%HPD={0.111466, 0.211634}]: 0.012208) [&95%HPD={0.126484, 0.220312}]: 0.076398, (Ziphius_cavirostris: 0.063261, Mesoplodon_bidens: 0.063261) [&95%HPD={0.0143248, 0.107555}]: 0.184842) [&95%HPD={0.197619, 0.317077}]: 0.014570) [&95%HPD={0.215415, 0.326395}]: 0.012215) [&95%HPD={0.22741, 0.333172}]: 0.095919, (Eubalaena_japonica: 0.189286, (Eschrichtius_robustus: 0.091937, (Balaenoptera_acutorostrata: 0.027956, Balaenoptera_bonaerensis: 0.027956) [&95%HPD={0.0088727, 0.0525905}]: 0.063982) [&95%HPD={0.0689268, 0.126244}]: 0.097349) [&95%HPD={0.139092, 0.28986}]: 0.181521) [&95%HPD={0.351576, 0.399448}]: 0.167324) [&95%HPD={0.518599, 0.567336}]: 0.031470) [&95%HPD={0.540433, 0.604126}]: 0.048025) [&95%HPD={0.583272, 0.648798}]: 0.031999, (Vicugna_pacos: 0.088892, (Camelus_dromedarius: 0.016271, (Camelus_bactrianus: 0.006185, Camelus_ferus: 0.006185) [&95%HPD={0.0019926, 0.0113381}]: 0.010086) [&95%HPD={0.0069837, 0.027573}]: 0.072622) [&95%HPD={0.0381445, 0.160738}]: 0.560733) [&95%HPD={0.619591, 0.67072}]: 0.198909) [&95%HPD={0.757415, 0.954589}]: 0.018277, ((((Megaderma_lyra: 0.415836, Craseonycteris_thonglongyai: 0.415836) [&95%HPD={0.345069, 0.478988}]: 0.102337, (Rhinolophus_sinicus: 0.409642, (Hipposideros_armiger: 0.156467, Hipposideros_galeritus: 0.156467) [&95%HPD={0.0600024, 0.245601}]: 0.253176) [&95%HPD={0.36854, 0.465255}]: 0.108531) [&95%HPD={0.454508, 0.576866}]: 0.065036, ((Macroglossus_sobrinus: 0.228220, (Pteropus_vampyrus: 0.046726, Pteropus_alecto: 0.046726) [&95%HPD={0.0165365, 0.0861247}]: 0.181494) [&95%HPD={0.145554, 0.307265}]: 0.009007, (Eidolon_helvum: 0.224454, Rousettus_aegyptiacus: 0.224454) [&95%HPD={0.142689, 0.305036}]: 0.012773) [&95%HPD={0.154933, 0.315271}]: 0.345982) [&95%HPD={0.52883, 0.642416}]: 0.055798, ((Noctilio_leporinus: 0.423726, ((Pteronotus_parnellii: 0.317426, Mormoops_blainvillei: 0.317426) [&95%HPD={0.239441, 0.386857}]: 0.050378, (Micronycteris_hirsuta: 0.284516, (Desmodus_rotundus: 0.266329, (Tonatia_saurophila: 0.215905, (Anoura_caudifer: 0.201735, (Carollia_perspicillata: 0.159451, Artibeus_jamaicensis: 0.159451) [&95%HPD={0.0922757, 0.23242}]: 0.042284) [&95%HPD={0.143326, 0.272853}]: 0.014171) [&95%HPD={0.154566, 0.282242}]: 0.050424) [&95%HPD={0.200892, 0.326998}]: 0.018187) [&95%HPD={0.217996, 0.349444}]: 0.083288) [&95%HPD={0.302667, 0.41664}]: 0.055922) [&95%HPD={0.353478, 0.498736}]: 0.117131, (Tadarida_brasiliensis: 0.460505, (((Lasiurus_borealis: 0.177522, (Pipistrellus_pipistrellus: 0.141818, Eptesicus_fuscus: 0.141818) [&95%HPD={0.0668883, 0.205783}]: 0.035704) [&95%HPD={0.107109, 0.248686}]: 0.070779, (Murina_feae: 0.182702, ((Myotis_myotis: 0.067753, Myotis_davidii: 0.067753)

[&95%HPD={0.0287573, 0.109376}]: 0.034662, (Myotis_lucifugus: 0.066590, Myotis_brandtii: 0.066590) [&95%HPD={0.0240335, 0.119677}]: 0.035825) [&95%HPD={0.0597132, 0.158011}]: 0.080287) [&95%HPD={0.123412, 0.243766}]: 0.065599) [&95%HPD={0.190366, 0.314825}]: 0.164451, (Miniopterus_natalensis: 0.032545, Miniopterus_schreibersii: 0.032545) [&95%HPD={0.0095001, 0.0618799}]: 0.380207) [&95%HPD={0.340887, 0.493685}]: 0.047753) [&95%HPD={0.389534, 0.524246}]: 0.080352) [&95%HPD={0.479309, 0.596402}]: 0.098152) [&95%HPD={0.590681, 0.673816}]: 0.227804) [&95%HPD={0.782292, 0.974046}]: 0.045966, ((Solenodon_paradoxus: 0.764841, (Erinaceus_europaeus: 0.679174, (Crocidura_indochinensis: 0.352859, Sorex_araneus: 0.352859) [&95%HPD={0.215047, 0.552378}]: 0.326316) [&95%HPD={0.604833, 0.788714}]: 0.085667) [&95%HPD={0.654586, 0.861713}]: 0.061751, (Uropsilus_gracilis: 0.488213, (Condylura_cristata: 0.296730, Scalopus_aquaticus: 0.296730) [&95%HPD={0.161184, 0.475494}]: 0.191483) [&95%HPD={0.353209, 0.645227}]: 0.338379) [&95%HPD={0.732018, 0.930188}]: 0.086185) [&95%HPD={0.8236, 1.00747}]: 0.079856, (((Ochotona_princeps: 0.571101, (Oryctolagus_cuniculus: 0.148927, Lepus_americanus: 0.148927) [&95%HPD={0.0487452, 0.278761}]: 0.422174) [&95%HPD={0.53569, 0.613394}]: 0.206790, (((Ctenodactylus_gundi: 0.552579, (Hystrix_cristata: 0.488232, (((Petromus_typicus: 0.217728, Thryonomys_swinderianus: 0.217728) [&95%HPD={0.129145, 0.308157}]: 0.154157, (Heterocephalus_glaber: 0.256455, Fukomys_damarensis: 0.256455) [&95%HPD={0.156294, 0.350722}]: 0.115430) [&95%HPD={0.301089, 0.431633}]: 0.082866, ((Cuniculus_paca: 0.302945, (Dasyprocta_punctata: 0.275222, ((Dolichotis_patagonum: 0.155809, Hydrochoerus_hydrochaeris: 0.155809) [&95%HPD={0.087489, 0.221641}]: 0.030066, (Cavia_aperea: 0.036067, (Cavia_tschudii: 0.004670, Cavia_porcellus: 0.004670) [&95%HPD={0.0014955, 0.0087264}]: 0.031397) [&95%HPD={0.0140088, 0.0629699}]: 0.149808) [&95%HPD={0.124107, 0.260454}]: 0.089347) [&95%HPD={0.206339, 0.362583}]: 0.027723) [&95%HPD={0.231483, 0.388308}]: 0.101796, (((Octodon_degus: 0.186975, Ctenomys_sociabilis: 0.186975) [&95%HPD={0.110503, 0.263967}]: 0.073738, (Myocastor_coypus: 0.169569, Capromys_pilorides: 0.169569) [&95%HPD={0.0980171, 0.24356}]: 0.091145) [&95%HPD={0.185265, 0.322497}]: 0.108520, (Chinchilla_lanigera: 0.277349, Dinomys_branickii: 0.277349) [&95%HPD={0.239621, 0.325227}]: 0.091884) [&95%HPD={0.315346, 0.420214}]: 0.035507) [&95%HPD={0.356555, 0.455203}]: 0.050010) [&95%HPD={0.404345, 0.505155}]: 0.033480) [&95%HPD={0.437998, 0.534471}]: 0.064347) [&95%HPD={0.519426, 0.584748}]: 0.088357, ((Aplodontia_rufa: 0.477924, (Xerus_inauris: 0.294307, (Marmota_marmota: 0.100212, (Spermophilus_dauricus: 0.056556, Ictidomys_tridecemlineatus: 0.056556) [&95%HPD={0.0211016, 0.0970406}]: 0.043655) [&95%HPD={0.0480706, 0.162392}]: 0.194095) [&95%HPD={0.19191, 0.408255}]: 0.183617) [&95%HPD={0.449262, 0.520073}]: 0.100370, (Glis_glis: 0.302150, (Muscardinus_avellanarius: 0.259363, Graphiurus_murinus: 0.259363) [&95%HPD={0.129266, 0.393147}]: 0.042787) [&95%HPD={0.188433, 0.441606}]: 0.276145) [&95%HPD={0.538221, 0.618004}]: 0.062642) [&95%HPD={0.61952, 0.659693}]: 0.016400, (((Nannospalax_galili: 0.492409, (Cricetomys_gambianus: 0.407860, (((Rattus_norvegicus: 0.152015, (Mus_pahari: 0.102920, (Mus_caroli: 0.064457, (Mus_musculus: 0.033780, Mus_spretus: 0.033780) [&95%HPD={0.0168867, 0.0524149}]: 0.030677) [&95%HPD={0.0416219, 0.0877621}]: 0.038463) [&95%HPD={0.0756162, 0.130146}]: 0.049095) [&95%HPD={0.132532, 0.164719}]: 0.159559, (Acomys_cahirinus: 0.246099, (Meriones_unguiculatus: 0.083513, Psammomys_obesus: 0.083513) [&95%HPD={0.0382044, 0.143985}]: 0.162586) [&95%HPD={0.181726, 0.313501}]: 0.065475) [&95%HPD={0.253135, 0.373801}]: 0.064480, (((Mesocricetus_auratus: 0.176054, Cricetulus_griseus: 0.176054) [&95%HPD={0.110178, 0.235918}]: 0.107012, (Ondatra_zibethicus: 0.181098, (Microtus_ochrogaster: 0.138048, (Ellobius_talpinus: 0.077071, Ellobius_lutescens: 0.077071) [&95%HPD={0.0386936, 0.125536}]: 0.060976) [&95%HPD={0.0813685, 0.186094}]: 0.043051) [&95%HPD={0.128695, 0.23196}]: 0.101967) [&95%HPD={0.235057, 0.337646}]: 0.019108, (Sigmodon_hispidus: 0.275919, (Peromyscus_maniculatus: 0.139310, Onychomys_torridus: 0.139310) [&95%HPD={0.0657437, 0.206707}]: 0.136610) [&95%HPD={0.222009, 0.330472}]: 0.026254) [&95%HPD={0.255019, 0.35354}]: 0.073880) [&95%HPD={0.33406, 0.42714}]: 0.031807) [&95%HPD={0.353946, 0.452722}]: 0.084549) [&95%HPD={0.446505, 0.547138}]: 0.067986, (Zapus_hudsonius: 0.366915, (Jaculus_jaculus: 0.209035, Allactaga_bullata: 0.209035) [&95%HPD={0.121852, 0.308246}]: 0.157880) [&95%HPD={0.258578, 0.461217}]: 0.193480) [&95%HPD={0.522926, 0.593098}]: 0.062811, (Castor_canadensis: 0.553033, (Perognathus_longimembris: 0.273854, (Dipodomys_stephensi: 0.058608, Dipodomys_ordii: 0.058608) [&95%HPD={0.0199858, 0.102104}]: 0.215246) [&95%HPD={0.170611, 0.383789}]: 0.279179) [&95%HPD={0.520246, 0.591473}]: 0.070173) [&95%HPD={0.59778, 0.646784}]: 0.034130) [&95%HPD={0.640843, 0.670503}]: 0.120554) [&95%HPD={0.718488, 0.858531}]: 0.042744, ((Tupaia_tana: 0.138827, Tupaia_chinensis: 0.138827) [&95%HPD={0.044148, 0.254641}]: 0.655431, (Galeopterus_variegatus: 0.741711, (((Daubentonia_madagascariensis: 0.436690, ((Lemur_catta: 0.147776, (Eulemur_flavifrons: 0.039845, Eulemur_fulvus: 0.039845) [&95%HPD={0.0142349, 0.0701445}]: 0.107930) [&95%HPD={0.076192, 0.213544}]: 0.124092, ((Cheirogaleus_medius: 0.177196, (Mirza_coquereli: 0.090532, Microcebus_murinus: 0.090532) [&95%HPD={0.0433141, 0.147726}]: 0.086664) [&95%HPD={0.108031, 0.236143}]: 0.077238, (Indri_indri: 0.131744, Propithecus_coquereli: 0.131744) [&95%HPD={0.0526057, 0.21009}]: 0.122690) [&95%HPD={0.192084, 0.321007}]: 0.017433) [&95%HPD={0.209656, 0.336816}]: 0.164822) [&95%HPD={0.351115, 0.523059}]: 0.070764, (Otolemur_garnettii: 0.392403, Nycticebus_coucang: 0.392403) [&95%HPD={0.364637, 0.430956}]: 0.115052) [&95%HPD={0.456047, 0.563004}]: 0.127371, ((((Ateles_geoffroyi: 0.157284, Alouatta_palliata: 0.157284) [&95%HPD={0.124969, 0.195464}]: 0.062562, ((Saimiri_boliviensis: 0.149150, (Cebus_capucinus: 0.008849, Cebus_albifrons: 0.008849) [&95%HPD={0.0028078, 0.0168338}]: 0.140302) [&95%HPD={0.115328, 0.185321}]: 0.026597, (Aotus_nancymaae: 0.169859, (Callithrix_jacchus: 0.102938, Saguinus_imperator: 0.102938) [&95%HPD={0.0528615, 0.159955}]: 0.066921) [&95%HPD={0.130089, 0.207681}]: 0.005888) [&95%HPD={0.139505, 0.213218}]: 0.044099) [&95%HPD={0.199895, 0.251065}]: 0.013800, (Callicebus_donacophilus: 0.174794, Pithecia_pithecia: 0.174794) [&95%HPD={0.0914254, 0.246119}]: 0.058852) [&95%HPD={0.208177, 0.268873}]: 0.153419, ((((Colobus_angolensis: 0.068858, Piliocolobus_tephrosceles: 0.068858) [&95%HPD={0.0334444, 0.121473}]: 0.018131, (Semnopithecus_entellus: 0.047710, ((Nasalis_larvatus: 0.036239, Pygathrix_nemaeus: 0.036239) [&95%HPD={0.0187344, 0.0572159}]: 0.002061, (Rhinopithecus_bieti: 0.012374, Rhinopithecus_roxellana: 0.012374) [&95%HPD={0.0042344, 0.0219599}]: 0.025926) [&95%HPD={0.0214357, 0.0601263}]: 0.009410) [&95%HPD={0.0276847, 0.0750195}]: 0.039279) [&95%HPD={0.0502518, 0.139881}]: 0.052457, ((Cercopithecus_neglectus: 0.057448, (Chlorocebus_sabaeus: 0.030856, Erythrocebus_patas: 0.030856) [&95%HPD={0.0125028, 0.0513545}]: 0.026592) [&95%HPD={0.0312832, 0.0838579}]: 0.027911, ((Macaca_nemestrina: 0.018457, (Macaca_fascicularis: 0.012330, Macaca_mulatta: 0.012330) [&95%HPD={0.0043497, 0.0220011}]: 0.006127) [&95%HPD={0.0082731, 0.0297255}]: 0.041818, (Papio_anubis: 0.046489, (Cercocebus_atys: 0.037106, Mandrillus_leucophaeus: 0.037106) [&95%HPD={0.0205531, 0.0533649}]: 0.009383) [&95%HPD={0.0305484, 0.0615532}]: 0.013786) [&95%HPD={0.0495976, 0.0738775}]: 0.025083) [&95%HPD={0.0625877, 0.114575}]: 0.054088) [&95%HPD={0.118191, 0.177132}]: 0.139297, (Nomascus_leucogenys: 0.167518, (Pongo_abelii: 0.141137, (Gorilla_gorilla: 0.071361, (Homo_sapiens: 0.057363, (Pan_paniscus: 0.014456, Pan_troglodytes: 0.014456) [&95%HPD={0.0045808, 0.027197}]: 0.042907) [&95%HPD={0.0498094, 0.0702619}]: 0.013997) [&95%HPD={0.0550564, 0.0913667}]: 0.069776) [&95%HPD={0.0874336, 0.196573}]: 0.026381) [&95%HPD={0.119502, 0.226781}]: 0.111226) [&95%HPD={0.244026, 0.328322}]: 0.108321) [&95%HPD={0.303002, 0.47541}]: 0.247761) [&95%HPD={0.588676, 0.665917}]: 0.106885) [&95%HPD={0.705533, 0.765992}]: 0.052546) [&95%HPD={0.733913, 0.886424}]: 0.026377) [&95%HPD={0.758058, 0.908386}]: 0.171999) [&95%HPD={0.881223, 1.11848}]: 0.045413, ((((Chrysochloris_asiatica: 0.562956, (Microgale_talazaci: 0.262229, Echinops_telfairi: 0.262229) [&95%HPD={0.163346, 0.403478}]: 0.300726) [&95%HPD={0.42984, 0.67108}]:

0.100038, (Elephantulus_edwardii: 0.597024, Orycteropus_afer: 0.597024) [&95%HPD={0.474308, 0.756856}]: 0.065970) [&95%HPD={0.561788, 0.789553}]: 0.072909, (Trichechus_manatus: 0.630516, (Loxodonta_africana: 0.596146, (Procavia_capensis: 0.078852, Heterohyrax_brucei: 0.078852) [&95%HPD={0.0589913, 0.10883}]: 0.517294) [&95%HPD={0.5563, 0.651162}]: 0.034370) [&95%HPD={0.584861, 0.696665}]: 0.105387) [&95%HPD={0.632385, 0.849178}]: 0.207192, (((Choloepus_didactylus: 0.035677, Choloepus_hoffmanni: 0.035677) [&95%HPD={0.0113256, 0.0647346}]: 0.413990, (Myrmecophaga_tridactyla: 0.182068, Tamandua_tetradactyla: 0.182068) [&95%HPD={0.122488, 0.257764}]: 0.267599) [&95%HPD={0.314216, 0.58746}]: 0.127646, (Dasypus_novemcinctus: 0.325553, (Chaetophractus_vellerosus: 0.198117, Tolypeutes_matacus: 0.198117) [&95%HPD={0.0803575, 0.316768}]: 0.127436) [&95%HPD={0.212665, 0.476845}]: 0.251759) [&95%HPD={0.481105, 0.661533}]: 0.365782) [&95%HPD={0.777667, 1.11263}]: 0.094952) [&95%HPD={0.907009, 1.18455}];

END;
